# Comprehensive taxon sampling and vetted fossils help clarify the time tree of shorebirds (Aves, Charadriiformes)

**DOI:** 10.1101/2021.07.15.452585

**Authors:** David Černý, Rossy Natale

## Abstract

Shorebirds (Charadriiformes) are a globally distributed clade of modern birds and, due to their ecological and morphological disparity, a frequent subject of comparative studies. While molecular phylogenies have been instrumental to resolving the suprafamilial backbone of the charadriiform tree, several higher-level relationships, including the monophyly of plovers (Charadriidae) and the phylogenetic positions of several monotypic families, have remained unclear. The timescale of shorebird evolution also remains uncertain as a result of extensive disagreements among the published divergence dating studies, stemming largely from different choices of fossil calibrations. Here, we present the most comprehensive non-supertree phylogeny of shorebirds to date, based on a total-evidence dataset comprising 336 ingroup taxa (89% of all extant species), 24 loci (15 mitochondrial and 9 nuclear), and 69 morphological characters. Using this phylogeny, we clarify the charadriiform evolutionary timeline by conducting a node-dating analysis based on a subset of 8 loci tested to be clock-like and 16 carefully selected, updated, and vetted fossil calibrations. Our concatenated, species-tree, and total-evidence analyses consistently support plover monophyly and are generally congruent with the topologies of previous studies, suggesting that the higher-level relationships among shorebirds are largely settled. However, several localized conflicts highlight areas of persistent uncertainty within the gulls (Laridae), true auks (Alcinae), and sandpipers (Scolopacidae). At shallower levels, our phylogenies reveal instances of genus-level nonmonophyly that suggest changes to currently accepted taxonomies. Our node-dating analyses consistently support a mid-Paleocene origin for the Charadriiformes and an early diversification for most major subclades. However, age estimates for more recent divergences vary between different relaxed clock models, and we demonstrate that this variation can affect phylogeny-based macroevolutionary studies. Our findings demonstrate the impact of fossil calibration choice on the resulting divergence time estimates, and the sensitivity of diversification rate analyses to the modeling assumptions made in time tree inference.

## 1 Introduction

Shorebirds (Charadriiformes) are an ecologically diverse and globally distributed order of approximately 380 species of neoavian birds (Boyd, 2019; Clements et al., 2019). Given the variation in habitat use, foraging mode, and behavior present in the group, the Charadriiformes have been a frequent subject of comparative research. A number of studies have investigated the origins and macroevolution of ecological, behavioral, or phenotypic traits in specific charadriiform genera or families, shedding light on questions such as the evolution of beak morphology in the Charadrii and Scolopacidae (Barbosa and Moreno, 1999), locomotor ecologies in auks and relatives (Smith and Clarke, 2012), migration behaviors in the *Charadrius* plovers (Joseph et al., 1999), and plumage in gulls and terns (Crochet et al., 2000; Bridge et al., 2005; Dufour et al., 2020). Despite these efforts, comparative analyses of shorebird morphology, ecology, and behavior at the ordinal level have been hampered by the lack of a robust, comprehensive time-calibrated phylogeny for the clade as a whole.

A division of Charadriiformes into three suborders has become well-established since molecular data started to replace or supplement the osteological (Strauch, 1978; Björklund, 1994; Chu, 1995), syringeal (Brown and Ward, 1990), integumentary (Jehl, 1968; Nielsen, 1975; Dove, 2000), and behavioral (Moynihan, 1959) characters that were used (often together, e.g. Chu, 1998; Jehl, 1968) to generate the earliest hypotheses of charadriiform phylogenetic relationships. This three-suborder structure consists of the Charadrii (plovers, thick-knees, oystercatchers, avocets), Scolopaci (sandpipers, jacanas, snipes), and the Lari (gulls, terns, skimmers, coursers). This basic structure was first identified by the DNA-DNA hybridization work of Sibley and Ahlquist (1990). Their phylogeny helped resolve the position of several taxa whose charadriiform affinities had been controversial, such as the auks (Verheyen, 1958; Gysels and Rabaey, 1964) and the plains-wanderer (Wetmore, 1960; Cracraft, 1981). The inclusion of the auks within Lari suggested by Sibley and Ahlquist (1990) corresponded to early hypotheses suggesting their close relationship to the gulls (Storer, 1960; Kozlova, 1961), while a sister-group relationship between the plains-wanderer and seedsnipes within Scolopaci corroborated the earlier morphological study of Olson and Steadman (1981). Topology-wise, Sibley and Ahlquist (1990) found the Scolopaci as the sister group of (Charadrii + Lari). Later studies using sequence data altered this topology by placing the Charadrii as sister to a clade formed by the Scolopaci and Lari, and by recovering the enigmatic buttonquails (Turnicidae), previously considered to be either gruiforms (Wetmore, 1960; Cracraft, 1981; Rotthowe and Starck, 1998) or a separate early-branching neoavian order (Sibley and Ahlquist, 1990), as early-diverging members of the Lari (Paton et al., 2003; Cracraft et al., 2004; Fain and Houde, 2004; Paton and Baker, 2006; Baker et al., 2007; Fain and Houde, 2007; Hackett et al., 2008; Hu et al., 2017; Prum et al., 2015).

Despite this general consensus, there remain several outstanding questions regarding higher-level charadriiform phylogenetic relationships, some of which concern the position of several monotypic families. The ibisbill (*Ibidorhyncha struthersii*) has been recovered as the sister group to the Recurvirostridae in morphological (Chu, 1995; Livezey, 2010) and supertree (Thomas et al., 2004) studies, while molecular analyses have allied it with the Haematopodidae (Baker et al., 2007; Chen et al., 2018) or a (Haematopodidae + Recurvirostridae) clade (hereafter referred to as Haematopodoidea following Cracraft, 2013; Burleigh et al. 2015). The monotypic crab plover (*Dromas ardeola*) was not represented in the early molecular phylogenetic studies on charadriiforms (Paton et al., 2003; Paton and Baker, 2006; Baker et al., 2007; Fain and Houde, 2007), and larger phylogenomic studies that did include it failed to sample shorebirds densely enough to unambiguously resolve its position within the order (Hackett et al., 2008; Reddy et al., 2017). Recent molecular evidence indicates a sister-group relationship of *Dromas* to the coursers and pratincoles (Glareolidae), albeit with varying degrees of support (Pereira and Baker, 2010; Burleigh et al., 2015; De Pietri et al., 2020; see also the supertree of Kimball et al., 2019). Finally, the monophyly of lapwings and plovers (Charadriidae) has been contested, with multiple studies finding the gray and golden plovers (genus *Pluvialis*) to be more closely related to the Haematopodoidea than to the rest of the family (Baker et al., 2007; Fain and Houde, 2007; Burleigh et al., 2015; Chen et al., 2018), or to fall outside of the clade formed by the Haematopodoidea and other plovers (Ericson et al., 2003). This contradicts the traditional morphological hypothesis of charadriid monophyly (Chu, 1995; Livezey, 2010) as well as the nuclear sequence analysis of Baker et al. (2012), who attributed the earlier molecular results to stochastic gene tree estimation error stemming primarily from the use of fast-evolving mitochondrial loci. However, phylogenies based on complete mitogenomes have since recovered *Pluvialis* within the Charadriidae (Hu et al., 2017; Chen et al., 2018), whereas the taxonomically comprehensive analysis of Burleigh et al. (2015), in which the position of *Pluvialis* was informed by three nuclear loci in addition to mtDNA data, again supported plover paraphyly.

The timescale of charadriiform evolution also remains uncertain due to incongruence among studies employing different types of molecular data and different interpretations of the fossil record. Notably, there is an almost threefold difference between the oldest (95% credible interval: 68–107 Ma; Paton et al., 2003) and the youngest (95% credible interval: 35.1–60.3 Ma; Prum et al., 2015) estimates of the age of the charadriiform root (Figure 1), with the two values implying drastically different scenarios for the tempo and mode of shorebird diversification. Early molecular dating studies based on mitochondrial DNA or small samples of nuclear loci placed the origin of crown shorebirds deep in the Late Cretaceous (Paton et al., 2002, 2003; Pereira and Baker, 2006), and in some cases suggested that many of their interfamilial divergences predated the Cretaceous–Paleogene (K–Pg) boundary (Baker et al., 2007; Brown et al., 2007, 2008; Pereira and Baker, 2008). These divergence time estimates are incompatible with the lack of any Cretaceous fossils that could be reliably attributed not only to charadriiforms (Smith, 2015), but even to neoavians in general (Field et al., 2020), and with phylogenomic evidence for an explosive origin of neoavian orders that largely (Ericson et al., 2006; Jarvis et al., 2014; Kimball et al., 2019; Kuhl et al., 2020) or perhaps entirely (Claramunt and Cracraft, 2015; Cracraft et al., 2015; Prum et al., 2015) postdated the K–Pg boundary, reflecting a rapid radiation into the ecological niches emptied by the Cretaceous–Paleogene mass extinction (Ksepka and Phillips, 2015; Suh, 2016; Berv and Field, 2017).

**Figure 1.**
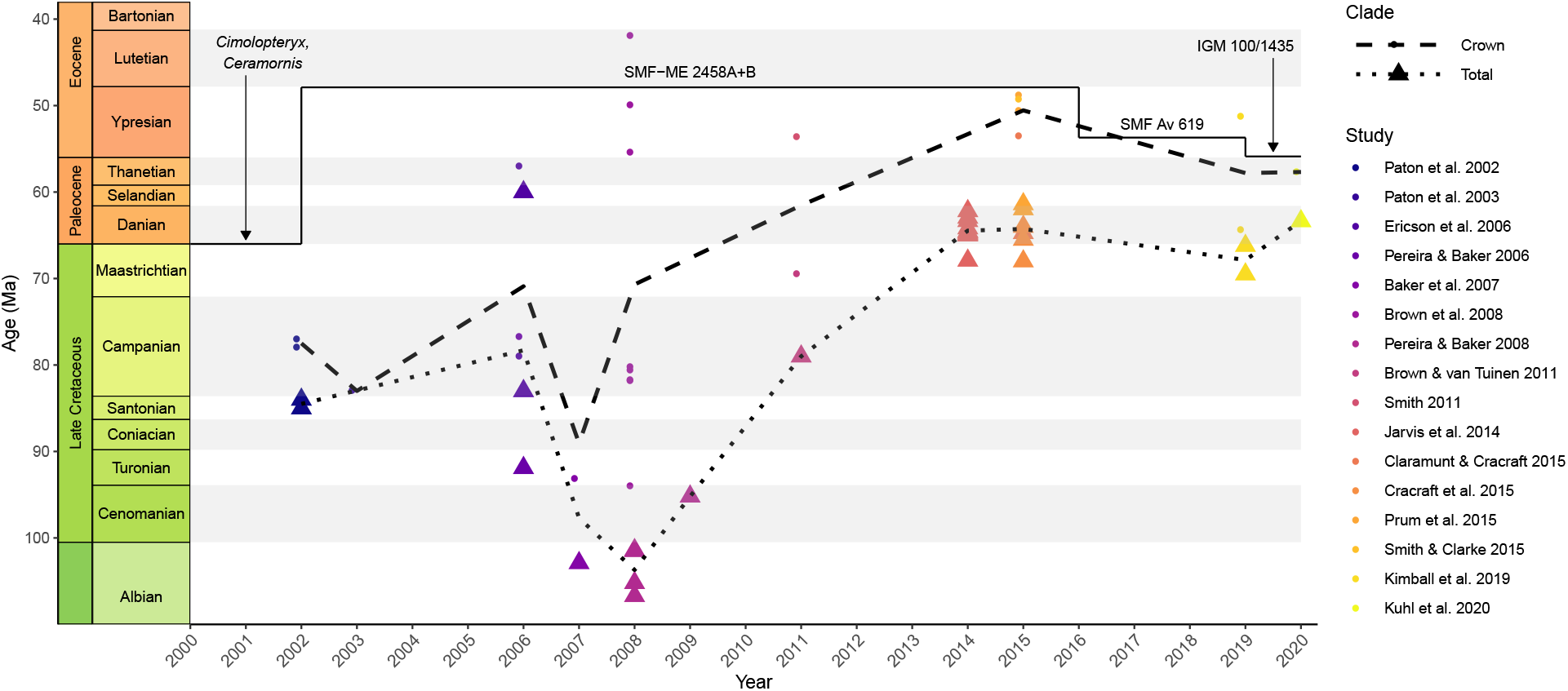
Previous age estimates for the Charadriiformes, plotted against the publication year and the oldest crown-charadriiform fossil known at the time (solid line). Mean estimates across different studies are shown separately for the total group (dotted line) and crown group (dashed line). After a long period during which the molecular divergence times drastically predated the known fossil record, the converse problem has started to occur with the advent of phylogenomic studies in the mid 2010s. See References for the full citations of the studies shown.

In part, the implausibly old divergence times inferred by early analyses can be attributed to the reliance on mitochondrial data (Brown and van Tuinen, 2011), which frequently overestimate node ages as a result of substitution saturation and small effective population sizes (Zheng et al., 2011; Smith and Klicka, 2013). However, inadequate calibration choices also play a role (Mayr, 2011; Smith, 2015). A number of early studies relied primarily on external calibrations phylogenetically distant from the clade of interest (Paton et al., 2002; Pereira and Baker, 2006), or re-used previous and excessively old divergence time estimates as secondary calibrations (Paton et al., 2003). The choice of calibration points within the Charadriiformes has been equally problematic, as the relevant taxa were drawn from obsolete fossil compilations without considering more recent re-assessments. Thus, Baker et al. (2007) calibrated the split between charadriiforms and their sister group with the ∼ 66 Ma old taxa *Ceramornis* and *Cimolopteryx*, which were regarded as charadriiforms by Brodkorb (1967) but subsequently re-evaluated as crown birds (Neornithes) of uncertain affinities by Hope (2002). Moreover, both fossils may be either latest Cretaceous or early Paleocene in age (Mayr, 2009), making them poorly constrained both phylogenetically and stratigraphically. Similar problems extended to nearly all calibrations used by Baker et al. (2007) and Pereira and Baker (2008) (see Mayr, 2011; Smith, 2011 for detailed criticisms), rendering the resulting divergence time estimates untrustworthy.

More recent node-dating analyses have generally inferred much younger divergence times for the Charadriiformes (Figure 1), estimating their origin to be younger than the K–Pg boundary (Jarvis et al., 2014; Kuhl et al., 2020) and often as young as Eocene in age (Claramunt and Cracraft, 2015; Prum et al., 2015; Kimball et al., 2019). While this shift to younger dates may have been aided by the transition to more slowly evolving and less saturated nuclear loci, as well as by the smaller branch length estimation error resulting from the use of larger quantities of sequence data (Yang and Rannala, 2005), the fact that a similar estimate was obtained by a study using a short mtDNA-dominated alignment along with carefully vetted calibrations (Smith and Clarke, 2015) points to calibration choice as a decisive factor. Indeed, a number of recent phylogenomic studies have cited and followed the “best practices” outlined by Parham et al. (2011), according to which calibrations should be assigned to nodes based on a list of apomorphies or the results of phylogenetic analysis, and explicit reasoning should be provided for the conversion of the available stratigraphic information into numeric ages. However, an overly conservative interpretation of these guidelines may have caused recent studies to over-correct and disregard pertinent fossil evidence, possibly resulting in the underestimation of divergence times (Figure 1). In the worst case, this bias may even give rise to “zombie lineages” (*sensu* Springer et al., 2017) whose estimated divergence time postdates their first appearance in the fossil record. For example, Jarvis et al. (2014) used the ∼32 Ma old *Boutersemia* to calibrate the divergence of the Charadriiformes from their sister group, despite the fact that an almost 50% older fossil had already been described by Mayr (2000) and assigned to the Charadriiformes based on apomorphies determined by outgroup comparison. Other fossils older than 32 Ma had been recovered as crown-group charadriiforms in a formal phylogenetic analysis by Smith (2011), demonstrating that even this more stringent criterion did not justify basing the calibration on *Boutersemia*.

Near-complete species-level phylogenies are increasingly available for many avian clades (e.g. Garcia-R et al. 2014; Marki et al. 2017; Olsson and Alstrom 2020), providing a robust basis for inferences ranging from diversification rate estimation to historical biogeography. In shorebirds, however, such phylogenies (summarized in Figure 2) remain subject to methodological shortcomings and limited data availability. With 227 species, the charadriiform matrix of Strauch (1978) still represents one of the largest morphological phylogenetic datasets ever constructed in terms of the number of taxa, but this early achievement has not been followed by subsequent morphological and molecular studies, whose taxon sampling has mostly remained either broad but sparse (Baker et al., 2007; Mayr, 2011) or dense but narrow (Pons et al., 2005; Smith and Clarke, 2015). As a result, attempts to construct a comprehensive shorebird phylogeny have so far relied on supertree techniques. The supertree of Thomas et al. (2004) succeeded at including all 350 then-recognized non-turnicid species, but aside from problems inherent to the method used (Gatesy and Springer, 2004; Bininda-Emonds, 2014), it also suffered from poor resolution and the reliance on obsolete source trees incompatible with the emerging consensus about shorebird phylogeny. Using a more advanced “backbone-and-patch” approach, Jetz et al. (2012) first inferred separate time trees for Charadrii, Scolopaci, Turnicidae, and the non-turnicid Lari from sequence data, and attached them to a phylogenomic backbone to produce a set of phylogenies comprising a total of 278 charadriiform species. These were then expanded to all 369 then-recognized species by using taxonomy to constrain the placement of those taxa for which no molecular data were available, and stochastically resolving the resulting polytomies. While accommodating uncertainty better than the approach of Thomas et al. (2004), this workflow, too, suffers from important drawbacks. The information about topology and divergence times present in the sequence data is not allowed to inform the backbone, and the placement of many taxa (*>* 25% of the extant charadriiform diversity) is not based on actual data and may reproduce the errors of previous taxonomies. Moreover, the use of birth-death polytomy resolvers may lead to unreliable downstream inferences (Rabosky, 2015; Weedop et al., 2019). The more recent phylogeny of Burleigh et al. (2015), based on a single molecular supermatrix, avoided these problems at the cost of reduced taxon sampling (272 charadriiform species; Figure 2).

**Figure 2.**
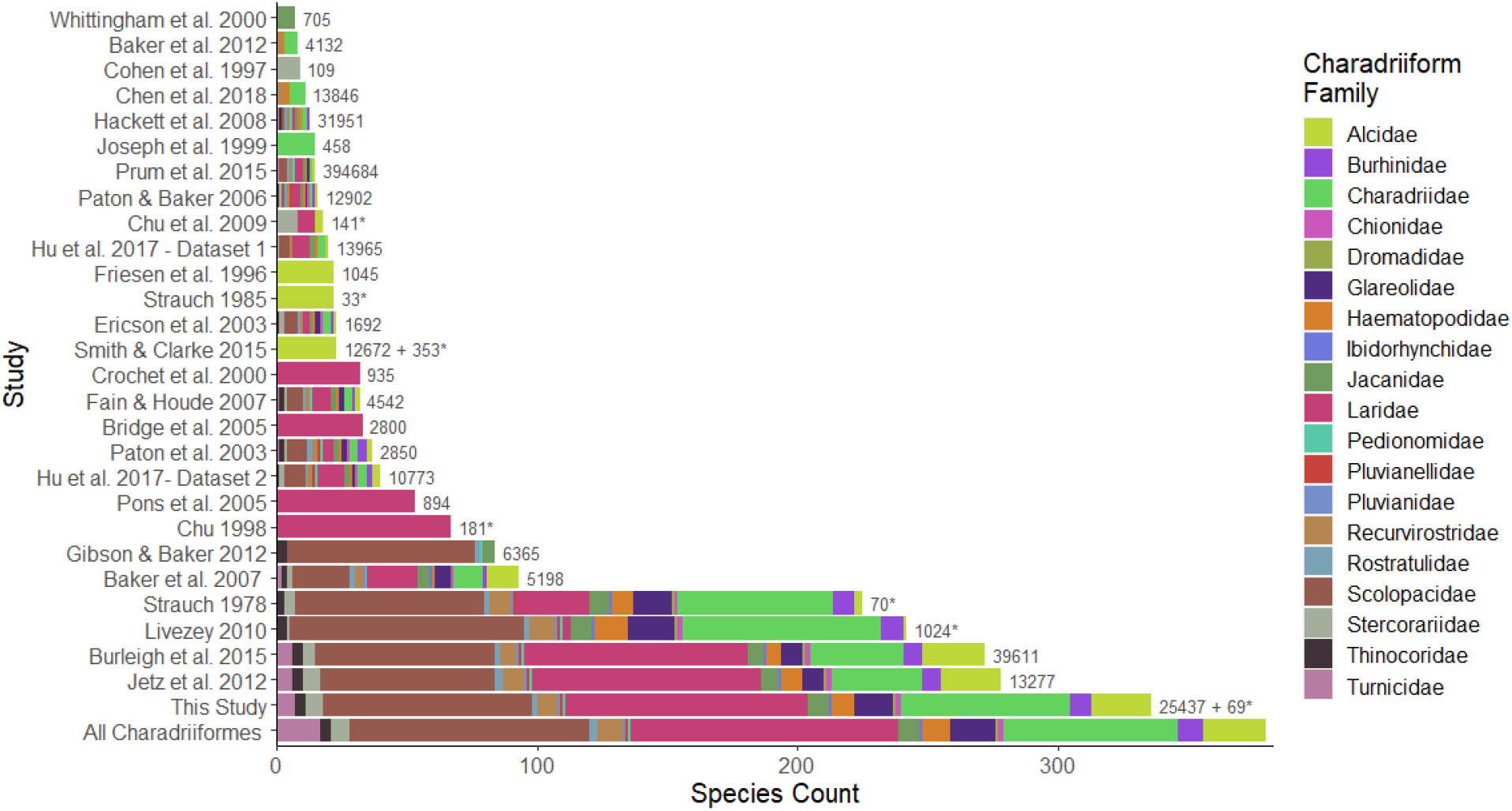
Taxonomic coverage of previous phylogenetic analyses of the Charadriiformes. Outgroups were not included in counts of family-level coverage. Counts of total recognized charadriiform families follow Boyd (2019). Bars are labeled with the number of characters used; asterisks denote morphological characters.

Here, we assemble the most comprehensive molecular dataset for the Charadriiformes to date, and combine it with a pre-existing morphological character matrix to estimate the phylogenetic interrelationships of 336 species of shorebirds (~89% of the extant diversity). This taxon sample substantially exceeds that of any previous study not based on supertrees or stochastic polytomy resolvers (Figure 2), making it possible to address outstanding areas of uncertainty due to insufficient sampling. Furthermore, we combine the resulting comprehensive total-evidence phylogeny with an up-to-date, extensively vetted set of 16 fossil calibrations to resolve the controversial timescale of shorebird evolution, and show how inferences about the tempo and mode of charadriiform diversification are sensitive to the estimated divergence times.

## 2 Material and methods

### 2.1 Data assembly and alignment

We obtained published sequences from GenBank (Benson et al., 2013) using Geneious (Kearse et al., 2012) or manual queries via the NCBI web interface. To ensure all available data was obtained, taxonomic coverage was further checked against a previous study attempting to comprehensively sample charadriiform species represented by molecular data (Burleigh et al., 2015). Taxonomy followed the Taxonomy in Flux (TiF) checklist (Boyd, 2019), which includes splitting of the traditionally broad genera *Charadrius*, *Larus*, and *Sterna*. A custom synonym dictionary and R script (R Core Team, 2019) were used to reconcile differences between the NCBI taxonomy and the TiF checklist.

To date, phylogenomic analyses have not conclusively identified the sister group of shorebirds (Hackett et al., 2008; Kimball et al., 2013; McCormack et al., 2013; Yuri et al., 2013; Jarvis et al., 2014; Burleigh et al., 2015; Prum et al., 2015). The Charadriiformes represent one of the six (Houde et al., 2019) to nine (Suh, 2016) major neoavian lineages whose interrelationships remain unresolved even with genome-scale data (Jarvis et al., 2014; Prum et al., 2015; Reddy et al., 2017), and which may constitute a hard polytomy (Suh et al., 2015; Suh, 2016). Here, we chose a gruiform species as the outgroup, as the Gruiformes were found to be the sister group of shorebirds in two recent genomescale analyses (Jarvis et al., 2014; Kuhl et al., 2020). This hypothesis is also supported by phylogenetic analyses of phenotypic data from extant taxa (McKitrick, 1991; Livezey and Zusi, 2007) and a high degree of morphological similarity between the early members of both clades (Musser and Clarke, 2020). To maximize data coverage for this outgroup, we specifically selected the Gray-crowned Crane (*Balearica regulorum*), a taxon for which both the complete nuclear genome (Zhang et al., 2014) and the complete mitochondrial genome (Krajewski et al., 2010) are available.

In assembling our dataset, we selected loci with the largest number of available sequences, and required at least 20 species to have available data for each gene to exclude low-coverage loci. The final sample included 2 mitochondrial ribosomal genes, 13 mitochondrial protein-coding genes, and 9 nuclear protein-coding genes represented by intronic and/or exonic sequences (Table 1). We aligned the sequences using MUSCLE (Edgar, 2004) as implemented in Geneious (Kearse et al., 2012) or as a standalone program. Alignments were visually inspected in AliView (Larsson, 2014) and manually edited if necessary. Reading frames in exonic sequences were identified using amino acid translation and were employed to check for poorly aligned sequences. A custom R script was used to exclude sequences consisting entirely of gaps or undetermined nucleotides.

**Table 1:** Information for the 24 loci used to construct the concatenated alignment. The table includes alignment length, number of sequences available (based on the TiF taxonomy and after the exclusion of conspecifics), and average pairwise identity.

| Code | Gene | Aligned length (bp) | Species sampled | Avg. pairwise identity (%) |
| --- | --- | --- | --- | --- |
| *Mitochondrial* |  |  |  |  |
| 12S | 12S ribosomal RNA | 1044 | 194 | 84.0 |
| 16S | 16S ribosomal RNA | 1735 | 113 | 82.5 |
| ATP6 | ATP synthase membrane subunit 6 | 675 | 84 | 83.0 |
| ATP8 | ATP synthase membrane subunit 8 | 175 | 73 | 79.7 |
| COX1 | Cytochrome *c* oxidase subunit I | 1552 | 248 | 85.9 |
| COX2 | Cytochrome *c* oxidase subunit II | 691 | 63 | 85.8 |
| COX3 | Cytochrome *c* oxidase subunit III | 784 | 57 | 86.9 |
| CytB | Cytochrome *b* | 1204 | 261 | 84.0 |
| ND1 | NADH dehydrogenase subunit 1 | 988 | 74 | 83.0 |
| ND2 | NADH dehydrogenase subunit 2 | 1059 | 210 | 81.7 |
| ND3 | NADH dehydrogenase subunit 3 | 352 | 92 | 84.0 |
| ND4 | NADH dehydrogenase subunit 4 | 1378 | 60 | 83.0 |
| ND4L | NADH dehydrogenase subunit 4L | 290 | 60 | 83.4 |
| ND5 | NADH dehydrogenase subunit 5 | 1832 | 92 | 83.8 |
| ND6 | NADH dehydrogenase subunit 6 | 527 | 70 | 83.2 |
| *Nuclear* |  |  |  |  |
| ADNH | Alcohol dehydrogenase 1, exon 5 to exon 6 | 883 | 54 | 90.0 |
| ALDOB | Fructose-bisphosphate aldolase B, exon 3 to exon 8 | 2217 | 27 | 84.8 |
| BDNF | Brain-derived neurotrophic factor | 689 | 22 | 97.5 |
| CMOS | Oocyte maturation factor Mos | 648 | 23 | 95.0 |
| FGB7 | Beta-fibrinogen, intron 7 | 1336 | 70 | 83.6 |
| GAPDH | Glyceraldehyde-3-phosphate dehydrogenase, intron 3 to intron 5 | 802 | 28 | 81.0 |
| MB2 | Myoglobin, intron 2 | 708 | 61 | 92.3 |
| NTF3 | Neurotrophin-3 | 730 | 23 | 96.9 |
| RAG1 | Recombination activating gene 1 | 3138 | 190 | 92.9 |

In addition to molecular data, we also employed the morphological matrix of Strauch (1978), which we first modified based on the recommendations of Chu (1995). After the modifications, the matrix comprised a total of 69 characters scored for 225 taxa, including 61 parsimony-informative characters and 8 autapomorphies. Unlike the molecular alignment, the morphological matrix did not include an outgroup.

### 2.2 Gene tree and species tree analyses

We subjected the 24 individual gene alignments to two iterations of tree searches to identify mislabeled and conspecific sequences. Preliminary maximum likelihood (ML) gene trees were inferred using RAxML v8.2.12 (Stamatakis, 2014) under the general time-reversible model with discrete gamma-distributed among-site rate variation (GTR+Γ), no codon position partitioning, and the slower but more thorough traditional search option (-f o). The resulting trees were inspected and accessions that violated the monophyly of well-established families were regarded as mislabeled. In addition, conspecific sequences were excluded at this stage; whenever the sequences were not identical, only the most complete one was retained.

The pruned alignments were then refined using MUSCLE and used for the second iteration of gene tree inference. Main tree searches followed the same settings as in the first iteration. Additionally, RAxML bootstrap analyses with 1000 pseudoreplicates were performed for all nuclear genes. A different treatment was applied to the mitochondrial loci, which constitute a single nonrecombinant unit or “superlocus” due to linkage, and are therefore not expected to provide independent estimates of the species tree (Reyes et al., 2004; Brown and van Tuinen, 2011; Richards et al., 2018). To account for this nonindependence, we concatenated the pruned and refined alignments for all mitochondrial loci using the Python package PHYLUCE (Faircloth, 2015). To find the best partitioning scheme for the mitochondrial genome as a whole, we initialized PartitionFinder 2 (Lanfear et al., 2016) with a scheme partitioned both by locus and by codon position for the protein-coding genes (41 partitions) and performed a greedy search using the Bayesian Information Criterion (BIC) to evaluate model fit. The set of candidate substitution models was restricted to those implemented in RAxML. The best-fit scheme consisted of 13 partitions, all of which favored either the GTR+Γ or GTR+Γ+I model. However, since the Γ and I parameters are not identifiable (Yang, 2014), we chose to analyze the mitogenome alignment with GTR+Γ assigned to all partitions. Except for partitioning, the RAxML settings for the main tree search and bootstrapping followed those applied to other gene tree analyses.

To accommodate potential gene tree conflict due to incomplete lineage sorting (ILS) or hybridization, we used ASTRAL-III v5.7.3 (Mirarab et al., 2014b; Zhang et al., 2018) to infer a species tree. Given a profile of unrooted and possibly only partially resolved gene trees, ASTRAL-III performs a limited tree search restricted to combinations of bipartitions observed in the source trees to find the species tree sharing the largest number of induced quartets with the profile. Assuming error-free gene trees, this method represents an estimator of the species tree topology that is statistically consistent even in the presence of heavy ILS which renders concatenation-based approaches positively misleading (Degnan and Rosenberg, 2006; Mirarab et al., 2014b,a). To minimize gene tree error, we collapsed all branches with bootstrap support of *<*10% using a custom R script. The cutoff was chosen based on previous simulation results showing this value to outperform both unfiltered analyses and more aggressive filtering (Zhang et al., 2018). The 1,000 bootstrap pseudoreplicates inferred for each of the 10 gene trees (9 nuclear loci and the mitogenome) were used to expand the search space but did not affect the quartet score, which was calculated from the ML estimates alone. We used the `-t 2` flag to annotate the species tree with internal branch lengths in coalescent units (inversely proportional to the amount of gene tree discordance if the latter were due entirely to ILS), the effective number of genes per branch (Sayyari and Mirarab, 2018), and local posterior probabilities (localPP; Sayyari and Mirarab, 2016). The localPP metric is a conservative measure of support that assumes error-free gene trees and the perfect accuracy of the four subtrees surrounding the focal branch. Its value depends on the frequency with which the branch appears in the input gene trees, as well as the total number of gene trees evaluated (Sayyari and Mirarab, 2016).

The resulting ASTRAL tree included multiple quadripartitions whose quartet support was lower than that of the alternatives, including two at the interfamilial level (involving *Pluvianus* and *Dromas*; see Results). To determine whether this was due to interaction between adjacent quadripartitions (whereby the inclusion of a better-supported alternative would decrease the quartet support of surrounding branches), or merely the failure of ASTRAL to sufficiently explore the search space, we quantified the support for alternative topologies differing in the positions of the two taxa. As there are three possible resolutions for each of the two quadripartitions of interest, with one of their 9 combinations already included in the best tree, we scored an additional 8 topologies using the -q flag and compared them to the best ASTRAL estimate in terms of overall quartet support. We used DiscoVista (Sayyari et al., 2018) to visualize the quartet frequencies and examine the relative strength of gene tree discordance with respect to the best-supported and alternative interfamilial relationships.

### 2.3 Concatenated analyses

We used PHYLUCE (Faircloth, 2015) to concatenate the pruned and refined alignments of all 24 genes, and calculated the partial decisiveness (Sanderson et al., 2010) of the resulting supermatrix using the Python package SUMAC (Freyman, 2015). To identify the best partitioning scheme, we provided PartitionFinder2 (Lanfear et al., 2016) with an initial scheme consisting of 94 partitions (2 for the ribosomal RNA genes, 39 for the three codon positions of the 13 mitochondrial protein-coding loci, 42 for the three codon positions of the 14 nuclear exons, and 11 for nuclear introns). A greedy search using the BIC score for model comparison found the best scheme to comprise 22 partitions. Having assigned a separate GTR+Γ model to each partition, we then used this scheme to perform another RAxML analysis as well as a bootstrap run of 1000 pseudoreplicates.

The use of resampling techniques, such as bootstrap, for measuring branch support has recently been criticized, since their application to phylogenomic-scale data is likely to result in high values even in the presence of substantial intra-dataset conflict (Salichos and Rokas, 2013; Suh, 2016). To take this criticism into account, we further estimated internode certainty (IC; Salichos and Rokas, 2013; Salichos et al., 2014; Kobert et al., 2016) for every branch of our ML tree. IC is an entropy-based measure of incongruence that is calculated for a reference tree based on a collection of (partial) alternative trees. It ranges from 1 (in the case of no conflict) through 0 (if the bipartition in the reference tree is present in the same percentage of trees as the most prevalent conflicting bipartition) to negative values (if there are more frequent alternatives to the bipartition present in the reference tree; Kobert et al. 2016). Since IC is only robust as a measure of support when the collection of alternative trees is sufficiently large (Salichos et al., 2014), we chose to use the bootstrap set for this purpose, and scored the ML tree accordingly using the `-f i -t` RAxML flags.

In addition to the ML analysis, we also performed Bayesian phylogenetic inference using ExaBayes v1.5 (Aberer et al., 2014) via the CIPRES portal (Miller et al., 2010). We used the best partitioning scheme described above, with a separate GTR+Γ substitution model for each partition and branch lengths linked across partitions. We placed flat Dirichlet priors on the exchangeability and state frequency vectors, a *U* (0, 200) prior on the shape parameter of the Γ distribution, and an Exp(10) prior on branch lengths (all defaults). A total of 4 independent runs were executed, each consisting of one cold and three incrementally heated Metropolis-coupled chains of 5 million generations. Convergence was assessed by ensuring that the average standard deviation of split frequencies (ASDSF) did not exceed 5% and by visually inspecting trends in scalar parameter values along each chain in Tracer v1.7 (Rambaut et al., 2018). This led us to exclude one of the 4 runs and set the burn-in proportion for the remaining three to 40%, which ensured that the effective sample sizes (ESS) exceeded 200 for all but 5 of the 246 continuous parameters. Finally, after the first 40% of trees were removed from each file, we combined the tree files from the three runs and used the program ‘consense’ from the ExaBayes suite to generate the extended majority-rule (MRE) consensus tree.

To identify which regions of the ML and Bayesian trees may have been affected by alignment incompleteness, we used the protocol introduced by McCraney et al. (2020) to calculate per-branch locus coverage, i.e., the number of genes informative (but not necessarily supportive) with respect to a given branch. We first pruned both trees down to the species sampled for each of the 24 genes, and used ASTRAL-III (Zhang et al., 2018) with the -q and -t 2 flags to score and annotate the original RAxML and ExaBayes trees based on the resulting set of subsampled trees. Since the pruned trees only differ from the reference trees in terms of taxon coverage rather topology, the effective number of loci computed by ASTRAL (defined as the number of gene trees that contain one of the three possible quartets around the branch of interest; Sayyari et al. 2018) is simply equal to per-branch locus coverage.

### 2.4 Combined analyses

We employed two different methods to expand our taxon sample by including species represented only by morphological data. First, we combined the concatenated alignment with the morphological character matrix of Strauch (1978), modified as described above. The resulting total-evidence supermatrix consisted of 27 partitions: 22 for the nucleotide data, delimited according to the best scheme described above; and 5 for the morphological characters, which were grouped into partitions according to the number of states (ranging from 2 to 6). This supermatrix was analyzed using RAxML-NG (Kozlov et al., 2019) after assigning the GTR model to the nucleotide partitions and appropriate M*k* models (with *k* again ranging from 2 to 6) to the morphological partitions. A Γ distribution discretized into 4 categories was used to account for among-site rate variation within all models except those applied to the 5-state and 6-state morphological partitions, which contained fewer characters than there were rate categories. Since the matrix of Strauch (1978) only contained variable characters, an ascertainment bias correction (Lewis, 2001) was applied to all morphological models. We conducted 100 tree searches (with 50 random and 50 parsimony starting trees) followed by a bootstrap analysis with 1000 pseudoreplicates run on the CIPRES Science Gateway (Miller et al., 2010). In these total-evidence (TE) analyses, the morphological characters helped determine the overall topology of the tree by informing the relationships among taxa represented by both morphological and molecular data.

Second, we performed an alternative analysis using the evolutionary placement algorithm (EPA; Berger and Stamatakis, 2010; Berger et al., 2011), in which the ML estimate from the concatenated analysis served as a fixed molecular scaffold, and the contribution of the morphological characters was limited to attaching to this scaffold the 31 species from Strauch’s (1978) matrix that lacked sequence data. To achieve this, we first pruned the 306-tip concatenated ML tree down to the 194 tips that were represented by both molecular and morphological data, and then used RAxML (-f u) to compute weights for the 69 characters based on their fit to this subsampled reference phylogeny, expressed in terms of per-site log likelihood scores (Berger et al., 2011). Since the weights were equal for all characters, we ran an unweighted EPA analysis using the -f v RAxML flag on the complete 306-tip scaffold. Next, we passed the resulting .jplace files to the program GAPPA (Czech et al., 2020) and grafted the morphology-only taxa onto the molecular scaffold at their most likely insertion positions. To obtain a strictly bifurcating topology comparable to the TE tree, we ran GAPPA with the --fully-resolve flag, allowing the edges of the scaffold tree to be split by the insertion edges according to the proximal length of the placements. Finally, we used a suite of likelihood tests implemented in IQ-TREE v1.6.12 (Nguyen et al., 2014), including the Shimodaira-Hasegawa (SH; Shimodaira and Hasegawa, 1999) and approximately unbiased (AU; Shimodaira, 2002) tests, to compare the fit of the TE and EPA trees to the combined dataset.

### 2.5 Fossil calibrations

We assembled a total of 16 calibrations (Table 2): 9 from a recently published, expert-vetted compendium (Smith, 2015), two that were utilized in a previous divergence time analysis (De Pietri et al., 2020), and 5 that were used here for the first time. Eight out of the 16 calibrations were described in or after 2010; 4 were described in or after 2015. All calibrations were thoroughly vetted to ensure compliance with the criteria of Parham et al. (2011). The 11 calibrations taken from earlier studies were revised following recent geochronological and phylogenetic studies; in all cases, this resulted in changes to their numeric ages, and in one case, the reassignment of a calibration to a different node (see Supplementary Information for details). We paid particular attention to the choice of the root calibration, since multiple fossils have been put forward as the earliest known shorebird remains (Mayr, 2000, 2016; Smith, 2015; Hood et al., 2019). In particular, a recent phylogenetic analysis suggested a crown-charadriiform affinity for at least two fossils dating to the early Eocene (Musser and Clarke, 2020). However, the resulting trees either exhibited a lack of resolution within Charadriiformes or contradicted well-established molecular results, rendering the placement of the fossils within the crown inconclusive.

**Table 2:** Fossil calibrations used for divergence time estimation. Specimen numbers are given for remains belonging to extant species and fossils not assigned to a named species. *t_L_* = minimum, *t_U_* = soft maximum (95th percentile), Cauchy = truncated Cauchy distribution with scale parameter *ct_L_* (Inoue et al., 2009), Soft-bounded uniform = uniform distribution *U* (*t_L_, t_U_*) with a power-decay left tail (0, *t_L_*) and an exponential-decay right tail (*t_U_*, *∞*) (Yang and Rannala, 2005). Detailed justification and additional references for the choice of calibrations and their numeric ages are provided in Supplementary Information.

| # | Fossil | Node calibrated | $t_L$ (Ma) | $t_U$ (Ma) | Density |
| --- | --- | --- | --- | --- | --- |
| 1 | CASG 71892 (*Uria lomvia*) | *Uria lomvia* + *U. aalge* | 2.58 | 44.01 | Cauchy ($c = 1.264$) |
| 2 | *Miocepphus bohaskai* | *Uria* + *Alle* | 18.1 | 57.62 | Cauchy ($c = 0.172$) |
| 3 | *Synthliboramphus rineyi* | *Synthliboramphus craveri* + *S. hypoleucus* | 1.73 | 43.89 | Cauchy ($c = 1.918$) |
| 4 | *Cepphus olsoni* | *Cepphus columba* + *C. carbo* | 6.6 | 44.48 | Cauchy ($c = 0.452$) |
| 5 | *Brachyramphus dunkeli* | *Brachyramphus marmoratus* + *B. brevirostris* | 1.73 | 43.89 | Cauchy ($c = 1.918$) |
| 6 | USNM 192994, USNM 215783<br>(*Fratercula* cff. *arctica*) | *Fratercula arctica* + *F. corniculata* | 3.92 | 32.03 | Cauchy ($c = 0.564$) |
| 7 | *Aethia barnesi* | Crown-group Fraterculinae | 6.6 | 44.48 | Cauchy ($c = 0.452$) |
| 8 | GCVP 5690 | Crown-group Alcoidea | 34.44 | 70.04 | Cauchy ($c = 0.081$) |
| 9 | *Mirolia* spp. | Crown-group Arenariinae | 12.6 | 37.77 | Cauchy ($c = 0.157$) |
| 10 | *Gallinago azovica* | *Gallinago* + *Coenocorypha* | 6.1 | 28.39 | Cauchy ($c = 0.288$) |
| 11 | *Elorius* spp., *Parvelorius* spp. | Crown-group Scolopacidae | 20.0 | 48.32 | Cauchy ($c = 0.111$) |
| 12 | *Nupharanassa tolutaria* | Crown-group Jacanoidea | 30.5 | 59.12 | Cauchy ($c = 0.074$) |
| 13 | *Oligonomus milleri* | Crown-group Thinocoroidea | 24.47 | 48.98 | Cauchy ($c = 0.079$) |
| 14 | NMB S.G.20252 | Crown-group Haematopodoidea | 20.0 | 68.50 | Cauchy ($c = 0.191$) |
| 15 | *Chionoides australiensis* | Crown-group Chionida | 24.76 | 68.94 | Cauchy ($c = 0.140$) |
| 16 | IGM 100/1435 | Crown-group Charadriiformes | 55.88 | 66.0 | Soft-bounded uniform |

To assess the confidence in the phylogenetic position of the early Eocene fossils, we re-analyzed the morphological character matrix of Musser and Clarke (2020) under three sets of “soft” topological constraints which fixed the relationships among most of the extant taxa but allowed the 8 included fossils to attach anywhere in the tree. The constraints enforced interfamilial relationships within all orders represented by more than two taxa (Anseriformes, Charadriiformes, Gruiformes) in addition to all applicable supraordinal relationships from three recent avian phylogenomic trees (Prum et al., 2015; Reddy et al., 2017; Kimball et al., 2019). Note that in the analyses of Musser and Clarke (2020), the interfamilial relationships within Charadriiformes were left unconstrained, and the supraordinal relationships were only selectively enforced. All three analyses were performed using MrBayes v3.2.6 (Ronquist et al., 2012b) under the M*k*+Γ substitution model and an Exp(10) branch length prior. For each analysis, we ran 4 independent replicates, each with 4 Metropolis-coupled chains (three of which were incrementally heated) of 20 million generations. After discarding the first 40% of states as burn-in, we verified that the analyses reached convergence based on the ASDSF (*<*0.01), ESS values for scalar parameters (*>*200), and potential scale reduction factor (PSRF; *∼*1.00). Finally, we used the MRE consensus tree to summarize the post-burn-in posterior sample.

A long-recognized problem of node-dating analyses is the fact that fossil calibrations can only provide a reliable lower bound on the age of any given clade (Yang and Rannala, 2005; Benton and Donoghue, 2006; Wilkinson et al., 2010). To avoid arbitrary upper bounds, we used a simple, Bayesian method devised by Hedman (2010) and first applied to fossil calibration design by Friedman et al. (2013). In this approach, the age of origin of a clade is informed by the sequence of the first appearance dates of its successive outgroups. The algorithm starts with the assumption that the age of the node connecting the most distant outgroup to the clade of interest is uniformly distributed between the first appearance of that outgroup and some arbitrary upper bound *t*_0_. The next outgroup diverges at a time that is uniformly distributed between its own first appearance date and the divergence time of the previous outgroup, integrating over the uncertainty in the latter. The process is then repeated until a non-uniform posterior distribution is obtained on the interval [*t_L_, t*_0_], where *t_L_* is the oldest known fossil belonging to the clade of interest (Hedman, 2010). Following Friedman et al. (2013), we took the conservative approach of excluding stratigraphically inconsistent outgroups (i.e., those that appear later in the fossil record despite diverging earlier). Together with the uncertain relationships within Neoaves (Jarvis et al., 2014; Prum et al., 2015; Suh, 2016) and the fact that few neornithine lineages predate the oldest known charadriiform occurrence from the Paleocene/Eocene boundary (Mayr, 2014; Ksepka et al., 2017), this required extending the outgroup sequence into the neornithine stem group. We used recent phylogenies of Mesozoic birds to construct several alternative sequences (Supplementary Information). After setting *t*_0_ = 160 Ma and evaluating the node age distributions at 1000 discrete time steps, we calculated the 95% credibility interval (CI) for the age of each calibrated node. We then averaged the 95% CI upper bounds across the different outgroup sequences to account for phylogenetic uncertainty, and used the resulting value as a soft maximum when designing the corresponding calibration density (Table 2).

While parameter-rich calibration densities offer substantial flexibility (Brown and van Tuinen, 2011), they also run the risk of being arbitrary, since it is frequently only the minimum and the maximum age that can be objectively determined for a given calibration (but see Wilkinson et al., 2010; Claramunt and Cracraft, 2015). Arbitrarily specified parameters can have a huge impact on the resulting node age posteriors (Inoue et al., 2009; Warnock et al., 2012). To avoid this, we constrained the truncated Cauchy densities implemented in the program MCMCTree (Yang, 2007; Inoue et al., 2009) to approximate two-parameter offset exponentials, which are fully determined by their minimum (offset) and soft maximum (95th percentile). Starting with a truncated Cauchy distribution of the form *t ∼ L*(*t_L_, p, c, p_L_*), we set *p_L_* (the probability of violating the lower bound *t_L_*) to 10*^−^*^300^ to approximate a hard minimum, and fixed *p* (the offset of the mode from the minimum) to 0 to ensure the monotonic decline of the density function after the fossil-based minimum. Finally, we analytically calculated *c* (parameter controlling the rate of the decline; Table 2) to ensure that 95% of the total probability mass would be located between the fossil-based hard minimum *t_L_* and the soft maximum *t_U_* estimated using the outgroup method. Since MCMCTree does not accept Cauchy densities at the root of the tree, a soft-bounded uniform density was used instead for the root calibration (Table 2).

### 2.6 Divergence time estimation

We used the program MCMCTree from the PAML v4.9j suite (Yang, 2007) to estimate node ages on the fixed topology of the total-evidence RAxML-NG tree based on the 16 fossil calibrations described above. We performed a number of preliminary analyses that either had difficulty reaching stationarity, even after very long Markov chain Monte Carlo (MCMC) runs, or were found to be computationally intractable due to the large number of partitions used (Supplementary Information). Comparisons with additional test runs sampling from the prior suggested that these problems were caused by the misspecification of the relaxed clock priors, large amounts of rate heterogeneity across the alignment, and nonrandom distribution of missing data across the tree. To mitigate these issues, we set out to identify a subset of our sequences evolving in a homogeneous and a relatively clock-like manner.

#### 2.6.1 Locus filtering

Using the R packages adephylo (Jombart et al., 2010), phangorn (Schliep, 2010), and phytools (Revell, 2012), we reimplemented the SortaDate pipeline (Smith et al., 2018) to carry out a multistep filtering workflow on the gene trees described in section 2.2. In SortaDate, individual gene trees are evaluated for their length (correlated with the rate of evolution), root-to-tip path length variance (a proxy for rate heterogeneity across lineages), and percentage of bipartitions shared with a reference tree (quantifying topological congruence with the species tree to be dated). We used the total-evidence RAxML-NG phylogeny as our reference tree, and after pruning it down to the taxon sample of a given gene tree, we calculated the percentage of shared bipartitions as 1 *− d/*(2*n −* 6), where *d* is the unnormalized Robinson-Foulds (RF) distance (Robinson and Foulds, 1981) and *n* is the number of tips represented by sequence data for the gene in question.

Despite our overall focus on rate homogeneity, we chose to include CytB (the least clock-like of all the genes examined: root-to-tip path length variance = 0.0248) in the filtered set of loci because of its high taxonomic coverage (Table 1). Next, we excluded four loci sharing fewer than 50% bipartitions with the reference tree, and ranked the remaining 19 loci from the most (7.72 × 10*^−^*^5^) to the least (0.0208) clock-like. We then kept adding loci in this order until the cumulative taxon sample included all 306 species for which at least some molecular data were available. The final dataset consisted of 8 loci comprising 10,883 base pairs: ADNH, ALDOB, COX1, CytB, GAPDH, ND3, NTF3, and RAG1.

#### 2.6.2 Relaxed clock hyperpriors

We used the program BASEML (Yang, 2007) bundled with MCMCTree to estimate the mean substitution rate for the 8-locus dataset under a strict molecular clock. A single calibration was placed at the root of the tree and assigned a point-mass prior set to the mean of the soft uniform density to be used for the subsequent divergence time analysis (Table 2). We used a time unit of 10 million years, resulting in a value of 6.108 assigned to the root prior. The maximum likelihood estimate of the substitution rate was 2.456 × 10*^−^*^9^ substitutions per site per year (ss*^−^*^1^y*^−^*^1^), which we used as the mean of a diffuse gamma hyperprior on the mean clock rate hyperparameter (rgene gamma) with shape *α* = 2, rate *β* = 81.44, and the 95% prior CI of [2.965 × 10*^−^*^10^, 6.852 × 10*^−^*^9^] ss*^−^*^1^y*^−^*^1^. Following McGowen et al. (2019), we placed a gamma hyperprior with shape *α* = 2 and rate *β* = 2 on the variance of the logarithm of the clock rate (sigma2 gamma). Assuming mean values for both hyperpriors, these choices specified a highly diffuse prior distribution of branch rates (95% prior CI: [2.641 × 10*^−^*^10^, 1.326 × 10*^−^*^8^] ss*^−^*^1^y*^−^*^1^), intended to accommodate the large amount of expected rate heterogeneity among genome types (nuclear vs. mitochondrial) and codon positions.

#### 2.6.3 Bayesian clock model selection

We used Bayes factors to compare two relaxed clock models implemented in MCMCTree. Specifically, we evaluated the marginal likelihoods and posterior probabilities of the uncorrelated or independent-rates (IR) model, in which branch rates are drawn from a lognormal distribution whose mean and variance are themselves estimated from the data as hyperparameters (Drummond et al., 2006), and the geometric Brownian motion or autocorrelated-rates (AR) model, in which the logarithm of the substitution rate diffuses along the tree following Brownian motion. Consequently, in the AR model, the rate at the midpoint of a given branch is drawn from the distribution with variance proportional to the length of that branch, and with a mean equal to the rate at the parent branch (Rannala and Yang, 2007). To estimate the marginal likelihoods, we used the stepping-stone sampling technique (Xie et al., 2011) as implemented in the R package mcmc3r (dos Reis et al., 2018). This method belongs to a broader class of path-sampling approaches, in which a number of unnormalized “power posterior” distributions of the form *f_β_* = (prior) × (likelihood)*^β^* are constructed along a path between the prior (*β* = 0) and the posterior (*β* = 1). The product of the ratios of the estimated normalizing factors of the successive power posteriors then approximates the marginal likelihood (Xie et al., 2011).

Because sampling from the power posteriors requires the computationally intensive calculation of exact likelihoods, the stepping-stone method is not practical for large datasets containing hundreds of tips and multiple calibrations (dos Reis et al., 2018; McGowen et al., 2019). We therefore chose to perform model comparisons on two subsets of the full 336-species sample. First, we generated a subset of 19 species by drawing one representative from every family. For the non-monotypic families, we selected the species with the highest sequence coverage in the filtered 8-locus alignment. Since the signal of rate autocorrelation is expected to decay over long periods of time, it may be difficult to detect from sparse phylogenies of old clades (Drummond et al., 2006; Brown and van Tuinen, 2011), running the risk that the 19-species tree may be biased toward the IR model. To alleviate this risk, we enlarged the first subset to 40 species, purposely using diversified sampling (Höohna et al., 2011) to break down long branches. Specifically, we identified the deepest split within each non-monotypic family, and drew a representative from that side of the split which was not yet represented in the 19-species subset. As before, if multiple species satisfied this condition, the highest-coverage one was selected. Finally, after sampling two species from each non-monotypic family following the aforementioned criteria, we kept adding the highest-coverage species not yet included in the subset regardless of their phylogenetic position, until a total of 40 species was reached.

We used 16 *β* points to construct the sampling path for the analysis of the 19-species dataset, drawing 20,000 samples from each power posterior at the frequency of one sample per 20 generations. For the 40-species dataset, we used 64 *β* points associated with shorter chains, such that each power posterior was represented by 6000 samples collected every 20 generations. Since only relative rather than absolute node ages were of interest at this stage of the analysis, we used a single narrow *U* [0.999, 1.001] calibration assigned to the root. The birth-death-sampling hyperparameters were fixed to `BDparas = 1 1 0.05` or `BDparas = 1 1 0.105` for the 19-species and 40-species datasets, respectively. In both cases, the sampling fraction corresponded to the proportion of the known extant charadriiform diversity (380 species) represented in a given tree. The substitution model and its associated priors, as well as the relaxed clock hyperpriors, were identical to those used for the main analysis.

#### 2.6.4 Main and sensitivity analyses

The main analyses were performed using a single HKY85+Γ model applied to the entire 8-locus alignment. We used 5 categories to discretize the Γ among-site rate distribution, assigning informative priors to its shape parameter *α* and to the transition/transversion ratio *κ* (`ncatG = 5, kappa gamma = 6 2, alpha gamma = 2 14`). The birth and death rates were both fixed to 1 species per the time unit of 10 million years following Marshall’s (2017) paleobiological “law” stating that the rates of speciation and extinction tend to be approximately equal, while the sampling fraction was set to the proportion of extant shorebird species included in the 336-species tree (`BDparas = 1 1 0.887`). For both the AR and IR models, we ran two MCMC analyses of 15,000 samples each, sampling every 250 generations. All analyses employed likelihood approximation based on the BASEML estimates of branch lengths (dos Reis and Yang, 2011). To diagnose convergence, we visually inspected the posterior traces from both runs in Tracer (Rambaut et al., 2018), determined the proportion of samples to be discarded as burnin (10%), and verified that after doing so, the ESS values exceeded 200 for all parameters in either run. We further used the ‘postProcParam’ utility from the ExaBayes suite (Aberer et al., 2014) to calculate the PSRF values between the two runs and the ESS values for both runs combined, ensuring that they were less than 1.01 and greater than 200 for all parameters, respectively. Finally, we concatenated the log files from both runs (excluding burnin) using LogCombiner v1.10 (Rambaut and Drummond, 2018), and summarized the combined MCMC sample in the form of posterior mean ages and 95% CIs. To assess the impact of prior interactions and the informativeness of the sequence data with regard to divergence times, we ran MCMC-Tree without data to sample from the joint prior. For both the AR and IR model, a single chain was used to draw 50,000 samples at the frequency of one sample per every 1000 generations.

Finally, to evaluate whether the results were influenced by the lack of partitioning across the relatively heterogeneous alignment, we performed additional analyses on a subset of the 8-locus dataset, which was divided into two partitions expected to exhibit less rate and process variation across sites. The “fast” partition comprised mitochondrial codon position 3, while the “slow” partition consisted of nuclear codon positions 1+2. An independent BASEML analysis was performed as described above to estimate the strict-clock rate for either partition, and the weighted average (by partition length) of the partition-specific rates was used to set up a gamma-Dirichlet hyperprior on the mean substitution rate, with another hyperprior of the same type assigned to the variance of its logarithm. These hyperpriors use a gamma distribution to specify the average value of both hyperparameters across partitions, and apportion their total value among partitions according to a Dirichlet distribution (dos Reis et al., 2014). We specified a flat distribution over all apportionment schemes by setting the concentration parameter to 1 (`rgene gamma = 2 55.92 1, sigma2 gamma = 2 2 1`). As before, a HKY85 model with 5 rate categories was specified for either partition, but a more diffuse prior was assigned to *α* to allow for smaller amounts of within-partition rate heterogeneity (`alpha gamma = 1 1`). All other priors were identical to the main analyses. The partitioned analyses were again conducted under both the AR and IR clock models, with likelihood approximation, two chains of 18,000 samples per analysis, and a sampling period of 250 generations. Convergence was assessed using the same criteria as that of the main analyses.

### 2.7 Macroevolutionary rate estimation

To infer the diversification dynamics of the Charadriiformes, we used Bayesian Analysis of Macroevolutionary Mixtures (BAMM) v2.6, a model-averaging approach that employs a time-scaled phylogeny to detect clade-specific shifts between distinct macroevolutionary regimes (Rabosky, 2014; Mitchell et al., 2018). The method relies on reversible-jump MCMC (rjMCMC) to sample models with different numbers of parameters, which correspond to within-regime rates of speciation and extinction. The number of regimes and the locations of the shifts between them are inferred from the data, and the estimates of macroevolutionary rates through time are marginalized over the models involving different regime numbers and shift configurations. To assess the sensitivity of the inferred shift configurations and marginal rates to the modeling choices made upstream in the divergence time estimation step, we performed separate BAMM analyses on both the AR and IR time trees. After taxonomic reconciliation, 388 stratigraphically unique species-level fossil occurrences associated with the tips of both time trees were located in the Paleobiology Database (http://www.paleobiodb.org; last accessed April 22, 2021) to inform the estimated extinction and fossil preservation rates (Mitchell et al., 2018).

We used the R package BAMMtools (Rabosky et al., 2014) to set the priors on the initial rates and the hyperprior on the exponential rate change parameter. The expected number of shifts was set to 1, and within-regime speciation rates were allowed to vary through time. The global sampling fraction was set to 0.887 following the same rationale as in the node dating analyses. For each tree, we performed two rjMCMC runs consisting of 4 Metropolis-coupled chains (one cold and three incrementally heated) of 50 million generations, sampling every 10,000 generations. After examining the posterior traces, the first 10% of samples were discarded as burnin, and convergence was assessed by calculating the ESS (*>* 200) and PSRF (*<* 1.01) using the R package coda (Plummer et al., 2006). We used BAMMtools to process the output, compare the prior and posterior probabilities of different diversification models, and calculate their Bayes factors relative to the best-supported model. Additionally, we computed the 95% credible set of rate shift configurations, and extracted the single maximum *a posteriori* (MAP) configuration. Finally, we summarized marginal macroevolutionary rates through time, calculated the mean rates inside and outside the shifted clade to determine the magnitude of the shift (Upham et al., 2020), and obtained the 95% CIs about the root and tip speciation rates within each regime to evaluate the support for rate variation over time.

## 3 Results

### 3.1 Data assembly and alignment

The final supermatrix obtained by concatenating all 24 loci (Table 1) included 25,437 sites from a single outgroup and 305 species of charadriiforms (Table S1), accounting for *∼*81% of the extant diversity (Dickinson and Remsen, 2013; Boyd, 2019; Clements et al., 2019). This concatenated alignment contained a total of 2,216,525 complete cells (excluding gaps and indeterminate residues), corresponding to 71.5% missing data. The number of genes per taxon (gene occupancy) ranged from 1 to all 24 in the outgroup and 22 in the highest-occupancy ingroup species (killdeer; *Charadrius vociferus*), with an average of 7.3 and a median of 5.5 (Supplementary Information, Fig. S2). Every ingroup species was represented by at least 275 (Olrog’s gull, *Larus atlanticus*; lava gull, *Leucophaeus fuliginosus*) and up to 22,660 (*Charadrius vociferus*) nucleotides (average = 7244, median = 5900). The dataset included representatives of all 19 extant families, of which 6 monotypic (Chionidae, Dromadidae, Ibidorhynchidae, Pedionomidae, Pluvianellidae, Pluvianidae) and 2 non-monotypic (Stercorariidae, Thinocoridae) families were sampled exhaustively. The partial decisiveness of our concatenated alignment was 0.87, meaning that the subtrees induced by the incomplete taxon coverage of the individual genes uniquely define 87% of all possible trees when combined. The inclusion of morphological data increased the number of sampled charadriiform species to 336 (~89% of the extant diversity), and the number of exhaustively sampled families to 9 by completing the sampling of the Jacanidae.

### 3.2 Gene tree and species tree analyses

The initial round of gene tree analyses described in section 2.2 resulted in the removal of 11 species-level duplicates and 18 potentially mislabeled sequences whose phylogenetic positions rendered well-supported families non-monophyletic. The final gene trees contained between 22 (BDNF) and 261 (CytB) tips (Table 1), with an average of 94 and a median of 70 tips per gene tree. On average, more species were included in the trees based on mitochondrial rather than nuclear loci (117 vs. 55 tips, respectively). Treating the entire mitochondrial genome as a single locus, as in the ASTRAL analyses, yielded a tree of 300 tips.

The average local posterior probability (localPP) across the branches of the ASTRAL species tree was 0.68. Almost 40.0% of branches were associated with localPP values greater than 0.7, a threshold considered to represent an adequate balance between accuracy (proportion of branches above the threshold that are correct) and recall (proportion of correct branches that are above the threshold) by Sayyari and Mirarab (2016). Only 7.3% of branches exceeded a more stringent threshold of localPP *>* 0.95. On average, the branches of the species tree were informed by 2.2 genes, with 55.8% of branches represented by at least 2 genes and 7.9% of branches informed by 5 or more genes. The effective number of genes was highest for the deepest branches in the tree, especially those connecting suborder and “parvorder”-level taxa. Using Bayesian correlation testing as implemented in the R package bayestestR (Makowski et al., 2019), we found decisive evidence (*sensu* Kass and Raftery, 1995) for a positive correlation between the localPP of a branch and its effective number of genes (*ρ* = 0.49, Bayes factor in favor of a nonzero *ρ* = 1.72 × 10^17^).

The ASTRAL tree supported the monophyly of the three charadriiform suborders (localPP: Charadrii = 0.60, Scolopaci = 0.87, Lari = 0.78), as well as the sister-group relationship of Scolopaci and Lari to the exclusion of Charadrii (localPP = 1). This was reflected in the generally high support for the corresponding nodes across the 10 gene trees (Figure 3). At the subordinal level, notable discordance between the species tree and gene trees was limited to the monophyly of Charadrii, which was strongly rejected by one locus (FGB7) and weakly rejected by 5 others (ADNH, BDNF, MB2, NTF3, RAG1). Relationships within suborders were mostly also in agreement with recent molecular phylogenies, and all of the “superfamily” or “parvorder”-level clades of Cracraft (2013) were strongly supported by 2–7 gene trees with little to no contradicting signal (Figure 3). The sister-group relationship between Haematopodidae and Recurvirostridae to the exclusion of the ibisbill (*Ibidorhyncha*) received a low support value (localPP = 0.42, equal to the support for the next best arrangement) but was only weakly contradicted by a single gene (RAG1), which supported a sister-group relationship between *Ibidorhyncha* and Haematopodidae instead. In contrast, the position of *Pluvialis* within Charadriidae (localPP = 0.51) was associated with substantial inter-gene conflict (Figure 3), with strong support from two loci (ALDOB, mitogenome) and strong opposition from three others, which either allied *Pluvialis* with Haematopodoidea (ADNH) or with a clade comprising Haematopodoidea and all other charadriids (GAPDH, MB2).

**Figure 3.**
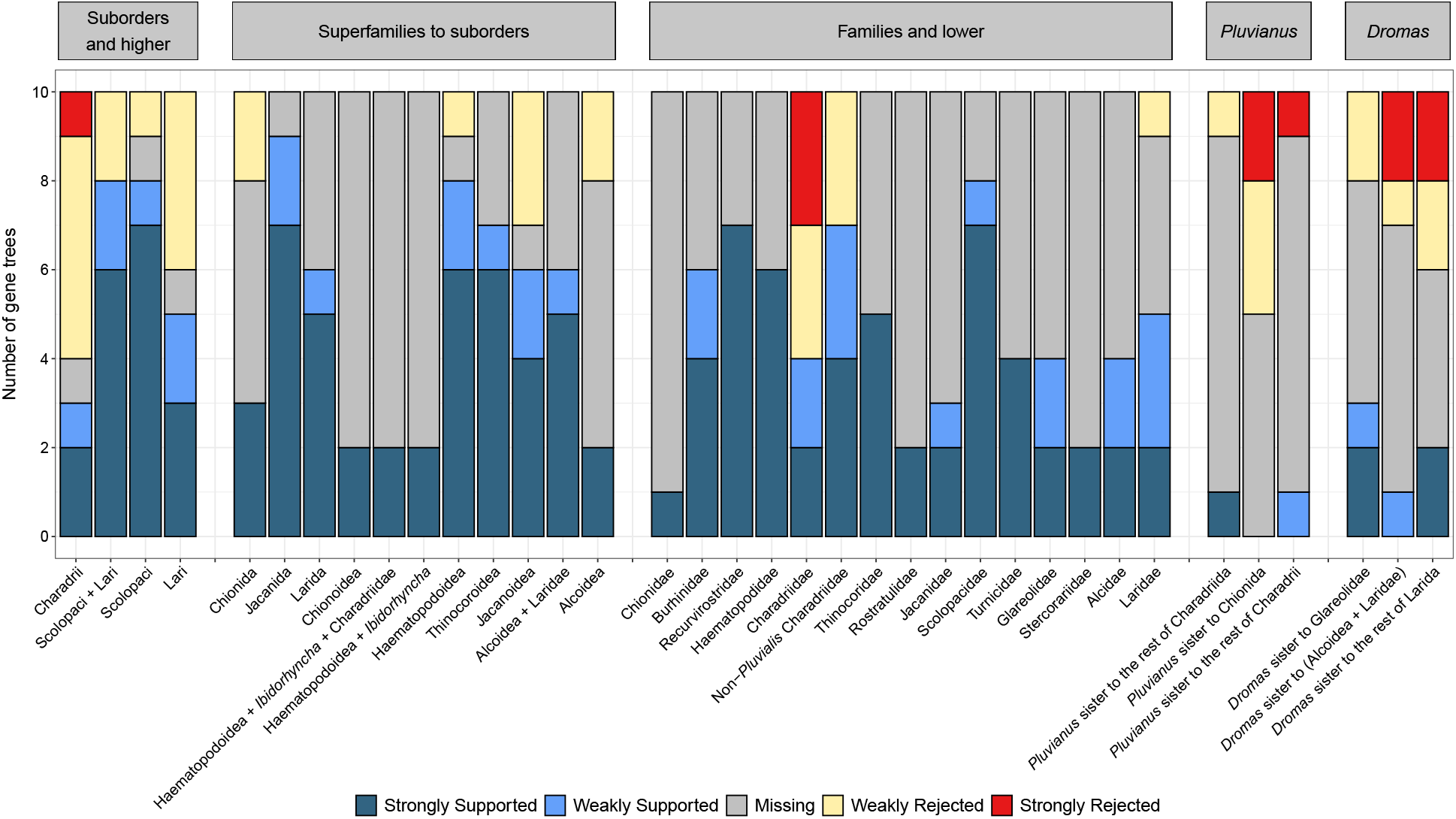
Gene tree support for higher-level clades of shorebirds summarized using DiscoVista. Strong support or rejection are defined by the focal or conflicting clade exceeding a 75% bootstrap threshold, respectively; missing data refers to loci that lacked the taxon sampling needed to evaluate a given branch. For *Pluvianus* and *Dromas*, two family-level taxa whose phylogenetic positions differed between the species-tree and concatenated analyses, all three possible resolutions of the relevant branch have been scored.

Two family-level relationships were subject to conflict between the ASTRAL tree on the one hand and concatenation-based results, as well as previous molecular phylogenies, on the other. These concerned the position of the monotypic families comprising the Egyptian plover (Pluvianidae) and the crab plover (Dromadidae) (Figure 4). The former taxon formed the sister group to the rest of Charadrii (localPP = 0.05), while the latter emerged sister to an (Alcoidea + Laridae) clade (localPP = 0.25), thus violating the monophyly of Charadriida and Glareoloidea *sensu* Cracraft (2013), respectively. Notably, despite the inclusion of these relationships in the final ASTRAL tree, both branches had better-supported alternative resolutions that agreed with the concatenation-based topology and previous analyses in placing *Pluvianus* as the earliest-diverging member of a monophyletic Charadriida (localPP = 0.92), and *Dromas* as the sister group of the pratincoles and coursers (Glareolidae) (localPP = 0.67). This result is also borne out by a detailed examination of the gene tree support for the alternative topologies (Figure 3). The conventional position of *Pluvianus* within Charadriida was strongly supported by a single gene (RAG1; bootstrap = 83%) and weakly opposed by one other locus (mitogenome; bootstrap = 56%) which favored the alternative quadripartition included in the species tree, whereas the inclusion of *Dromas* within Glareoloidea received support from three loci (FGB7: 90%, MB2: 78%, RAG1: 56%) while being contradicted by two others that instead weakly supported a relationship to the auk and gull clade (BDNF: 34%, mitogenome: 43%). However, the failure to include the better-supported alternative positions in the species tree was not due to an inadequate exploration of the search space, as constrained searches performed on topologies enforcing alternative resolutions for the two quadripartitions in question all yielded quartet scores 0.013–0.065% lower than that of the original tree. Enforcement of the conventional position incurred a greater quartet score reduction for *Dromas* (0.043%, averaged over the three possible resolutions for the branch attaching to *Pluvianus*) than for *Pluvianus* (0.036%, averaged over the three possible resolutions for the branch attaching to *Dromas*).

**Figure 4.**
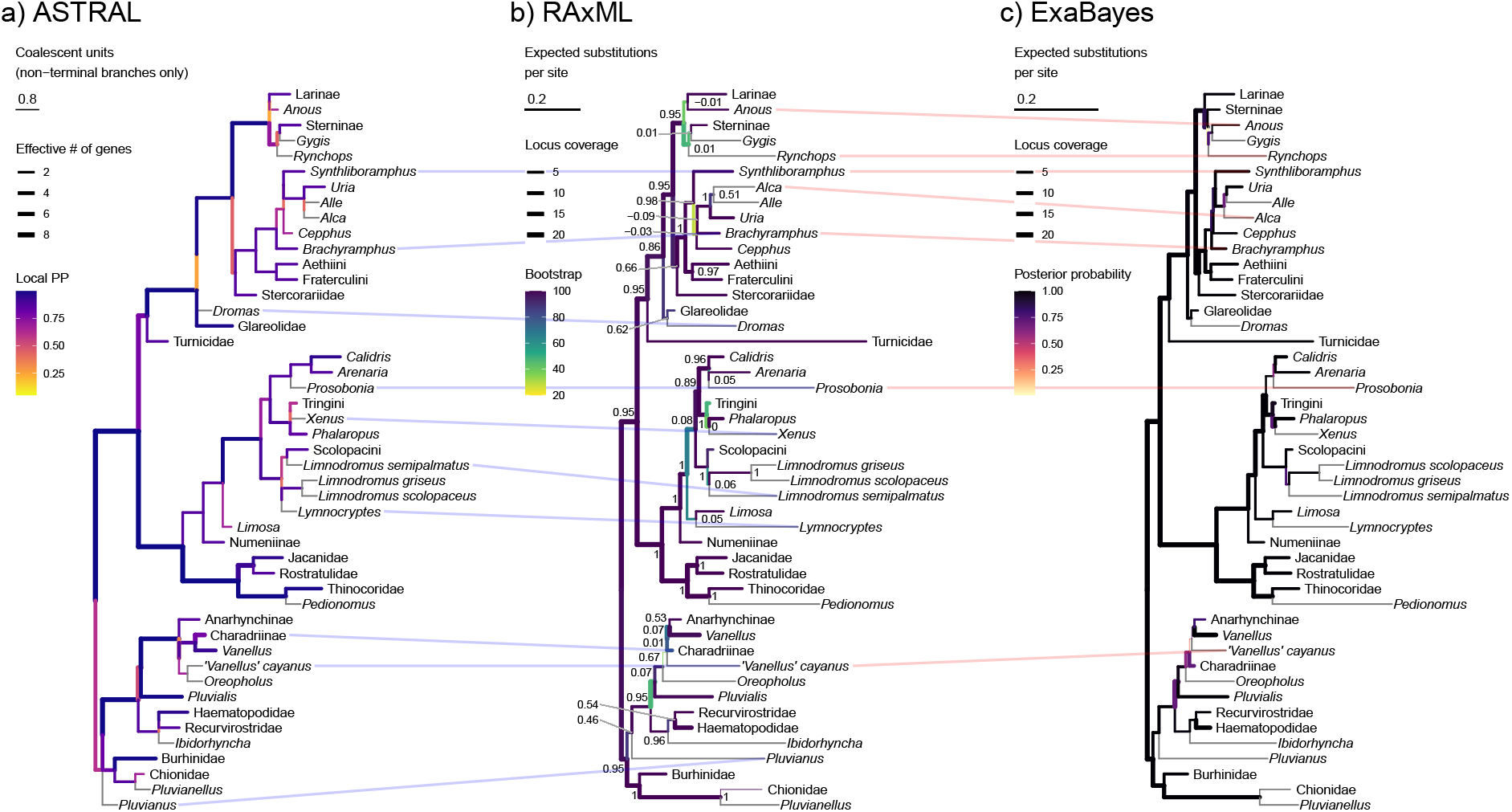
Phylogenetic estimates for the Charadriiformes obtained from species-tree (a), concatenated maximum likelihood (b), and concatenated Bayesian (c) methods. Higher-level clades have been collapsed except when subject to topological conflict; branches subtending terminal taxa are shown in gray. The node labels of the RAxML tree denote internode certainty (IC) values; the light blue and red lines connect taxa with conflicting placements between a given pair of trees. Note that the effective number of genes for the ASTRAL tree is calculated out of a maximum of 10 (mitochondrial genome treated as a single unit), while the per-branch locus coverage of the concatenation-based trees is calculated out of 24 (mitochondrial loci analyzed separately). The full versions of all three trees are given in Supplementary Information (Figs. S3–S8).

### 3.3 Concatenated analyses

The average internode certainty (IC) value across the branches of the RAxML tree was 0.59; of the 304 internal branches present in the unrooted tree, 10.5% had negative IC values (indicating that a conflicting bipartition had greater prevalence in the bootstrap set than the bipartition present in the maximum likelihood tree), and 88.8% had strictly positive internode certainties (indicating that the bipartition in the RAxML tree occurred in the bootstrap set more frequently than any conflicting bipartition). Nearly all the branches associated with negative IC values represented internal relationships within species-rich genera such as *Calidris*, *Larus*, *Ochthodromus*, and *Vanellus*. The average bootstrap support (BS) across the RAxML tree was 79%, with 69.6% of branches exceeding the threshold of BS = 70%. In contrast, the IC value exceeded by the same percentage of branches only amounted to 0.26. On average, the branches of the RAxML tree were informed by 5.9 genes. It should be noted that this value is not directly comparable to that given for the ASTRAL tree, as mitochondrial loci were considered separately when quantifying the locus coverage across the concatenation-based trees. Over 87% of branches were informed by 2 or more loci, with 52% of branches represented by at least 5 loci. We found decisive evidence that the locus coverage of a branch was positively correlated with both its internode uncertainty (*ρ* = 0.24, Bayes factor = 801.3) and its bootstrap support (*ρ* = 0.21, BF = 128.8).

The average posterior probability on the ExaBayes tree was 0.86, with 65% of nodes exceeding the threshold of PP = 0.95. The average locus coverage of a branch in the ExaBayes tree was 5.8 genes, with 86% of branches informed by at least 2 loci and 51% of branches represented by 5 or more loci. Bayesian correlation analysis found decisive evidence for a relationship between the posterior probability of a branch and its locus coverage (*ρ* = 0.28, BF = 4.03 × 10^4^). The average posterior probability of the 50 bipartitions present in the ExaBayes tree but not the RAxML tree was 0.46, while the average bootstrap and internode certainty of the conflicting branches from the RAxML tree amounted to 40% and 0.03, respectively.

The RAxML and ExaBayes analyses of the concatenated alignment agreed on all family-level interrelationships, which were also consistent with previous molecular results and – except for the positions of *Pluvianus* and *Dromas* – with the topology of the species tree (Figure 4). The monophyly of Haematopodoidea to the exclusion of *Ibidorhyncha* received strong support from both approaches (BS = 88%, PP = 1), while the monophyly of the plovers was considerably less robust (BS = 47%, PP = 0.71). Both analyses found strong support for the sister-group relationships between *Dromas* and Glareolidae (BS = 89%, PP = 1) and between *Pluvianus* and the rest of Charadriida (BS = 87%, PP = 1), in contrast to the ASTRAL tree. Major areas of disagreement between the RAxML and ExaBayes topologies included the intrafamilial relationships within the Laridae and the Alcidae. In the former case, both approaches found the gulls (Larinae) outside of a clade including the true terns (Sterninae), white terns (*Gygis*), and the skimmers (*Rynchops*), but disagreed on the position of the noddies (*Anous*), which were allied with the gulls in the RAxML phylogram (BS = 40%) but with *Gygis* in the ExaBayes consensus tree (PP = 0.77). Similarly, among the auks, the interrelationships of *Brachyramphus*, *Synthliboramphus*, the true guillemots (*Cepphus*), and the true auks and murres (*Alca*, *Alle*, *Uria*) differed between the two methods, but with poor support values (BS *<* 50%, PP *<* 0.9) in both cases.

Topological inconsistencies between the concatenated analyses and the ASTRAL estimate were similarly limited to branches that were poorly supported in all three trees (Figure 4). In contrast to previous studies (Baker et al., 2007; Burleigh et al., 2015), both concatenation-based trees found the jacksnipe (*Lymnocryptes*) within Limosinae rather than Scolopacinae (BS = 63%, PP = 1). The species tree weakly upheld a scolopacine affinity for the jacksnipe (localPP = 0.42) but rendered the dowitchers (*Limnodromus*) paraphyletic with respect to the rest of the subfamily (localPP = 0.58; Figure 4). Within the Charadriidae, RAxML and ExaBayes both found the tawny-throated dotterel (*Oreopholus*) to represent the second earliest divergence after *Pluvialis* (BS = 36%, PP = 0.60), consistent with several earlier analyses (Baker et al., 2012; Burleigh et al., 2015). In the species tree, this position was instead occupied by a moderately well-supported clade (localPP = 0.67) uniting *Oreopholus* with the pied lapwing (*“Vanellus” cayanus*), whose lack of a close relationship to other lapwings (*Vanellus*) was also borne out by the concatenated analyses (Figure 4). Aside from this limited conflict, the species-tree and concatenation-based approaches yielded highly similar topologies, as evidenced by the fact that the RAxML–ExaBayes (0.165), RAxML–ASTRAL (0.165), and ExaBayes–ASTRAL (0.221) RF distances were all significantly smaller than the average distance between any of the three trees and 1000 sequence-only topologies from the Jetz et al. (2012) pseudoposterior (RAxML: 0.304, 95% CI: 0.273–0.333; ExaBayes: 0.328, 95% CI: 0.300–0.356; ASTRAL: 0.313, 95% CI: 0.285–0.341). To a lesser extant, this was also true for their distances from the supermatrix-based tree of Burleigh et al. (2015) (RAxML = 0.267, ExaBayes = 0.286, ASTRAL = 0.267).

### 3.4 Combined analyses

The fully resolved tree generated by the evolutionary placement algorithm (EPA) differed from the total-evidence (TE) tree in the positions of 21 out of the 31 species without sequence data (Supplementary Information, Fig. S9). Neither tree perfectly matched the preexisting taxonomy of the clade. Both analyses found the beach stone-curlew (*Burhinus magnirostris*), often placed in the separate genus *Esacus* (Boyd, 2019; Clements et al., 2019), to be deeply nested among other *Burhinus* species. The EPA tree rendered the lapwings (*Vanellus*) polyphyletic by allying the Senegal lapwing (*V. lugubris*) with *Oreopholus*, while the TE analysis weakly favored the polyphyly of the jacana genus *Actophilornis* by inferring a sister-group relationship between *A. albinucha* and *Microparra* (Figure 5). However, the relationships within the Jacanidae were essentially ambiguous in the TE tree, and none of the nodes separating the two species of *Actophilornis* received a bootstrap support value greater than 50%. The same was true of most of the nodes present in the TE tree but not the EPA tree (average BS = 38.4%), and indeed of most of the nodes connecting the morphology-only species to the rest of the tree (average BS = 46.2%). In terms of RF distances, the TE and EPA topologies were substantially closer to each other (RF distance = 0.201) than to a sample of 1000 all-taxa trees from the Jetz et al. (2012) pseudoposterior (TE: 0.520, 95% CI: 0.488–0.550; EPA: 0.510, 95% CI: 0.478–0.541). Despite this high degree of congruence between both analyses, the TE tree was found to fit the combined data significantly better than the EPA tree (Table 3).

**Figure 5.**
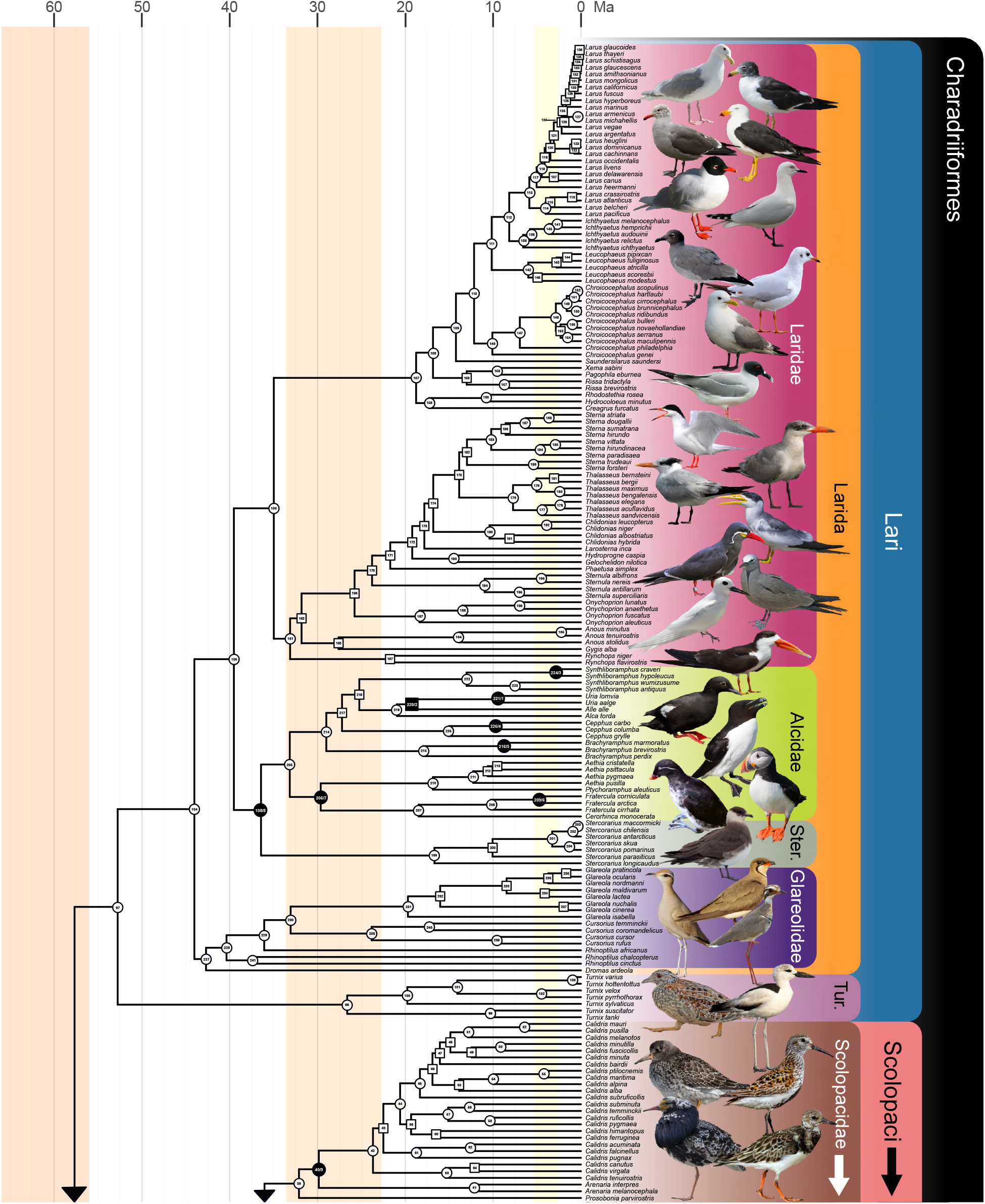
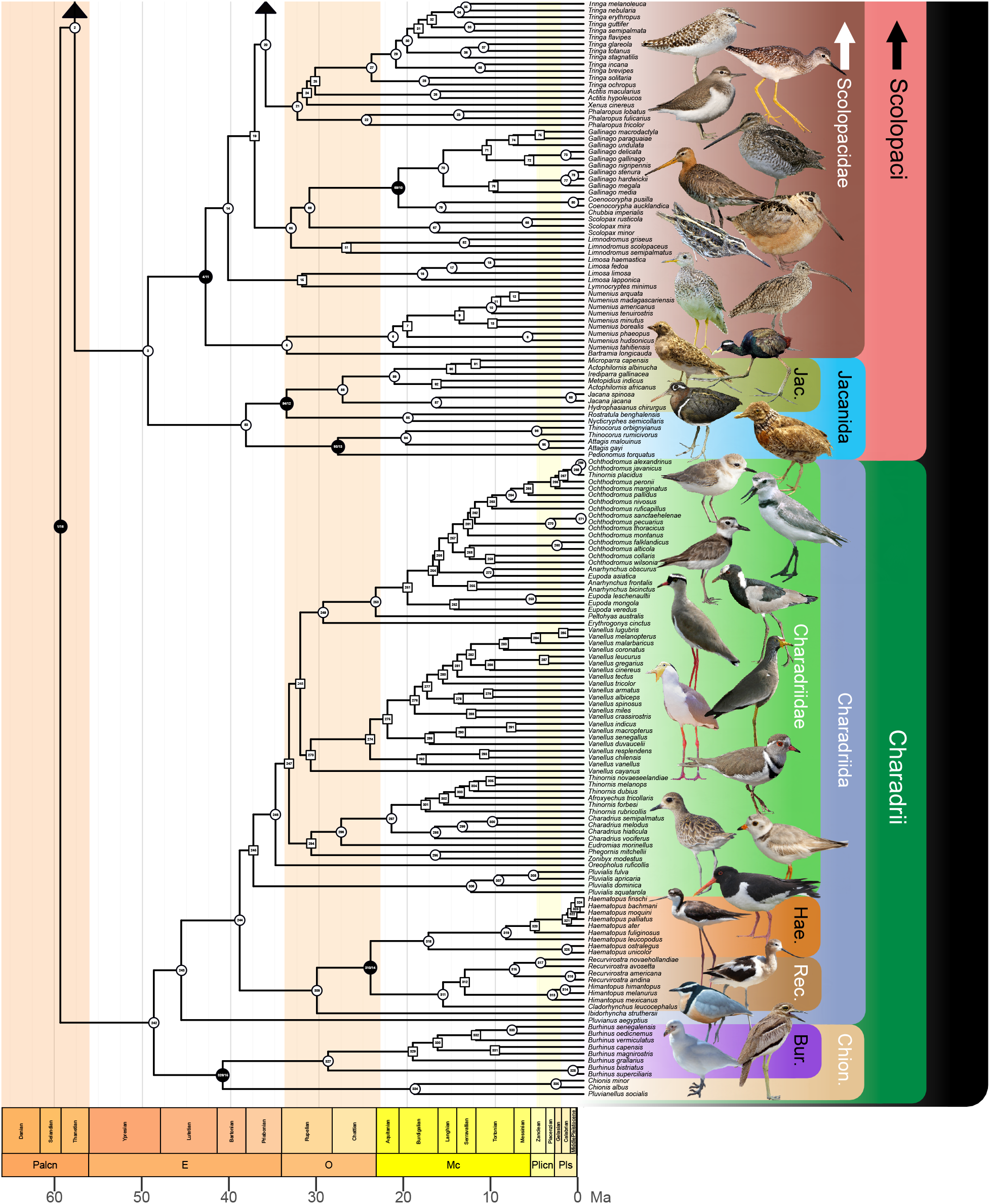
Time-calibrated phylogeny of 336 species of shorebirds based on the Bayesian node-dating analysis of 8 clock-like loci under the independent-rates relaxed clock and a fixed maximum-likelihood topology inferred using RAxML-NG from 24 genes and 69 morphological characters. The figure and this caption continue on the opposite page. Nodes with bootstrap support ≥70% are indicated by circles; nodes with bootstrap support *<*70% are indicated by squares. Fossil-calibrated nodes are shown in black; the second of the two numbers corresponds to that in Table 2. Shaded tabs represent higher-level clades; background shading indicates geochronological epochs. Ma = million years ago; Ster. = Stercorariidae; Tur. = Turnicidae; Jac. = Jacanidae; Hae. = Haematopodidae; Rec. = Recurvirostridae; Bur. = Burhinidae; Chion. = Chionida; Palcn = Paleocene; E = Eocene; O = Oligocene; Mc = Miocene; Plicn = Pliocene; Pls = Pleistocene. Representative species are illustrated next to their lineages; see Supplementary Information (Table S2) for full image credits.

**Table 3:** Likelihood support for the total-evidence (TE) and evolutionary placement algorithm (EPA) trees calculated on the combined dataset using IQ-TREE. Significance levels (*p*) and the inclusion in (+) or significant exclusion from (−) the 95% confidence set are given for the Kishino-Hasegawa (KH; Kishino and Hasegawa, 1989), Shimodaira-Hasegawa (SH; Shimodaira and Hasegawa, 1999), and approximately unbiased (AU; Shimodaira, 2002) tests; *c* (ELW) denotes confidence as measured using expected likelihood weight (Strimmer and Rambaut, 2002).

| Tree | IQ-TREE log likelihood | *p* (KH) | *p* (SH) | *p* (AU) | *c* (ELW) |
| --- | --- | --- | --- | --- | --- |
| TE | −412147.336 | 0.979 + | 1 + | 0.985 + | 0.978 + |
| EPA | −412224.722 | 0.0213 − | 0.0213 − | 0.0154 − | 0.0219 − |

The inclusion of morphological data in the TE analysis also changed the backbone relationships among the species represented by sequence data. When pruned down to the taxon sample of the concatenated tree, the TE tree differed from the former in 43 nodes (RF distance = 0.142); however, these were generally poorly supported in both the concatenated RAxML tree (average BS = 38.2%, average IC = 0.025) and in the TE tree (average BS = 34.6%). Out of the 43 nodes present in the concatenated RAxML tree but not in the TE tree, 33 were also absent from the concatenated ExaBayes tree. The TE tree and the concatenated trees agreed on all essential features of higher-level charadriiform phylogeny, including the sister-group relationship between *Ibidorhyncha* and Haematopodoidea and the monophyly of Charadriidae (Figure 5), and topological conflict was largely restricted to intrageneric relationships within species-rich genera such as *Glareola*, *Larus*, or *Ochthodromus*. Notable deviations from the concatenation-based topologies included the monophyly of the lapwings (*Vanellus*), with *V. cayanus* sister to the rest of the clade, but the support for this result was extremely weak (BS = 9%).

### 3.5 Fossil calibrations

The Bayesian re-analyses of the morphological matrix of Musser and Clarke (2020) robustly supported a charadriiform affinity for specimen IGM 100/1435 from the Paleocene/Eocene boundary of Mongolia, which predates the previous oldest known remains of the clade (Figure 1). Under all three topological constraints, IGM 100/1435 emerged as a crown-group charadriiform and specifically as a total-group member of the Chionoidea (average PP = 0.909; Supplementary Information, Fig. S10), a position also supported by some of the original, partially constrained parsimony analyses (Musser and Clarke, 2020). For the purposes of calibration design, we took the conservative approach of associating the specimen with the least inclusive clade to which it could be assigned with a PP ≥ 0.95, a condition satisfied only by the entire charadriiform crown group (Table 2). When constructing the outgroup sequences, which require taxa to be associated with branches rather than nodes, the fossil was treated as *incertae sedis* within the total group of Charadrii. The analyses also weakly (average PP = 0.504) but consistently placed SMF Av 619, another early Eocene shorebird record (Figure 1), within the total group of Larida. However, the least inclusive node to which the specimen could be assigned with a PP *≥* 0.95 was again the charadriiform crown group, rendering it redundant as a potential calibration. To account for phylogenetic uncertainty, we averaged the posterior probabilities of the membership of IGM 100/1435 within the shorebird crown across all three analyses, and used the complement (1 *−* average PP = 0.024) as the left-tail probability for the corresponding calibration density (i.e., the probability that the true age of the clade is younger than the soft minimum defined by IGM 100/1435).

### 3.6 Divergence time estimation

Under the IR model, the 2-partition analysis failed to reach the target ESS (*>* 200) and PSRF (*<* 1.01) for the ages of two of the calibrated nodes (calibrations 1 and 4; nodes 221 and 226 in Figure 5). We therefore refer primarily to the unpartitioned results, although the estimates were broadly similar between the two analyses. Figure 5 shows the posterior mean node ages obtained from the unpartitioned analysis under the IR model, supporting a mid-Paleocene origin for the charadriiform crown group at 59.3 Ma (95% posterior CI = 55.4–64.8 Ma). This result was robust to the choice of the relaxed clock model (autocorrelated rates: mean = 58.1 Ma, 95% posterior CI = 55.0–63.0 Ma) and the partitioning scheme (2-partition analysis, AR: mean = 58.3 Ma, 95% posterior CI = 55.1–63.6 Ma; IR: mean = 60.9 Ma, 95% posterior CI = 55.9–66.0 Ma), with all analyses yielding relatively narrow credibility intervals that did not extend into the Cretaceous. Almost half of the families (IR: 8 out of 19; AR: 9 out of 19) had Eocene mean stem ages, indicating an early period of steady diversification. The estimated timeline suggests that the current diversity of the three most species-rich families (Laridae: 105 species in the TiF checklist, Scolopacidae: 97 species, Charadriidae: 67 species) has been accumulated over long periods of time, as the posterior distributions of their crown ages were concentrated in the Eocene (Laridae, IR: 30.2–39.8 Ma; Scolopacidae, IR: 37.4–48.6 Ma; Charadriidae, IR: 31.5–43.3 Ma), substantially predating those of the other families with the exception of Glareolidae.

**Table 4:**
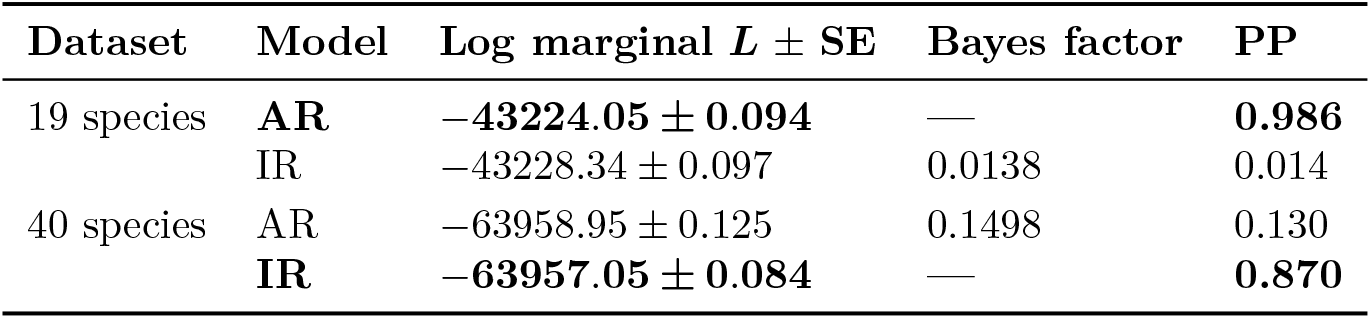
Bayes factor comparison of the relaxed clock models. AR = autocorrelated rates; IR = independent rates; Log marginal *L* ± SE = logarithm of the marginal likelihood with standard error; PP = model posterior probability assuming a flat prior over models. For each dataset, the model with the highest marginal likelihood is shown in bold; note that the Bayes factors are given relative to the preferred model.

The Bayesian clock model selection yielded ambiguous results (Table 5), favoring the AR model over the IR model in the 19-species dataset (PP = 0.986, BF = 72.6, strong evidence *sensu* Kass and Raftery, 1995) but flipping in favor of the IR model in the 40-species dataset (PP = 0.870, BF = 6.7, positive evidence). Clock model choice had little impact on the ages of the deepest nodes, but resulted in considerable differences between the date estimates for some of the more recent divergences (Figure 6). On average, the IR posterior mean ages were 1.80 Myr younger compared to the AR estimates, although the greatest difference between the two models occurred at a node that was much older under independent rates (node 241 in Figure 5: IR mean = 37.4 Ma, AR mean = 20.8 Ma). Notably, all of the 37 nodes whose posterior CIs did not overlap between the two clock models were younger under the IR model (Figure 6); these nodes were all concentrated within the gull subfamily (Larinae). The difference was even more pronounced in the partitioned analysis (IR means on average younger by 5.52 Myr; 62 nodes with non-overlapping posterior CIs all younger under the IR model). The 95% posterior CIs tended to be narrower under the IR model (average width = 8.67 Myr) than under the AR model (average width = 9.88 Myr). Compared to the IR model, outliers with particularly diffuse posterior age distributions were more prominent but mostly associated with the same nodes when the AR model was employed; the three widest 95% posterior CIs (range of widths: 27.1– 35.4 Myr) were found at nodes 197, 238, and 241 under autocorrelated rates (Figure 6), and at nodes 197, 238, and 239 under independent rates (range of widths: 21.9–23.4 Myr). Bayesian simple linear regression of the 95% posterior CI onto the posterior mean (with a suppressed intercept and an improper uniform prior on the slope) yielded lines with a slope of 0.458 for the IR model and 0.471 for the AR model (Supplementary Information, Fig. S11), indicating the amount by which the 95% posterior CI widens for every 1 Ma added to the mean age (dos Reis et al., 2018). In the 2-partition analysis, the 95% posterior CIs were wider under the AR model (average width = 12.69 Myr) but not the IR model (average width = 8.34 Myr; note that this value is not sensitive to the exclusion of the two nodes whose ages failed to converge). However, the regression lines were steeper for both models (Supplementary Information, Fig. S11), suggesting that the inclusion of a greater amount of sequence data in the unpartitioned analyses helped reduce the uncertainty associated with the divergence time estimates.

**Figure 6.**
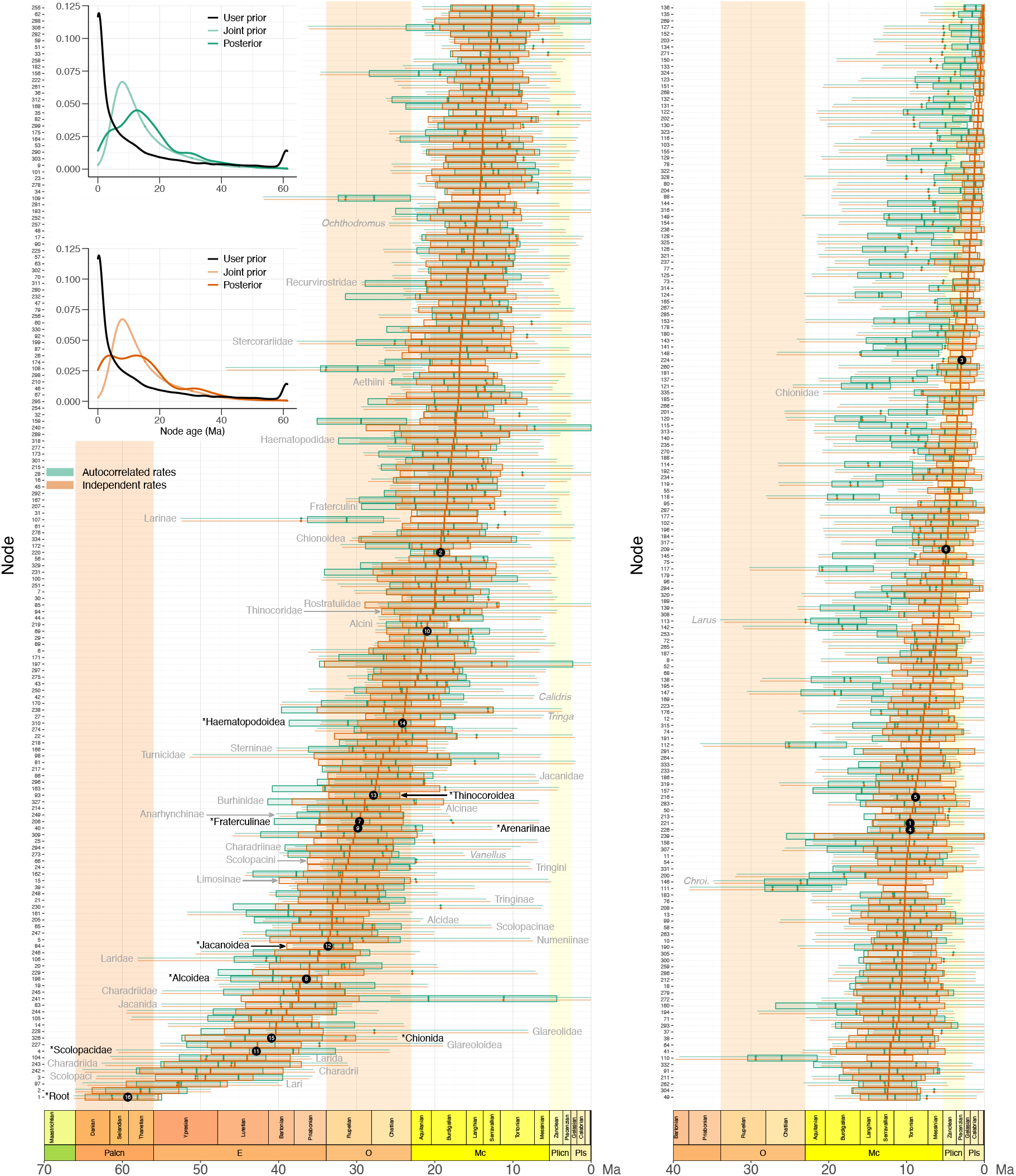
Comparison of the prior and posterior node age distributions under the AR versus IR relaxed clock models. Node numbers on the left correspond to those in Figure 5. Lines and boxes represent the 95% prior and posterior credibility intervals, respectively (with prior means indicated by diamonds and posterior means indicated by vertical lines). Calibrated nodes are highlighted by black circles; the calibration number inside each circle corresponds to that in Table 2. Clade labels are provided for suprageneric taxa and major genera; the labels for calibrated nodes are in black and denoted by asterisk, while other nodes are labeled in gray. Insets show the kernel densities of node ages under the AR versus IR models; the density for the user-specified prior was calculated from 1000 simulated trees.

**Table 5:**
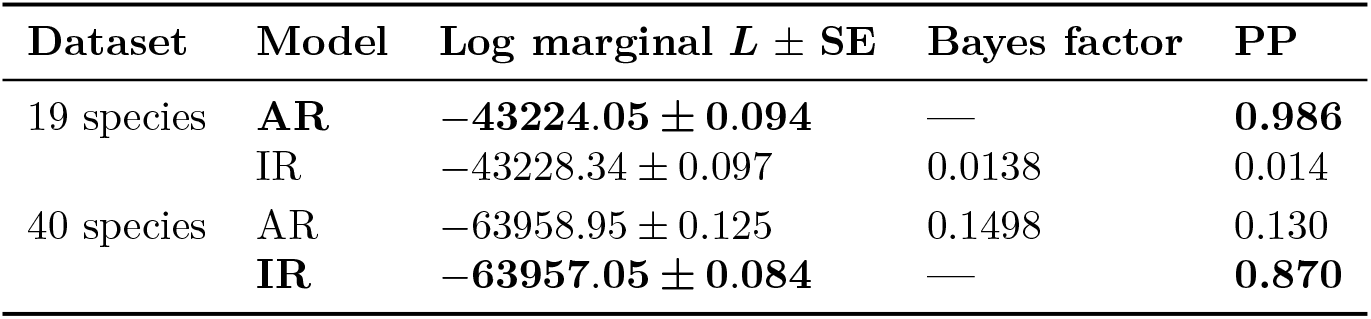
Bayes factor comparison of the relaxed clock models. AR = autocorrelated rates; IR = independent rates; Log marginal *L* ± SE = logarithm of the marginal likelihood with standard error; PP = model posterior probability assuming a flat prior over models. For each dataset, the model with the highest marginal likelihood is shown in bold; note that the Bayes factors are given relative to the preferred model.

Although MCMCTree employs conditional rather than multiplicative calibration construction, thus ensuring compatibility between the user-specified calibration densities and the birth-death prior (Yang and Rannala, 2005; Heled and Drummond, 2011), the joint (effective) prior density on the age of a node can still deviate from the specified calibration density because of the requirement that ancestral nodes be older than their descendants (dos Reis et al., 2018; Su et al., 2021). We observed this effect for several calibration densities assigned to nested nodes (Figure 7). In particular, the joint prior on the age of the sandpipers (Scolopacidae) appears to have been pushed into the past by the calibration densities assigned to two of its subclades (Arenariinae and *Gallinago* + *Coenocorypha*), with its 95% prior CI ranging from 25.8–60.6 Ma, as opposed to 20.0–48.3 Ma for the user-specified density (Table 2). However, the fact that the joint priors of the two subclades were bimodal suggests that the effect may have been bidirectional, with the ancestral node pulling the ages of the nested nodes deeper into the past (Figure 7). The same phenomenon also affected another pair of nested calibrations within Fraterculinae (6 and 7) but did not impact calibrations 8 (assigned to a clade comprising calibrations 1 through 7) and 2 (assigned to the node immediately above calibration 1), possibly due to the fact that their user-specified densities were much older than those of any of the calibrated nodes descended from them (Table 2). Similar patterns of calibration interactions were also observed under the AR model (Supplementary Information, Fig. S12).

**Figure 7.**
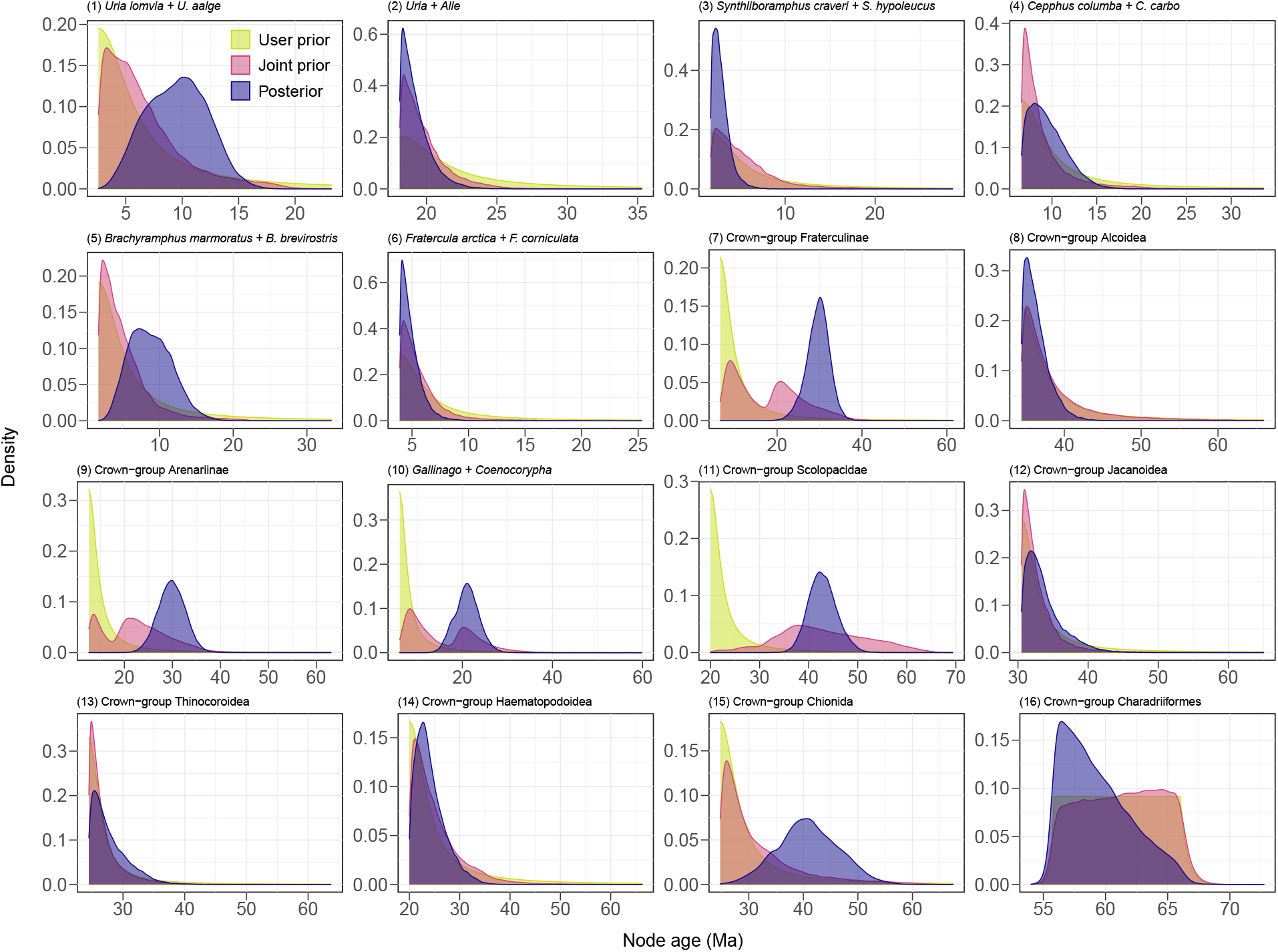
Probability density functions for the ages of the 16 calibrated nodes. “User prior” = user-specified calibration densities (truncated Cauchy for calibrations 1 through 15, soft-bounded uniform for calibration 16); “Joint prior” = effective prior resulting from calibration interactions under the IR model; “Posterior” = marginal posterior under the IR model.

Consistent with other recent studies (Chazot et al., 2019; Su et al., 2021), we found that the sequence data were able to inform the estimated dates, as indicated by the divergence of node age posteriors from the corresponding joint priors (Figure 7). This distinction was particularly notable for the root age (calibration 16). Although the joint prior placed probabilities of 0.055 (IR) or 0.057 (AR) on ages predating the K–Pg boundary (66.0 Ma), approximating the user-specified 5% fraction, the posterior probability of a pre-K–Pg origin of shorebirds only amounted to 0.0095 or 0.0035 under the IR and AR models, respectively. Conversely, the probability of a post-Paleocene origin more than doubled from 0.021 (IR) or 0.020 (AR) under the joint prior (user-specified prior probability: 0.024) to 0.044 and 0.099 under the IR and AR posteriors, with the mean and median shifting toward the present by 2.0 and 2.6 Myr (IR) or 3.3 and 4.0 Myr (AR), respectively. In agreement with recent findings (Brown and Smith, 2018), we found greater differences between the induced prior and the posterior for uncalibrated nodes than for calibrated nodes, indicating that the ability of the data to override the prior was real but limited. Relative prior-posterior displacement, defined here as |*a*_po_ *− a*_pr_ *|/*[(*a*_po_ + *a*_pr_)*/*2] (where *a*_pr_ is the prior mean and *a*_po_ is the posterior mean), was greater for the uncalibrated nodes (IR: 0.530, AR: 0.356) than for the calibrated ones (IR: 0.206, AR: 0.237). Similarly, the average shrinkage of the 95% posterior CIs relative to the 95% prior CIs was less prominent for the nodes with calibrations (IR: 37.8%, AR: 37.1%) than for those without them (IR: 57.9%, AR: 51.8%) (Figure 6).

### 3.7 Macroevolutionary rate estimation

Using the time tree inferred under the AR model, we found no evidence for clade-specific diversification regimes that significantly differed from that of the order as a whole (Figure 8a). The 95% credibility set contained a single configuration without rate shifts, and the model consisting of a single macroevolutionary regime was favored over multi-regime alternatives (PP = 0.86, BF = 3.74 relative to the next best model with one shift). Within this regime, we found strong support for a treewide diversification slowdown, with nonoverlapping 95% CIs about the mean net diversification rates at the root (0.144–0.303 sp*·*Myr*^−^*^1^, i.e., species per million years) and at the tips (0.037–0.053 sp*·*Myr*^−^*^1^).

**Figure 8.**
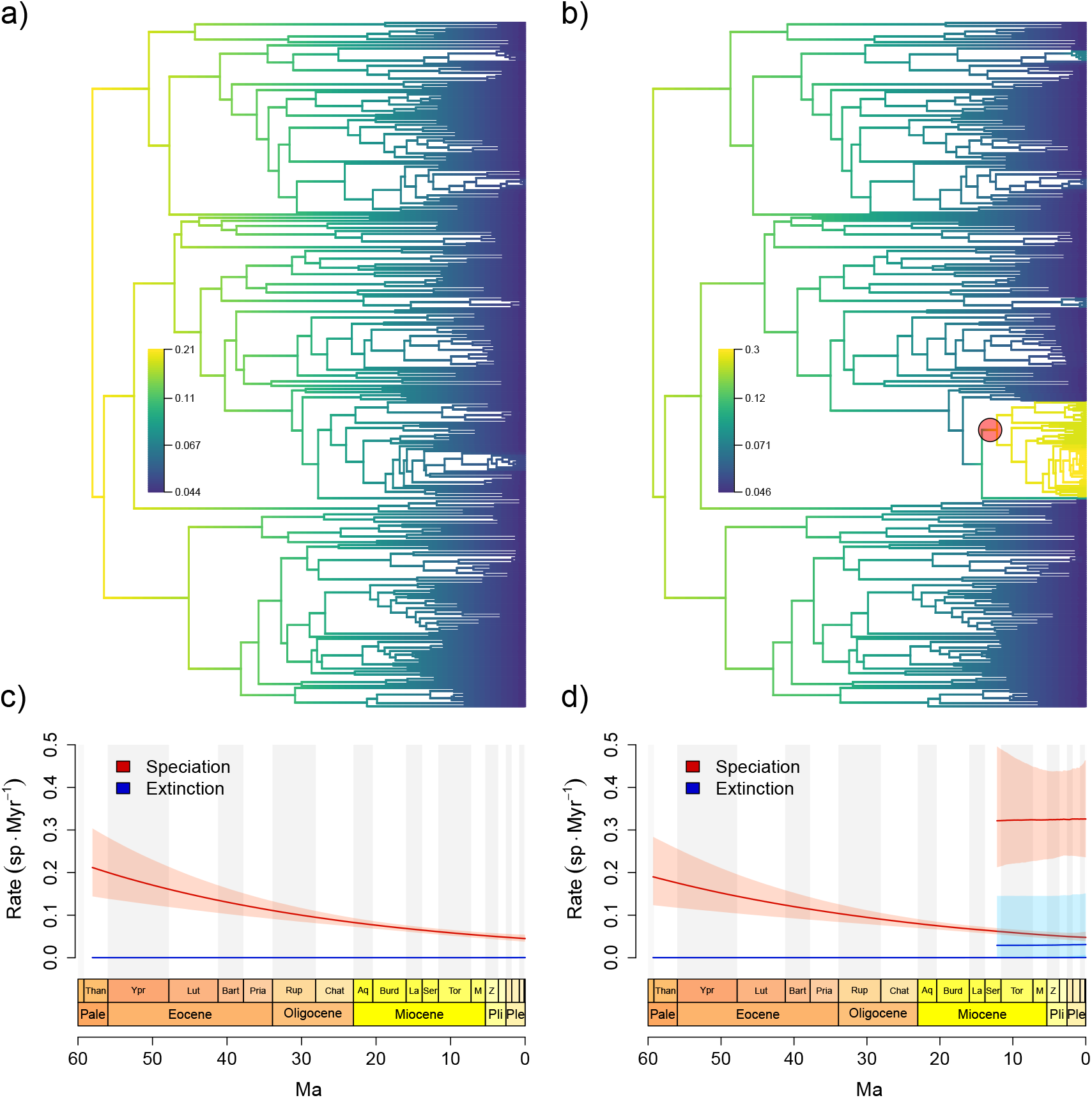
Top: phylorate plots for the AR (a) and IR (b) trees with branches colored by the net diversification rate (in sp Myr*^−^*^1^) and rate shifts denoted by red circles. Bottom: corresponding rate-through-time (RTT) plots showing the mean speciation and extinction rates with their associated 95% credibility intervals (c, d). The RTT plot for the IR tree (d) shows the rates for both the background regime and the shifted regime corresponding to the gull clade highlighted in (b).

In contrast, when estimating diversification dynamics from the tree based on the IR relaxed clock, the best supported model included a rate shift associated with the clade comprising the gull genera *Larus*, *Ichthyaetus*, *Leucophaeus*, and *Chroicocephalus* (node 110 in Figure 5). The shift subtending this node, or a shift subtending the node immediately above it (node 109), consistently appeared in all the 22 configurations included in the 95% credibility set, and represented the only shift present in the MAP configuration (PP = 0.67; Figure 8b). The model-averaged time-weighted mean rate of the (*Larus* + *Chroicocephalus* clade exceeded that of the background regime by a factor of 5.0 for speciation and 4.5 for net diversification. The elevated net diversification rates estimated for the gull clade were nearly constant through time (root 95% CI: 0.168–0.445 sp Myr*^−^*^1^, tip 95% CI: 0.187–0.404 sp Myr*^−^*^1^), whereas the background regime again exhibited declining diversification (root 95% CI: 0.123–0.283 sp Myr*^−^*^1^, tip 95% CI: 0.038–0.058 sp Myr*^−^*^1^) (Figure 8d). Configurations containing additional shifts were also sampled relatively frequently on the IR tree, and enjoyed non-negligible support. The cumulative posterior probability of models involving 2–7 shifts amounted to 0.34, and the Bayes factor of the single-shift model compared to the next best model with 2 shifts was 1.36 (“barely worth mentioning” following the criteria of Kass and Raftery, 1995). In addition to the upshift associated with the gulls, other frequently sampled shifts occurred within the genera *Haematopus* (nodes 319–321 in Figure 5; present in 6 out of the top 22 configurations with a cumulative PP of 0.065) and *Stercorarius* (nodes 201 or 202; present in 4 out of the top 22 configurations with a cumulative PP of 0.035). All of the rate shifts present in the 95% credible set of configurations represented accelerations (upshifts).

## 4 Discussion

### 4.1 Congruence and conflict in higher-level charadriiform relationships

The topologies inferred by our concatenated, species-tree, and total-evidence analyses are broadly consistent with previous estimates based on molecular data (Ericson et al., 2003; Paton et al., 2003; Paton and Baker, 2006; Baker et al., 2007; Fain and Houde, 2007; Hackett et al., 2008; Prum et al., 2015; Hu et al., 2017). Our results support the three-suborder division of the Charadriiformes into Charadrii, Scolopaci, and Lari; the position of the morphologically aberrant buttonquails (Turnicidae) as the earliest-diverging lineage within the Lari; and most of the previously proposed interfamilial relationships (Baker et al., 2007; Burleigh et al., 2015; Prum et al., 2015). In conjunction with previous work, these findings indicate that most of the higher-order charadriiform relationships are robustly resolved. However, several localized areas of uncertainty persist despite the comprehensive taxon sampling employed here.

Most notably, our ASTRAL species tree contradicts both previous phylogenies of the clade (Baker et al., 2007; Pereira and Baker, 2010; Burleigh et al., 2015; De Pietri et al., 2020) and our own concatenated analyses in placing the crab plover (*Dromas ardeola*) closer to the auks and gulls than to the pratincoles and coursers, and in finding the Egyptian plover (*Pluvianus aegyptius*) outside of a group comprising the Chionida and the plover–oystercatcher assemblage, as opposed to allying it with the latter clade (Figure 4). Although close examination of the branches in question revealed the presence of better-supported alternatives consistent with the concatenation-based results, constrained searches failed to improve on the quartet score of the unconstrained estimate. We hypothesize that the relationships involving the two taxa interacted with other regions of the tree in such a manner that their unconventional positions increased global quartet support despite decreasing it locally. This phenomenon was recently noted by Rabiee and Mirarab (2020), who showed that constrained ASTRAL searches can reveal well-supported clades absent from the main ASTRAL tree at the cost of decreasing the overall quartet score. As a result, we do not consider the relationships of *Dromas* and *Pluvianus* to represent genuine instances of gene tree/species tree discordance, but their different resolutions in the species-tree and concatenated analyses may indicate the presence of such conflict in neighboring regions of the tree.

We found weak but consistent support for the monophyly of typical plovers (Charadriidae), as opposed to previous studies that have suggested their paraphyly with respect to Haematopodoidea and *Ibidorhyncha* (Ericson et al., 2003; Baker et al., 2007; Fain and Houde, 2007; Burleigh et al., 2015; but see Baker et al., 2012; Hu et al., 2017). The consistent grouping of the golden and gray plovers (*Pluvialis*) with the rest of the Charadriidae in our species-tree as well as concatenated analyses suggests that plover paraphyly is not an artifact of gene tree/species tree discordance, in agreement with Baker et al. (2012). Instead, our analyses support the interpretation of this result as a stochastic error stemming from the use of a small number of loci that happen to incorrectly resolve the short branch connecting *Pluvialis* to the rest of the plovers. Contrary to the suggestion of Baker et al. (2012) that mitochondrial loci may have been especially prone to inferring the incorrect relationship due to their high mutational variance, we observed no difference between nuclear and mitogenomic data in this regard. In fact, the nuclear loci used in this study mostly favored paraphyly (5 out of the 8 applicable loci), which was supported by the gene trees for RAG1 (also used by Ericson et al., 2003 and Baker et al., 2007), MB2 (used by Ericson et al., 2003), as well as ADNH, FGB7, and GAPDH (all three also used by Fain and Houde, 2007). In contrast, mitochondrial loci narrowly supported charadriid monophyly (8 out of the 15 loci), as did the mitochondrial genome as a whole when analyzed as a single unit. This finding is consistent with the previous observation of Chen et al. (2018) that increased locus sampling tends to overturn plover paraphyly in mtDNA analyses (see also Hu et al., 2017). In conjunction with previous work, our results suggest that plover monophyly is supported by both nuclear and mitochondrial genomes (Baker et al., 2012; Hu et al., 2017; Chen et al., 2018), and underline the need for extensive gene sampling to eliminate stochastic error.

Our study helps indicate directions for future research by identifying regions of the shorebird tree that could not be confidently resolved using the relatively small number of loci employed here. One such area of uncertainty represents the interrelationships of the five major lineages comprising the Laridae (often classified as separate subfamilies; Cracraft, 2013; Boyd, 2019): the gulls (Larinae), true terns (Sterninae), skimmers (*Rynchops*), noddies (*Anous*), and white terns (*Gygis*), whose relationships have so far proved elusive (Chu, 1998; Paton and Baker, 2006; Baker et al., 2007; Fain and Houde, 2007; Burleigh et al., 2015; Hu et al., 2017). This uncertainty is expected given the combination of very short internodes connecting the five taxa and the relatively long branches subtending them, a feature characteristic of ancient rapid radiations (Lanyon, 1988; Whitfield and Lockhart, 2007). However, our node-dating analyses show that the corresponding divergences unfolded over a period of approximately 6–7 Myr, suggesting they were not so rapid as to be intractable. As a result, longer alignments relying on data from ultraconserved elements (Faircloth et al., 2012), exon capture (Bragg et al., 2016), or whole-genome sequencing (Jarvis et al., 2014; Zhang et al., 2014) may be more successful at resolving the larid radiation. The same is likely to be true of the interrelationships of the six extant true auk genera (*Alca*, *Alle*, *Uria*, *Synthliboramphus*, *Brachyramphus*, and *Cepphus*), where disagreements occur both among earlier studies (Baker et al., 2007; Smith and Clarke, 2015) and among the different analytical approaches employed here (Figure 4). Finally, topological conflict is also rampant among the sandpipers (Scolopacidae), especially with regard to the position of the jacksnipe (*Lymnocryptes*), the monophyly of the dowitchers (*Limnodromus*), the membership of the Polynesian sandpipers (*Prosobonia*) in the Arenariinae *sensu* Banks (2012) (*Arenaria* + *Calidris*), and the exact sequence of divergences within the Tringinae (Figure 4). While the last two of these relationships are again associated with extremely short internodes and have been uncertain in previous studies (Baker et al., 2007; Cibois et al., 2012; Gibson and Baker, 2012; De Pietri et al., 2020), the differences between the species-tree and concatenated analyses detected for *Limnodromus* and *Lymnocryptes* hint at the presence of other problems, such as gene tree estimation error or genuine gene tree/species tree discordance.

Despite maximizing taxon sampling based on the available molecular and morphological data, our study still contains gaps in taxonomic coverage that suggest where additional sampling effort should be directed. Of the 13 non-monotypic charadriiform families, the buttonquails (Turnicidae) had the lowest coverage in the present study (7 out of 17 species), and currently available sequence data do not even make it possible to test whether *Turnix* is monophyletic with respect to the monotypic *Ortyxelos*. We suggest that sequencing the latter genus as well as additional *Turnix* species should be a priority for future studies seeking to improve taxon sampling within the Charadriiformes.

### 4.2 Taxonomic implications

While the higher-level phylogeny of the Charadriiformes is generally well-established, the comprehensive taxon sampling of our study helps reveal instances of nonmonophyly at the genus level. Despite the low support values often associated with shallower relationships in our trees, several of these results are robust enough to motivate changes to the current taxonomies of the clade. Our findings lend support to classifying the double-banded courser as the sole member of the genus *Smutsornis*, following Clements et al. (2019), in contrast to other recent treatments that consider it a species of *Rhinoptilus* (*“R.” africanus*; Dickinson and Remsen, 2013; Boyd, 2019; De Pietri et al., 2020). Although at least one molecular phylogeny showed the species to be nested within a monophyletic *Rhinoptilus* (Cohen, 2011), our concatenated and total-evidence analyses find moderate to strong support (RAxML BS/IC = 87%/0.564, ExaBayes PP = 0.73, TE BS = 85%) for a node uniting it with *Cursorius* and *Glareola* instead, in agreement with Burleigh et al. (2015) and with earlier observations noting its morphological and behavioral differences from the *Rhinoptilus* coursers (del Hoyo and Collar, 2014).

Our analyses yield a scolopacid topology that is highly similar to the phylogeny of Gibson and Baker (2012), as expected given their reliance on much of the same sequence data. Accordingly, our findings are consistent with the changes implemented by recent taxonomies in response to the latter study, including the expansion of the genera *Calidris* and *Tringa* to ensure their monophyly (Dickinson and Remsen, 2013; Boyd, 2019; Clements et al., 2019), and the reassignment of the imperial snipe to a separate genus (*Chubbia*) to preserve the monophyly of *Gallinago* (Boyd, 2019). In contrast, the narrow concept of the genus *Charadrius* and the reassignment of most of its former species to the resurrected genera *Afroxyechus*, *Eupoda*, and *Ochthodromus*, while adopted by the taxonomy employed here (Boyd, 2019) to reflect recent phylogenetic findings (Barth et al., 2013; Dos Remedios et al., 2015), does not fit our results substantially better than alternative taxonomies (Dickinson and Remsen, 2013; Clements et al., 2019).

We corroborate previous findings (Barth et al., 2013; Burleigh et al., 2015; Dos Remedios et al., 2015) showing that most of the species traditionally assigned to *Charadrius* are more closely related to *Anarhynchus*, *Erythrogonys*, and the Vanellinae than to the type (*C. hiaticula*), thus violating the monophyly of the genus and of the subfamily Charadriinae as traditionally defined (Cracraft, 2013). We also find *Anarhynchus* to be deeply nested within the clade consisting of most of the former *Charadrius* species (“CRD II” *sensu* Dos Remedios et al., 2015; Anarhynchinae *sensu* Boyd, 2019), although its exact position with respect to the latter differs here from the earlier analyses despite no appreciable differences in taxon sampling. Our results support the reassignment of “*Charadrius*” *bicinctus* to *Anarhynchus* in light of their sister-group relationship (ASTRAL localPP = 0.44, RAxML BS/IC = 53%/0.065, ExaBayes PP = 0.56, TE BS = 49%), and show a moderately well-supported (RAxML BS/IC = 71%/0.327, ExaBayes PP = 0.98, TE BS = 62%) early-diverging clade consisting of the greater sand plover (“*C.*” *leschenaultii*), oriental plover (“*C.*” *veredus*), and mountain plover (“*C.*” *montanus*), for which the name *Eupoda* was resurrected by Boyd (2019). However, instead of allying the Caspian plover (“*C.*” *asiaticus*) with the rest of the genus *Eupoda* (of which it is the type species) and the New Zealand plover (“*C.*” *obscurus*) with *Anarhynchus*, as in Barth et al. (2013) and Dos Remedios et al. (2015), our concatenated and total-evidence analyses show both taxa to form a robust clade (RAxML BS/IC = 80%/0.305, ExaBayes PP = 1, TE BS = 82%) that is more closely related to *Ochthodromus* than to either *Anarhynchus* or *Eupoda* (RAxML, TE), or even deeply nested within *Ochthodromus* (ExaBayes). Interestingly, the ASTRAL species tree does recover *Eupoda* and *Anarhynchus sensu* Boyd (2019) as monophyletic with localPP values of 0.97 and 0.31, respectively. To safeguard generic monophyly under both of the competing hypotheses, we suggest combining the genera *Anarhynchus*, *Eupoda*, and *Ochthodromus*, in which case *Anarhynchus* Quoy and Gaimard 1830 takes the priority. The subfamily Anarhynchinae would then comprise the genera *Anarhynchus*, *Erythrogonys*, and *Peltohyas*, which are universally recognized by major taxonomies (Dickinson and Remsen, 2013; Boyd, 2019; Clements et al., 2019).

The second major assemblage of the former *Charadrius* species (“CRD I” *sensu* Dos Remedios et al., 2015; Charadriinae *sensu* Boyd, 2019) is rendered nonmonophyletic by the inclusion of the genera *Thinornis* and *Afroxyechus*. Like Barth et al. (2013) and Dos Remedios et al. (2015), we find that the former genus is itself nonmonophyletic, as its two species (*T. novaeseelandiae* and *T. rubricollis*) span a clade that also includes two plovers usually assigned to *Charadrius* (“*C.*” *dubius* and “*C.*” *forbesi*) as well as the black-fronted dotterel, occasionally placed in its own separate genus (“*Elseyornis*” *melanops*). These results therefore support the expansion of *Thinornis* following Boyd (2019). However, we find the three-banded plover (“*Afroxyechus*” *tricollaris*) to be nested within the expanded *Thinornis* as well, contrasting with the results of Dos Remedios et al. (2015) who found it outside of the (*Charadrius* + *Thinornis*) clade. The latter result was extremely weakly supported (PP = 0.47) and only appeared in the concatenated analysis despite its absence from any of the individual gene trees, whereas we find strong support for the inclusion of “*A.*” *tricollaris* within *Thinornis* (ASTRAL localPP = 0.85, RAxML BS/IC = 78%/0.581, ExaBayes PP = 1, TE BS = 63%), a position also favored by 5 out of the 6 gene trees of Dos Remedios et al. (2015). We therefore suggest that the species be reassigned to this genus as a new combination, *Thinornis tricollaris*. More surprisingly, another species formerly recovered within and reassigned to *Thinornis*, the long-billed plover (“*T.*” *placidus*; see Dos Remedios et al., 2015; Boyd, 2019), is found here to be deeply nested within *Ochthodromus* (*Anarhynchus* in our preferred taxonomy), possibly as a result of differences in the sampling of the mitochondrial genome (4 genes used by Dos Remedios et al., 2015 vs. 15 genes used here). We refrain from making taxonomic recommendations for this species pending results from analyses with broader locus sampling.

Our final recommendation concerns the pied lapwing (“*Vanellus*” *cayanus*), whose affinity to other lapwings (*Vanellus*) remains doubtful (Figure 4). Of the analyses performed here, only the total-evidence tree supported its inclusion within the genus, but with virtually no bootstrap support (9%), and only as the sister species to all other members of the genus. In contrast, other analyses variously placed the pied lapwing in a sister-group relationship with *Oreopholus* (ASTRAL; localPP = 0.67), all charadriids other than *Oreopholus* and *Pluvialis* (RAxML; BS/IC = 36%/0.011), or the (Vanellinae + Anarhynchinae) clade (ExaBayes; PP = 0.29). To account for this range of hypotheses, we propose to resurrect the genus *Hoploxypterus* Bonaparte 1856 for the species. The sub-family Vanellinae, currently redundant in most taxonomies with respect to *Vanellus*, could then represent a useful name for the (*Vanellus* + *Hoploxypterus*) group, should it prove to be monophyletic.

### 4.3 A new timeline for charadriiform evolution

The mid-Paleocene origin of the charadriiform crown inferred here is consistent with an explosive radiation of neoavian lineages in the wake of the end-Cretaceous mass extinction (Ericson et al., 2006; Suh, 2016; Berv and Field, 2017), and (unlike the early mitogenomic timescales; Paton et al. 2002; Pereira and Baker 2006; Baker et al. 2007) does not require positing a long period during which the clade supposedly diversified but failed to leave behind a fossil record. The near-complete absence of support for a Cretaceous origin of shorebirds (PP *<*0.01) in this study is remarkable, since our root calibration was designed to allow for this possibility (Table 2), and consequently saw one of the greatest shifts between the effective prior and the posterior (Figure 7). At the same time, our age estimates for the charadriiform root (IR mean: 59.3 Ma, AR mean: 58.1 Ma) are older than the early Eocene dates favored by many recent studies (Smith, 2011: 53.6 Ma; Claramunt and Cracraft, 2015: 53.5 Ma; Prum et al., 2015: 48.8–50.6 Ma; Smith and Clarke, 2015: 49.3 Ma; cf. Figure 1). Given the recent evidence for the crown-charadriiform affinities of fossil specimens from the early Eocene of Virginia (SMF Av 619; Mayr, 2016) and the Paleocene–Eocene boundary of Mongolia (IGM 100/1435; Hood et al., 2019), presented by Musser and Clarke (2020) and corroborated by our constrained Bayesian re-analyses of their dataset, these earlier estimates must now be viewed as contradicting the known fossil record. The fact that the fossil evidence most relevant to the age of the charadriiform root only emerged one year before the completion of this study (Figure 1) illustrates the fast pace at which calibrations continue to be superseded and replaced, and serves as a reminder of the need to use up-to-date fossil information in calibration design (Ksepka et al., 2015; Marjanovíc, 2021).

Our timeline for shorebird evolution is compatible with the positions of both SMF Av 619 and IGM 100/1435 within the charadriiform crown, despite the fact that their inclusion therein was not strictly enforced: our soft-bounded root age prior placed nonzero probability on ages younger than those of the two fossils, and this probability increased after analyzing the sequence data (section 3.6). However, while our re-analyses of the Musser and Clarke (2020) dataset found weak but consistent support for the placement of the two fossils within the total groups of Larida and Chionoidea, respectively, our divergence time estimates preclude their inclusion in these clades. Our point estimates for the Larida/Turnicidae divergence are very slightly younger (IR mean: 52.8 Ma, AR mean: 52.5 Ma) than SMF Av 619 (54.17–53.7 Ma based on calcareous nannoplankton zonation; Anthonissen and Ogg, 2012; Mayr, 2016), whose age is nevertheless still included in the corresponding 95% credibility intervals. Our dating of the Chionoidea/Burhinidae divergence (IR mean: 40.9 Ma, AR mean: 44.3 Ma) is substantially younger than the estimated age of IGM 100/1435 (~55.88 Ma; see Supplementary Information), which was even excluded from the relevant 95% CIs under both relaxed clock models. Should more evidence emerge for a deeply nested phylogenetic position of early Eocene fossils, the timescale presented here would have to be altered by shifting the root age even deeper into the past, and/or by positing a more rapid succession of the interfamilial divergences.

In addition to the origin of the order, our timescale also diverges from recent phylogenomic studies with respect to the age of individual charadriiform subclades. Compared to the pseudoposterior of Jetz et al. (2012), our posterior mean ages inferred under the IR model were so young as to be excluded from the 95% pseudoposterior CIs for Chionida (40.9 Ma vs. 41.9–66.1 Ma), Charadriidae (37.4 Ma vs. 37.5–59.2 Ma), Thinocoroidea (27.9 Ma vs. 28.9–48.7 Ma), Jacanoidea (33.7 Ma vs. 33.8–54.2 Ma), and Rostratulidae (20.0 Ma vs. 22.9–40.5 Ma); for the last three clades, this was also true under the AR model, in addition to Jacanida (35.3 Ma vs. 37.0–58.1 Ma) and Jacanidae (24.7 Ma vs. 27.0–45.1 Ma). For Larida and its constituent families (Alcidae, Glareolidae, Laridae, Stercorariidae) and superfamilies (Alcoidea, Glareoloidea), the difference was even more pronounced but opposite in direction, as we found all of these clades to be significantly older than suggested by the Jetz et al. (2012) time tree distribution. Regardless of the clock model used, not only their means but also their entire 95% CIs fell outside of those derived from the Jetz et al. (2012) pseudoposterior, which produced mean ages for the Alcidae and the Stercorariidae that were less than half as old as the IR posterior means yielded by the present study (16.0 vs 33.2 Ma and 7.4 vs 16.7 Ma, respectively). Our estimates for the ages of Alcoidea, the (Alcoidea + Laridae) clade, and Larida also exceed those of other recent analyses (Claramunt and Cracraft, 2015; Prum et al., 2015; Kuhl et al., 2020), although they are not as old as suggested by early studies that relied on obsolete calibrations (Pereira and Baker, 2008).

The difference can be largely explained by our use of calibration 8 (Table 2), representing a late Eocene pan-alcid of uncertain affinities whose position within the clade was nevertheless supported by a formal phylogenetic analysis (Smith, 2011). The post-Eocene dates suggested for the Alcidae/Stercorariidae divergence (or even more inclusive clades) by recent studies therefore exemplify the “zombie lineage” problem described by Springer et al. (2017), in which molecular divergence times turn out to be younger than the known fossil record allows. The same underestimation of divergence times was recently reported for the Gruiformes by Musser et al. (2019), and may be relatively widespread as a result of efforts to correct for the implausibly old divergences yielded by earlier studies by means of overly stringent calibration choice. Conversely, the criteria employed here that allowed calibration 8 to be used could be criticized as too lax given the fragmentary nature of the material and the long temporal gap separating it from the next oldest pan-alcid occurrence. Ultimately, this problem may only be resolved by total-evidence tip-dating analyses (Ronquist et al., 2012a; Heath et al., 2014), which co-estimate tree topology and divergence times for extant and fossil taxa alike while allowing the phylogenetic position of the fossils to be informed by both their morphology and their stratigraphic age. Such analyses unfortunately remain computationally prohibitive for datasets as taxon-rich as ours. In the absence of a viable alternative to node-dating, the question of the age of Alcoidea illustrates the overwhelming influence of calibration choice on the outcomes of divergence time estimation – a statistical problem that is not fully identifiable without fossil data (Rannala, 2016).

Our ability to infer the timescale of charadriiform evolution with confidence is limited by the discordance between the results based on two different relaxed clock models, and by the lack of unambiguous support for one model over the other. Unfortunately, Bayes factor model comparisons cannot be conducted on phylogenies with hundreds of tips, since the likelihood approximation that makes MCMCTree analyses tractable for trees of such size is not valid for parameter values far removed from the likelihood peak, which are frequently visited in the course of marginal likelihood estimation (dos Reis and Yang, 2011; dos Reis et al., 2018). As a result, model comparisons have to be restricted to subsets of the full tree that are small enough for exact likelihood calculation to remain feasible. While previous studies that made use of such restricted analyses obtained consistent results under different subsampling schemes (dos Reis et al., 2018; McGowen et al., 2019), we find the preferred model to differ between the 19-species and 40-species schemes employed here (Table 5). Moreover, the direction of the change in model preference is unexpected, since the signal for rate autocorrelation should be more difficult to detect in sparsely sampled trees with long periods of time separating individual branching events (Drummond et al., 2006; Brown and van Tuinen, 2011). Here, it was the sparser 19-species dataset that favored rate autocorrelation over the independent-rates model. The resulting incongruence is substantial, as the autocorrelated-rates model estimates much older ages for many of the shallower nodes (Figure 6), raising concerns about the effect of this difference on downstream inferences (see below). Without more efficient marginal likelihood estimators to help choose between competing relaxed clock models, the more recent half of the shorebird evolutionary timeline remains subject to considerable uncertainty.

Recent node-dating studies have discouraged the use of truncated Cauchy calibration densities, finding them to be more prone to truncation and consequent deviation from the user-specified prior than simple uniform densities (dos Reis et al., 2018; Su et al., 2021). Additionally, they have been criticized for being overly informative to the extent that the sequence data may not be able to meaningfully update them (Su et al., 2021), or for being so heavy-tailed as to allow calibration interactions to pull the corresponding joint prior too deep into the past (dos Reis et al., 2018). Here, we have only observed these effects on a limited scale. The user-specified prior was generally close to the joint prior, and both often (though not always) deviated from the posterior (Figure 7), in both cases exhibiting the behavior one would ideally expect from a calibrated analysis. Nevertheless, instances of the effective prior being pulled deeper into the past were observed for calibrations 7 and 9–11 (Figure 7), an effect likely attributable to interactions between the user-specified densities assigned to nested clades. For approximately half of the calibrated nodes, there was little movement between the joint prior and the posterior, although this failure to update the effective prior was not as pervasive as recently reported (Brown and Smith, 2018). Moreover, the small number of loci employed here likely did not exhaust the ability of molecular data to update node age priors. Plotting 95% CI width against posterior mean node ages (“infinite-sites plots” *sensu* Rannala and Yang, 2007) reveals considerable scatter about the regression line (Supplementary Information, Fig. S11), indicating that uncertainty in the estimated divergence times cannot be attributed solely to the fossil calibrations, but includes sequence-data sampling error as well (Rannala and Yang, 2007; Inoue et al., 2009; dos Reis et al., 2018; McGowen et al., 2019). We therefore expect that the use of longer alignments will improve not only topological inference within the Charadriiformes, but also the precision of their estimated divergence times.

### 4.4 Tempo and mode of shorebird diversification

Our BAMM analyses of shorebird macroevolutionary dynamics yielded drastically different results depending on the time tree used (Figure 8). The inference based on the independent-rates tree showed that a clade comprising 4 genera of gulls (including a total of 48 species; Boyd, 2019) entered a new diversification regime characterized by accelerated rates of speciation and extinction that have not appreciably declined since the clade’s origin. This scenario is consistent with the findings of Jetz et al. (2012), who identified a gull clade of similar composition and size (44 species) as the single fastest-diversifying group of extant birds. Their estimate of the net diversification rate was even higher (0.74 vs. 0.29 sp Myr*^−^*^1^) as a result of a younger origin inferred for the clade (4.6 vs 12.2 Ma). In contrast, the BAMM analysis based on the autocorrelated-rates tree found little evidence for clade-specific diversification regimes, or any notable speed-up within the gulls (Figure 8a).

A close examination shows that this lack of congruence is due entirely to the drastically different divergence times estimated for the gull radiation by the two relaxed clocks. Both the IR and AR models infer similar ages for Laridae as a whole (IR mean: 35.0 Ma, AR mean: 40.3 Ma), but disagree on the ages of the gulls (Larinae; IR mean: 18.8 Ma, AR mean: 31.3 Ma), the shifted node (*Chroicocephalus* + *Larus*; IR mean: 12.2 Ma, AR mean: 26.1 Ma), and most of the individual larine genera, including the species-rich *Larus* (IR mean: 5.9 Ma, AR mean: 18.7 Ma). In all these cases, the 95% posterior CIs failed to overlap under the two clock models (Figure 6). This conflict appears to follow directly from the different assumptions made by the two models about the distribution of branch rates across the tree. The AR model, which disfavors sudden shifts, prefers to assign similar rates to the parent and daughter branches, and consequently ended up distributing divergence times evenly between the origin of Laridae and the present (Supplementary Information, Fig. S13 and Table S1). In contrast, under the IR model, each branch rate represents an independent draw from a lognormal distribution, so that the highest-probability rates can differ drastically even between the parent and daughter branches if required by the data (Drummond et al., 2006). The strong impact of clock model choice on the estimated macroevolutionary rates contrasts with previous simulation-based research finding the latter to be largely unaffected by the choice between strict and independent-rates relaxed clocks (Sarver et al., 2019), and suggests that the sensitivity of diversification rate analyses to the assumptions made in time tree inference should be widely explored in empirical systems.

## 5 Conclusions

The densely sampled, time-calibrated, total-evidence tree of shorebirds presented here is a major step toward understanding the evolution of one of the most ecomorphologically diverse and species-rich clades of non-passerine birds. It represents a substantial advance over earlier studies that were not consistent with the fossil record of the group, did not include as many species, or only did so without informing their placement by character data. We expect that the availability of a generally well-resolved phylogeny accounting for nearly nine tenths of the extant diversity of the clade will greatly facilitate future comparative, biogeographical, and macroecological studies of shorebirds. In addition to highlighting areas of robust support, which span nearly the entire suprafamilial backbone of the charadriiform tree, we also identify regions of persisting uncertainty to be prioritized by future analyses with increased taxon and locus sampling. Our study demonstrates the importance of new fossil evidence for inferring evolutionary timescales, and serves as a cautionary note about the impact of modeling choices on downstream inferences, with implications for the Charadriiformes and beyond.

## Funding

The authors received no funding for this work.

## Acknowledgments

We are grateful to Mario dos Reis, Siavash Mirarab, and Alexis Stamatakis for their help with the methods and software used in this study. We further thank the wildlife photographers whose work features in this paper for making their art available under Creative Commons licenses, Simon Ho for providing us with the time trees used to generate Figure 1, and Susan Kidwell for sharing stratigraphic information relevant to calibration design. Early drafts of the manuscript were improved by comments from John Bates and Graham Slater. Marginal likelihood estimation was performed on the Midway2 Research Computing Cluster at the University of Chicago.

## Competing interests

There are no competing interests to declare.

## Data accessibility

All GenBank accession numbers, alignments, tree files, configuration files, and R scripts are available from the Dryad Digital Repository: http://dx.doi.org/10.5061/dryad. nnnnnnnnn.

## Notes

### Competing Interest Statement

The authors have declared no competing interest.

