## Supplementary Material for "Comprehensive taxon sampling and vetted fossils help clarify the time tree of shorebirds (Aves, Charadriiformes)"

**3 Supplementary Methods** **72**

**4 Supplementary Figures and Tables** **74**

**5 Image Credits** **91**

**References** **99**

### 1 Fossil Calibrations

#### 1.1 Calibrations used

**Calibration 1**

**Node calibrated.** MRCA of *Uria aalge* and *Uria lomvia.*

**Fossil taxon.** *Uria lomvia* (Linnaeus, 1758).

**Specimen.** CASG 71892 (referred specimen; Olson, 2013), California Academy of Sciences, San Francisco, CA, USA.

**Lower bound.** 2.58 Ma.

**Phylogenetic justification.** As in Smith (2015).

**Age justification.** The status of CASG 71892 as the oldest known record of either of the two spp. of Uria was recently confirmed by the review of Watanabe et al. (2016). The younger of the two marine transgressions at the Tolstoi Point corresponds to the Bigbendian transgression (Olson, 2013), which contains the Gauss-Matuyama magnetostratigraphic boundary (Kaufman and Brigham-Grette, 1993). Attempts to date this reversal have been recently reviewed by Ohno et al. (2012); Singer (2014), and Head (2019). In particular, Deino et al. (2006) were able to tightly bracket the age of the reversal using high-precision $^{40}Ar/^{39}Ar$ dating of two tuffs in normally and reversely magnetized lacustrine sediments from Kenya, obtaining a value of 2.589 ± 0.003 Ma. Applying a +0.8% correction (based on the Fish Canyon sanidine standard astronomically calibrated to 28.201 Ma) yields an age of 2.61 Ma, which is followed by Singer (2014). However, Ohno et al. (2012) suggested there may have been a problem in the calibration of the $^{40}Ar/^{39}Ar$-dated tufts, and used relative paleointensity estimates for IODP Site U1314 to derive a midpoint age of 2.587 $\pm \geq$0.005 Ma for the Gauss–Matuyama reversal, a value accepted by Head (2019) and also adopted here (inclusive of error).

**Outgroup sequence.** IGM 100/1435 (Calibration 16), GCVP 5690 (Calibration 8), *Miocepphus bohaskai* (Calibration 2), Calibration 1.

**Outgroup age sequence.** 55.88, 34.44, 18.1, 2.58.

**95% soft upper bound.** 44.01 Ma.

**Calibration 2**

**Node calibrated.** MRCA of *Uria aalge* and *Alle alle.*

**Fossil taxon.** *Miocepphus bohaskai* Wijnker and Olson 2009.

**Specimen.** USNM 237142 (paratype; Wijnker and Olson, 2009), Smithsonian Institution, National Museum of Natural History, Washington DC, USA.

**Lower bound.** 18.1 Ma.

**Phylogenetic justification.** As in Smith (2015).

**Age justification.** Wijnker and Olson (2009, Figure 2) show the stratigraphic range of *M. bohaskai* to include the "Popes Creek Sand Member" of the Calvert Fm., which they date to approx. 19.8–19.2 Ma. The name "Popes Creek Sand Member" has since been discarded (Ward and Andrews, 2008); according to Weems and George (2013) it corresponds to unit C of the Fairhaven Mbr. and to the lower "Newport News unit" of Powars and Scott (1999). This unit spans dinocyst zones DN2b–DN2c (Weems et al., 2017). The top of zone DN2c and the base of calcareous nannoplankton zone NN3 are generally close in age (Browning et al., 2013), with various correlation charts showing the former to equal the latter (Edwards et al., 2010, Figure 29), slightly predate it (Perez et al., 2018, Figure 1), or slightly postdate it (Browning et al., 2013, Figure 6; McCarthy et al., 2013, Figure 2). According to the numeric ages given by McCarthy et al. (2013), the base of DN2b corresponds to 19.4 Ma and the top of DN2c to 18.1 Ma. An alternative dating is provided by Weems and George (2013, Figure 2), who show the top of the Faihaven C unit to correspond to the end of the early Hemingfordian NAMLA, dated at 17.5 Ma in accordance with Hilgen et al. (2012). However, this correlation is not explicitly established in the paper. We therefore prefer the dating of McCarthy et al. (2013) here. Note that Smith (2015) uses the end date of the Burdigalian instead, considering the age determination of the *M. bohaskai* fossils to be too uncertain, as it is based on biostratigraphy rather than radiometric dates. However, if this criterion were consistently applied, nearly all calibrations recommended by Smith (2015) would have to be rejected, as it is rare for radiometric samples to be available directly from the site of interest. Biostratigraphic correlation to localities for which radiometrically derived dates are available often represents the only option, and the best practices of Parham et al. (2011) allow for such chains of inferences as long as they are made explicit.

**Outgroup sequence.** IGM 100/1435 (Calibration 16), GCVP 5690 (Calibration 8), Calibration 2.

**Outgroup age sequence.** 55.88, 34.44, 18.1.

**95% soft upper bound.** 57.62 Ma.

**Calibration 3**

**Node calibrated.** MRCA of *Synthliboramphus craveri* and *Synthliboramphus hypoleucus*.

**Fossil taxon.** *Synthliboramphus rineyi* Chandler 1990.

**Specimen.** UCMP 61590/5566 (holotype; Chandler, 1990), University of California Museum of Paleontology, Berkeley, CA, USA.

**Lower bound.** 1.73 Ma.

**Phylogenetic justification.** Smith (2011a, Figure 7.7) sampled the same four extant *Synthliboramphus* species as our total-evidence tree and recovered the same topology for them, while showing *S. rineyi* to be sister to *S. hypoleucus*. As a result, this calibration can be directly re-used in our tree for the same node.

**Age justification.** The uncertain provenance of *S. rineyi* within the San Diego Formation means that the youngest possible date for the formation as a whole should be used. Unfortunately, the top of the San Diego Fm. is poorly constrained (Buczek et al., 2020). Smith (2015) used an age range of 3.6–1.5 Ma, the upper bound of which is based on Wagner et al.'s (2001) date for nonmarine facies from the lower part of the formation in Chula Vista, while the lower bound likely refers to Deméré (1983) – see Buczek et al. (2020, Figure 1) for an overview of previous estimates. Deméré's (1983) date was also used as the lower bound by Vendrasco et al. (2012), who took into account additional studies from the early 2000s to provide the oft-cited age range of 4.2–1.5 Ma (e.g., see Racicot et al., 2014). Several other studies have cited Vendrasco et al. (2012) in support of other lower bounds without clear justification: Velez-Juarbe (2017) cited the study in support of a "Zanclean to Gelasian" age, despite the fact that the 1.5 Ma lower bound implies a Calabrian age for the top of the formation (as explicitly noted, for example, by Smith, 2011b), and Boessenecker et al. (2019) incorrectly cited it as reporting an age range of 4.2–1.8 Ma. Recently, Buczek et al. (2020) used strontium isotope dating to suggest that the San Diego Fm. may be older than assumed based on biostratigraphy, deriving estimates consistent with a Zanclean–Piacenzian age (4.95–2.75 Ma). Unfortunately, it is not clear if their choice of localities was intended to span the entire duration of the formation. In the absence of this information, we follow the microfossil biostratigraphic estimate of Deméré (1983). Note that the value of 1.5 Ma given by the author was intended to reflect the age of the "*Emiliania annula* subzone", corresponding to subzone CN13a of Okada and Bukry (1980). Anthonissen and Ogg (2012) place the CN13a/CN13b subzonal boundary at 1.73 Ma, which is the minimum age used here.

**Outgroup sequence.** IGM 100/1435 (Calibration 16), GCVP 5690 (Calibration 8), *Miocepphus bohaskai* (Calibration 2), Calibration 3.

**Outgroup age sequence.** 55.88, 34.44, 18.1, 1.73.

**95% soft upper bound.** 43.89 Ma.

**Calibration 4**

**Node calibrated.** MRCA of *Cepphus columba* and *Cepphus carbo*.

**Fossil taxon.** *Cepphus olsoni* Howard 1982.

**Specimen.** LACM 107032 (holotype; Howard, 1982), Natural History Museum of Los Angeles County, Los Angeles, CA, USA.

**Lower bound.** 6.6 Ma.

**Phylogenetic justification.** Smith (2011a) and Smith and Clarke (2015) sampled all three extant *Cepphus* species in addition to the extinct *Cepphus olsoni* and *Pseudocepphus teres*, but their interrelationships varied depending on the analysis used. These included: (1) a polytomy between a (*C. olsoni* + *C. carbo*) clade, *C. columba*, and *C. grylle* in a parsimony analysis of 353 morphological characters and 11,601 bp scored for 3 extinct and 23 extant alcids (Smith, 2011a, Figure 6.7); (2) a sister-group relationship between (*C. olsoni* + *C. carbo*) and (*C. columba* + *C. grylle*) clades in a time-free Bayesian analysis of 353 morphological characters and 12,672 bp scored for 28 extinct and 52 extant charadriiforms (Smith and Clarke, 2015, Supplementary Figure A1); and (3) a sister-group relationship between a (*C. olsoni* + *C. carbo*) clade and *C. columba* to the exclusion of *C. grylle* in a parsimony analysis of the same dataset as in (2) (Smith and Clarke, 2015, Figure 3), as well as in a node-dating Bayesian analysis of the same dataset as in (2) but with 27 of the 28 extinct taxa removed (Smith and Clarke, 2015, Supplementary Figure A2). The third topology matches our total-evidence results and the first is still compatible with them, suggesting that *C. olsoni* can be directly re-used as a calibration in our tree.

**Age justification.** The dating of the lower unit of the San Mateo Formation was recently reviewed by Smith (2015) and Boessenecker et al. (2019). Domning and Deméré (1984) noted the presence of *Aepycamelus* in the lower assemblage (referred to as San Luis Rey River Local Fauna; SLRRLF), a taxon that reached the limit of its chronologic range in the late early Hemphillian (Hh2) (Tedford et al., 2004). A number of radiometric dates for localities correlated to Hh2 were provided by Tedford et al. (2004), although the provenance of some of the samples has been questioned (Kelly, 2013). Along with Tedford et al.'s (2004) Figure 6.2, these dates are likely the basis for the values of 6.7 Ma (Carrasco et al., 2007; Smith, 2015) or 6.8 Ma (Kelly and Secord, 2009; Gustafson, 2012) commonly cited in the literature for the top of Hh2. The standard Neogene timescale of Hilgen et al. (2012, Figure 29.9) gives a value of 6.6 Ma, which is followed here.

**Outgroup sequence.** IGM 100/1435 (Calibration 16), GCVP 5690 (Calibration 8), *Miocepphus bohaskai* (Calibration 2), Calibration 4.

**Outgroup age sequence.** 55.88, 34.44, 18.1, 6.6.

**95% soft upper bound.** 44.48 Ma.

**Calibration 5**

**Node calibrated.** MRCA of *Brachyramphus marmoratus* and *Brachyramphus brevirostris.*

**Fossil taxon.** *Brachyramphus dunkeli* Chandler 1990.

**Specimen.** SDSNH 24573/2971 (holotype; Chandler, 1990), San Diego Natural History Museum, San Diego, CA, USA.

**Lower bound.** 1.73 Ma.

**Phylogenetic justification.** Smith (2011a, Figures 7.6, 7.7) sampled all three extant *Brachyramphus* species and recovered the same topology for them as our total-evidence tree while showing *B. dunkeli* to be the sister group of *B. marmoratus*, meaning that this calibration can be directly re-used in our tree for the same node.

**Age justification.** As for Calibration 3.

**Outgroup sequence.** IGM 100/1435 (Calibration 16), GCVP 5690 (Calibration 8), *Miocepphus bohaskai* (Calibration 2), Calibration 5.

**Outgroup age sequence.** 55.88, 34.44, 18.1, 1.73.

**95% soft upper bound.** 43.89 Ma.

**Calibration 6**

**Node calibrated.** MRCA of *Fratercula arctica* and *Fratercula corniculata.*

**Fossil taxon.** *Fratercula* cff. *arctica* (Linnaeus, 1758).

**Specimens.** USNM 192994 and USNM 215783 (referred specimens; Olson and Rasmussen, 2001), National Museum of Natural History, Smithsonian Institution, Washington, DC, USA. Note that specimen USNM 490887, on which the corresponding calibration was based by Smith (2015), was assigned to *F.* aff. *cirrhata* rather than *F.* aff. *arctica* by Olson and Rasmussen (2001). The referral of various Lee Creek Mine specimens to *Fratercula* as opposed to *Cerorhinca* is supported by characters of the coracoid, humerus, and tarsometatarsus (Olson and Rasmussen, 2001, 280), while their assignment to *Fratercula* aff. *arctica* rather than *F.* aff. *cirrhata* is based on differences in the coracoid and tibiotarsus (Olson and Rasmussen, 2001, 282). Therefore, from the Lee Creek Mine specimens described by Olson and Rasmussen (2001) as *F.* aff. *arctica*, we specifically choose USNM 192994 (an incomplete coracoid) and USNM 215783 (a distal tibiotarsus) as the basis for the calibration.

**Lower bound.** 3.92 Ma.

**Phylogenetic justification.** As in Smith (2015).

**Age justification.** Smith (2015) details why *Fratercula* aff. *arctica* and other contemporary remains from the Yorktown Formation (*Fratercula* aff. *cirrhata*) provide the oldest well-established record of Fraterculini, as opposed to Miocene fossils with less certain affinities. The Lee Creek Mine locality belongs to the Sunken Meadow Member of the Yorktown Formation (Gibson and Geisler, 2009), for which a number of age estimates are available; see Johnson et al. (2017) and Boessenecker et al. (2019) for recent reviews. Smith (2015) relies on the ostracod biostratigraphic evidence reported by Hazel (1983), who assigned Lee Creek to the *Orionina vaughani* biozone, and noted that glautonite collected from this biozone at Grove Creek, Virginia, was dated to $4.4 \pm 0.2$ Ma using K/Ar dating. This is consistent with the model of Krantz (1991), who correlated the transgressive event associated with the depositon of the Sunken Meadow Member with deep-ocean $\delta^{18}$O records to derive an age of 4.5–4.4 Ma. While the *Orionina vaughani* biozone spans calcareous nannofossil zones NN12 through NN15, the youngest part of the Yorktown Formation at Lee Creek is no younger than its middle (Hazel, 1983, 97). Based on this information and Hazel's (1983) Figure 4, Marx and Fordyce (2015) suggested bracketing the age of Lee Creek using the top of biozone NN14, thus deriving a minimum age of 3.92 Ma following Anthonissen and Ogg (2012). This justification was accepted by Boessenecker et al. (2019), and it is also followed here.

**Outgroup sequence.** IGM 100/1435 (Calibration 16), GCVP 5690 (Calibration 8), *Miocepphus bohaskai* (Calibration 2), *Aethia barnesi* (Calibration 7), Calibration 6.

**Outgroup age sequence.** 55.88, 34.44, 18.1, 6.6, 3.92.

**95% soft upper bound.** 32.03 Ma.

**Calibration 7**

**Node calibrated.** Crown-group Fraterculinae (MRCA of *Aethia cristatella* and *Fratercula arctica*).

**Fossil taxon.** *Aethia barnesi* Smith 2014.

**Specimen.** LACM 107031, (holotype; Smith, 2014), Natural History Museum of Los Angeles County, Los Angeles, CA, USA.

**Lower bound.** 6.6 Ma.

**Phylogenetic justification.** As in Smith (2015).

**Age justification.** As for Calibration 4.

**Outgroup sequence.** IGM 100/1435 (Calibration 16), GCVP 5690 (Calibration 8), *Miocepphus bohaskai* (Calibration 2), Calibration 7.

**Outgroup age sequence.** 55.88, 34.44, 18.1, 6.6.

**95% soft upper bound.** 44.48 Ma.

**Calibration 8**

**Node calibrated.** Crown-group Alcoidea (MRCA of *Alca torda* and *Stercorarius parasiticus*).

**Fossil taxon.** Total-group Alcidae *incertae sedis* Chandler and Parmley 2003.

**Specimen.** GCVP 5690, Georgia College Vertebrate Paleontology collection, Georgia College and State University, Milledgeville, GA, USA.

**Lower bound.** 34.44 Ma.

**Phylogenetic justification.** As in Smith (2015).

**Age justification.** The primary evidence for the age of the Hardie Mine fossils comes from the locality's dinocyst assemblage, which has been compared to nearby Georgia and South Carolina samples placed in calcareous nannofossil zone NP19–20 (Parmley and Alan, 2003), and accordingly dated at 36.0–34.2 Ma based on Berggren et al. (1995). Parmley et al. (2006, 343) narrowed down this range to 35.5–34.5 Ma based on unspecified "[n]ew vertebrate and invertebrate faunal evidence"; this was possibly intended to correspond to the middle Chadronian NALMA (North American land mammal age), dated at 35.7–34.7 Ma by Prothero and Emry (2004). However, this correlation was not made explicit, and more recent reviews of the Hardie Mine mammal remains only regard the locality as Chadronian or older (Westgate, 2012). Accordingly, we base our estimate on the higher-resolution dinocyst evidence, and use the revised cycle-calibrated age range of 36.97–34.44 Ma for the NP19–20 zone derived by Anthonissen and Ogg (2012).

**Outgroup sequence.** IGM 100/1435 (Calibration 16), Calibration 8.

**Outgroup age sequence.** 55.88, 34.44.

**95% soft upper bound.** 70.04 Ma.

**Calibration 9**

**Node calibrated.** Crown-group Arenariinae (MRCA of *Arenaria interpres* and *Calidris canutus*).

**Fossil taxa.** *Mirolia brevirostrata* Ballmann 2004, *Mirolia dubia* Ballmann 2004, *Mirolia parvula* Ballmann 2004, ?*Mirolia mascalidris* Ballmann 2004.

**Specimens.** 1970 XVIII Steinberg (collective designation of the holotypes of *M. brevirostrata*, *M. dubia*, *M. parvula*, and ?*M. mascalidris*; Ballmann, 2004), Bayerische Staatssammlung für Paläontologie und historische Geologie, Munich, Germany.

**Lower bound.** 12.6 Ma.

**Phylogenetic justification.** Ballmann (2004) assigned the genus *Mirolia* to "Calidridinae", or the group comprising *Calidris* and the monotypic genera *Eurynorhynchus*, *Limicola*, *Micropalama*, *Philomachus*, and *Tryngites*. Since the traditional *Calidris sensu stricto* is not monophyletic with respect to these genera (Gibson and Baker, 2012; see also our total-evidence tree), the latter have been merged into *Calidris* (Banks, 2012; Boyd, 2019). This renders Calidridinae *sensu* Ballmann 2004 (= Calidrini [*sic*] *sensu* Cracraft, 2013) redundant with respect to the enlarged genus, hereafter referred to as *Calidris sensu lato*. As noted by Zelenkov and Kurochkin (2015), these taxonomic changes may require *Mirolia* to be merged into *Calidris* as well.

*Mirolia* has been previously used to calibrate the crown group of *Calidris sensu lato* (Smith, 2015) as well as the divergence between Scolopacinae on the one hand and Arenariinae (*sensu* Banks, 2012) plus Tringinae on the other (De Pietri et al., 2020b). Both decisions are poorly justified. Ballmann (2004, Figure 3) showed *M. brevirostrata* to exhibit 17 features of the coracoid, humerus, and tarsometatarsus that he regarded as typical but not necessarily diagnostic of "Calidridinae". He also provided a list of four mandibular characters distinguishing *Calidris sensu lato* from Tringinae (Ballmann, 2004, Figure 4), two of which (a long processus retroarticularis and a large insertion area for m. depressor mandibulae) could be ascertained from ?*M. mascalidris* (Ballmann, 2004, Figure 8). The presence of these characters indicates that *Mirolia* is more closely related to *Calidris* than to Tringinae, and its assignment to a node predating the divergence of these two taxa, as in De Pietri et al. (2020b), is therefore overly conservative. The four characters cited by Smith (2015) in support of assigning *Mirolia* to the crown group of "Calidridinae" were intended to suggest a close relationship between the fossil genus and "*Philomachus*" (= *Calidris pugnax*) as well as "*Tryngites*" (= *Calidris subruficollis*) to the exclusion of other extant "calidridines" (Ballmann, 2004, 111). However, these two taxa are no more closely related to each other than to other species of *Calidris* (Baker et al., 2007; Gibson and Baker, 2012; see also our total-evidence tree), and the characters that unite them are consequently either homoplastic or plesiomorphic for *Calidris* or a large subclade thereof. The latter option is plausible given that two of the characters in question that relate to bill shape (the shallow and narrow origin of m. depressor mandibulae corresponding to a relatively shorter processus retroarticularis; short bill with moderate dorsal bar reinforcement) may reflect a lack of adaptations for tactile feeding, and Ballmann (2004, 112) considered *C. pugnax* and *C. subruficollis* to "still represent in our time the evolutionary level of *Mirolia*". Accordingly, *Mirolia* cannot be associated with any specific subclade of *Calidris sensu lato*, and its position on the stem of the genus cannot be ruled out. We thus use it to calibrate the divergence of *Calidris sensu lato* from its sister taxon (*Arenaria*), which corresponds to the crown group of Arenariinae *sensu* Banks 2012.

Notably, Ballmann (2004) did not explicitly compare *Mirolia* to *Arenaria* and *Prosobonia*, now known to represent the closest living relatives of *Calidris sensu lato*: in the traditional morphological classifications referenced by the author (Jehl, 1968; Strauch, 1978; Zusi, 1984), *Prosobonia* was regarded as a member of Tringinae, while *Arenaria* was treated as constituting a subfamily of its own. In light of these missing comparisons, a more conservative assignment of the fossil to the total group of (Arenariinae + *Prosobonia*), equivalent to the crown group of (Arenariinae + Tringinae), may also be justifiable. Here, we prefer a more deeply nested position for *Mirolia* in line with Ballmann's (2004) original assessment.

**Age justification.** The status of the four species of *Mirolia* as the oldest known representatives of Arctic sandpipers (*Calidris sensu lato*) was recently affirmed by De Pietri et al. (2020b). The upper bound on the age of *Mirolia* is provided by the asteroid impact that created a crater subsequently filled by a long-term evaporative saline lake from whose deposits the fossils were described (Ballmann, 2004; Moncunill-Solé et al., 2019). A high-precision $^{40}$Ar/$^{39}$Ar date of 14.808 ± 0.038 Ma for the impact was recently provided by (Schmieder et al., 2018a); Rocholl et al. (2018) used orbital tuning and paleomagnetic constraints to propose two alternative ages of 14.870 and 14.609 Ma, favoring the former (but see the response by Schmieder et al., 2018b. In contrast, the minimum age is only poorly constrained. The Nördlinger Ries lake is estimated to have lasted 0.3–2 Myr (Moncunill-Solé et al., 2019), and its vertebrate fossils are thought to be restricted to its final freshwater stage (Arp, 2006), implying a minimum age close to 12.8 Ma. This is compatible with the correlation chart of Arp et al. (2016, Figure 2), who show the relevant deposits to be at least 12.7 Ma old while rejecting the hypothesis of their deposition during a discrete freshwater stage. Contemporary mammal remains belong to Neogene mammal unit MN6 (Ballmann, 2004), which is estimated to range from 14.2 Ma to 13.1–12.6 Ma based on faunal correlations or from 14.2 Ma to 14.1 Ma based on reference localities (Hilgen et al., 2012). Since Nördlinger Ries is not a reference locality for the unit and its assignment to MN6 has been determined based on faunal correlations, only the former range is appropriate. Hence, we choose 12.6 Ma as the conservative lower bound on the age of *Mirolia*.

**Outgroup sequence.** IGM 100/1435 (Calibration 16), GCVP 5690 (Calibration 8), *Nupharanassa tolutaria* (Calibration 12), *Elorius* and *Parvelorius* (Calibration 11), Calibration 9.

**Outgroup age sequence.** 55.88, 34.44, 30.5, 20.0, 12.6.

**95% soft upper bound.** 37.77 Ma.

**Calibration 10**

**Node calibrated.** Total-group *Gallinago* (MRCA of *Gallinago gallinago* and *Coenocorypha aucklandica*).

**Fossil taxon.** *Gallinago azovica* Zelenkov and Panteleyev 2015.

**Specimen.** ZIN PO 7299 (holotype; Zelenkov and Panteleyev, 2015), Zoological Institute of the Russian Academy of Sciences, Saint Petersburg, Russia.

**Lower bound.** 6.1 Ma.

**Phylogenetic justification.** The holotype of *G. azovica* was referred to the Scolopacidae based on a combination of characters (Zelenkov and Panteleyev, 2015); of these, the absence of the foramen nervi supracoracoidei only characterizes the Scolopaci as a whole (and may not be apomorphic even for the latter clade; De Pietri and Mayr, 2012). The fossils also display a scar separating the shaft from the procoracoid process, which is unique to *Calidris sensu lato*, *Limnodromus*, and *Gallinago* among extant scolopacids; since these three taxa do not form an exclusive clade, the character is most parsimoniously regarded as independently acquired in all three taxa. Importantly, Zelenkov and Panteleyev (2015) explicitly note that this character is missing in *Coenocorypha*, which forms (along with *Chubbia*) the sister group of *Gallinago* in our total-evidence tree. The specimen can further be distinguished from *Limnodromus* based on the shape of the base of the procoracoid process (wide rather than narrow) and from *Calidris sensu lato* based on a lower-positioned tuberculum brachiale. Zelenkov and Panteleyev (2015) did not note whether these additional characters represent apomorphies or plesiomorphic retentions. While *G. azovica* has not been included in a phylogenetic analysis, other paleornithologists have considered its inclusion within *Gallinago* to be sufficiently robust to use it as a calibration (De Pietri et al., 2020b).

**Age justification.** The fossils of *G. azovica* come from the Morskaya-2 locality, dated to the middle Turolian (European reference levels: MN12–?MN13) by Titov and Tesakov (2013). A numeric age estimate of 7.5–7.1 Ma provided by Zelenkov et al. (2017) is unsupported by references but likely based on the correlation chart of Titov and Tesakov (2013, Figure 24.2), which places the locality at the boundary of the Maeotian and Pontian Easten Paratethys regional stages. As noted by Titov and Tesakov (2013), two paleomagnetic ages are available for the base of the Pontian, and thus for the minimum age of the Morskaya-2 fauna. Of these, Titov and Tesakov (2013, Figure 24.2) prefer the older (7.1 Ma), while Hilgen et al. (2012, Figure 29.8) prefer the younger (6.1 Ma). Here, we conservatively use the younger age as well, contrary to De Pietri et al. (2020b), who set the age of *G. azovica* to 7.0 Ma.

**Outgroup sequence.** IGM 100/1435 (Calibration 16), GCVP 5690 (Calibration 8), *Nupharanassa tolutaria* (Calibration 12), *Elorius* and *Parvelorius* (Calibration 11), *Mirolia* spp. (Calibration 9), Calibration 10.

**Outgroup age sequence.** 55.88, 34.44, 30.5, 20.0, 12.6, 6.1.

**95% soft upper bound.** 28.39 Ma.

**Calibration 11**

**Node calibrated.** Crown-group Scolopacidae (MRCA of *Scolopax rusticola* and *Numenius arquata*).

**Fossil taxa.** *Elorius paludicola* Milne-Edwards 1868, ?*Elorius limosoides* De Pietri and Mayr 2012, *Parvelorius gracilis* (Milne-Edwards, 1868), ?*Parvelorius calidris* De Pietri and Mayr 2012.

**Specimens.** MNHN Av. 9503 (lectotype of *Elorius paludicola*; De Pietri and Mayr, 2012), Muséum national d'histoire naturelle, Paris, France; NMB S.G. 6698, NMB S.G. 4016, and NMB MA 672 (paratypes of ?*Elorius limosoides*; De Pietri and Mayr, 2012), Naturhistorisches Museum Basel, Basel, Switzerland; MNHN Av. 9528 (lectotype of *Parvelorius gracilis*; De Pietri and Mayr, 2012), Muséum national d'histoire naturelle, Paris, France; NMB S.G. 16551 (holotype of ?*Parvelorius calidris*; De Pietri and Mayr, 2012), Naturhistorisches Museum Basel, Basel, Switzerland.

**Lower bound.** 20.0 Ma.

**Phylogenetic justification.** According to De Pietri and Mayr (2012, 1190), *Elorius paludicola* can be "confidently assigned" to the Scolopacidae based on the small foramen vasculare distale at the distal end of the tarsometatarsus. However, this character state is plesiomorphic within the Charadriiformes, and can only distinguish the taxon from the Jacanidae and Rostratulidae (Mayr, 2011b); the other two lineages of the Scolopaci (Thinocoridae and Pedionomidae) were likely excluded from the comparison because of their restricted present-day distribution. Within the Scolopacidae, *E. paludicola* shares an open canal for the tendon of m. flexor digitorum longus with *Limosa* and some species of *Tringa* (De Pietri and Mayr, 2012, 1190, 1192). The latter trait is also shared by ?*Elorius limosoides* (De Pietri and Mayr, 2012, 1191), which is assumed to be congeneric with *E. paludicola* based on the overall similarity of the tarsometatarsus (*ibid.*), and which further shares with *Limosa* a very narrow sulcus extensorius on the distal end of the tibiotarsus, a trait that distinguishes the two taxa from both earlier-diverging (*Numenius*) and more deeply nested (Tringinae, *Calidris sensu lato*) members of the Scolopacidae (De Pietri and Mayr, 2012, 1192). *Parvelorius* (especially ?*P. calidris*) exhibits a well-developed second (dorsal) fossa pneumotricipitalis (De Pietri and Mayr, 2012, 1194), a feature noted to be apomorphic for the Scolopacidae and otherwise only occurring in the Thinocoridae among the Scolopaci (Mayr, 2011b). Along with the presence of a hook-like rather than blunt tip of the tuberculum ventrale of the humerus (as in *Calidris*, *Arenaria*, and *Gallinago*, but different from *Numenius* and *Limosa*) in *P. gracilis* (De Pietri and Mayr, 2012, 1193), these traits suggest that *Elorius* and *Parvelorius* represent crown-group scolopacids diverging after *Numenius* (possibly closely related to *Limosa* or on the stem lineage of the clade comprising the Scolopacinae, Tringinae, and Arenariinae), although we note that supporting this hypothesis with an explicit list of derived characters is difficult, both because of the paucity of morphological apomorphies for the Scolopacidae (De Pietri and Mayr, 2012) and because of the fragmentary nature of the material. Therefore, we conservatively treat all four taxa as crown scolopacids of uncertain affinities.

**Age justification.** All fossils referred to *Elorius* and *Parvelorius* come from localities jointly known as Saint-Gérand-le-Puy, which are generally dated to the early Miocene, although a late Oligocene age cannot be excluded for some of them (De Pietri et al., 2011a; De Pietri and Mayr, 2012). A Miocene age for the two taxa is further supported by the fact that "the best part of [their] material" comes from the Montaigu-le-Blin quarry (De Pietri and Mayr, 2012, 1179), the type locality of mammal Neogene unit MN2a (Mourer-Chauviré, 2000). In terms of numeric ages, De Pietri et al. (2011a) date the Saint-Gérand-le-Puy avifauna to 22.5–20.5 Ma, citing Steininger (1999) in support of this estimate. However, Steininger (1999) correlated the top of MN2 to the base of chron C6r, which is also consistent with the more recent review of Agustí et al. (2001), and which would correspond to a numerical age of 20.04 Ma (Ogg, 2012). The 20.5 Ma minimum age estimate was perhaps derived from the Aquitanian/Burdigalian boundary (20.44 Ma; Hilgen et al., 2012), which was suggested to approximately correspond to the Agenian/Orleanian (MN2/MN3) boundary by Steininger et al. (1989). More recently, Hilgen et al. (2012, Figure 29.9) estimated the top of MN2 to ∼19.5 Ma; however, the MN scale of Hilgen et al. (2012) has been criticized for disregarding large-bodied mammals or only considering them at the genus level (Casanovas-Vilar, 2017). An alternative estimate of 19.88 Ma was proposed by Ruiz-Sánchez et al. (2012) based on radiometric evidence from the Ebro Basin and magnetostratigraphy. The numeric age estimate could potentially be further refined by setting the lower bound to the MN2a/MN2b subzonal boundary, which is, however, even more poorly constrained than the MN2(b)/MN3 boundary. Sen (1997, Figure 1) showed the top of MN2a to correspond to the base of chron C6AAn (21.16 Ma; Hilgen et al., 2012); note that Head et al. (2016) interpreted the figure as showing a correlation with the top of C6AAn instead (21.08 Ma; Hilgen et al., 2012). In contrast, Hilgen et al. (2012, Figure 29.9) date the type locality for MN2a (Montaigu-le-Blin) to 21.7–20.0 Ma. The lower end of this range is here used as a conservative estimate of the minimum age for *Elorius* and *Parvelorius*, as it is also congruent with the magnetostratigraphically and radiometrically constrained end date for MN2 as a whole (Agustí et al., 2001; Ruiz-Sánchez et al., 2012).

**Discussion.** Several possible occurrences of the Scolopacidae predate the early Miocene *Elorius* and *Parvelorius*. One such record consists of an uncatalogued specimen from the Vachères limestones of Luberon (early Oligocene, European reference level MP24: Riamon et al., 2020) described by Roux (2002). The scolopacid affinity of the specimen is considered to be likely (De Pietri and Mayr, 2012), but was only established based on the overall shape of the acrocoracoid process rather than derived characters (Mayr, 2009, 91). Moreover, the specimen was not linked to any particular lineage within the scolopacid crown, meaning that a stem-scolopacid position cannot be ruled out. Such a position would make it redundant with respect to Calibration 12, assigned to the jacanid-rostratulid split.

Another possible scolopacid predating the Saint-Gérand-le-Puy taxa, specimen FLS 367 064 from the latest Oligocene of France (Mourer-Chauviré et al., 2004), was referred to Scolopaci based on the absence of the foramen nervi supracoracoidei, but this character is no longer considered to represent an unambiguous apomorphy of the clade (De Pietri and Mayr, 2012). The assignment of FLS 367 064 to the Scolopacidae was based on overall morphology of the coracoid rather than a list of apomorphies. Among the scolopacids, the remains were noted to be particularly similar to the genus *Phalaropus* based on the overall shape of the acrocoracoid process; however, the phylogenetic distribution of the similarities listed was not commented upon, and it is unclear whether they represent apomorphies or symplesiomorphies, or even well-defined characters. Given this lack of unambiguous derived characters and extremely fragmentary nature of the specimen, which only includes the omal extremity of the left coracoid, we consider *Elorius* and *Parvelorius* to be the earliest well-supported crown-group members of the Scolopacidae.

**Outgroup sequence.** IGM 100/1435 (Calibration 16), GCVP 5690 (Calibration 8), *Nupharanassa tolutaria* (Calibration 12), Calibration 11.

**Outgroup age sequence.** 55.88, 34.44, 30.5, 20.0.

**95% soft upper bound.** 48.32 Ma.

**Calibration 12**

**Node calibrated.** Crown-group Jacanoidea (MRCA of *Jacana jacana* and *Rostratula benghalensis*).

**Fossil taxon.** *Nupharanassa tolutaria* Rasmussen et al. 1987.

**Specimen.** DPC 2580 (holotype; Rasmussen et al., 1987), Duke Lemur Center, Durham, NC, USA.

**Lower bound.** 30.5 Ma.

**Phylogenetic justification.** Smith (2015) recommends using the younger congener *N. bulotorum* with the justification that its jacanid affinities have been confirmed using a phylogenetic analysis, unlike those of *N. tolutaria*. However, *N. tolutaria* has never been suggested to represent anything but a jacana, and has been used as a calibration by other paleornithologists (De Pietri et al., 2020b), who regarded it as an "unequivocal jacanid". Moreover, Figure 6 of Rasmussen et al. (1987) shows that it exhibits the same enlarged distal vascular foramen on the tarsometatarsus that was optimized as a jacanid apomorphy in the analysis that supported a jacanid position for *N. bulotorum* (Smith, 2011a). Hence, we consider the use of *N. bulotorum* too conservative. Note that the choice between the two species of *Nupharanassa* matters, since they come from two different quarries within the Jebel Qatrani Formation, of which Quarry E (yielding *N. tolutaria*) is substantially older.

**Age justification.** Unfortunately, correlating the paleomagnetic record to the global standard has proved difficult. Seiffert (2006), whose dating was followed by Smith (2015), placed Quarry E "near the bottom of Chron C12r" (age range: 33.157–31.034 Ma according to Vandenberghe et al., 2012). This led to numerical estimates of 33.0 (Stidham and Smith, 2015) or 32.5 Ma (Coster et al., 2015) appearing in the literature. However, in the correlation suggested by Underwood et al. (2013, Figure 5), Seiffert's bottom of C12r corresponds to the bottom of C11r (age range: 30.591–29.970 Ma according to Vandenberghe et al., 2012). Since Parham et al. (2011) recommend using the youngest possible age of the fossil as the hard minimum, we follow Coster et al. (2015) in using 30.5 Ma as the numerical age of Quarry E, and thus of *N. tolutaria*.

**Outgroup sequence.** IGM 100/1435 (Calibration 16), GCVP 5690 (Calibration 8), Calibration 12.

**Outgroup age sequence.** 55.88, 34.44, 30.5.

**95% soft upper bound.** 59.12 Ma.

**Calibration 13**

**Node calibrated.** Crown-group Thinocoroidea (MRCA of *Thinocorus rumicivorus* and *Pedionomus torquatus*).

**Fossil taxon.** *Oligonomus milleri* De Pietri et al. 2015.

**Specimen.** SAMA P27976 (holotype; De Pietri et al., 2015), South Australian Museum, Adelaide, SA, Australia.

**Lower bound.** 24.47 Ma.

**Phylogenetic justification.** De Pietri et al. (2015) referred *Oligonomus* to the Pedionomidae based on a combination of 7 characters, of which one is unique to the clade among the Scolopaci, and can therefore be optimized as its unambiguous apomorphy: the absence of a recess in sulcus musculi supracoracoidei, medially, below the facies articularis clavicularis. The remaining 6 characters may represent pedionomid apomorphies that also arose convergently in other lineages of Scolopaci, or symplesiomorphies that were lost in some of its subclades.

**Age justification.** The only known specimen of *Oligonomus* was collected from Etadunna Formation faunal zone B, correlated to chron C7r by Megirian et al. (2010). This corresponds to an age of 24.76–24.47 Ma based on the timescale of Vandenberghe et al. (2012).

**Outgroup sequence.** IGM 100/1435 (Calibration 16), GCVP 5690 (Calibration 8), *Nupharanassa tolutaria* (Calibration 12), Calibration 13.

**Outgroup age sequence.** 55.88, 34.44, 30.5, 24.47.

**95% soft upper bound.** 48.98 Ma.

**Calibration 14**

**Node calibrated.** Crown-group Haematopodoidea (MRCA of *Haematopus ostralegus* and *Recurvirostra avosetta*).

**Fossil taxon.** Total-group Haematopodidae *incertae sedis* De Pietri et al. 2013.

**Specimen.** NMB S.G.20252, Naturhistorisches Museum Basel, Basel, Switzerland.

**Lower bound.** 20.0 Ma.

**Phylogenetic justification.** NMB S.G.20252 was referred to the clade including the Charadriidae, Recurvirostridae, and Haematopodidae on the basis of four cranial characters, of which three (the presence of a distinct second opening caudal of foramen nervi maxillomandibularis, a functional processus basipterygoidei, and of well-developed fonticuli occipitales) were optimized as apomorphies of this clade in the phylogenetic analyses of Mayr (2011b, Figure 3). Note that in our total-evidence tree, this clade also includes *Ibidorhyncha*, which was not examined by Mayr (2011b) nor by De Pietri et al. (2013), but which was noted to also possess a functional processus basipterygoidei by Strauch (1978). In addition, NMB S.G.20252 exhibits two characters which were otherwise only present in species of *Haematopus* among the charadriiform taxa examined by De Pietri et al. (2013), and as such can be regarded as unambiguous apomorphies of the Haematopodidae: a deep crescent-shaped depression dorsal to the crista nuchalis transversa, and the foramen magnum displaying a square-like shape with a straight dorsal rim in occipital view.

**Age justification.** As for Calibration 11.

**Outgroup sequence.** IGM 100/1435 (Calibration 16), Calibration 14.

**Outgroup age sequence.** 55.88, 20.0.

**95% soft upper bound.** 68.50 Ma.

**Calibration 15**

**Node calibrated.** Crown-group Chionida (MRCA of *Chionis albus* and *Burhinus grallarius*).

**Fossil taxon.** *Chionoides australiensis* De Pietri et al. 2016a.

**Specimen.** SAM P41458 (holotype; De Pietri et al., 2016a), South Australian Museum, Adelaide, SA, Australia.

**Lower bound.** 24.76 Ma.

**Phylogenetic justification.** *Chionoides australiensis* can be referred to Chionoidea (Chionidae + *Pluvianellus*) on the basis of the following four apomorphies of the coracoid: (1) ventral deflection of the medial portion of the acrocoracoid ('tuberculum brachiale') relative to the shaft, (2) ventral portion of the facies articularis clavicularis exhibiting a rounded rather than dorsoventrally flat and medially protruding surface, (3) processus acrocoracoideus high (omally projecting) relative to facies articularis clavicularis, (4) ventral area adjacent to the facies articularis humeralis separated from the ligamental insertion area on the tuberculum brachiale by a prominent ridge (De Pietri et al., 2016a). Based on its age (predating the previous molecular dating estimates of the divergence between the Chionidae and *Pluvianellus*) and the fact that the fossil displays a mosaic of chionid and pluvianellid traits, De Pietri et al. (2016a) regarded *Chionoides* as a probable stem-group representative of the Chionoidea.

**Age justification.** *Chionoides* comes from Faunal Zone A at Lake Palankarinna, which represents the oldest fossil assemblage of the Etadunna Formation and can be correlated with chrons C7Ar and C7An (Megirian et al., 2010). The minimum age given here therefore corresponds to the end of chron C7An according to Vandenberghe et al. (2012).

**Discussion.** *Chionoides supersedes* the previous oldest record of Chionida (Chionoidea + Burhinidae), the burhinid *Genucrassum bransatensis* from the latest Oligocene (European reference level MP30) of France (De Pietri et al., 2016a), recommended as a calibration by Smith (2015).

**Outgroup sequence.** IGM 100/1435 (Calibration 16), Calibration 15.

**Outgroup age sequence.** 55.88, 24.76.

**95% soft upper bound.** 68.94 Ma.

#### Calibration 16

**Node calibrated.** Crown-group Charadriiformes (MRCA of *Charadrius hiaticula* and *Larus marinus*).

**Fossil taxon.** Charadrii (?Chionida) total-group *incertae sedis* Hood et al. 2019.

**Specimen.** IGM 100/1435, Mongolian Institute of Geology, Ulaanbaatar, Mongolia.

**Soft lower bound.** 55.88 Ma.

**Phylogenetic justification.** Hood et al. (2019) originally concluded that the fossil cannot be assigned to crown-group charadriiforms, and can only be placed within the charadriiform total group (Pan-Charadriiformes). However, the phylogenetic analysis of Musser and Clarke (2020) found it within the crown under one of the three constraints used, and specifically as the sister group of *Chionis* (note that under the other two constraints, the trees were too poorly resolved to distinguish between crown-group and total-group affinities). Our re-analyses of the Musser and Clarke (2020) matrix using Bayesian inference and more stringent versions of the original constraints recovered the (*Chionis* + IGM 100/1435) clade with moderate support under all three constraints (Figure A.10); however, since the support for deeper nodes was extremely poor, we choose to conservatively identify IGM 100/1435 only as a member of total-group Charadrii, and hence a crown-charadriiform.

**Age justification.** All avian fossils described by Hood et al. (2019) are reported to have been found 'in and around "Quarry 2" of Dashzėvėg et al. (1998)' at the Tsagan Khushu locality of the Naranbulag (also transliterated as "Naran Bulak") Formation (Hood et al., 2019, 2). The authors consider this site to lie at the base of the Bumban Member of the formation, consistent with a number of earlier publications (Dashzėvėg and Russell, 1988; Lucas and Kondrashov, 2004; Mao et al., 2017). The Bumban Member conformably overlies the Naran Member (Dashzėvėg et al., 1998), and the boundary between the two units corresponds to the boundary between the Gashatan and Bumbanian Asian Land Mammal Ages (ALMAs) (Bowen et al., 2002; Hwang et al., 2010). The age of the latter boundary was estimated at 55.7–54.97 Ma by Bowen et al. (2002). Later publications treated these estimates as absolute dates and used their deviation from the currently accepted beginning of the Eocene (see below) to claim that the Gashatan–Bumbanian transition may have taken place in the early Eocene (Hwang et al., 2010) instead of exactly coinciding with the Paleocene–Eocene boundary (Zelenkov, 2018). However, the lower bound of 54.97 Ma simply corresponded to a contemporary age estimate (Wing et al., 1999) for the negative carbon ($\delta^{13}$C) isotope excursion that defines the Paleocene–Eocene boundary (Vandenberghe et al., 2012). (Note that while Wing et al. (1999) are cited for the 54.97 Ma date by multiple studies – e.g., Ting et al. (2003) – this value may in fact represent a misreading of the estimate obtained under their age model 1, which was 15 kyr younger.) Consequently, insofar as the correlations proposed by Bowen et al. (2002) are accepted, the Gashatan–Bumbanian boundary must be treated as coinciding with the Paleocene–Eocene boundary, and any change to the dating of the latter also applies to the dating of the former.

Unfortunately, the precisely dated Paleocene–Eocene boundary provides only the upper (rather than lower) bound on the age of the Tsagan Khushu Quarry 2 fossils. Hood et al. (2019) date the specimen to ∼55 Ma, but this value appears to be arbitrary and possibly based on the obsolete numeric ages given by Bowen et al. (2002). A better justified lower bound could be based on the end of the Bumbanian ALMA, which is, however, poorly constrained: Vandenberghe et al. (2012, Figure 28.10) estimate it to be as young as ∼52.0 Ma, while Wang et al. (2010) suggest it to be as old as 54.8 Ma. Moreover, the whole Bumban Member of the Naranbulag Formation falls within the first subdivision of the Bumbanian (Missiaen, 2011; though note that the subdivision of the Bumbanian into three biochrons was doubted by Lucas and Kondrashov, 2004), and Tsagan Khushu Quarry 2 lies at the very bottom of the Bumban Member (Dashzėvėg and Russell, 1988, Figure 2). Therefore, using the end of the Bumbanian as the minimum age of IGM 100/1435 would underestimate its true age far more than the use of the Paleocene–Eocene boundary would overestimate it. To account for this issue while simultaneously avoiding arbitrary adjustments, we set the minimum age for Calibration 16 to the youngest plausible date for the Paleocene–Eocene boundary, derived from the astronomically calibrated estimate of Westerhold et al. (2018) inclusive of error (55.93 ± 0.05 Ma). The resulting value of 55.88 Ma may slightly underestimate the true age of the boundary, as it lies outside of the error bars of another recent astrochronologic estimate (56.01 ± 0.05 Ma; Zeebe and Lourens, 2019). We consider this desirable, since the Tsagan Khushu Quarry 2 fossils postdate the boundary somewhat. Moreover, unlike the lower bounds of the remaining calibrations, the lower bound of Calibration 16 is treated as soft, allowing the analysis to sample ages younger than the set minimum. This mitigates the risk of bias that is otherwise associated with insufficiently conservative lower bounds (Parham et al., 2011).

**Outgroup sequence.** N/A.

**Outgroup age sequence.** N/A.

**95% soft upper bound.** 66.0 Ma.

#### 1.2 Rejected calibrations

The fossil evidence listed below was considered for inclusion in the calibration set but rejected on the grounds of phylogenetic uncertainty, stratigraphic uncertainty, or redundancy with respect to calibrations already present in the set. Since some of these fossils have been either proposed or actually employed as age constraints by previous calibration compendia and node-dating analyses (Jarvis et al., 2014; Claramunt and Cracraft, 2015; Smith, 2015; Kimball et al., 2019; De Pietri et al., 2020b), we provide detailed reasons for their exclusion from the calibration set used in this study.

##### *Becassius charadriioides*

**Potential node calibrated.** Crown-group Glareoloidea (MRCA of *Glareola pratincola* and *Dromas ardeola*).

**Fossil taxon.** *Becassius charadriioides* De Pietri and Mayr 2012.

**Specimen.** NMB SG. 12784 (holotype; De Pietri and Mayr, 2012), Naturhistorisches Museum Basel, Basel, Switzerland.

**Lower bound.** 20.0 Ma.

**Discussion.** *Becassius charadriioides* was originally described as a member of Scolopaci with uncertain affinities (De Pietri and Mayr, 2012) based on its possession of a transverse ridge across the incisura capitis humeri, a character found to be apomorphic for Scolopaci by Strauch (1978) and Mayr (2011b). The fossils of *Becassius* were collected from the Saint-Gérand-le-Puy and Saulcet localities (De Pietri et al., 2020a), both of which fall within the MN2 zone at the latest (De Pietri et al., 2011a,b), corresponding to a minimum age of 20.0 Ma (see the discussion of the age of Calibration 11). In a recent reassessment, De Pietri et al. (2020a) referred the right humerus that constitutes the holotype of *Becassius* to Glareolidae based on a combination of 9 characters which exhibit a highly homoplastic distribution within Charadriiformes, and which occur together in some but not all glareolids. None of the characters was unique to Glareolidae among the Lari, and at least five were also present among the Scolopaci. A combination of five additional characters ascertained from referred coracoids was also reported to support a glareolid affinity for *Becassius*, but this referral was only based on the size and relative abundance of the elements (De Pietri et al., 2020a). Given the homoplastic distribution of the relevant traits among shorebirds, and the absence of a phylogenetic analysis polarizing the corresponding character state transitions by means of outgroup comparison, it is difficult to establish which (if any) represent glareolid apomorphies. The evidence linking *Becassius* to Glareolidae therefore fails to satisfy the criteria outlined by Parham et al. (2011), precluding the taxon from calibrating the (Glareolidae + *Dromas*) node.

***Boutersemia* spp.**

**Potential node calibrated.** Crown-group Glareoloidea (MRCA of *Glareola pratincola* and *Dromas ardeola*).

**Fossil taxa.** *Boutersemia belgica* Mayr and Smith 2001 and *Boutersemia parvula* Mayr and Smith 2001.

**Specimens.** IRScNB Av 41 (holotype of *B. belgica*; Mayr and Smith, 2001) and IRScNB Av 46 (holotype of *B. parvula*; Mayr and Smith, 2001), Institut Royal des Sciences Naturelles de Bruxelles, Brussels, Belgium.

**Lower bound.** 32.02 Ma.

**Discussion.** *Boutersemia* was referred to the Glareolidae on the basis of the following two apomorphies of the tarsometatarsus: foramen vasculare distale (1) very large and (2) situated on the bottom of a well-developed, broad groove. Both characters have a homoplastic distribution among shorebirds and are also found in the Charadriidae, but *Boutersemia* can be distinguished from the latter by its retention of the plesiomorphically distinct fossa metatarsi I (Mayr and Smith, 2001). However, the referral to the Glareolidae was considered tentative by the authors, and was not supported when *B. belgica* was included in the parsimony and Bayesian phylogenetic analyses of Smith and Clarke (2015), both of which found it to be a jacanid. This alternative position is still relevant to calibration design, since *Boutersemia* would then constitute the earliest record of Jacanidae. The fossils of both *B. belgica* and *B. parvula* were collected from the Boutersem Sand Member of the Borgloon Formation, whose dating was recently reviewed by Mayr et al. (2019a). The unit can be correlated to calcareous nannofossil zone NP22 (32.92–32.02 Ma; Anthonissen and Ogg, 2012) based on dinocysts collected from the area, and to European reference level MP21 based on its mammalian fauna. The latter assignment would imply a slightly older minimum age of 32.6 Ma (Becker, 2009; see also Vandenberghe et al., 2012, Figure 28.10), which also agrees with the date assigned to the top of the Ruisbroek Member (equivalent to Boutersem Sand; Mayr et al., 2019a) by Hooker et al. (2009, Figure 3), although both Becker (2009) and Hooker et al. (2009) give a duration for the NP22 zone in their correlation charts that differs from the more recent estimate of Anthonissen and Ogg (2012). The minimum age of 32.6 Ma was also used in a previous divergence time analysis for a calibration based on a different Boutersem avian fossil (Stervander et al., 2019). However, even with the more conservative lower bound of 32.02 Ma, *Boutersemia* would still supersede *Nupharanassa tolutaria* (see above) as the oldest known jacanoid. Nevertheless, we do not regard the results of Smith and Clarke (2015) as sufficiently robust to allow using *Boutersemia* to calibrate the jacanid-rostratulid split. *B. belgica* was the least complete taxon scored by Smith and Clarke (2015), with 98.3% missing data, leaving only 6 characters in which the fossil was identical to the extant jacana *Hydrophasianus chirurgis* (Smith, 2011a). Most recently, De Pietri et al. (2020a) corroborated the referral of *Boutersemia* to Glareolidae, considering it to be more robust than originally suggested by Mayr and Smith (2001). However, this conclusion was based on overall morphology rather than a list of apomorphies, and since it contradicts the only phylogenetic analysis performed so far, we consider the affinities of *Boutersemia* uncertain.

##### *Eocliffia primaeva*

**Potential node calibrated.** Total-group Turnicidae (MRCA of *Turnix sylvaticus* and *Larus marinus*).

**Fossil taxon.** *Eocliffia primaeva* Mourer-Chauviré et al. 2017.

**Specimen.** EC7 Ep1 (holotype; Mourer-Chauviré et al., 2017) and EC7 Ep4 (paratype consisting of a right tarsometatarsus; Mourer-Chauviré et al., 2017), Geological Survey of Namibia, Windhoek, Namibia.

**Lower bound.** 18.7 Ma.

**Discussion.** Synapomorphies of the Turnicidae mainly pertain to the coracoid (Mayr and Knopf, 2007), which is unknown for *Eocliffia*, and Mourer-Chauviré et al. (2017) support the turnicid affinities of the taxon on the basis of overall comparisons rather than an explicit list of shared derived characters. However, two turnicid apomorphies can be identified from the description: (1) a hypotarsus with an enclosed bony canal for the tendon of the musculus flexor digitorum longus (Mayr, 2011b), which can be ascertained from specimen EC7 Ep4, and (2) the carpal trochlea with a subcircular outline (Zelenkov et al., 2016), which can be ascertained from the holotype. The former character is potentially problematic, as it is highly homoplastic within Charadriiformes and also occurs in *Pluvianus*, Burhinidae, and Scolopaci (Mayr, 2011b); given the early-diverging positions of *Pluvianus* and Burhinidae within Charadrii and of Turnicidae within Lari, it could even constitute a charadriiform symplesiomorphy. In contrast, the subcircular ventral outline of the trochlea carpalis is otherwise only present in the distantly related Burhinidae (Zelenkov et al., 2016). While *Eocliffia* has not been included in a phylogenetic analysis, it has never been linked to a clade other than buttonquails, and other paleornithologists have considered its relationship to the Turnicidae to be sufficiently robust to use it as a calibration (De Pietri et al., 2020b).

While the buttonquail affinities of *Eocliffia* may be sufficiently well-supported to permit its use as a calibration, the fossiliferous limestones of the Sperrgebiet are too poorly constrained for the taxon to supersede *Turnipax oechslerorum* (see below) as the oldest known stem-turnicid. Mourer-Chauviré et al. (2017) consider the locality to be Bartonian, in accordance with Pickford et al. (2014) and Pickford (2015). This is based on evidence from rodent fauna, which appears to be more "primitive" at Eocliff than at Quarry BQ-2 of the Birket Qarun Formation (Fayum, Egypt), a locality correlated to calcareous nannoplankton zone NP19–20. Moreover, Eocliff contains the same rodent taxa as the nearby Silica North locality, which is unconformably overlain by the marine Langental beds also dated to NP19–20 (Dauteuil et al., 2018; Godinot et al., 2018). Morales and Pickford (2018) allowed for the possibility that given the uncertainty of the Quarry BQ-2 correlation, Eocliff might be Priabonian rather than Bartonian, but no younger. However, this was doubted by Sallam and Seiffert (2016), whose tip-dating analysis favored a late Oligocene age for the Silica North rodent *Prepomonomys bogenfelsi* that was also recorded from Eocliff (Pickford, 2018, Table 1). The authors further noted that the presence of an advanced anthracotheriid as well as tenrecoids intermediate between late Eocene forms from Fayum and early Miocene species suggested the site could not be older than latest Priabonian, and possible as young as late Oligocene. Sallam and Seiffert (2019) expanded on these findings by showing that the posterior tip age distributions of both *P. bogenfelsi* and another Eocliff rodent were strongly concentrated in the earliest Miocene despite the use of broad uniform tip age priors, and that Bayes factors provided very strong evidence in favor of this dating relative to placing both taxa in the Bartonian. This is also consistent with the dating proposed by Marivaux et al. (2014), who suggested a Miocene age for the Silica North and Silica South localities (assumed to be contemporaneous with Eocliff; Pickford et al., 2014).

In the absence of stratigraphic information more decisive than mammalian faunal correlations, we use the lower bound of the 95% highest posterior interval about the tip age of *Prepomonomys bogenfelsi* (22.3–18.7 Ma) from the analysis by Sallam and Seiffert (2019) as the strict minimum age of *Eocliffia*. Given the resulting extremely broad range of possible dates (late middle Eocene to early Miocene; 41.2–18.7 Ma), there is no firm evidence that *Eocliffia* predates the early Oligocene *Turnipax oechslerorum*, and the taxon should not be used as a calibration (*contra* De Pietri et al., 2020b).

##### *Laricola elegans*

**Potential node calibrated.** Total-group Laridae (MRCA of *Larus marinus* and *Alca torda*).

**Fossil taxon.** *Laricola elegans* (Milne-Edwards, 1868).

**Specimen.** MNHN Av.4134 (lectotype of *Laricola elegans*; De Pietri et al., 2011a), Muséum national d'histoire naturelle, Paris, France.

**Lower bound.** 20.0 Ma.

**Discussion.** Smith (2011a, Figure 8.5) found *L. elegans* to be indistinguishable from *Larus marinus* in the characters the former could be coded for, and therefore deeply nested within Laridae. However, De Pietri et al. (2011a) referred new cranial material to the taxon and included the resulting combined OTU in a phylogenetic analysis that yielded two most parsimonious trees, which recovered *L. elegans* either within the crown or on the stem of "Laromorphae" (= Laridae *sensu* Boyd, 2019). Given this phylogenetic uncertainty, *L. elegans* can only reliably calibrate the larid total group, i.e., the split between Laridae and Alcoidea, as also noted by Kimball et al. (2019). However, this renders the resulting calibration redundant with respect to Calibration 8, which is older and calibrates a more deeply nested node. This holds true regardless of whether the minimum age of *L. elegans* is more conservatively based on that of the Saint-Gérand-le-Puy area in general (here assumed to be 20.0 Ma; see above), a possibility favored by Kimball et al. (2019) based on the lack of information about the exact provenance of the *L. elegans* fossils described by De Pietri et al. (2011a), or on the age of layer Créchy 1–2 in particular (Smith, 2015), from which remains of *L. elegans* were reported by Mourer-Chauviré et al. (2004). The latter site was correlated by Hugueney et al. (2003) to the Swiss locality Brochene Fluh 53, spanning chrons C6Cn.2r through C6Cn.1r (Kempf et al., 1999) and thus a time period of 23.23–22.75 Ma (Ogg, 2012). As even this older age estimate is substantially younger than the more deeply nested Calibration 8, we leave both the crown group and the total group of Laridae uncalibrated.

***Turnipax oechslerorum***

**Potential node calibrated.** Total-group Turnicidae (MRCA of *Turnix sylvaticus* and *Larus marinus*).

**Fossil taxon.** *Turnipax oechslerorum* Mayr and Knopf 2007.

**Specimen.** SMF Av 506a+b (holotype; Mayr and Knopf, 2007), Forschungsinstitut Senckenberg, Frankfurt am Main, Germany.

**Lower bound.** 29.62 Ma.

**Discussion.** *T. oechslerorum* represents the older of the two species of *Turnipax* and can be referred to the Turnicidae based on two coracoid apomorphies (Mayr and Knopf, 2007); note that the other six characters listed by (Smith, 2015) as turnicid apomorphies in fact represent plesiomorphies and autapomorphies distinguishing Turnipax from extant buttonquails. While the phylogenetic position of the taxon is sufficiently well-established to permit its use as a calibration, its age renders it redundant with respect to Calibration 8. The known material of *T. oechslerorum* derives from the Grube Unterfeld clay pit (formerly referred to as Frauenweiler), which can be correlated to calcareous nannoplankton zone NP23 (Maxwell et al., 2016; Micklich et al., 2016), corresponding to an age of 32.02–29.62 Ma (Anthonissen and Ogg, 2012). *T. oechslerorum* is thus substantially younger than GCVP 5690, dated to zone NP19–20. Since the divergence between the Turnicidae and the rest of the Lari necessarily predates the scolopacid–alcid split in our tree, a calibration based on *T. oechslerorum* would be superfluous.

***Vanolimicola longihallucis***

**Potential node calibrated.** Crown-group Jacanoidea (MRCA of *Jacana jacana* and *Rostratula benghalensis*).

**Fossil taxon.** *Vanolimicola longihallucis* Mayr 2017b.

**Specimen.** SMNK.PAL 8683a+b (holotype; Mayr, 2017b), Staatliches Museum für Naturkunde, Karlsruhe, Germany.

**Lower bound.** 47.41 Ma.

**Discussion.** *Vanolimicola* is known from a single poorly preserved partial skeleton found in the lacustrine deposits of Grube Messel near Darmstadt, Germany (Mayr, 2017b). The fossil belongs to a small, long-legged wading bird with a needle-like bill, and its morphology generally agrees with extant Charadriiformes, which would have significant implications for calibration design given the early to middle Eocene age of the material. The dating of the Messel site was recently reviewed by Lenz et al. (2015), who obtained $^{40}$Ar/$^{39}$Ar ages of 48.27 $\pm$ 0.22 Ma and 48.11 $\pm$ 0.22 Ma (depending on the value used for the Fish Canyon sanidine standard) for a basaltic clast from a tuff layer below the first lacustrine sediments. Along with the estimated duration of 640 kyr for the main fossil-bearing oil shale deposits ("Middle Messel Formation"; MMF) and the calibration of the site's high-resolution palynological record to astronomical solutions, these data indicate an age of 47.61 Ma (La2010d solution) or 47.41 Ma (La2010a solution) for the top of the MMF. Lenz et al. (2015) considered the latter value to be more likely, which is also consistent with the recommendation of Parham et al. (2011) that the youngest plausible date be used when designing age constraints.

In his description of the taxon, Mayr (2017b) noted several features that link *Vanolimicola* with jacanas, including a very long hallux and the humerus with a weakly developed processus supracondylaris dorsalis. If these characters proved to be phylogenetically informative, *Vanolimicola* would represent the earliest known named charadriiform species and the first appearance of Jacanidae in the fossil record, exceeding the age of the next oldest known jacanid, the early Oligocene *Nupharanassa tolutaria* (see Calibration 12), by more than 50%. However, Mayr (2017b) considered the referral of *Vanolimicola* to Jacanidae to be only poorly supported, and noted similarities between *Vanolimicola* and *Songzia*, a taxon which has been repeatedly recovered as a crown-group gruiform by recent phylogenetic analyses (Musser et al., 2019; Musser and Clarke, 2020; see also our re-analyses of the Musser and Clarke, 2020 dataset in Figure A.10). Accordingly, we consider *Vanolimicola* to be of uncertain phylogenetic position, and thus unsuitable for use as a calibration.

##### *Wilaru tedfordi*

**Potential node calibrated.** Crown-group Burhinidae (MRCA of "*Esacus*" *magnirostris* and *Burhinus bistriatus*).

**Fossil taxon.** *Wilaru tedfordi* Boles et al. 2013.

**Specimen.** SAM P48925 (holotype; Boles et al., 2013), South Australian Museum, Adelaide, Australia.

**Lower bound.** 24.76 Ma.

**Discussion.** In the original description, *Wilaru* was referred to the Burhinidae based on a number of characters pertaining to the humerus, coracoid, and femur, although without explicitly indicating that these were supposed to be apomorphic for the clade (Boles et al., 2013). The authors also reported several traits shared by *Wilaru* and the extant genus "*Esacus*" to the exclusion of *Burhinus* (humerus with a second, dorsally located fossa pneumotripicitalis; scapula with a laterally folded facies articularis clavicularis; carpometacarpus with a fossa infratrochlearis that is elongated along the proximocranial-distocaudal axis rather than round; and a laterally deeper hypotarsus) while noting that their phylogenetic importance was unclear. If interpreted as evidence for a sister-group relationship between the two taxa, these characters would support a position of *Wilaru* within the crown-group Burhinidae. Note that while our total-evidence topology shows "*Esacus*" to be nested within *Burhinus* (Figure A.9), this would strengthen rather than contradict the inclusion of *Wilaru* within the burhinid crown clade. In contrast, Claramunt and Cracraft (2015, Table S1) opted for a more conservative treatment of the taxon as a total-group burhinid, and accordingly used it to calibrate Chionida as a whole. Depending on which of these two options is followed, *Wilaru* would either complement or supersede the coeval *Chionoides australiensis* (Calibration 15). The fossils of *W. tedfordi* were recovered from the Pinpa Local Fauna of the Namba Formation as well as faunal zone B of the Etadunna Formation. The former unit is older, as Woodburne et al. (1994) correlated it with Etadunna Formation faunal zone A, which yielded the fossils of *C. australiensis* (see above), and this correlation has been followed by all recent studies (Beck et al., 2020; Thorn et al., 2021; note that the latter study gives an age of 25.7–25.5 Ma for Etadunna Formation faunal zone A based on the original dating of Woodburne et al., 1994, as opposed to the more recent magnetostratigraphic evidence of Megirian et al., 2010 used here).

More recently, however, De Pietri et al. (2016b) reinterpreted the taxon as a late-surviving presbyornithid (i.e., a representative of the total-group Anseriformes) based on a combination of 15 morphological characters, and showed that much of the character evidence used by Boles et al. (2013) to support a burhinid and charadriiform affinity for the taxon was erroneous, including the presence of the second fossa pneumotripicitalis. This reassessment was subsequently corroborated by a number of phylogenetic analyses, although all of these were based on a single character matrix that did not include charadriiforms (Worthy et al., 2017; Tambussi et al., 2019; Agnolín, 2021). In accordance with these results, we consider *Wilaru* to be a pan-anseriform, and exclude it from our calibration set.

#### 2 Outgroup sequences

Except for Calibration 16, whose soft upper bound was set to 66.0 Ma to reflect our strong prior belief that crown-group charadriiforms did not originate before the K–Pg boundary, all of our calibrations employ soft upper bounds calculated using the outgroup-based Bayesian algorithm of Hedman (2010). Our approach to outgroup sequence construction generally followed the protocol introduced by Friedman et al. (2013) and elaborated by subsequent studies (e.g., Alfaro et al., 2018). Accordingly, our sequences are restricted to "stratigraphically consistent" outgroups, each of which is older than or of the same age as the immediately preceding outgroup, but younger than or equal in age to the next outgroup in the sequence. Similar to Alfaro et al. (2018), we considered four classes of eligible outgroups: (1) taxa inside the group of interest (here, Charadriiformes) that were themselves used as primary calibrations (i.e., minimum age constraints); (2) taxa inside the group of interest that were not employed as primary calibrations; (3) a sequence of taxa outside the group of interest yielding a stratigraphically consistent pattern of first appearance dates; and (4) a single hard maximum $t_0$ believed to predate all of the divergences under consideration. Charadriiform outgroups from classes (1) and (2) are specific to each primary calibration, and as such are listed directly below each of the 15 non-root calibrations in Section 1.1. Non-charadriiform constraints from classes (3) and (4) form a fixed series of ages appended to each of the charadriiform sequences, and are reported below.

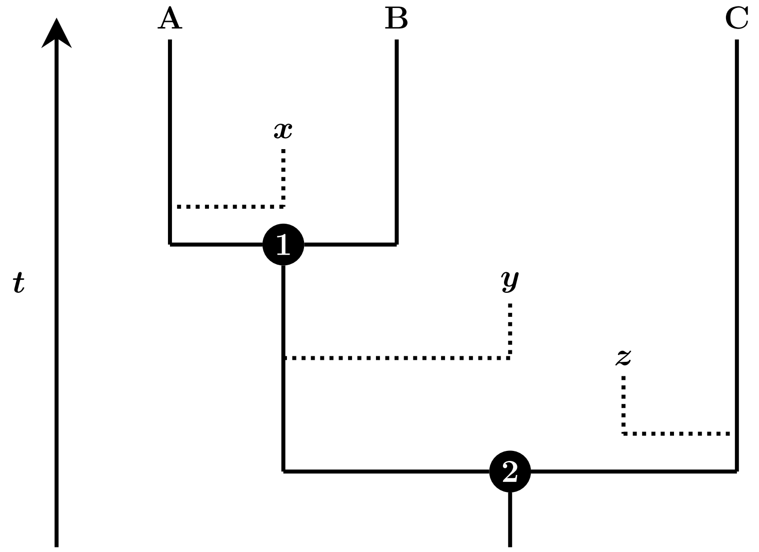

**Figure A.1:** A phylogeny of extant taxa (A, B, C) and several fossils ($x$, $y$, $z$) whose position along the time axis indicates relative age. The oldest fossil within a given clade calibrates its initial split. Fossil $x$ constrains the split between A and B (calibration 1); $z$ constrains the split between (A + B) and C (calibration 2). Fossil $y$ cannot be used as a calibration, as it is superseded by the older fossil $z$, but it can still serve as an outgroup to calibration 1.

We attempted to include as many informative fossils as possible in the set of primary calibrations (1), leaving only a few fossils as possible members of class (2). Nevertheless, there are cases in which a fossil will not be eligible for inclusion in the set of primary calibrations, but can still provide valuable information when included in the outgroup sequence. Consider the pair of nested calibrations depicted in Figure A.1. Two fossils are available to calibrate the split between taxa (A + B) and C: $y$, belonging to the stem group of (A + B), and $z$, belonging to the stem group of C. Since $y$ is younger than $z$, it will be discarded as redundant, and the split will be calibrated by $z$ instead (calibration 2). However, since $y$ is more closely related to (A + B) than $x$ is, it can still be used in the outgroup sequence of the calibration assigned to that node (calibration 1, based on $x$), in conjunction with $z$. Accordingly, when constructing our outgroup sequences, we also considered those fossils that were excluded from the primary calibration set due to redundancy (*Laricola elegans* and *Turnipax oechslerorum*; see Section 1.2). However, these did not prove to be stratigraphically consistent, and our analysis ended up including no class (2) outgroups.

Nested pairs of primary calibrations can be problematic when the "outer" calibration is well-established as a member of the more inclusive clade, but its exact position therein remains unknown. The former allows it to be used as a calibration in its own right, but for the taxon to serve as a class (1) outgroup to the more deeply nested ("inner") calibration, the latter information is required. This problem affected Calibration 8, Calibration 11, and Calibration 16, and we addressed it on a case-specific basis. Specimen GCVP 5690 (Calibration 8) was found within crown-group Alcinae in a phylogenetic analysis by Smith (2011a), who nevertheless considered the polarization of the single character supporting this placement to be unclear, and preferred to treat the fossil as a total-group Alcidae *incertae sedis*. For outgroup construction purposes, we effectively assumed it to be a stem-alcid, capable of serving as an outgroup both to the primary calibrations within Alcinae (Calibrations 1–5) and to those within Fraterculinae (Calibrations 6, 7). *Elorius* and *Parvelorius* (Calibration 11) have never been included in a phylogenetic analysis, but extensive comparisons suggest that they most closely resemble the early-diverging *Numenius* and *Limosa*, and humeral plesiomorphies indicate their exclusion from the clade comprising Scolopacinae, Tringinae, and Arenariinae (De Pietri and Mayr, 2012). Accordingly, we considered them capable of serving as an outgroup to the arenariine Calibration 9 and the scolopacine Calibration 10. Finally, specimen IGM 100/1435 (Calibration 16) was strongly supported as a crown-group charadriiform and, additionally, weakly supported as a member of Chionida in our re-analyses of the Musser and Clarke (2020) character matrix (Figure A.10). We followed the weakly supported result during outgroup sequence construction, and additionally chose to assume that IGM 100/1435 was less closely related to extant Chionidae than *Chionoides* (Calibration 15). We were thus able to use Calibration 16 as a class (1) outgroup to all other primary calibrations, including those within Lari (Calibrations 1–8), Scolopaci (Calibrations 9–13), Charadriida (Calibration 14), as well as Calibration 15.

Due to the unique phylogenetic context of this study, we further chose to split Alfaro et al.'s (2018) third class of outgroups into two categories. Charadriiform outgroups among Neornithes (crown birds) comprise a limited number of candidate lineages, and determining the first appearance date of any given lineage requires a thorough review of the paleontological literature that employs qualitative assignments to extant clades more often than formal phylogenetic analyses. In contrast, outside of Neornithes (within the bird stem group), new lineages are constantly described and their relationships tested using explicit phylogenetic analyses, which, however, differ from one another with respect not only to the resulting topologies but also taxon sampling. As a result, quite apart from topological conflict, constructing a single consensus outgroup sequence is nearly impossible because some of the potentially relevant taxa have never been included in a single analysis together, and their placement in the sequence vis-à-vis each other cannot be established. We dealt with this issue by considering neornithine and non-neornithine outgroups separately, and by using multiple recent densely sampled phylogenies for the latter. This required evaluating a much larger number of outgroups than has been the case in previous studies (60 compared to 16 in Friedman et al., 2013 and 22 in Alfaro et al., 2018) despite the resulting sequences being shorter (minimum length = 8 outgroups, compared to 13 in Friedman et al., 2013 and 10 in Alfaro et al., 2018). As demonstrated by Hedman (2010), five stratigraphically consistent outgroups are sufficient for robust estimates.

In summary, the soft upper bounds of all non-root calibrations were calculated using a combination of (1) a calibration-specific sequence of other primary calibrations ($n = 15$), with (2) several additional charadriiform taxa evaluated but ultimately not used ($n = 0$); (3$a$) neornithine outgroups common to all primary calibrations ($n = 2$); (3$b$) non-neornithine outgroups arranged in multiple alternative sequences, each of which was appended to the base sequence comprising (1) through (3$a$) ($n = 58$); and (4) a single hard upper bound on all calculations ($n = 1$), set equal to 160 Ma here. This approximately corresponds to the age of the Tiaojishan Formation, which yielded the oldest known dinosaur fossils preserving pennaceous feathers (Chu et al., 2016), and predates even the oldest previous molecular estimates for the origin of Charadriiformes by $> 50$ Myr. In practice, estimated ages are relatively insensitive to this maximum value (Hedman, 2010).

#### 2.1 Neornithine outgroups

Although the Charadriiformes are deeply nested within crown-group birds (Neornithes *sensu* Cracraft, 1986) (Cracraft, 1988; Sibley and Ahlquist, 1990; Mayr and Clarke, 2003; Livezey and Zusi, 2007; Hackett et al., 2008; Jarvis et al., 2014; Prum et al., 2015), the number of neornithine fossils that can be incorporated into the outgroup sequences of charadriiform calibrations is relatively low. This is mainly due to the uncertain interrelationships among the major lineages of Neoaves – a clade that includes all neornithines other than gamefowl (Galliformes), waterfowl (Anseriformes), and ostriches and kin (Palaeognathae) – and the fact that there are relatively few neornithine fossils from the Paleocene or the Cretaceous (i.e., predating the earliest known crown-group charadriiform occurrence from the earliest Eocene) that can be confidently assigned to specific extant lineages (Mayr, 2014, 2017a). The oldest known neornithine fossils, and the only ones to reliably predate the K–Pg boundary, belong to the Galloanserae (gamefowl and waterfowl) rather than the earlier-diverging Palaeognathae, whose earliest representatives date from either the latest Maastrichtian or the earliest Paleocene (Parris and Hope, 2002; Nesbitt and Clarke, 2016). Given our decision to exclude stratigraphically inconsistent outgroups, this precludes Palaeognathae from contributing to the outgroup sequences considered here. Although the temporal gap between the oldest known galloanserans from the early Maastrichtian (see below) and the oldest known charadriiforms from the Paleocene–Eocene boundary (see above) may be bridged by a variety of neoavian taxa (e.g., *Berruornis*, *Gradiornis*, *Lithoptila*, *Neogaeornis*, *Novacaesareala*, *Palaeotringa*, *Polarornis*, *Protoplotus*, *Qianshanornis*, *Telmatornis*, *Tytthostonyx*, *Walbeckornis*; Hope, 2002; Mayr, 2014, 2017a), these are nearly always poorly constrained phylogenetically and, in some cases, stratigraphically. Moreover, even if the fossils in question could be accurately dated and reliably attributed to specific neaovian subclades, the persistent uncertainty about higher-level neoavian phylogeny (Suh, 2016; Reddy et al., 2017; Houde et al., 2019) makes it unclear how many distinct positions in an outgroup sequence they could occupy without rendering each other redundant due to stratigraphic inconsistencies. Here, we thus take a conservative approach and select only two neornithine outgroups to be appended to all charadriiform calibrations: one from Neoaves and another one from Galloanserae, listed in this order below.

##### Additional outgroup 1

**Fossil taxon.** *Tsidiiyazhi abini* Ksepka et al. 2017.

**Specimen.** NMMNH P-54128 (holotype; Ksepka et al., 2017), New Mexico Museum of Natural History and Science, Albuquerque, NM, USA.

**Lower bound.** 62.221 Ma.

**Phylogenetic justification.** Based on a constrained phylogenetic analysis of 111 morphological characters, Ksepka et al. (2017) found the taxon to be a sandcoleid, i.e., a stem-group member of the Coliiformes (mousebirds). Mayr (2018) opined that the phylogenetic position of *Tsidiiyazhi* required further study, and Mayr et al. (2019b) compared it to messelasturids.

However, this alternative phylogenetic position would still fall within Neoaves and, even more narrowly, Telluraves (landbirds), and as such would not affect the validity of treating *Tsidiiyazhi* as a neoavian outgroup to the Charadriiformes.

**Age justification.** A precise dating of the locality that yielded the type material of *T. abini* was provided in the original description of the taxon (Ksepka et al., 2017), and is followed here without modifications. The site can be constrained to magnetochron C27n, corresponding to an age of 62.517–62.221 Ma according to the geomagnetic time scale of Ogg (2012).

**Discussion.** We consider *Tsidiiyazhi abini* to represent the oldest well-established non-charadriiform neoavian, superseding the mid-Paleocene stem-penguin *Waimanu manneringi* from the Waipara Greensand of New Zealand, which has seen extensive use in previous node-dating studies (Slack et al., 2006; Pacheco et al., 2011; Gibb et al., 2013; Claramunt and Cracraft, 2015; Prum et al., 2015; Boast et al., 2019). Given the doubts expressed about the mousebird affinities of *Tsidiiyazhi* (see above), *Waimanu* still represents the oldest known neoavian that can be uncontroversially linked to a specific "order-level" clade. However, its fossils are younger than those of *Tsidiiyazhi*, rendering it unsuitable as an alternative or additional outgroup. An age of 61.6–60.5 Ma is commonly cited for the taxon (Mayr and Scofield, 2014; Ksepka and Clarke, 2015; Blokland et al., 2019). A slightly different but consistent date of "about 61" Ma (early late Teurian stage) is given by Mayr et al. (2017b) for fossils found 11 m above the *W. manneringi* type locality, and by Mayr et al. (2017a) for fossils found 13 m above it. Blokland et al. (2019) expand these age ranges to 61.5–59 Ma. The first Waipara fossil described from strata older than the type horizon of *W. manneringi* is the pseudodontorn *Protodontopteryx ruthae*, whose fossils were found at least 3 m below the *W. manneringi* holotype and were estimated to be 62–61.5 Myr old (Mayr et al., 2021). Perhaps the most accurate estimate of the age of *W. manneringi* is given by Benton et al. (2015) based on the correlation between revised dinoflagellate zonation and magnetostratigraphy; their conservative minimum is 60.2 Ma. Regardless of the exact date used, *W. manneringi* is uncontroversially younger than *T. abini*, and hence considered here to be superseded by the latter taxon.

Under several phylogenetic hypotheses, both *W. manneringi* and *T. abini* could be used as successive outgroups to the Charadriiformes. Specifically, this would require shorebirds to be more closely related to penguins than to landbirds (the "Aequorlitornithes hypothesis"; Prum et al., 2015), a relationship only recovered in Prum et al. (2015) as well as some of the secondary analyses of Jarvis et al. (2014; see their Figs. 4C,D and S13), and suggested to be driven by the use of protein-coding sequences that artificially unites taxa with similar life history traits (Jarvis et al., 2014; Reddy et al., 2017). In contrast, both *Tsidiiyazhi* and *Waimanu* would represent the same outgroup lineage if landbirds and penguins were more closely related to each other than to shorebirds (Jarvis et al., 2014; Kuhl et al., 2020), with *T. abini* superseding *W. manneringi* as the oldest known representative of such a clade. Similarly, if shorebirds were more closely related to landbirds than to penguins (the "Litoritelluraves hypothesis"; Yuri et al., 2013), as suggested by Hackett et al. (2008), Kimball et al. (2013), and Reddy et al. (2017), the two taxa could potentially occupy two distinct positions in the same outgroup sequence, but the more closely related and older *T. abini* would still supersede the more distant but younger *W. manneringi*. Currently, it is unclear which of these three hypotheses is correct, and there is a non-negligible chance that landbirds, shorebirds, and a waterbird clade including penguins represent constituent lineages of an irresolvable, "hard" polytomy at or close to the base of Neoaves (Suh, 2016; Houde et al., 2019).

As the earliest known representative of Neoaves, *Tsidiiyazhi* may itself be superseded by *Tytthostonyx glauconiticus* Olson and Parris 1987 from the lower part of the Hornerstown Formation of New Jersey. The age of this unit is notoriously poorly constrained (Ksepka et al., 2017; Mayr, 2017a), with the latest Maastrichtian or the earliest Paleocene both considered possible (Olson and Parris, 1987; Parris and Hope, 2002). Benton et al. (2015) favored the latter option, which has been followed by later studies (Maisch, 2020). A recent detailed review by Wiest et al. (2016) concluded that the K–Pg boundary may lie immediately below, immediately above, or even within the "Main Fossiliferous Layer" (MFL) of the basal Hornerstown Formation, suggesting that the numerical age of 66.0 $\pm$ 0.1 Ma assigned to the relevant fossils by Benton et al. (2015) is appropriate. Regardless of stratigraphic uncertainty, there is therefore little doubt that *Tytthostonyx* substantially predates *Tsidiiyazhi*. However, the phylogenetic position of the taxon remains uncertain. Originally suggested to be a procellariiform (Olson and Parris, 1987), it was reinterpreted as a possible tropicbird by Bourdon et al. (2008); however, Mayr (2015) and Mayr and Scofield (2016) tentatively supported the original assignment to Procellariiformes based on a single apomorphy (distal humerus with an absent sulcus scapulotricipitalis). While both positions are compatible with its use here, and while the broader neoavian affinities of *Tytthostonyx* have never been doubted, neither have they been supported by a formal phylogenetic analysis or an explicit list of apomorphies – a fact that is unsurprising given the fragmentary nature of the material (Olson and Parris, 1987) and the paucity of neoavian morphological synapomorphies (Mayr, 2011a). Consequently, most reviews conservatively concluded that the taxon is in need of further study (Mayr, 2009; Kaiser and Dyke, 2011), and calibration compendia as well as node-dating analyses have implicitly (Prum et al., 2015; Kimball et al., 2019) or explicitly (Benton et al., 2015; Ksepka and Clarke, 2015; Smith and Ksepka, 2015) avoided its use as an age constraint.

As a result, we regard *T. abini* as the only taxon that (1) has well-established neoavian affinities; (2) predates the earliest known fossil remains attributable to the Charadriiformes; and (3) is not superseded by any other taxa satisfying conditions (1) and (2) under any plausible phylogenetic hypothesis for neoavian interrelationships.

**Additional outgroup 2**

**Fossil taxon.** *Teviornis gobiensis* Kurochkin et al. 2002.

**Specimen.** PIN 44991-1 (holotype; Kurochkin et al., 2002), Paleontological Institute of the Russian Academy of Sciences, Moscow, Russia.

**Lower bound.** 69 Ma.

**Phylogenetic justification.** *Teviornis* was originally described as an anseriform based on a single apomorphy identified by a previous phylogenetic analysis, and as a presbyornithid based on a unique combination of three characters of the carpometacarpus (Kurochkin et al., 2002). However, this referral was questioned by Clarke and Norell (2004), who pointed out that the taxon lacked the synapomorphies of Neornithes, Neognathae, and Galloanserae, and that the single character supporting its anseriform affinities was plesiomorphic when a broader taxon sample was considered. This skepticism about the position of *Teviornis* within Anseriformes or even Neornithes was echoed by most subsequent studies (O'Connor et al., 2011; Ksepka and Phillips, 2015; Braun et al., 2019), but recent detailed reassessments of the taxon corroborated its inclusion in the waterfowl total group (Zelenkov and Kurochkin, 2015; De Pietri et al., 2016b). In particular, De Pietri et al. (2016b) explicitly addressed the arguments of Clarke and Norell (2004) and provided four more characters linking the taxon to presbyornithids. Most importantly, *Teviornis* was recovered as a pan-anseriform in a phylogenetic analysis of 700 characters by Hartman et al. (2019). In contrast, there is no phylogenetic analysis supporting an alternative position for the taxon; the only other analysis including *Teviornis* of which we are aware, Cau et al. (2017, Extended Data Figure 9), found it in a large ornithuromorph polytomy that could only be resolved upon its removal.

**Age justification.** The only known material of *Teviornis* derives from the Guriliin (= Gurilyn) Tsav locality of the lower Nemegt Formation in the Nemegt Basin, Ömnögovi Province, Mongolia (Kurochkin et al., 2002). Unfortunately, the age of the Nemegt Formation is poorly constrained (Weishampel et al., 2008; Ksepka and Clarke, 2015). Weishampel et al. (2008) cite a study (unavailable to us) that correlated a different locality within the Nemegt Basin (Hermiin Tsav) to chron C32n, and using this information as well as the assumption of lateral continuity, derive an age of 72.0–70.8 Ma for Bügiin Tsav, a site located 7 km from Guriliin Tsav and presumed to be coeval with it (Kurochkin et al., 2002). Following a more recent age range of 73.649–71.449 Ma for C32n (Ogg, 2012), the Nemegt Formation would span the Campanian–Maastrichtian boundary (72.1 ± 0.2 Ma), consistent with a number of studies considering it to be late Campanian through early Maastrichtian in age (see Lillegraven and McKenna, 1986 and references therein). In their calibration compendia, Benton and Donoghue (2006) and Benton et al. (2009) cited Lillegraven and McKenna (1986) in support of assigning the Nemegt Formation specifically to the early Maastrichtian; however, this is not unambiguously borne out by the latter study. The authors then converted this stratigraphic range into a numeric age range of 70.6 ± 0.6 to 69.6 ± 0.6 Ma (Benton and Donoghue, 2006; Benton et al., 2009) but gave no supporting references for these values. The upper bound corresponds to the Campanian–Maastrichtian boundary in the timescale of Ogg et al. (2004, Figure 19.1), but the source for the lower bound is unclear, especially since the lower and upper Maastrichtian are not formally recognized units with a well-defined substage boundary (Ogg et al., 2004; Radmacher et al., 2014). Nevertheless, both the stratigraphy and dating of Benton and Donoghue (2006) and Benton et al. (2009) are consistent with later studies, which have generally assigned the Nemegt Formation to the early Maastrichtian (Kim et al., 2018; Funston et al., 2020) and dated it at "approx. 69 Ma" (Zanno and Makovicky, 2013), 70–69 Ma (Ksepka and Clarke, 2015), or 71–69 Ma (Kim et al., 2018). While these values are likely just as arbitrary as the numeric ages cited by Benton and Donoghue (2006) and Benton et al. (2009), their general congruence motivates the use of 69 Ma as the lower bound on the age of *Teviornis*.

**Discussion.** Due to the presumed uncertainty about its neornithine affinities, *Teviornis* was not used as a calibration by any node-dating study we are aware of. Most of the recent divergence time analyses instead relied on the late Maastrichtian *Vegavis* (see Additional outgroup 54) or even younger fossils (*Waimanu manneringi* in the case of Prum et al., 2015) as the oldest known well-supported crown-group bird. Paradoxically, recent evidence suggests that the status of *Vegavis* as a neornithine is less certain than that of the early Maastrichtian *Teviornis*, indicating that the latter taxon is more suitable as a calibration not only on stratigraphic but also phylogenetic grounds. The deeply nested position of *Vegavis* within crown-group Anseriformes inferred by Clarke et al. (2005) was questioned early on by Mayr (2013), who regarded the taxon as a likely neornithine but pointed out that its sister-group relationship to Anatidae was supported by a single character complex known to be prone to homoplasy. Several subsequent analyses found *Vegavis* to be a stem-group rather than crown-group anseriform (Agnolín et al., 2017; Worthy et al., 2017; Tambussi et al., 2019; Agnolín, 2021), and Mayr et al. (2018) regarded even its position within Galloanserae as uncertain. Indeed, several phylogenetic analyses focusing on Mesozoic birds found *Vegavis* outside of the (*Gallus* + *Anas*) clade, and their failure to sample other crown birds left even its neornithine affinities ambiguous (O'Connor et al., 2011; Zheng et al., 2018). Most recently, McLachlan et al. (2017) and Field et al. (2020) presented well-sampled phylogenetic analyses that placed *Vegavis* outside of Neornithes, indicating its unsuitability as a calibration for molecular dating studies. In contrast, the inclusion of Presbyornithidae within total-group Anseriformes is supported by all recent analyses (Clarke et al., 2005; Agnolín et al., 2017; Worthy et al., 2017; Tambussi et al., 2019; Field et al., 2020). Membership within the total group of Galloanserae is also well-established for the recently described *Asteriornis* (Field et al., 2020); at 66.7 Ma, this taxon is possibly slightly older than *Vegavis* but appreciably younger than *Teviornis*.

Several fossils even more ancient than *Teviornis* have been proposed to represent galloanserans or at least neornithines. *Austinornis lentus* is widely considered to be the oldest known taxon for which neornithine affinities are plausible (Mayr, 2009, 2014; Mitchell et al., 2015); it was recovered in a trichotomy with two crown-group galliforms in an analysis by Clarke (2004). However, *Austinornis* is known from a single tarsometatarsus fragment that could have only been scored for 9 out of the 202 characters used (95.5% missing data), of which just one was optimized as a galliform synapomorphy. No synapomorphies of more inclusive clades (Neornithes or Neognathae) could be ascertained from the specimen, and Clarke (2004, 149) cautioned against its use in molecular dating studies. Moreover, the specimen lacks exact provenance data Clarke (2004, 53), casting doubt on its Cretaceous age. The source for the age of 85 Ma commonly attributed to *Austinornis* (e.g., Mitchell et al., 2015; Braun et al., 2019) is also uncertain. Mitchell et al. (2015) cite Myers (2010) in support of this estimate, but as was pointed out by Cracraft et al. (2015), that study makes no mention of the taxon. However, it was cited in connection with *Austinornis* by Mayr (2014), and it does put the age of the Austin Chalk at 88–82 Ma (Myers, 2010, 1072); 85 Ma was probably chosen simply as the midpoint of that interval. Similar concerns also apply to putative neornithine material from the Late Cretaceous of Patagonia (Agnolín et al., 2006; Agnolín and Novas, 2012) and indeed to all pre-Maastrichtian remains once thought to belong to crown-group birds (Hope, 2002). Unlike *Austinornis*, these have either never been included in a formal phylogenetic analysis, or their inclusion in such failed to corroborate the hypothesized neornithine affinities. The latter scenario is exemplified by *Palintropus* (see Additional outgroup 45), considered to be a galliform by Hope (2002) but repeatedly found outside of Neornithes in later studies (Longrich et al., 2011; McLachlan et al., 2017; Hartman et al., 2019).

Following the reasoning above, we consider *Teviornis* to represent the earliest known taxon with well-established galloanseran affinities. After the completion of this study, this conclusion was seconded by Marjanović (2021), whose justification generally agrees with that given here. Based on unpublished magnetostratigraphic evidence and assumptions about the relative ages of the Nemegt and Baruungoyot Formations, Marjanović (2021) correlated the former unit to the lower half of chron C31 (approx. 71.4–69.9 Ma); his final recommended date of 71 Ma disregards the guidelines outlined by Parham et al. (2011), according to which the youngest plausible date should be applied to each minimum age constraint.

#### 2.2 Non-neornithine outgroups

To ensure that each calibration is associated with at least seven stratigraphically consistent outgroups, we extended the outgroup sequences beyond Neornithes into the avian stem group. Specifically, we used nine phylogenies extensively sampling non-neornithine representatives of Avialae, sourced from eight recent studies: McLachlan et al. (2017); Field et al. (2018); Zheng et al. (2018); Hartman et al. (2019); Kundrát et al. (2019); Cordes-Person et al. (2020); Wang et al. (2020b); Wang et al. (2020c). Stratigraphic ranges for the taxa represented in the topologies employed here were obtained from the literature and converted into numeric ages following detailed justifications reported below. Consistent with the treatment applied to other calibrations and outgroups, the youngest plausible date was assigned to each taxon following Parham et al. (2011).

Specimen numbers are generally not given for these additional outgroups, as this information is either redundant (many Mesozoic bird taxa are known only from the holotype) or unavailable, since the studies in question generally did not report which specimens their character codings were based on. An exception was made for unnamed taxa, or in cases where a single specimen is known to predate all other material assigned to the taxon. The phylogenetic justification entry reported for the previously listed calibrations is here supplanted by an explicit reference to a published topology; however, several assumptions about phylogeny were made in the process of converting tree topologies into outgroup sequences. Specifically, the suprageneric clades Confuciusornithiformes, Enantiornithes, and Hesperornithiformes were assumed to be monophyletic, as they have been recovered as such in all recent phylogenies we are aware of. Consequently, the oldest known species from each of these clades was chosen to represent it in the relevant outgroup sequence, even if this species was not itself sampled in a given tree. A similar assumption had to be made for tree tips which represented potentially polytypic genera but which were only referred to by their generic names (e.g., *Archaeopteryx*, *Jeholornis*, *Lithornis*), since the relevant studies did not always explicitly state whether they (1) considered the genera in question to be monotypic, (2) used composite terminals based on several species of the same genus, or (3) used only a single species for each terminal but omitted its specific epithet. In all such cases, our assumptions are explicitly stated and justified.

For the purposes of outgroup sequence construction, each topology was encoded as a list of taxa ranked by their phylogenetic distance from Neornithes, with "1" representing the immediate sister group of the avian crown clade. This list was extended as far from the crown as needed to obtain five stratigraphically consistent non-neornithine outgroups. Taxa that formed a clade to the exclusion of Neornithes were treated as having the same rank; similarly, an identical rank was assigned to all the constituent lineages of a polytomy. This treatment yielded preliminary outgroup sequences, from which the final sequences were obtained by selecting only the oldest taxon of a given rank. For clarity, both the preliminary and final outgroup sequence are listed for each tree topology. Finally, to obtain the final soft maximum ages (reported for each calibration in Section 1.1), we appended each of the nine 5-taxon non-neornithine outgroup sequences to the "base" sequence of each calibration (consisting of charadriiform outgroups where applicable, and of the two non-charadriiform neornithine outgroups listed above), and calculated the unweighted mean of the upper bounds of the resulting 95% credibility intervals.

All taxa that form a part of a preliminary outgroup sequence under at least one of the nine phylogenetic hypotheses are listed below in alphabetical order, with their corresponding numeric ages and age justifications. The sequences themselves are presented at the end of the list, with stratigraphically consistent entries of each preliminary sequence highlighted using gray shading. For the preliminary sequences, both the original taxon label (used in the tree figure on which the sequence is based) and a "reconciled" taxon label are given for each outgroup. The reconciled names correspond to the species listed below and are usually identical to the original labels up to the specific epithet, except for the three suprageneric clades listed above, for which the reconciled names are those of the oldest known representative(s).

**Additional outgroup 3**

**Fossil taxon.** *Ambiortus dementjevi* Kurochkin 1982.

**Minimum age.** 125.0 Ma.

**Age justification.** *Ambiortus* is known from the Khurilt Ulaan Bulag locality of the Andaikhudag Formation, Bayanhongor Province, Mongolia (O'Connor and Zelenkov, 2013), which can be dated to the Hauterivian–Barremian based on its mollusk, ostracod, fish, and insect fauna (Khand et al., 2000). This stratigraphic range has been converted here to a numeric age range following the latest version of the International Chronostratigraphic Chart (ICS v2020/03; `http://stratigraphy.org/ICSchart/ChronostratChart2020-03.pdf`).

**Additional outgroup 4**

**Fossil taxon.** *Antarcticavis capelambensis* Cordes-Person et al. 2020.

**Minimum age.** 70.8 Ma.

**Age justification.** The only known material of *Antarcticavis* derives from the Cape Lamb Member of the Snow Hill Island Formation at Cape Lamb, Vega Island, West Antarctica (Cordes-Person et al., 2020). The specimen was found approximately 20 m above a marker horizon dated at 71.0 ± 0.2 Ma by Crame et al. (1999) based on strontium isotope stratigraphy, indicating its close temporal proximity to this datum (Cordes-Person et al., 2020).

**Additional outgroup 5**

**Fossil taxon.** *Apsaravis ukhaana* Norell and Clarke 2001.

**Minimum age.** 71.9 Ma.

**Age justification.** *Apsaravis* is known from the Ukhaa Tolgod locality of the Bayn Dzak Member of the Djadokhta Formation, Ömnögovi Province, Mongolia (Clarke and Norell, 2002). This unit is considered to be late Campanian in age (Dashzėvėg et al., 2005; Dingus et al., 2008; Hasegawa et al., 2009), with magnetostratigraphic evidence suggesting its deposition during the end of chron C33 and chron C32 (Dashzėvėg et al., 2005), potentially corresponding to a minimum age of 71.449 Ma (Ogg, 2012). However, Dashzėvėg et al. (2005) cautioned that this correlation to the geomagnetic timescale was only tentative. The minimum age used here is therefore based on the end of the Campanian according to ICS v2020/03 (inclusive of error).

**Additional outgroup 6**

**Fossil taxon.** *Archaeopteryx* sp. Rauhut et al. 2018.

**Specimen.** DNWK 02924 (referred specimen; Rauhut et al., 2018), Dinosaurier Freiluftmuseum Altmühltal, Denkendorf, Bayern, Germany.

**Minimum age.** 151.2 Ma.

**Age justification.** The oldest known remains of *Archaeopteryx* consist of the recently described 12th skeletal specimen from the Öchselberg Member of the Painten Formation near Schamhaupten, Bavaria, Germany (Rauhut et al., 2018). This unit predates the Altmühltal and Mörnsheim Formations that yielded all the previously described fossils of the genus (variously attributed to one or several species; Kundrát et al., 2019), and can be correlated to the *eigeltingense* ammonite biohorizon, whose base coincides with the Kimmeridgian–Tithonian boundary (Schweigert, 2007). Given the proximity of the Painten specimen to this boundary, its date (according to ICS v2020/03 and inclusive of error) is used here as the minimum age constraint on *Archaeopteryx*.

**Additional outgroup 7**

**Fossil taxon.** *Archaeorhynchus spathula* Zhou and Zhang 2006.

**Minimum age.** 121.8 Ma.

**Age justification.** Material of *Archaeorhynchus* has been described from both the Yixian Formation and the overlying Jiufotang Formation in Liaoning Province, China (Zhou et al., 2013). The stratigraphic subdivision and absolute ages of these two units have been the subject of a number of recent studies (He et al., 2004; Yang et al., 2007; Zhu et al., 2007; Chang et al., 2009; Wang et al., 2016b; Chang et al., 2017; Zhong et al., 2021). Given the lack of more precise information about the stratigraphic provenance of the relevant fossils, the minimum age used here for the Yixian remains of *Archaeorhynchus* is based on the date of 122.1 ± 0.3 Ma estimated for a tuff from the lowermost part of the Jiufotang Formation using high-precision $^{40}Ar/^{39}Ar$ dating (Chang et al., 2009). After the completion of this study, Zhong et al. (2021) suggested that the duration of the Yixian Formation was considerably shorter than hypothesized by Chang et al. (2009), with a minimum age of

124.122 ± 0.048 Ma based on the high-precision U–Pb dating of zircons collected from tuff layers in the Jin-Yang basin.

##### Additional outgroup 8

**Fossil taxon.** *Archaeornithura meemannae* Wang et al. 2015a.

**Minimum age.** 130.7 Ma.

**Age justification.** *Archaeornithura* is known from the *Protopteryx* horizon in the Sichakou basin, Fengning County, Hebei Province, China, which can be referred to the lower part of the Huajiying Formation (Wang et al., 2015a). The widely reported date of 130.7 Ma for this horizon is based on the weighted mean of $^{40}Ar/^{39}Ar$ ages for an interbedded tuff layer located 6 m below the bird fossil-bearing shales (He et al., 2006).

##### Additional outgroup 9

**Fossil taxon.** *Bellulornis rectusunguis* (Wang et al. 2016a).

**Minimum age.** 119.6 Ma.

**Age justification.** The known material of *Bellulornis* derives from the Jiufotang Formation of Jianchang County, Liaoning Province, China (Wang et al., 2016a). Given the lack of more precise information about its stratigraphic provenance, the age of *Bellulornis* cannot be narrowed down beyond the range spanned by the Jiufotang Formation as a whole. However, this age range is itself uncertain, as Chang et al. (2009) questioned the previously proposed $^{40}Ar/^{39}Ar$ date of 110.59 ± 0.52 Ma, based on a basalt collected from Tebch, Inner Mongolia (Eberth et al., 1993), on the grounds that the correlation between the Jiufotang Formation strata in Inner Mongolia and in Liaoning remains uncertain. Here, we use a minimum age of 119.6 Ma, derived from a combined U–Pb and $^{40}Ar/^{39}Ar$ estimate for an interbedded tuff layer located approximately 1.5 m above the main fossil-bearing shale at the Shangheshou locality in Liaoning (He et al., 2004). This estimate represents the original source for the 120 Ma date widely attributed to the Jiufotang bird fossils (Wang et al., 2014a; Hu et al., 2015; Wang et al., 2017b), but it fails to account for the known ∼1% discrepancy between the U–Pb and $^{40}Ar/^{39}Ar$ methods (Chang et al., 2009).

##### Additional outgroup 10

**Fossil taxon.** *Ceramornis major* Brodkorb 1963.

**Minimum age.** 66.2 Ma.

**Age justification.** The fossils of *Ceramornis* are documented from UCMP locality V5620 of the Lance Formation, Niobrara County, Wyoming (Hope, 2002). Based on the position of the site within the formation and assumed sedimentation rates, Longrich et al. (2011) estimated the corresponding bird fossils to predate the end-Cretaceous extinction event by 200,000 years. Combined with the recent estimate of 66.043 ± 0.011/0.043 Ma for the age of the K–Pg boundary (Renne et al., 2013), this results in the value employed here.

##### Additional outgroup 11

**Fossil taxon.** *Changzuiornis ahgmi* Huang et al. 2016.

**Minimum age.** 119.6 Ma.

**Age justification.** *Changzuiornis* is known from the Sihedang locality near Lingyuan, Liaoning Province, China (Huang et al., 2016). There is some uncertainty as to whether this locality belongs to the Yixian or Jiufotang Formation (Yao et al., 2019): the original description of the taxon reported the latter (Huang et al., 2016), while numerous other studies on Mesozoic birds assigned the site to the older Yixian Formation (Zhou et al., 2014a; O'Connor et al., 2016; Hu and O'Connor, 2017; Wang and Zhou, 2018). Shao et al. (2018) cite a recent (2016), untranslated Chinese reference placing the Sihedang beds within the "third member" of the Jiufotang Formation. Since the minimum age should be based on the youngest plausible estimate, we assume for calibration purposes that the assignment of the Sihedang locality to the younger unit is correct, and given the paucity of radiometric dates available for the Jiufotang Formation, we use the same numeric age estimate for Sihedang as for the entire formation. For the source of this numeric age, see Additional outgroup 9.

##### Additional outgroup 12

**Fossil taxon.** *Chaoyangia beishanensis* Hou and Zhang 1993.

**Minimum age.** 119.6 Ma.

**Age justification.** *Chaoyangia* is known from the Xidagou locality of the Jiufotang Formation, near the town of Boluochi, Chaoyang County, Liaoning Province, China (O'Connor and Zhou, 2013). Zhang et al. (2007) assigned the Xidagou Bed to the top of the middle member of the Jiufotang Formation, whose exact stratigraphy and subdivision nevertheless remain uncertain. The minimum age applied to the fossils from this locality is thus the same as that used for other Jiufotang taxa, and follows the reasoning given for Additional outgroup 9.

**Additional outgroup 13**

**Fossil taxon.** "*Cimolopteryx*" *maxima* Brodkorb 1963.

**Minimum age.** 66.0 Ma.

**Age justification.** The holotype of "*C.*" *maxima*, which represents the only known specimen of the taxon according to Longrich et al. (2011, *contra* Hope, 2002), is known from UCMP locality V5711 of the Lance Formation, Niobrara County, Wyoming (Hope, 2002). The position of this site within the Lance Formation is not precisely documented, but Longrich et al. (2011) consider its age to be within 650,000 years of the K–Pg boundary based on the assumption that the Lance Formation was deposited over a period of 1.3 Myr, similar to the Hell Creek Formation. Combined with the estimate of Renne et al. (2013) for the K–Pg boundary, this yields an age range of 66.65–66.0 Ma, whose lower bound is used here as the minimum age for the taxon.

**Additional outgroup 14**

**Fossil taxon.** "*Cimolopteryx*" *minima* Brodkorb 1963.

**Minimum age.** 66.0 Ma.

**Age justification.** The holotype and only known specimen of "*C.*" *minima* derives from UCMP locality V5003 of the Lance Formation, Niobrara County, Wyoming (Hope, 2002). Longrich et al. (2011) offer the same reasoning for the age of this site as for UCMP V5711 (see Additional outgroup 13), resulting in the same minimum age being used here.

**Additional outgroup 15**

**Fossil taxon.** "*Cimolopteryx*" *petra* Hope 2002.

**Minimum age.** 66.0 Ma.

**Age justification.** The holotype and only known specimen of "*C.*" *petra* is known from UCMP locality V5711 of the Lance Formation, Niobrara County, Wyoming (Hope, 2002). For the reasoning behind the numeric age assigned to this site, see Additional outgroup 13.

**Additional outgroup 16**

**Fossil taxon.** *Cimolopteryx rara* Marsh 1892.

**Minimum age.** 66.0 Ma.

**Age justification.** The holotype of *C. rara*, which represents the only known specimen of the taxon according to Longrich et al. (2011, *contra* Hope, 2002), derives from an unknown locality within the type area of the Lance Formation, Niobrara County, Wyoming (Hope, 2002). Given the lack of more precise information about the stratigraphic provenance of the fossil, the date used here is based on the age range estimated by Longrich et al. (2011) for the Lance Formation as a whole (67.3–66.0 Ma).

##### Additional outgroup 17

**Fossil taxon.** *Dingavis longimaxilla* O'Connor et al. 2016.

**Minimum age.** 119.6 Ma.

**Age justification.** The holotype and only known specimen of *Dingavis* was described from the Sihedang locality (Jehol Group), Liaoning Province, China (O'Connor et al., 2016). For the reasoning behind the numeric age assigned to this locality, see Additional outgroup 11.

##### Additional outgroup 18

**Fossil taxa.** *Enaliornis barretti* Seeley 1876, *Enaliornis sedgwicki* Seeley 1876, *Enaliornis seeleyi* Galton and Martin 2002.

**Minimum age.** 100.5 Ma.

**Age justification.** The three species of *Enaliornis* jointly represent the oldest known representatives of Hesperornithiformes (Bell and Chiappe, 2016). All three are known from multiple specimens found in the Cambridge Greensand, Cambridgeshire, UK (Galton and Martin, 2002), now considered to represent the basalmost member of the West Melbury Marly Chalk Formation (Machalski, 2018; Barrett and Bonsor, 2021). Galton and Martin (2002) reported that the strata yielding *Enaliornis* fossils span either the entire *Stoliczkaia dispar* ammonite biozone or just the top *Mortoniceras perinflatum* subzone, suggesting a latest Albian age for the taxon. Recently, the subzones of the *Stoliczkaia dispar* zone were elevated to zone status and reported to span an age range of 101.72–100.91 Ma, somewhat predating the Albian–Cenomanian boundary (Ogg et al., 2012). Machalski (2018) suggested that the Cambridge Greensand fauna may extend into the earliest Cenomanian based on a reexamination of the phosphatized specimens of the ammonite *Schloenbachia varians*, but this was regarded as unlikely by Barrett and Bonsor (2021), who considered most of the vertebrate material found in the Cambridge Greensand Member to have been reworked from the underlying upper Albian Gault Formation. Here, the minimum age of *Enaliornis* is conservatively based on the Albian–Cenomanian boundary as dated by ICS v2020/03.

**Additional outgroup 19**

**Fossil taxon.** *Eoconfuciusornis zhengi* Zhang et al. 2008.

**Minimum age.** 130.7 Ma.

**Age justification.** *Eoconfuciusornis* represents the oldest known member of Confuciusornithiformes (Navalón et al., 2018). Its fossils derive from the *Protopteryx horizon* in the Sichakou basin (Fengning County, Hebei Province, China), which was assigned to the Dabeigou Formation in the original description of the taxon (Zhang et al., 2008) but subsequently reassigned to the overlying Huajiying Formation (Jin et al., 2008). For the reasoning behind the numeric age assigned to this horizon, see Additional outgroup 8.

**Additional outgroup 20**

**Fossil taxon.** *Eogranivora edentulata* Zheng et al. 2018.

**Minimum age.** 122.6 Ma.

**Age justification.** The known material of *Eogranivora* derives from the Dawangzhangzi locality of the Yixian Formation near Lingyuan, Liaoning Province, China (Zheng et al., 2018). The Dawangzhangzi Beds were considered to represent one of the four major divisions of the Yixian Formation (Zhou et al., 2003) and correspond to the "Daxinfangzi Bed" of other authors (Jin et al., 2008; Sun et al., 2011) or to the lower part of the undivided Upper Yixian Formation, located below the Huanghuashan Unit (Zhong et al., 2021). In their overview, Sun et al. (2011) report three isotopic dates obtained from the relevant strata: an andesite $^{40}$Ar/$^{39}$Ar age of 122.9 $\pm$ 0.3 Ma (Smith et al., 1995), a zircon U–Pb age of 124.4 $\pm$ 1.1 Ma (Zhang et al., 2006), and another zircon U–Pb age of 124.4 $\pm$ 1.4 Ma (Meng et al., 2008). Based on this range of values, Sun et al. (2011) conservatively dated the Dawangzhangzi Beds at 125.8–122.6 Ma, an estimate that is followed here. After the completion of this study, Zhong et al. (2021) reported an U–Pb age of 124.122 $\pm$ 0.048 Ma for a sample from the overlying Huanghuashan Unit, suggesting that our minimum age may be too conservative.

**Additional outgroup 21**

**Fossil taxon.** *Eopengornis martini* Wang et al. 2014b.

**Minimum age.** 130.7 Ma.

**Age justification.** *Eopengornis* is known from the *Protopteryx* horizon of the Huajiying Formation in the Sichakou basin, Fengning County, Hebei Province, China (Wang et al., 2014b). The stratigraphic position of this unit within the Jehol Group renders *Eopengornis* the oldest taxon with well-established enantiornithine affinities. Two other species from the same horizon, *Protopteryx fengningensis* and *Cruralispennia multidonta*, were also described as enantiornithines (Wang et al., 2017a), but their inclusion within the clade is less certain. *Protopteryx* may fall outside of the clade uniting enantiornithines and ornithuromorphs (Cau et al., 2017) or within Ornithuromorpha (Hartman et al., 2019), whereas *Cruralispennia* was recently inferred to be the earliest-diverging ornithuromorph (Wang et al., 2020c). Therefore, we choose *Eopengornis* alone to stand in for Enantiornithes in our outgroup sequences as the oldest known member of the group. For the reasoning behind the numeric age assigned to the *Protopteryx* horizon, see Additional outgroup 8.

**Additional outgroup 22**

**Fossil taxon.** *Gansus yumenensis* Hou and Liu 1984.

**Minimum age.** 114.1 Ma.

**Age justification.** *Gansus* is known from the Xiagou Formation of the Changma Basin, Gansu Province, China (You et al., 2006). Based on the cyclostratigraphic analysis of samples collected from the Zhangye Basin, Liu et al. (2017) estimated the age of the Xiagou Formation at 120.2–114.1 Ma. This range is compatible with the less precise $^{40}Ar/^{39}Ar$ estimates of 112.8 ± 3.4 Ma and 118.8 ± 3.6 Ma obtained by Li and Yang (2004) for samples collected directly from the Changma Basin.

**Additional outgroup 23**

**Fossil taxon.** *Hollanda luceria* Bell et al. 2010.

**Minimum age.** 71.449 Ma.

**Age justification.** *Hollanda* is known from the Hermiin (= Khermeen) Tsav locality in the western Nemegt Basin, Ömnögovi Province, Mongolia, which belongs to the "Middle Red Bed" of the Baruungoyot (= Barun Goyot) Formation (Bell et al., 2010). The relationship of the Baruungoyot Formation to other units outcropping in the southern Gobi Desert remains uncertain. It conformably underlies the Nemegt Formation, but the existence of an approximately 25 m thick interfingering interval in the central Nemegt and the presence of multiple taxa within both formations suggests that their faunas and ecosystems were partly coeval (Eberth et al., 2009; Fanti et al., 2018; Funston et al., 2018; Nakajima et al., 2018). The physical contact between the Baruungoyot and Djadokhta Formations has not yet been discovered (Khand et al., 2000; Dingus et al., 2008), and both units are either considered to be approximately coeval (Dingus et al., 2008; Bell et al., 2010), or the stratigraphically lower Djadokhta Formation is regarded as somewhat older (Khand et al., 2000; Dashzėvėg et al., 2005). Taken together, these observations suggest that *Hollanda* is likely somewhat older than *Teviornis* (Additional outgroup 2) but younger than or contemporary to *Apsaravis* (Additional outgroup 5). However, note that after the completion of this study, Jerzykiewicz et al. (2021) proposed that all three formations – Djadokhta, Baruungoyot, and Nemegt – were largely coeval, with the differences among their faunal assemblages reflecting different environments rather than temporal succession.

Bell et al. (2010) based their estimate of 75–71 Ma for the age of *Hollanda* on the magnetostratigraphic data reported by Dashzėvėg et al. (2005), which, however, pertained to the Bayn Dzak Member of the Djadokhta Formation, outcropping at the Flaming Cliffs and Khren Tsav localities. The tentative referral of Hermiin Tsav to chron C32n cited by Weishampel et al. (2008) is broadly compatible with the interval suggested by Dashzėvėg et al. (2005) for Bayn Dzak (comprising the end of C33 in addition to C32), but has the added advantage of being more precise and pertaining directly to the locality under consideration. We therefore use the date given for the top of C32n by Ogg (2012) as the minimum age for *Hollanda*.

##### Additional outgroup 24

**Fossil taxon.** *Hongshanornis longicresta* Zhou and Zhang 2005.

**Minimum age.** 125.0 Ma.

**Age justification.** The holotype of *Hongshanornis* (IVPP V14533) was described from the Shifo locality of the Yixian Formation, Ningcheng County, Inner Mongolia Autonomous Region, China (Zhou and Zhang, 2005). An additional specimen (DNHM D2945/6) was collected from the exposures of the same formation at Dawangzhangzi near Lingyuan, Liaoning Province (Chiappe et al., 2014). As confirmed by Zheng et al. (2014), the former locality in fact refers to the same site as the "Xisanjia locality" that also yielded the remains of *Tianyuornis* (see Additional outgroup 54), Xisanjia being the name of a village and Shifo referring to the corresponding township (Jarzembowski et al., 2015). This horizon can be correlated to the Jianshangou Unit of western Liaoning (Jarzembowski et al., 2015; Bi et al., 2018), the validity of which is recognized in most of the subdivisions proposed for the formation (Zhou et al., 2003; Jin et al., 2008; Chang et al., 2017; Zhong et al., 2021). The Jianshangou Unit is stratigraphically lower and older than the Dawangzhangzi Beds (Zhou et al., 2003; Sun et al., 2011), suggesting that the holotype predates the referred specimen. The radioisotopic ages obtained for tuffs immediately above the fossil-bearing bed of the Jianshangou Unit were recently recalculated by Chang et al. (2017), resulting in a weighted mean of 125.22 $\pm$ 0.22 Ma. This range is also consistent with two additional dates reported by Yang et al. (2007) and Meng et al. (2008) (as cited in Chang et al., 2017 and Sun et al., 2011, respectively), and its lower bound is therefore used here as the minimum age for *Hongshanornis*. After the completion of this study, Zhong et al. (2021) obtained a U–Pb age of 125.457 $\pm$ 0.27 Ma (combined analytical, tracer, and decay constant uncertainties) for another tuff sample above the main fossil-bearing layer of the Jianshangou Unit, which is also compatible with the minimum employed here.

**Additional outgroup 25**

**Fossil taxon.** *Iaceornis marshi* Clarke 2004.

**Minimum age.** 80.5 Ma.

**Age justification.** The holotype and only known specimen of *Iaceornis* is known from the Smoky Hill Chalk Member of the Niobrara Formation in Gove County, Kansas (Clarke, 2004). The minimum age is based on the assumption that the material derives from the "*Hesperornis* zone", which represents the youngest subunit of the Smoky Hill Chalk Member that is known to yield fossils of *Ichthyornis*-like birds (Carpenter, 2008). The numeric value is based on Carpenter (2008, Figure 1).

**Additional outgroup 26**

**Fossil taxon.** *Ichthyornis dispar* Marsh 1872.

**Minimum age.** 80.5 Ma.

**Age justification.** As for *Iaceornis* (Additional outgroup 25). Note that this is likely a severe underestimate, since material as old as Cenomanian has been referred to *I. dispar* (specimens SMNH P2077.67, SMNH P2077.111, SMNH P2077.112, SMNH P2487.5 from the Belle Fourche Formation, Saskatchewan; Clarke, 2004). Moreover, the holotype of the taxon (YPM 1450) itself likely comes from the *Cladoceramus undulatoplicatus* inoceramid biozone, whose top yielded a $^{40}$Ar/$^{39}$Ar age of 84.88 $\pm$ 0.28 Ma (Obradovich, 1993), recently recalibrated to 85.84 $\pm$ 0.37 Ma by Sageman et al. (2014) and to 85.409 $\pm$ 0.282 Ma by Fowler (2017). While a specimen-level phylogenetic analysis of *Ichthyornis*-like birds has not been conducted and the conspecificity of the Cenomanian material with *I. dispar* is uncertain (Tokaryk et al., 1997; Clarke, 2004), the apomorphies shared by the specimens from various levels of the Smoky Hill Chalk suggest that the divergence of the *Ichthyornis* lineage from more crownward birds substantially predates the minimum plausible age derived here. Nevertheless, using the age of the Cenomanian–Turonian boundary (93.9 Ma) instead of 80.5 Ma for *Ichthyornis* has no appreciable effect on the resulting calibration upper bounds (range of differences: 0.01–0.52 Myr), and for the sake of consistency with the treatment of other outgroups, we therefore employ the latter minimum here.

**Additional outgroup 27**

**Fossil taxon.** Ichthyornithes *incertae sedis* ("Ornithurine D" of Longrich et al., 2011).

**Specimens.** AMNH 22002, American Museum of Natural History, New York City, NY, USA; RSM P2992.11, Royal Saskatchewan Museum, Regina, SK, Canada; and UCMP 187207, University of California Museum of Paleontology, Berkeley, CA, USA.

**Minimum age.** 66.0 Ma.

**Age justification.** Longrich et al. (2011) referred three specimens to their "ornithurine D": RSM P2992.11 from the Frenchman Formation of Saskatchewan; UCMP 187207 from UCMP locality V84145 of the Hell Creek Formation in Montana; and AMNH 22002 (only mentioned in passing) from the Lance Formation of Wyoming. The minimum ages of all three specimens coincide with one another and with the K–Pg boundary. The entire Frenchman Formation lies within the Cretaceous portion of chron C29r (McIver, 2002), and can thus be constrained to the last 259 ± 52 kyr of the Cretaceous (Sprain et al., 2018). The duration of the Hell Creek Formation was substantially longer (Longrich et al., 2011: 1.3 Myr; Sprain et al., 2018: 1.8 Myr), but the site that yielded specimen UCMP 187207 can also be assigned to chron C29r, resulting in the same narrow range of ages. Finally, the Lance Formation was assumed by Longrich et al. (2011) to have been deposited over the same period as the Hell Creek Formation, and also extends to the K–Pg boundary. Therefore, the end of the Cretaceous provides the minimum age for all the "ornithurine D" fossils.

##### Additional outgroup 28

**Fossil taxon.** *Iteravis huchzermeyeri* Zhou et al. 2014a.

**Minimum age.** 119.6 Ma.

**Age justification.** *Iteravis* is known from the Sihedang locality (Jehol Group) near Lingyuan, Liaoning Province, China (Zhou et al., 2014a). For the reasoning behind the numeric age assigned to this locality, see Additional outgroup 11. Note that following Wang et al. (2018), we treat *Iteravis huchzermeyeri* as the senior synonym of "*Gansus zheni*" Liu et al. 2014, whose holotype was described from the same locality and formation.

##### Additional outgroup 29

**Fossil taxon.** *Jeholornis curvipes* Lefèvre et al. 2014.

**Minimum age.** 122.6 Ma.

**Age justification.** In their recent revision of the Jeholornithiformes, Wang et al. (2020d) recognized three valid species of *Jeholornis*: *J. prima*, *J. curvipes*, and *J. palmapenis*. Of these, *J. palmapenis* is restricted to the younger Jiufotang Formation (O'Connor et al., 2012) and *J. curvipes* is restricted to the older Yixian Formation (Lefèvre et al., 2014). The type species *J. prima* has been repeatedly reported to occur in both formations (Zhou and Zhang, 2007; Li et al., 2010), but no catalogue numbers were provided for the alleged Yixian specimens, and the information was likely based on the assumption that the Yixian taxon *Jixiangornis orientalis* represented a junior synonym of *J. prima*, which was rejected by Wang et al. (2020d). However, a Yixian specimen of *J. prima* (BMNHC PH780) from

Xiaoyugou, Chaoyang County, Liaoning Province, China, was illustrated by Chiappe and Meng (2016). In the absence of more precise information about the stratigraphic provenance of this specimen, *J. prima* is still likely superseded as the oldest known representative of Jeholornithiformes by *J. curvipes*, whose fossils derive from the "Dakangpu Member" of the Yixian Formation, considered by Lefèvre et al. (2014) to be equivalent to the Dawangzhangzi Beds. Therefore, for the purposes of outgroup sequence construction, *Jeholornis* is here taken to refer to *J. curvipes* where no specific epithet is given (Zheng et al., 2018), and *J. curvipes* replaces *J. prima* in phylogenies that explicitly refer to the latter species (Field et al., 2018). This assumes the monophyly of a clade uniting *J. curvipes* and *J. prima* to the exclusion of other birds, a result repeatedly supported by formal phylogenetic analyses (Lefèvre et al., 2014; Wang et al., 2020d). For the reasoning behind the numeric age assigned to the Dawangzhangzi Beds, see Additional outgroup 20.

##### Additional outgroup 30

**Fossil taxon.** *Jianchangornis microdonta* Zhou et al. 2009.

**Minimum age.** 119.6 Ma.

**Age justification.** *Jianchangornis* is known from the Jiufotang Formation of Jianchang County, Liaoning Province, China (Zhou et al., 2009). For the reasoning behind the numeric age assigned to the Jiufotang Formation, see Additional outgroup 9.

##### Additional outgroup 31

**Fossil taxon.** *Juehuaornis zhangi* Wang et al. 2015b.

**Minimum age.** 119.6 Ma.

**Age justification.** In the original description of the taxon (Wang et al., 2015b), the fossil material of *Juehuaornis* was reported to derive from the exposures of the Jiufotang Formation in the township of Sanjiazi, Lingyuan, Liaoning Province, China. Hu and O'Connor (2017) instead suggested the nearby Sihedang locality as its provenance, based on the color and lithology of the slab (Hu Han, pers. comm.). For the reasoning behind the numeric age assigned to the Sihedang locality, see Additional outgroup 11.

##### Additional outgroup 32

**Fossil taxon.** *Khinganornis hulunbuirensis* Wang et al. 2020c.

**Minimum age.** 120.49 Ma.

**Age justification.** *Khinganornis* is known from the Pigeon Hill locality of the upper Longjiang Formation near the town of Baoshan, Hulunbuir City, Inner Mongolia Autonomous Region, China (Wang et al., 2020c). U–Pb dating of zircon grains from volcanic tuff samples found in the *Khinganornis* fossil-bearing bed constrains the age of the taxon with high precision to 121.23 ± 0.74 Ma (Wang et al., 2019a). The value employed here corresponds to the lower bound of this range.

##### Additional outgroup 33

**Fossil taxon.** *Limenavis patagonica* Clarke and Chiappe 2001.

**Minimum age.** 72.1 Ma.

**Age justification.** *Limenavis* is known from the Salitral Moreno locality (Río Negro Province, Argentina), belonging to the "Lower Member" of the Allen Formation (Clarke and Chiappe, 2001). Different lines of biostratigraphic evidence suggest either a middle Campanian or an early Maastrichtian age for the lower Allen Formation (Clarke and Chiappe, 2001), a range that remains widely cited in the literature without further comment (e.g., González et al., 2020). Here, we follow Head (2015) in using the Campanian–Maastrichtian boundary boundary to derive a minimum age for Allen Formation taxa, even though this value overestimates the true minimum somewhat.

##### Additional outgroup 34

**Fossil taxon.** *Lithornis celetius* Houde 1988.

**Minimum age.** 60.21 Ma.

**Age justification.** The genus *Lithornis* includes six valid, named species, of which the oldest is the middle Paleocene *L. celetius* (Stidham et al., 2014; Nesbitt and Clarke, 2016). The known material of *L. celetius* derives from the Bangtail Quarry of the Fort Union Formation in the western Crazy Mountain Basin, Park County, Montana (Nesbitt and Clarke, 2016). The Bangtail Quarry can be assigned to the earliest substage (Ti1) of the Tiffanian North American Land Mammal Age (NALMA) according to Lofgren et al. (2004). Jarvis et al. (2014, Supplementary Material: 37) used this assignment to derive a minimum age of 60.6 Ma for *Lithornis*, citing Secord (2008) in support of this value. However, this date was in fact intended to correspond to the base rather than the top of Ti1; accordingly, the minimum age should be 60.21 Ma instead (the latest possible date assuming a 1 Myr hiatus near the base of chron C26r). Secord et al. (2006) also offered another potential constraint on the end of Ti1 based on a volcanic ash bed positioned 6–10 m below a site correlated to Ti2, which was dated to 61.06 ± 0.33 Ma by Belt et al. (2004). Here, we use the younger of the two dates as the latest possible age for *L. celetius*.

An even older fossil possibly referable to *Lithornis* has been reported from the Crosswicks Creek locality of the Hornerstown Formation (Main Fossiliferous Layer, 66.0 ± 0.1 Ma; see the discussion under Additional outgroup 1). The material consists of an incomplete right scapula (specimen NJSM 15065) described as being *Lithornis*-like (Parris and Hope, 2002). This was partially corroborated by Nesbitt and Clarke (2016), who found the specimen to possess a laterally hooked acromion, optimized in their phylogenetic analyses as a synapomorphy of Lithornithidae (a putative clade including *Lithornis*, *Calciavis*, *Paracathartes*, and *Pseudocrypturus*). However, another recent analysis found "lithornithids" to be paraphyletic with respect to crown-group paleognaths (Yonezawa et al., 2017), in which case the character would be most parsimoniously interpreted as a pan-paleognath symplesiomorphy. Given the fragmentary nature of the material, its affinities are here considered to be too uncertain to allow extending the stratigraphic range of *Lithornis* to the K–Pg boundary.

##### Additional outgroup 35

**Fossil taxon.** *Longicrusavis houi* O'Connor et al. 2010.

**Minimum age.** 122.6 Ma.

**Age justification.** *Longicrusavis* is known from the Dawangzhangzi locality of the Yixian Formation near Lingyuan, Liaoning Province, China (O'Connor et al., 2010). For the reasoning behind the numeric age assigned to the Dawangzhangzi Beds, see Additional outgroup 20.

##### Additional outgroup 36

**Fossil taxon.** *Maaqwi cascadensis* McLachlan et al. 2017.

**Minimum age.** 71.939 Ma.

**Age justification.** The known material of *Maaqwi* was collected from the intertidal exposures of the Northumberland Formation along the northwestern shore of Hornby Island, British Columbia, Canada (McLachlan et al., 2017). According to McLachlan and Haggart (2018, Figure 2), the boundary between the Northumberland Formation and the conformably overlying Geoffrey Formation at Hornby Island lies within magnetochron C32n.2n, whose top (as dated by Ogg, 2012) is used here to constrain the minimum age of *Maaqwi*.

##### Additional outgroup 37

**Fossil taxon.** "*Martinavis*" *saltariensis* Walker and Dyke 2009.

**Minimum age.** 71.5 Ma.

**Age justification.** "*M.*" *saltariensis* is known from the El Brete locality of the Lecho Formation, La Candelaria Department, Salta Province, Argentina (Walker and Dyke, 2009). A volcanic tuff sample from the overlying Yacoraite Formation, in a layer positioned 27.6 m above its contact with the Lecho Formation, yielded a U–Pb age of 71.9 ± 0.4 Ma (Marquillas et al., 2011). The lower end of this range constrains the minimum age of "*M.*" *saltariensis*.

##### Additional outgroup 38

**Fossil taxon.** "*Martinavis*" *whetstonei* Walker and Dyke 2009.

**Minimum age.** 71.5 Ma.

**Age justification.** "*M.*" *whetstonei* was described in the same publication and from the same formation and locality as "*M.*" *saltariensis*: El Brete, Lecho Formation, Argentina (Walker and Dyke, 2009). For the reasoning behind the numeric age assigned to the El Brete avifauna, see Additional outgroup 37.

##### Additional outgroup 39

**Fossil taxon.** *Mystiornis cyrili* Kurochkin et al. 2011.

**Minimum age.** 113.0 Ma.

**Age justification.** The known material of *Mystiornis* derives from the Shestakovo-1 locality of the Ilek Formation (= Ilekskaya Svita) in Tchebulinski District, Kemerovskaya Oblast, Russia (Kurochkin et al., 2011). The age of the Ilek Formation is poorly constrained (Averianov et al., 2018), with estimates ranging from Valangian based on mollusk and ostracod biostratigraphy to Albian based on palynology (Golovneva and Shchepetov, 2010). However, the overlying Kiya Formation can be dated to the late Albian based on phytostratigraphy (Golovneva and Shchepetov, 2010), suggesting a somewhat older age for the Ilek Formation. Averianov et al. (2018) considered a Barremian age to be most likely but not definitive. Based on this information, we use the Aptian–Albian boundary as dated by ICS v2020/03 to derive a minimum age for *Mystiornis*.

##### Additional outgroup 40

**Fossil taxon.** Ornithurae *incertae sedis* ("Ornithurine A" of Longrich et al., 2011).

**Specimens.** RSM P1927.936, Royal Saskatchewan Museum, Regina, SK, Canada; UCMP 53962 and UCMP 53963, University of California Museum of Paleontology, Berkeley, CA, USA; uncatalogued specimen, American Museum of Natural History, New York City, NY, USA.

**Minimum age.** 66.2 Ma.

**Age justification.** The four known specimens of "Lancian ornithurine A" all derive from the Lance Formation of Wyoming (UCMP localities V5620 and V5711) or from the Frenchman Formation of Saskatchewan (Longrich et al., 2011). While the exact stratigraphic position of the other localities is unclear, and the minimum age of the corresponding fossils therefore cannot be constrained beyond the K–Pg boundary, UCMP V5620 predates the boundary by ∼200,000 years (see Additional outgroup 10), yielding the minimum age used here.

##### Additional outgroup 41

**Fossil taxon.** Ornithurae *incertae sedis* ("Ornithurine B" of Longrich et al., 2011).

**Specimen.** UCMP 129143, University of California Museum of Paleontology, Berkeley, CA, USA.

**Minimum age.** 66.0 Ma.

**Age justification.** The only specimen referred by Longrich et al. (2011) to "Lancian ornithurine B" is known from UCMP locality V75178 of the Hell Creek Formation, Wild Horse Basin, Garfield County, Montana (Longrich et al., 2011; Arens et al., 2014). The Hell Creek exposures in Garfield County can be assigned to the Cretaceous portion of chron C29r, resulting in an age range of at most 66.31–66.0 Ma (Sprain et al., 2018), whose lower bound is used here as the minimum age for the taxon.

##### Additional outgroup 42

**Fossil taxon.** Ornithurae *incertae sedis* ("Ornithurine C" of Longrich et al., 2011).

**Specimens.** MOR 2918, Museum of the Rockies, Bozeman, MT, USA; SDSM 64281A and SDSM 64281B, South Dakota School of Mines, Rapid City, SD, USA; UCMP 175251 and UCMP 187208, University of California Museum of Paleontology, Berkeley, CA, USA; and YPM PU 17020, Yale Peabody Museum, New Haven, CT, USA.

**Minimum age.** 66.0 Ma.

**Age justification.** The specimens referred by Longrich et al. (2011) to their "Lancian ornithurine C" mostly derive from the Hell Creek Formation of Montana and South Dakota (UCMP locality V93126 and other, unspecified localities) and, in the case of YPM PU 17020, from the Lance Formation of Wyoming (Longrich et al., 2011, Table S2). UCMP 187208, however, is known from the Fort Union Formation, which overlies the Hell Creek Formation in Montana and can be dated to the early Paleocene (Fowler, 2017). As the minimum age of the taxon should be based on the youngest plausible date for its oldest known occurrence, the value used here is based on the K–Pg boundary, following the same reasoning as for Additional outgroup 27.

**Additional outgroup 43**

**Fossil taxon.** Ornithurae *incertae sedis* ("Ornithurine E" of Longrich et al., 2011).

**Specimens.** AMNH 13011, American Museum of Natural History, New York City, NY, USA; and USNM 181923, United States National Museum, Washington, DC, USA.

**Minimum age.** 66.0 Ma.

**Age justification.** Longrich et al. (2011) attributed two specimens to their "Lancian ornithurine E": USNM 181923 from UCMP locality V5622 of the Lance Formation, Wyoming, and AMNH 13011 from a different locality (UCMP V5711) of the same unit. (Note that AMNH 13011 was erroneously listed as "USNM 13011" in their Table S2, but the correct specimen designation is given in their Figure 3 and in Hope, 2002). For the reasoning behind the numeric age assigned to these sites, see Additional outgroups 13 and 27.

**Additional outgroup 44**

**Fossil taxon.** Ornithurae *incertae sedis* ("Ornithurine F" of Longrich et al., 2011).

**Specimens.** ACM 12359, Amherst College Museum, Amherst, MA, USA; and UCMP 53957, University of California Museum of Paleontology, Berkeley, CA, USA.

**Minimum age.** 66.2 Ma.

**Age justification.** Longrich et al. (2011) referred two fossils to their "Lancian ornithurine F", both of which come from the Lance Formation of Wyoming. While the exact stratigraphic provenance of ACM 12359 within this unit remains unknown (Longrich et al., 2011, Table S2), and its minimum age thus cannot be constrained beyond the K–Pg boundary, specimen UCMP 53957 derives from locality UCMP V5620, which predates the boundary by ∼200,000 years (see Additional outgroup 10), yielding the minimum age used here.

**Additional outgroup 45**

**Fossil taxon.** *Palintropus* sp. Hope 2002.

**Specimens.** RTMP 88.116.1 and RTMP 86.36.126 (referred specimens; Hope, 2002), Royal Tyrrell Museum of Palaeontology, Drumheller, AB, Canada.

**Minimum age.** 75.762 Ma.

**Age justification.** Hope (2002, 377) recognized *Palintropus* as a "polymorphic species or group of species", and divided the material assigned to the genus between *Palintropus retusus* from the upper Maastrichtian Lance Formation (with YPM 503 as the holotype and only known specimen) and *Palintropus* sp., represented by two coracoids from the upper Campanian Dinosaur Park Formation (RTMP 88.116.1 and RTMP 86.36.126) and a single scapula from the mid-Campanian Foremost Formation (RTMP 86.146.11). However, the three derived characters Hope (2002) identified as diagnostic of the genus only pertained to the coracoid, and therefore could not be ascertained for the Foremost Formation scapula. Longrich (2009) expanded the array of fossils referred to *Palintropus* and refined Hope's (2002) taxonomy, dividing her undetermined *Palintropus* sp. into "*Palintropus* species A" (comprising RTMP 86.36.126 and two other specimens assigned to the species based on their size) and "*Palintropus* species B" (comprising RTMP 88.116.1, RTMP 86.146.11, and five other specimens). However, like Hope (2002), Longrich (2009) only referred RTMP 86.146.11 to *Palintropus* because of the size of the specimen, and because it articulated well with the known coracoid. His revised diagnosis of the genus was again restricted to coracoid characters, and the assignment of RTMP 86.146.11 to *Palintropus* was described as tentative. Despite potentially extending the stratigraphic range of the genus into the middle Campanian, the referral of RTMP 86.146.11 to *Palintropus* is here considered to be too uncertain to allow the use of the specimen as an age constraint.

Within a single publication, Longrich (2009) variously assigned specimen RTMP 88.116.1 to the Dinosaur Park Formation (DPF; his Figure 13), consistent with Hope (2002), or to the Oldman Formation (p. 172) that underlies the DPF in Dinosaur Provincial Park. The Onefour locality, given as the provenance of the specimen (p. 172), was also inconsistently assigned either to the DPF (Figure 1) or to the Oldman Formation (p. 172). Campbell et al. (2019) support the latter assignment, but clarify that Oldman exposures at Onefour are coeval with the DPF as exposed in the Dinosaur Provincial Park area. Therefore, the uncertainty about the stratigraphic provenance of the fossil has no bearing on its age, which can be based on estimates available for the DPF in either case. In an exhaustive review of the geochronology of the Late Cretaceous Western Interior of North America, Fowler (2017, Table S2) provided a number of recalibrated radiometric dates for the DPF. Aside from two outliers of $< 74$ Ma and $> 77$ Ma, which are incompatible with the overall stratigraphic chart proposed by the author (Fowler, 2017, Table S1), all of these range from 76.5 to 75.5 Ma. This range of dates is also compatible with the two new $^{40}Ar/^{39}Ar$ ages of 76.39 Ma and $76.10 \pm 0.5$ Ma reported by the author, which were obtained from tuffs positioned 36 and 61.5 meters above the Oldman–DPF contact, respectively (Fowler, 2017, Table S1). The lowest non-outlier estimate (inclusive of error) is used here as the minimum age for *Palintropus*.

##### Additional outgroup 46

**Fossil taxon.** *Parahongshanornis chaoyangensis* Li et al. 2011.

**Minimum age.** 119.6 Ma.

**Age justification.** *Parahongshanornis* is known from the Jiufotang Formation exposures in the village of Yuanjiawa, Dapingfang Town, Chaoyang County, Liaoning Province, China (Li et al., 2011). For the reasoning behind the numeric age assigned to the Jiufotang Formation, see Additional outgroup 9.

##### Additional outgroup 47

**Fossil taxon.** *Patagopteryx deferrariisi* Alvarenga and Bonaparte 1992.

**Minimum age.** 83.4 Ma.

**Age justification.** *Patagopteryx* is known from the Boca del Sapo locality of the Bajo de la Carpa Formation (formerly classified as a member of a larger Río Colorado Formation) in Neuquén Province, Argentina (Chiappe, 2002). Following the magnetostratigraphic evidence reported by Dingus et al. (2000), Chiappe (2002) proposed an early to middle Campanian age for *Patagopteryx* (83.5–79.5 Ma); however, these data actually pertained to the conformably overlying Anacleto Formation, and the Bajo de la Carpa Formation itself is likely older (Leanza and Hugo, 2001). In his review of Neuquén Group stratigraphy, Garrido (2010) concurred with this assessment as well as other earlier studies, and suggested a Santonian age for the unit. The minimum age of *Patagopteryx* used here is therefore based on the Santonian–Campanian boundary, as dated by ICS v2020/03 (inclusive of error).

##### Additional outgroup 48

**Fossil taxon.** *Piscivoravis lii* Zhou et al. 2014b.

**Minimum age.** 119.6 Ma.

**Age justification.** *Piscivoravis* is known from the Jiufotang Formation exposures near the village of Xiaotaizi, Liaoning Province, China (Zhou et al., 2014b). For the reasoning behind the numeric age assigned to this formation, see Additional outgroup 9.

##### Additional outgroup 49

**Fossil taxon.** *Qinornis paleocenica* Xue 1995.

**Minimum age.** 62.22 Ma.

**Age justification.** *Qinornis*, possibly representing the only non-neornithine bird surviving into the Cenozoic (Mayr, 2009; Hartman et al., 2019), is known from the Fangou Formation of Shaanxi Province, China (Mayr et al., 2013). This unit can be referred to the Shanghuan

Asian Land Mammal Age (ALMA), whose lower boundary coincides with the end of magnetochron C27n (Vandenberghe et al., 2012; Wang et al., 2019b), yielding the minimum age employed here.

**Additional outgroup 50**

**Fossil taxon.** *Sapeornis chaoyangensis* Zhou and Zhang 2002.

**Specimen.** DNHM-D3078 (referred specimen; Gao et al., 2012), Dalian Natural History Museum, Dalian, Liaoning Province, China.

**Minimum age.** 125.0 Ma.

**Age justification.** Multiple species classified in up to four different genera (*Sapeornis*, "*Omnivoropteryx*", "*Didactylornis*", "*Shenshiornis*") have been included in the family "Sapeornithidae" or "Omnivoropterygidae", but the corresponding fossils likely represent a single species, *Sapeornis chaoyangensis* (Gao et al., 2012; Chiappe and Meng, 2016; Mayr, 2017a). Most of the material referred to *Sapeornis* derives from the Jiufotang Formation, but a single specimen (DNHM-D3078) is known from the older Yixian Formation (Gao et al., 2012; Wang et al., 2017b). Specifically, the fossil was collected from a site near Huludao City, Jianchang County, Liaoning Province, China, which can be attributed to the Jianshangou Unit. For the reasoning behind the numeric age assigned to this unit, see Additional outgroup 24.

**Additional outgroup 51**

**Fossil taxon.** *Schizooura lii* Zhou et al. 2012.

**Minimum age.** 119.6 Ma.

**Age justification.** *Schizooura* is known from the Jiufotang Formation of Jianchang County, Liaoning Province, China (Zhou et al., 2012). For the reasoning behind the numeric age assigned to this formation, see Additional outgroup 9.

**Additional outgroup 52**

**Fossil taxon.** *Songlingornis linghensis* Hou 1997.

**Minimum age.** 119.6 Ma.

**Age justification.** The known material of *Songlingornis* derives from the Xidagou locality of the Jiufotang Formation, near the town of Boluochi, Chaoyang County, Liaoning Province, China (Hou, 1997; O'Connor and Zhou, 2013). For the reasoning behind the numeric age assigned to this locality, see Additional outgroup 12.

**Additional outgroup 53**

**Fossil taxon.** *Tianyuornis cheni* Zheng et al. 2014.

**Minimum age.** 125.0 Ma.

**Age justification.** *Tianyuornis* is known from the Xisanjia locality of the Yixian Formation in the township of Shifo, Chifeng City, Ningcheng County, Inner Mongolia Autonomous Region, China (Zheng et al., 2014). For the reasoning behind the numeric age assigned to this locality, see Additional outgroup 24.

**Additional outgroup 54**

**Fossil taxon.** *Vegavis iaai* Clarke et al. 2005.

**Minimum age.** 66.5 Ma.

**Age justification.** The holotype of *V. iaai* (MLP 93-I-3-1) derives from locality VEG9303 at Cape Lamb, Vega Island, West Antarctica (Clarke et al., 2005), which can be assigned to the lowermost subunit (SBM1) of the Sandwich Bluff Member of the López de Bertodano Formation (Roberts et al., 2014). Referred specimen MACN-PV 19.748 was found in association with the holotype, and can thus be assigned to the same subunit (Clarke et al., 2016). Both fossils are likely slightly older than specimen SDSM 78247 of *Vegavis* sp., described by West et al. (2019) from locality V2005-3, which is positioned approximately 12 m above the horizon that yielded the fossils of *V. iaai* and can be referred to subunit SBM2. Ksepka and Clarke (2015) noted that estimating the age of these older (then only known) specimens is difficult due to the uncertain correlation between the Vega and Seymour Island sections of the López de Bertodano Formation. They provided an older date (67.5 Ma) based on $^{87}Sr/^{86}Sr$ chronology, and a younger date (66.5 Ma) based on the proximity of the *Vegavis* horizon to the base of the "*Manumiella bertodano*" dinoflagellate biozone (now known as the *Alterbidinium longicornutum* biozone; Bowman et al., 2016). Roberts et al. (2014) regarded these estimates as reliable in their revision of Sandwich Bluff stratigraphy. Ksepka and Clarke (2015) argued that the younger of their two dates should be used for calibration purposes, and we follow their choice here.

**Additional outgroup 55**

**Fossil taxon.** *Vorona berivotrensis* Forster et al. 1996.

**Minimum age.** 69.6 Ma.

**Age justification.** The holotype of *Vorona* was found in quarry MAD93-18 of the Anembalemba Member of the Maevarano Formation, near the village of Berivotra in northwestern Madagascar, and several additional specimens tentatively referred to the same species have since been described from the same general area (Forster et al., 1996; O'Connor and Forster, 2010). Abramovich et al. (2003) concluded that the terrestrial vertebrate assemblage of the Anembalemba Member in the Berivotra area is at least 69.6 Ma old and possibly as old as the Campanian–Maastrichtian boundary, since the overlying marine beds of the Berivotra Formation can be dated to planktic foraminiferal zone CF6 (69.6–69.1 Ma) while still being separated from the Maevarano Formation by a 9-meter interval without datable microfossils. This date is therefore used here to constrain the minimum age of *Vorona*.

**Additional outgroup 56**

**Fossil taxon.** *Xinghaiornis lini* Wang et al. 2013a.

**Minimum age.** 119.6 Ma.

**Age justification.** The holotype of *Xinghaiornis* was reported from the Sihetun locality of the Yixian Formation, within the town of Shangyuan, Beipiao City, Liaoning Province, China (Wang et al., 2013a). However, based on lithology of the slab as well as the color and overall preservation of the fossils, O'Connor et al. (2016) suggested that the material came from the Sihedang locality instead, which may belong to the younger Jiufotang Formation. To derive the youngest plausible age for the taxon, we follow this latter possibility here. For the reasoning behind the numeric age assigned to the Sihedang locality, see Additional outgroup 11.

**Additional outgroup 57**

**Fossil taxon.** *Yanornis martini* Zhou and Zhang 2001.

**Minimum age.** 119.6 Ma.

**Age justification.** The type species of *Yanornis* is known from the Jiufotang Formation of Chaoyang City, Liaoning Province, China (Zhou and Zhang, 2001; Wang et al., 2020a). A putative second species of the genus, "*Y.*" *guozhangi*, was described from the exposures of the Yixian Formation in Jinzhou City, Liaoning Province, China (Wang et al., 2013b). A subsequent study failed to corroborate the supposed differences between "*Y.*" *guozhangi* and *Y. martini*, suggesting that the former was a junior synonym of the latter (Wang et al., 2020a), which would extend the stratigraphic range of *Y. martini* into the older Yixian Formation. However, in the only phylogenetic analysis we are aware of where the two species were included as separate terminals, "*Y.*" *guozhangi* was found to be more closely related to *Chaoyangia* and *Schizooura* than to *Y. martini* Hartman et al. (2019). We therefore consider *Y. martini* to be known exclusively from the Jiufotang Formation, and its minimum age is constrained accordingly (see Additional outgroup 9).

**Additional outgroup 58**

**Fossil taxon.** *Yixianornis grabaui* Zhou and Zhang 2001.

**Minimum age.** 119.6 Ma.

**Age justification.** *Yixianornis* is known from the exposures of the Jiufotang Formation near the town of Qianyang, Yi County, Jinzhou City, Liaoning Province, China (Zhou and Zhang, 2001). For the reasoning behind the numeric age assigned to the Jiufotang Formation, see Additional outgroup 9.

**Additional outgroup 59**

**Fossil taxon.** *Yumenornis huangi* Wang et al. 2013c.

**Minimum age.** 114.1 Ma.

**Age justification.** The holotype and only known specimen of *Yumenornis* derives from the Xiagou Formation exposures near the township of Changma, Yumen City, Gansu Province, China (Wang et al., 2013c). For the reasoning behind the numeric age assigned to the Xiagou Formation, see Additional outgroup 22.

**Additional outgroup 60**

**Fossil taxon.** *Zhongjianornis yangi* Zhou et al. 2010.

**Minimum age.** 119.6 Ma.

**Age justification.** The known material of *Zhongjianornis* derives from the Jiufotang Formation of Jianchang County, Liaoning Province, China (Zhou et al., 2010). For the reasoning behind the numeric age assigned to this formation, see Additional outgroup 9.

**Outgroup sequence 1**

**Reference phylogeny.** Field et al., 2018, Supplementary Tree 15.

**Preliminary outgroup sequence.**

| Rank | Original taxon label | Reconciled taxon label | Age (Ma) |
|---|---|---|---|
| 1 | *Iaceornis marshii* [*sic*] | *Iaceornis marshi* | 80.5 |
| 2 | [Multiple taxa of Hesperornithiformes] | *Enaliornis barretti* + *Enaliornis sedgwicki* + *Enaliornis seeleyi* | 100.5 |
| 2 | *Apsaravis ukhaana* | *Apsaravis ukhaana* | 71.9 |
| 2 | *Ichthyornis dispar* | *Ichthyornis dispar* | 80.5 |
| 3 | *Iteravis huchzermeyeri* | *Iteravis huchzermeyeri* | 119.6 |
| 3 | *Gansus zheni* | | |
| 3 | *Gansus yumenenis* | *Gansus yumenensis* | 114.1 |
| 3 | *Changzuiornis ahgmi* | *Changzuiornis ahgmi* | 119.6 |
| 4 | *Yanornis martini* | *Yanornis martini* | 119.6 |
| 5 | *Yixianornis grabaui* | *Yixianornis grabaui* | 119.6 |
| 5 | *Songlingornis linghensis* | *Songlingornis linghensis* | 119.6 |

**Final outgroup sequence.** *Yixianornis grabaui* (Additional outgroup 58), *Yanornis martini* (Additional outgroup 57), *Iteravis huchzermeyeri* (Additional outgroup 28), *Enaliornis* spp. (Additional outgroup 18), *Iaceornis marshi* (Additional outgroup 25), *Teviornis gobiensis* (Additional outgroup 2), *Tsidiiyazhi abini* (Additional outgroup 1).

**Outgroup age sequence.** 119.6, 119.6, 119.6, 100.5, 80.5, 69, 62.221.

##### Outgroup sequence 2

**Reference phylogeny.** Field et al., 2018, Supplementary Tree 17.

**Preliminary outgroup sequence.**

| Rank | Original taxon label | Reconciled taxon label | Age (Ma) |
|---|---|---|---|
| 1 | [Multiple taxa of Hesperornithiformes] | *Enaliornis barretti* + *Enaliornis sedgwicki* + *Enaliornis seeleyi* | 100.5 |
| 2 | *Ichthyornis dispar* | *Ichthyornis dispar* | 80.5 |
| 3 | *Gansus yumenensis* | *Gansus yumenensis* | 114.1 |
| 4 | *Iteravis huchzermeyeri* | *Iteravis huchzermeyeri* | 119.6 |
| 5 | *Tianyuornis cheni* | *Tianyuornis cheni* | 125.0 |
| 5 | *Archaeornithura meemannae* | *Archaeornithura meemannae* | 130.7 |
| 5 | *Parahongshanornis chaoyangornis* [*sic*] | *Parahongshanornis chaoyangensis* | 119.6 |
| 5 | *Hongshanornis longicresta* | *Hongshanornis longicresta* | 125.0 |
| 5 | *Longicrusavis houi* | *Longicrusavis houi* | 122.6 |
| 5 | *Songlingornis linghensis* | *Songlingornis linghensis* | 119.6 |
| 5 | *Piscivoravis lii* | *Piscivoravis lii* | 119.6 |
| 5 | *Yixianornis grabaui* | *Yixianornis grabaui* | 119.6 |
| 5 | *Yanornis martini* | *Yanornis martini* | 119.6 |
| 6 | *Patagopteryx defarrariisi* [*sic*] | *Patagopteryx deferrariisi* | 83.4 |
| 6 | *Vorona berivotrensis* | *Vorona berivotrensis* | 69.6 |
| 6 | *Bellulornis rectusunguis* | *Bellulornis rectusunguis* | 119.6 |
| 6 | *Schizooura lii* | *Schizooura lii* | 119.6 |
| 7 | *Jianchangornis microdonta* | *Jianchangornis microdonta* | 119.6 |
| 8 | *Archaeorhynchus spathula* | *Archaeorhynchus spathula* | 121.8 |
| 9 | [Multiple taxa of Enantiornithes] | *Eopengornis martini* | 130.7 |

**Final outgroup sequence.** *Eopengornis martini* (Additional outgroup 21), *Archaeornithura meemannae* (Additional outgroup 8), *Iteravis huchzermeyeri* (Additional outgroup 28), *Gansus yumenensis* (Additional outgroup 22), *Enaliornis* spp. (Additional outgroup 18), *Teviornis gobiensis* (Additional outgroup 2), *Tsidiiyazhi abini* (Additional outgroup 1).

**Outgroup age sequence.** 130.7, 130.7, 119.6, 114.1, 100.5, 69, 62.221.

**Outgroup sequence 3**

**Reference phylogeny.** Wang et al., 2020c, Figure 10.

**Preliminary outgroup sequence.**

| Rank | Original taxon label | Reconciled taxon label | Age (Ma) |
|---|---|---|---|
| 1 | [Multiple taxa of Hesperornithiformes] | *Enaliornis barretti* + *Enaliornis sedgwicki* + *Enaliornis seeleyi* | 100.5 |
| 2 | *Ichthyornis* | *Ichthyornis dispar* | 80.5 |
| 3 | *Patagopteryx* | *Patagopteryx deferrariisi* | 83.4 |
| 3 | *Vorona* | *Vorona berivotrensis* | 69.6 |
| 3 | *Apsaravis* | *Apsaravis ukhaana* | 71.9 |
| 4 | *Gansus yumenensis* | *Gansus yumenensis* | 114.1 |
| 5 | *Changzuiornis* | *Changzuiornis ahgmi* | 119.6 |
| 5 | *Iteravis* | *Iteravis huchzermeyeri* | 119.6 |
| 5 | *Khinganornis* | *Khinganornis hulunbuirensis* | 120.49 |
| 6 | *Ambiortus* | *Ambiortus dementjevi* | 125.0 |
| 6 | *Piscivoravis* | *Piscivoravis lii* | 119.6 |
| 6 | *Yanornis* | *Yanornis martini* | 119.6 |
| 6 | *Yixianornis* | *Yixianornis grabaui* | 119.6 |
| 7 | *Hongshanornis* | *Hongshanornis longicresta* | 125.0 |
| 7 | *Longicrusavis* | *Longicrusavis houi* | 122.6 |
| 7 | *Parahongshanornis* | *Parahongshanornis chaoyangensis* | 119.6 |
| 7 | *Archaeornithura* | *Archaeornithura meemannae* | 130.7 |
| 7 | *Tianyuornis* | *Tianyuornis cheni* | 125.0 |

**Final outgroup sequence.** *Archaeornithura meemannae* (Additional outgroup 8), *Ambiortus dementjevi* (Additional outgroup 3), *Khinganornis hulunbuirensis* (Additional outgroup 32), *Gansus yumenensis* (Additional outgroup 22), *Enaliornis* spp. (Additional outgroup 18), *Teviornis gobiensis* (Additional outgroup 2), *Tsidiiyazhi abini* (Additional outgroup 1).

**Outgroup age sequence.** 130.7, 125.0, 120.49, 114.1, 100.5, 69, 62.221.

##### Outgroup sequence 4

**Reference phylogeny.** Kundrát et al., 2019, Appendix 14.

**Preliminary outgroup sequence.**

| Rank | Original taxon label | Reconciled taxon label | Age (Ma) |
|---|---|---|---|
| 1 | *Lithornis* | *Lithornis celetius* | 60.21 |
| 2 | *Limenavis patagonica* | *Limenavis patagonica* | 72.1 |
| 3 | *Iaceornis marshii* [*sic*] | *Iaceornis marshi* | 80.5 |
| 4 | *Ichthyornis* | *Ichthyornis dispar* | 80.5 |
| 5 | [Multiple taxa of Hesperornithiformes] | *Enaliornis barretti* + *Enaliornis sedgwicki* + *Enaliornis seeleyi* | 100.5 |
| 6 | *Apsaravis ukhaana* | *Apsaravis ukhaana* | 71.9 |
| 7 | *Yixianornis* | *Yixianornis grabaui* | 119.6 |

**Final outgroup sequence.** *Yixianornis grabaui* (Additional outgroup 58), *Enaliornis* spp. (Additional outgroup 18), *Ichthyornis dispar* (Additional outgroup 26), *Iaceornis marshi* (Additional outgroup 25), *Limenavis patagonica* (Additional outgroup 33), *Teviornis gobiensis* (Additional outgroup 2), *Tsidiiyazhi abini* (Additional outgroup 1).

**Outgroup age sequence.** 119.6, 100.5, 80.5, 80.5, 72.1, 69, 62.221.

##### Outgroup sequence 5

**Reference phylogeny.** Wang et al., 2020b, Figure 6.

**Preliminary outgroup sequence.**

| Rank | Original taxon label | Reconciled taxon label | Age (Ma) |
|---|---|---|---|
| 1 | [Multiple taxa of Hesperornithiformes] | *Enaliornis barretti* + *Enaliornis sedgwicki* + *Enaliornis seeleyi* | 100.5 |
| 2 | *Ichthyornis* | *Ichthyornis dispar* | 80.5 |
| 3 | *Gansus* | *Gansus yumenensis* | 114.1 |
| 4 | *Iteravis* | *Iteravis huchzermeyeri* | 119.6 |
| 5 | *Yixianornis* | *Yixianornis grabaui* | 119.6 |
| 6 | *Piscivoravis* | *Piscivoravis lii* | 119.6 |

**Final outgroup sequence.** *Piscivoravis lii* (Additional outgroup 48), *Yixianornis grabaui* (Additional outgroup 58), *Iteravis huchzermeyeri* (Additional outgroup 28), *Gansus yumenensis* (Additional outgroup 22), *Enaliornis* spp. (Additional outgroup 18), *Teviornis gobiensis* (Additional outgroup 2), *Tsidiiyazhi abini* (Additional outgroup 1).

**Outgroup age sequence.** 119.6, 119.6, 119.6, 114.1, 100.5, 69, 62.221.

##### Outgroup sequence 6

**Reference phylogeny.** Zheng et al., 2018, Figure 6.

**Preliminary outgroup sequence.**

| Rank | Original taxon label | Reconciled taxon label | Age (Ma) |
|---|---|---|---|
| 1 | [Multiple taxa of Hesperornithiformes] | *Enaliornis barretti*<br>+ *Enaliornis sedgwicki*<br>+ *Enaliornis seeleyi* | 100.5 |
| 1 | *Limenavis* | *Limenavis patagonica* | 72.1 |
| 2 | *Hollanda* | *Hollanda luceria* | 71.449 |
| 2 | *Ichthyornis* | *Ichthyornis dispar* | 80.5 |
| 3 | *Songlingornis* | *Songlingornis linghensis* | 119.6 |
| 3 | *Yanornis* | *Yanornis martini* | 119.6 |
| 3 | *Yixianornis* | *Yixianornis grabaui* | 119.6 |
| 3 | *Parahongshanornis* | *Parahongshanornis chaoyangensis* | 119.6 |
| 3 | *Archaeornithura* | *Archaeornithura meemannae* | 130.7 |
| 3 | *Longicrusavis* | *Longicrusavis houi* | 122.6 |
| 3 | *Hongshanornis* | *Hongshanornis longicresta* | 125.0 |
| 3 | *Ambiortus* | *Ambiortus dementjevi* | 125.0 |
| 3 | *Apsaravis* | *Apsaravis ukhaana* | 71.9 |
| 3 | *Gansus* | *Gansus yumenensis* | 114.1 |
| 4 | *Dingavis* | *Dingavis longimaxilla* | 119.6 |
| 4 | *Iteravis* | *Iteravis huchzermeyeri* | 119.6 |
| 4 | *Chaoyangia* | *Chaoyangia beishanensis* | 119.6 |
| 4 | *Xinghaiornis* | *Xinghaiornis lini* | 119.6 |
| 4 | *Eogranivora* STM35-3 | *Eogranivora edentulata* | 122.6 |
| 4 | *Zhongjianornis* | *Zhongjianornis yangi* | 119.6 |
| 4 | *Varona* [*sic*] | *Vorona berivotrensis* | 69.6 |
| 4 | *Schizooura* | *Schizooura lii* | 119.6 |
| 4 | *Jianchangornis* | *Jianchangornis microdonta* | 119.6 |
| 5 | *Archaeorhynchus* | *Archaeorhynchus spathula* | 121.8 |
| 5 | *Patagopteryx* | *Patagopteryx deferrariisi* | 83.4 |
| 6 | [Multiple taxa of Enantiornithes] | *Eopengornis martini* | 130.7 |
| 7 | *Sapeornis* | *Sapeornis chaoyangensis* | 125.0 |
| 7 | *Didactylornis* | | |
| 8 | [Multiple taxa of Confuciusornithiformes] | *Eoconfuciusornis zhengi* | 130.7 |
| 9 | *Jeholornis* | *Jeholornis curvipes* | 122.6 |
| 10 | *Archaeopteryx* | *Archaeopteryx* sp. | 151.2 |

**Final outgroup sequence.** *Archaeopteryx* sp. (Additional outgroup 6), *Eoconfuciusornis zhengi* (Additional outgroup 19), *Eopengornis martini* (Additional outgroup 21), *Archaeornithura meemannae* (Additional outgroup 8), *Enaliornis* spp. (Additional outgroup 18), *Teviornis gobiensis* (Additional outgroup 2), *Tsidiiyazhi abini* (Additional outgroup 1).

**Outgroup age sequence.** 151.2, 130.7, 130.7, 130.7, 100.5, 69, 62.221.

##### Outgroup sequence 7

**Reference phylogeny.** Hartman et al., 2019, Figure S2.

**Preliminary outgroup sequence.**

| Rank | Original taxon label | Reconciled taxon label | Age (Ma) |
|---|---|---|---|
| 0 | *saltariensis* [as a paleognath] | "*Martinavis*" *saltariensis* | 71.5 |
| 0 | *Limenavis* [as a paleognath] | *Limenavis patagonica* | 72.1 |
| 1 | *Lithornis* | *Lithornis celetius* | 60.21 |
| 2 | *Apsaravis* | *Apsaravis ukhaana* | 71.9 |
| 3 | *Palintropus* | *Palintropus* sp. | 75.762 |
| 3 | *Iaceornis* | *Iaceornis marshi* | 80.5 |
| 3 | *Qinornis* | *Qinornis paleocenica* | 62.22 |
| 4 | *Eogranivora* | *Eogranivora edentulata* | 122.6 |
| 5 | *Xinghaiornis* | *Xinghaiornis lini* | 119.6 |
| 6 | [Multiple taxa of Hesperornithiformes] | *Enaliornis barretti* + *Enaliornis sedgwicki* + *Enaliornis seeleyi* | 100.5 |
| 6 | *Ichthyornis* | *Ichthyornis dispar* | 80.5 |
| 6 | *Mystiornis* | *Mystiornis cyrili* | 113.0 |
| 7 | *Piscivoravis* | *Piscivoravis lii* | 119.6 |
| 7 | *Juehuaornis* | *Juehuaornis zhangi* | 119.6 |
| 7 | *Dingavis* | *Dingavis longimaxilla* | 119.6 |
| 7 | *whetstonei* | "*Martinavis*" *whetstonei* | 71.5 |
| 7 | *Ambiortus* | *Ambiortus dementjevi* | 125.0 |
| 7 | *Gansus* | *Gansus yumenensis* | 114.1 |
| 7 | *Yumenornis* | *Yumenornis huangi* | 114.1 |
| 7 | *Iteravis* | *Iteravis huchzermeyeri* | 119.6 |
| 8 | *Longicrusavis* | *Longicrusavis houi* | 122.6 |
| 8 | *Parahongshanornis* | *Parahongshanornis chaoyangensis* | 119.6 |
| 8 | *Tianyuornis* | *Tianyuornis cheni* | 125.0 |

**Final outgroup sequence.** *Tianyuornis cheni* (Additional outgroup 53), *Ambiortus dementjevi* (Additional outgroup 3), *Eogranivora edentulata* (Additional outgroup 20), *Iaceornis marshi* (Additional outgroup 25), *Limenavis patagonica* (Additional outgroup 33), *Teviornis gobiensis* (Additional outgroup 2), *Tsidiiyazhi abini* (Additional outgroup 1).

**Outgroup age sequence.** 125.0, 125.0, 122.6, 80.5, 72.1, 69, 62.221.

**Outgroup sequence 8**

**Reference phylogeny.** McLachlan et al., 2017, Figure 7B.

**Preliminary outgroup sequence.**

| Rank | Original taxon label | Reconciled taxon label | Age (Ma) |
|---|---|---|---|
| 1 | Ornithurine F | Ornithurae *incertae sedis* (“Ornithurine F” of Longrich et al., 2011) | 66.2 |
| 1 | Ornithurine C | Ornithurae *incertae sedis* (“Ornithurine C” of Longrich et al., 2011) | 66.0 |
| 1 | Ornithurine B | Ornithurae *incertae sedis* (“Ornithurine B” of Longrich et al., 2011) | 66.0 |
| 1 | *Cimolopteryx maxima* | “*Cimolopteryx*” *maxima* | 66.0 |
| 1 | *Ceramornis major* | *Ceramornis major* | 66.2 |
| 1 | Ornithurine E | Ornithurae *incertae sedis* (“Ornithurine E” of Longrich et al., 2011) | 66.0 |
| 1 | Ornithurine A | Ornithurae *incertae sedis* (“Ornithurine A” of Longrich et al., 2011) | 66.2 |
| 1 | *Cimolopteryx petra* | “*Cimolopteryx*” *petra* | 66.0 |
| 1 | *Cimolopteryx minima* | “*Cimolopteryx*” *minima* | 66.0 |
| 1 | *Cimolopteryx rara* | *Cimolopteryx rara* | 66.0 |
| 2 | *Maaqwi cascadensis* | *Maaqwi cascadensis* | 71.939 |
| 2 | *Vegavis iaai* | *Vegavis iaai* | 66.5 |
| 3 | *Laceornis* [*sic*] *marshii* [*sic*] | *Iaceornis marshi* | 80.5 |
| 4 | Ornithurine D | Ichthyornithes *incertae sedis* (“Ornithurine D” of Longrich et al., 2011) | 66.0 |
| 4 | *Ichthyornis dispar* | *Ichthyornis dispar* | 80.5 |
| 5 | [Multiple taxa of Hesperornithiformes] | *Enaliornis barretti* + *Enaliornis sedgwicki* + *Enaliornis seeleyi* | 100.5 |
| 6 | *Palintropus retusus* | *Palintropus* sp. | 75.762 |
| 6 | *Apsaravis ukhaana* | *Apsaravis ukhaana* | 71.9 |
| 7 | *Yixianornis grabaui* | *Yixianornis grabaui* | 119.6 |
| 7 | *Songlingornis linghensis* | *Songlingornis linghensis* | 119.6 |
| 7 | *Yanornis martini* | *Yanornis martini* | 119.6 |

**Final outgroup sequence.** *Yixianornis grabaui* (Additional outgroup 58), *Enaliornis* spp. (Additional outgroup 18), *Ichthyornis dispar* (Additional outgroup 26), *Iaceornis marshi* (Additional outgroup 25), *Maaqwi cascadensis* (Additional outgroup 36), *Teviornis gobiensis* (Additional outgroup 2), *Tsidiiyazhi abini* (Additional outgroup 1).

**Outgroup age sequence.** 119.6, 100.5, 80.5, 80.5, 71.939, 69, 62.221.

##### Outgroup sequence 9

**Reference phylogeny.** Cordes-Person et al., 2020, Supplementary material 6.

**Preliminary outgroup sequence.**

| Rank | Original taxon label | Reconciled taxon label | Age (Ma) |
|---|---|---|---|
| 1 | *Antarcticavis* | *Antarcticavis capelambensis* | 70.8 |
| 2 | *Ichthyornis* | *Ichthyornis dispar* | 80.5 |
| 3 | [Multiple taxa of Hesperornithiformes] | *Enaliornis barretti* + *Enaliornis sedgwicki* + *Enaliornis seeleyi* | 100.5 |
| 4 | *Apsaravis* | *Apsaravis ukhaana* | 71.9 |
| 5 | *Gansus* | *Gansus yumenensis* | 114.1 |
| 6 | *Iteravis* | *Iteravis huchzermeyeri* | 119.6 |

**Final outgroup sequence.** *Iteravis huchzermeyeri* (Additional outgroup 28), *Gansus yumenensis* (Additional outgroup 22), *Enaliornis* spp. (Additional outgroup 18), *Ichthyornis dispar* (Additional outgroup 26), *Antarcticavis capelambensis* (Additional outgroup 4), *Teviornis gobiensis* (Additional outgroup 2), *Tsidiiyazhi abini* (Additional outgroup 1).

**Outgroup age sequence.** 119.6, 114.1, 100.5, 80.5, 70.8, 69, 62.221.

### 3 Supplementary Methods

In addition to the analyses described in the main text, we attempted to perform divergence time estimation on longer alignments and under more complex partitioning schemes. We first explored the possibility of using the same alignment and partitioning scheme as in our time-free analyses (25,437 sites, 22 partitions). However, the initial calculation of branch length maximum likelihood estimates (MLEs) in baseml found 112 branches to have lengths of $> 10$ expected substitutions per site, which are implausible outside of randomly generated sequences. We interpreted this as a result of overparameterization, stemming from the inability to simultaneously estimate a large quantity of branch length parameters from multiple short partitions. When attempting to use the resulting MLEs and Hessian regardless, MCMCTree aborted on failure to reset log likelihoods. Following the developers' recommendation at `http://groups.google.com/g/pamlsoftware/c/VOsptqwnhuc/m/onR9jqlyBgAJ`, we solved this issue in subsequent analyses by commenting out line 4155 of the `mcmctree.c` source file and recompiling:

```
if (fabs(lnL - lnpData(data.lnpDi)) > 0.001) {
    printf("\n%12.6f = %12.6f?  Resetting lnL\n", lnL, lnpData(data.lnpDi));
    lnL = lnpData(data.lnpDi);
    /* exit(-1); */
}
```

Next, we attempted an unpartitioned analysis of the full alignment, running 5 replicates for both the independent-rates (IR) and autocorrelated-rates (AR) clock models. All analyses were performed using a time unit of 1 Myr, setting the gamma-Dirichlet hyperprior on the mean substitution rate to `rgene_gamma = 2 427.55 1 1` and the hyperprior on log rate variance ($\sigma^2$) to `sigma2_gamma = 1 3 1`. Despite the analyses completing 5 million generations and running for up to $> 600$ hours each, the replicates failed to reach stationarity. Under the IR model, three chains were exploring higher likelihoods than the remaining two, and these sampled substantially different values for the log rate variance hyperparameter. Combining the two high-likelihood chains that sampled similar values for $\sigma^2$ yielded 21 node age parameters with effective sample sizes (ESS) $< 200$ and a potential scale reduction factor (PSRF) of $> 1.05$. Under the AR model, a single chain sampled noticeably higher likelihoods than the other four but failed to attain ESS $> 200$ for the ages of 30 out of the 335 nodes. The distribution of the node ages with low effective sample sizes differed between the two relaxed clocks; under the IR model, they were mostly associated with tips without sequence data, whose "attachment times" were entirely determined by the birth-death prior. In addition, running the analyses without data to sample from the prior revealed that the AR analyses consistently failed to reach stationarity for $\sigma^2$.

In response to these problems, we changed the time unit to 10 Myr following the recommendation that all ages in the tree be in the range of 0.01–10 units (Yang, 2020, 42), and relaxed the $\sigma^2$ hyperprior, increasing both its mean (1 vs. 1/3) and variance (1/2 vs. 1/9). Most importantly, we subsampled the original 24-locus alignment to a subset of 8 loci selected for clocklikeness, as described in the main text. This was motivated by the effort to both improve the performance of the analyses and limit the amount of rate heterogeneity present in the data, which should aid in the estimation of the relaxed clock hyperparameters.

We employed four different subsampling and partitioning strategies. In strategy (1), the 8-locus alignment was partitioned by genome (nuclear vs. mitochondrial) and codon position (1, 2, 3), with a separate partition for nuclear introns (7 partitions, 10,883 bp). In strategy (2), only the 3 mitochondrial loci (COX1, CytB, ND3) were used and partitioned by codon position (1+2 vs. 3), resulting in 2 partitions with a total length of 3113 bp. In strategy (3), the 8-locus alignment was partitioned by genome (nuclear vs. mitochondrial) and codon position (1+2 vs. 3), but only the "fast" mitochondrial-3 and "slow" nuclear-1+2 partitions were retained, yielding 2 partitions with a total length of 4262 bp. Finally, in strategy (4), the entire 8-locus alignment was used and left unpartitioned (length = 10,883 bp). The mean substitution rate hyperprior was calculated separately for each strategy as the mean of partition-specific baseml estimates weighted by partition length: (1) `rgene_gamma = 2 93.85 1`, (2) `rgene_gamma = 2 39.74 1`, (3) `rgene_gamma = 2 55.92 1`, (4) `rgene_gamma = 2 81.44`. Strategies (4) and (3) correspond to the primary and sensitivity analyses described in the main text, respectively.

Analytical strategies (1) and (2) were affected by substantial performance issues, convergence issues, or both, and their results were not summarized. The time required for the strategy (1) MCMCTree runs to reach the target length of 5 million generations (20,000 samples at the frequency of 1 sample per every 250 generations) was estimated at 5 and 8 months for the AR and IR analyses, respectively. Both analyses were terminated after 720 and 793 hours of runtime, respectively, after the AR analyses started to sample impossibly ancient root ages ($> 4 \times 10^4$ and $> 7 \times 10^4$ Ma) and large positive log likelihoods, probably due to undiagnosed numeric instabilities. Strategy (2) was less computationally intensive, with the runtime required to reach 5 million generations estimated at $\sim$1.5 months for the AR analyses and over 2 months for the IR analyses. However, a check performed after more than two weeks of runtime showed that the runs were not promising, since many parameters did not even begin to approach the effective sample size of 200 (IR: minimum ESS = 1.59, AR: minimum ESS = 2.78). Accordingly, both the IR and AR analyses performed under strategy (2) were terminated after 581 and 428 hours, respectively. The remaining analyses reached convergence for all (strategy 4) or all but two (strategy 3) parameters as assessed by the ESS and PSRF criteria, and were summarized as reported in the main text. A detailed comparison of the results is included in Table A.1.

### 4 Supplementary Figures and Tables

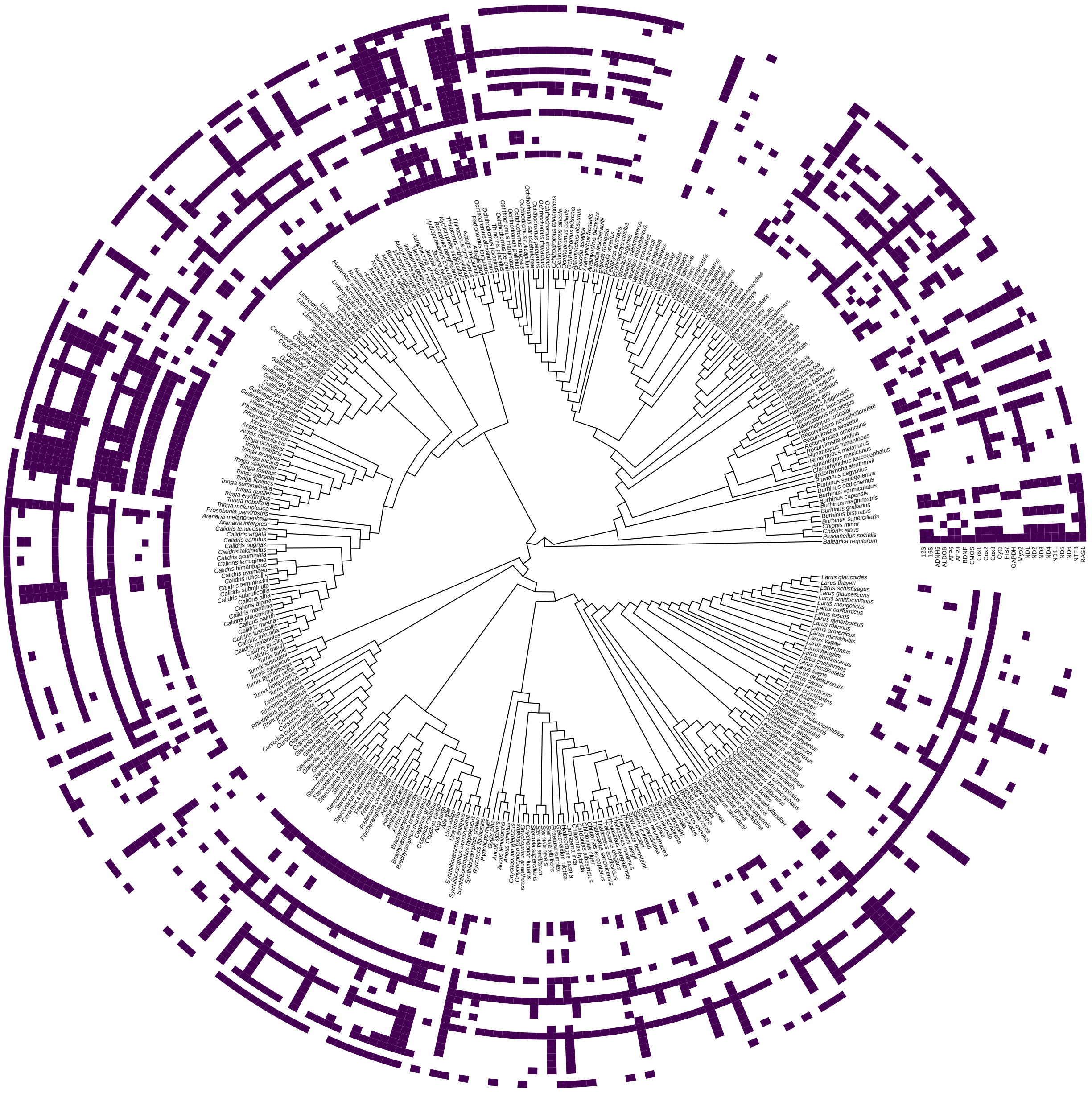

**Figure A.2:** Occupancy map for the 24 loci employed in the concatenated analyses, plotted next to the tips of the 337-species total-evidence (RAxML-NG) tree. Filled cells indicate the presence of sequence data for a given gene; white cells indicate missing data. The phylogenetic positions of the 31 species without sequence data were informed by morphology.

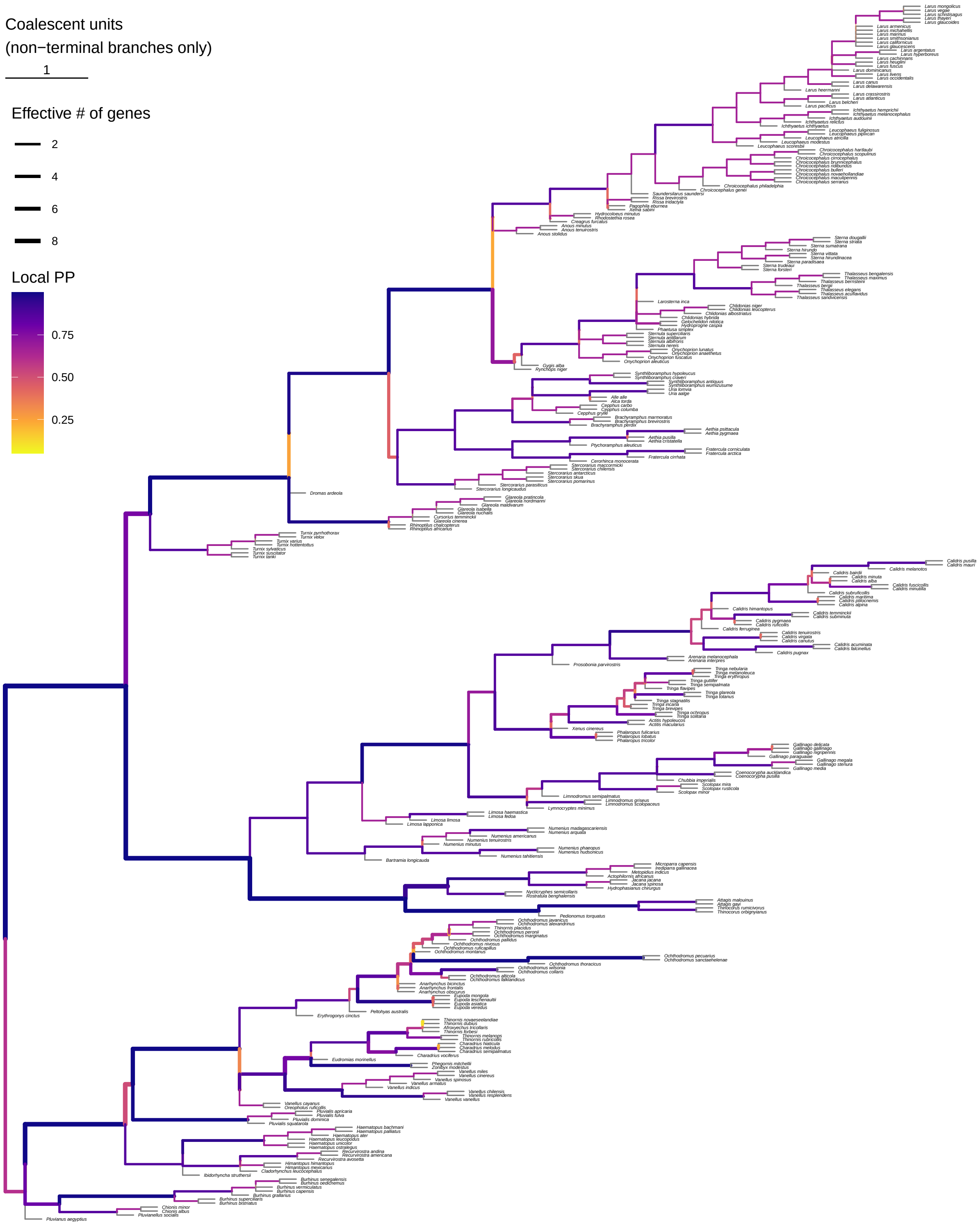

**Figure A.3:** Full 305-tip species tree inferred using ASTRAL-III, plotted as a phylogram with branch lengths in coalescent units. Local PP = local posterior probability. The effective number of genes is calculated out of a maximum of 10 (mitochondrial genome treated as a single unit).

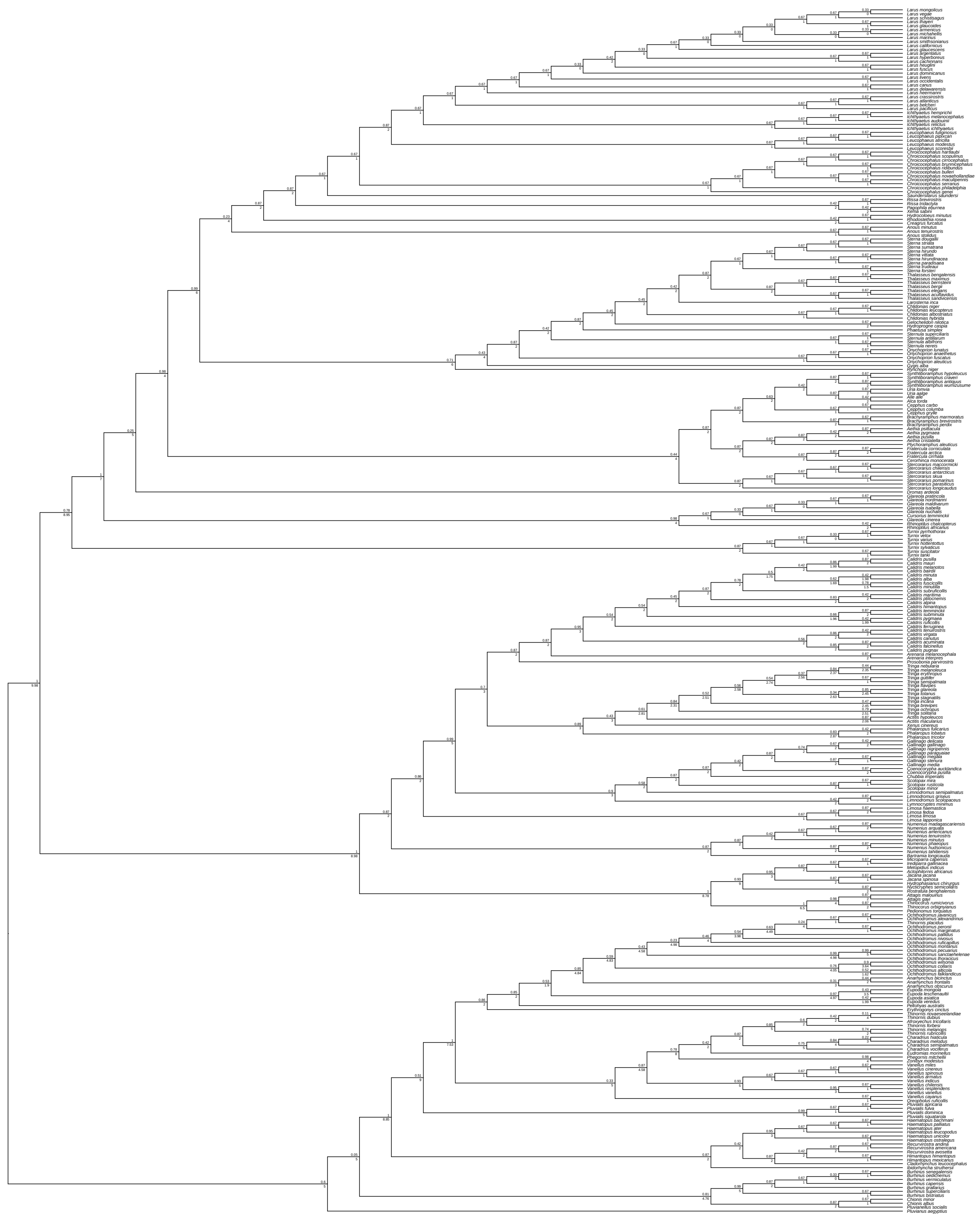

**Figure A.4:** Full 305-tip species tree inferred using ASTRAL-III, plotted as a cladogram for easier viewing of the branching sequence and branch annotations. Numbers above each node indicate local posterior probabilities; numbers below each node indicate the effective number of genes calculated out of a maximum of 10 (mitochondrial genome treated as a single unit).

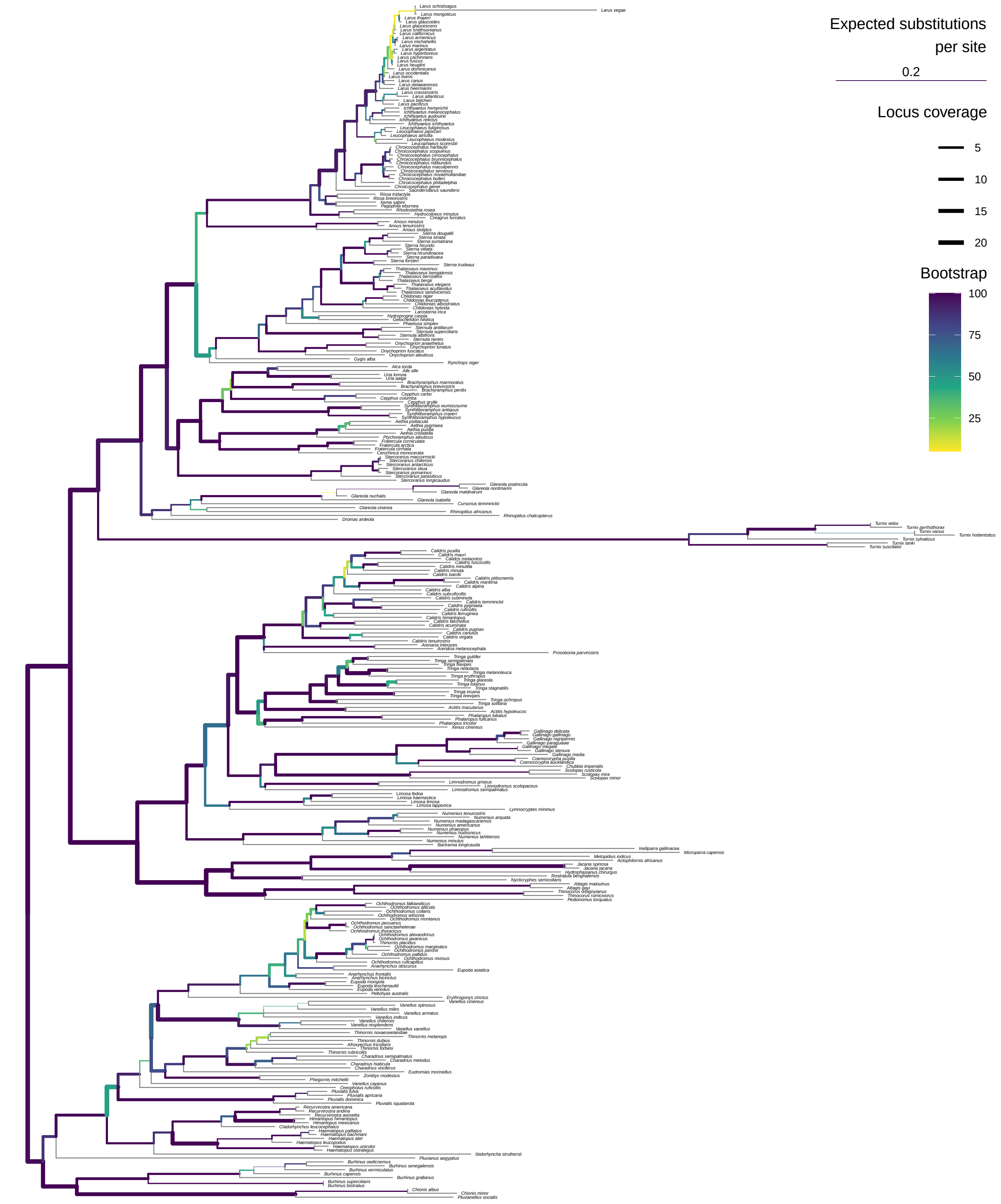

**Figure A.5:** Full 305-tip phylogeny inferred using a RAxML analysis of concatenated data, plotted as a phylogram with branch lengths in units of expected substitutions per site. The per-branch locus coverage is calculated out of a maximum of 24 (mitochondrial loci analyzed separately).

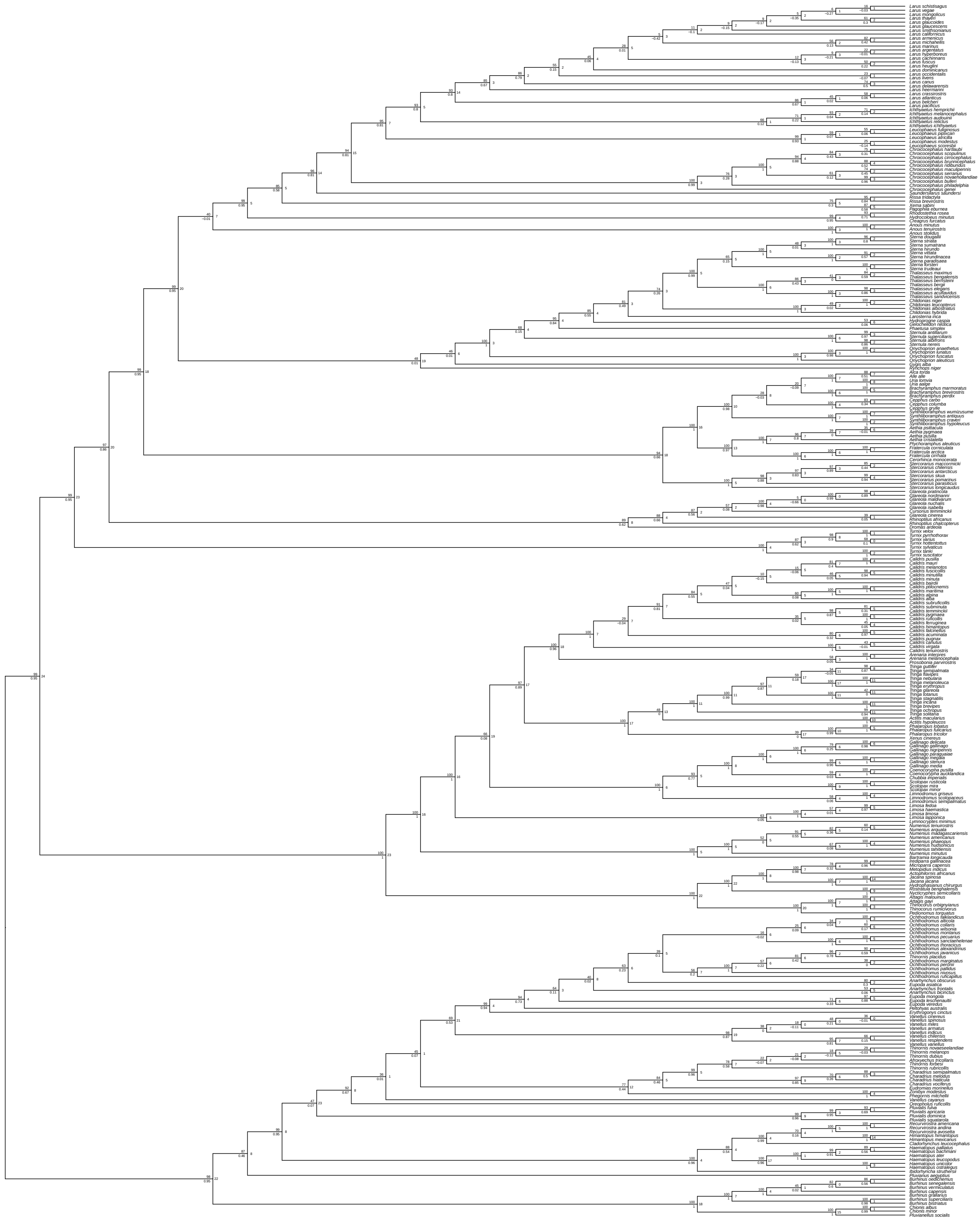

**Figure A.6:** Full 305-tip phylogeny inferred using a RAxML analysis of concatenated data, plotted as a cladogram for easier viewing of the branching sequence and branch annotations. Numbers above each node indicate bootstrap support; numbers below each node indicate internode certainty; numbers to the right of each node indicate the per-branch locus coverage calculated out of a maximum of 24 (mitochondrial loci analyzed separately).

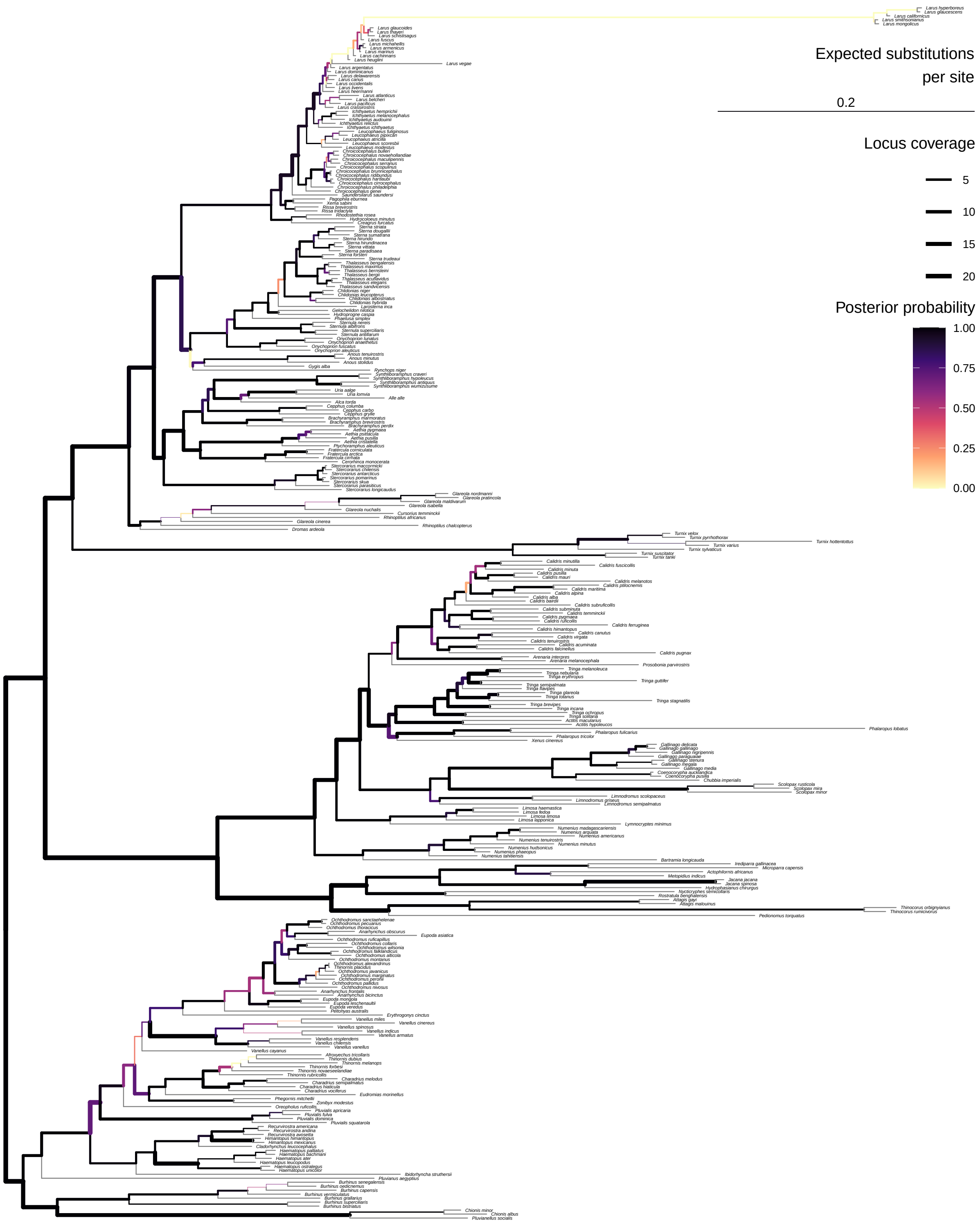

**Figure A.7:** Full 305-tip phylogeny inferred using an ExaBayes analysis of concatenated data, plotted as a phylogram with branch lengths in units of expected substitutions per site. The per-branch locus coverage is calculated out of a maximum of 24 (mitochondrial loci analyzed separately).

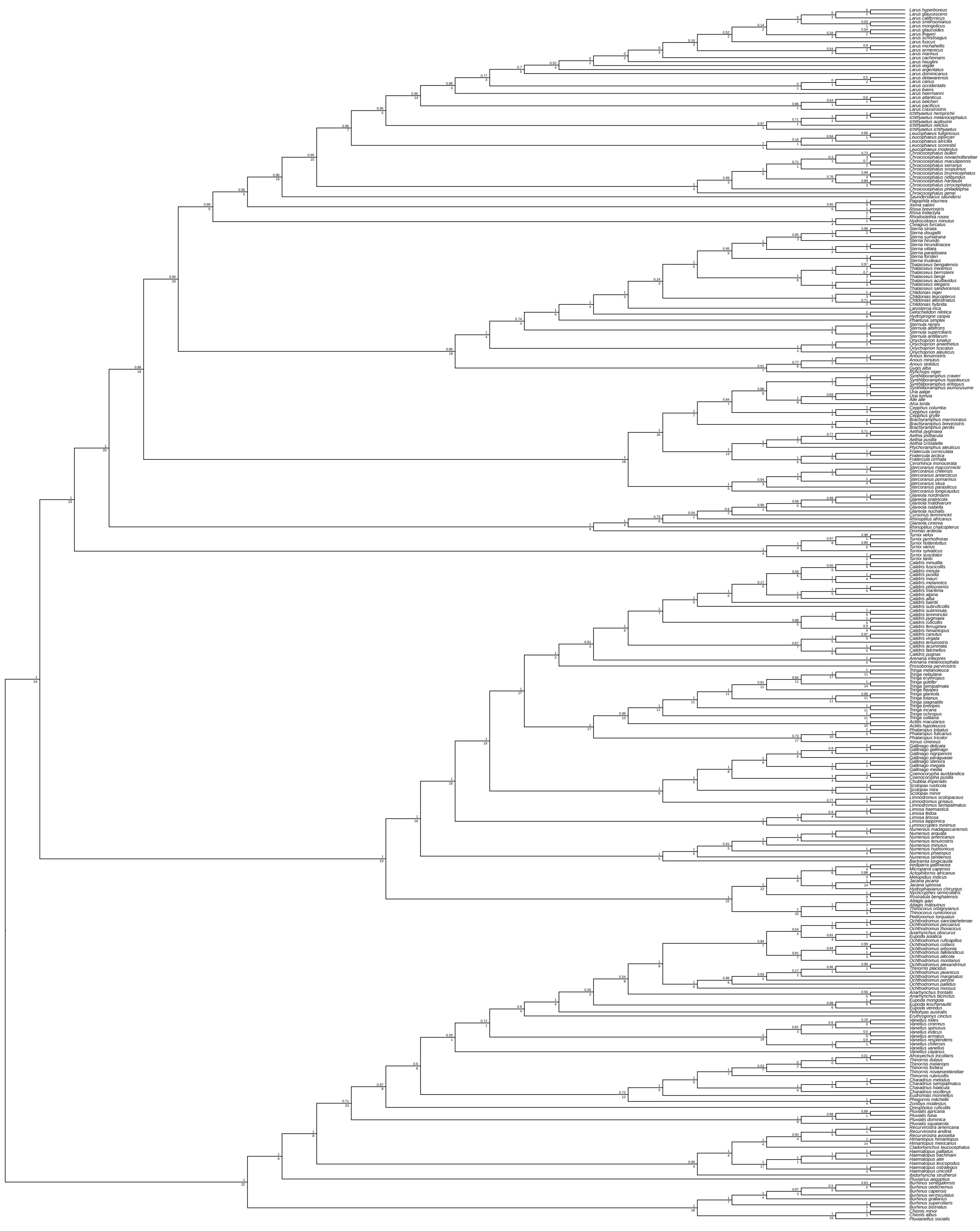

**Figure A.8:** Full 305-tip phylogeny inferred using an ExaBayes analysis of concatenated data, plotted as a cladogram for easier viewing of the branching sequence and branch annotations. Numbers above each node indicate posterior probabilities; numbers below each node indicate the per-branch locus coverage calculated out of a maximum of 24 (mitochondrial loci analyzed separately).

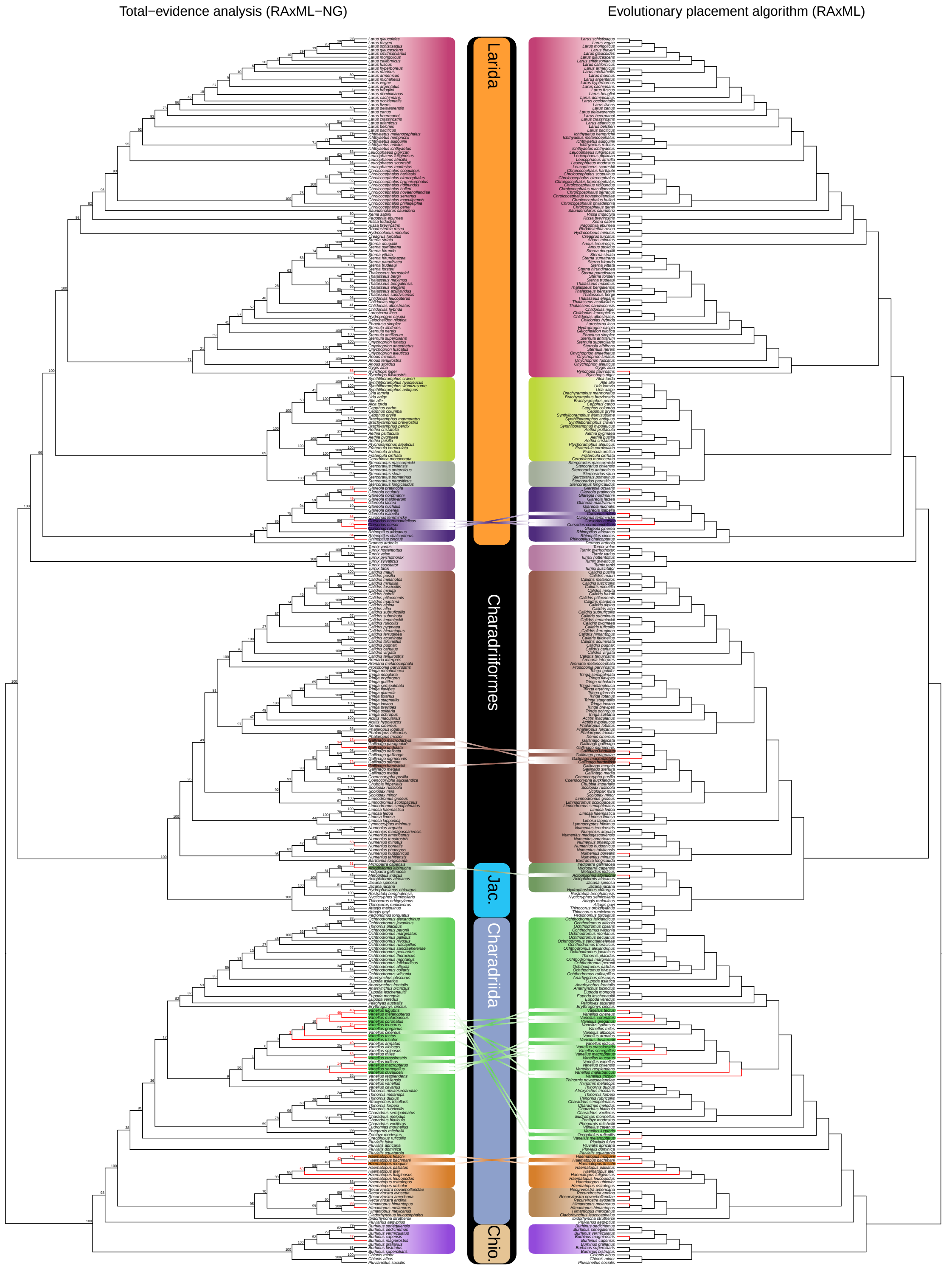

**Figure A.9:** Comparison of topologies resulting from the total-evidence (TE) and evolutionary placement algorithm (EPA) analyses of combined data, plotted as cladograms for easier viewing of the branching sequence. Numbers above each node of the TE tree indicate bootstrap support (BS). Red indicates tips without sequence data and BS values for the nodes at which they attach to the tree. Conflicting positions of the morphology-only tips are indicated with lines crossing the midline. Jac. = Jacanida; Chio. = Chionida.

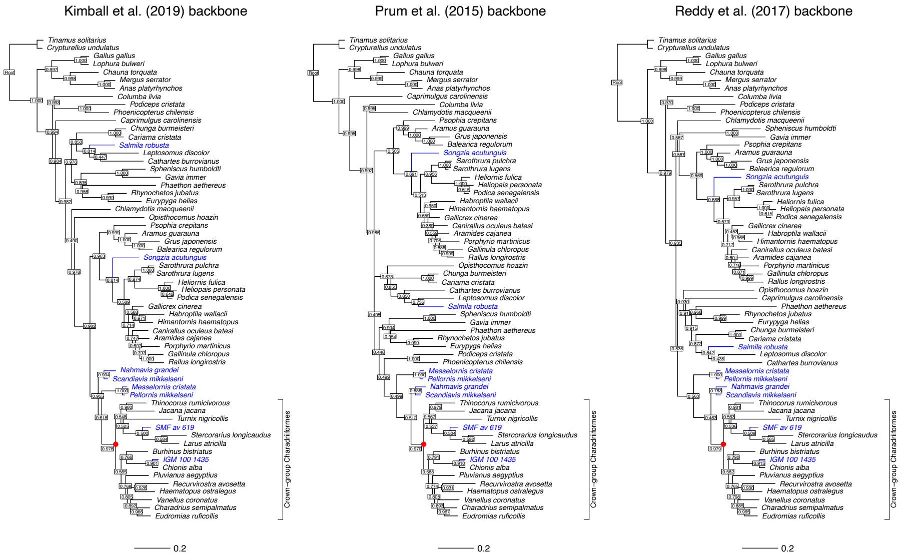

**Figure A.10:** Phylogenies inferred using Bayesian re-analyses of the morphological data from Musser and Clarke (2020) under topological constraints derived from recent phylogenomic studies. Node labels indicate posterior probabilities; filled red circles denote the node corresponding to crown-group Charadriiformes. Extant and fossil taxa are shown in black and blue, respectively. The scale bar represents 0.2 expected substitutions per character.

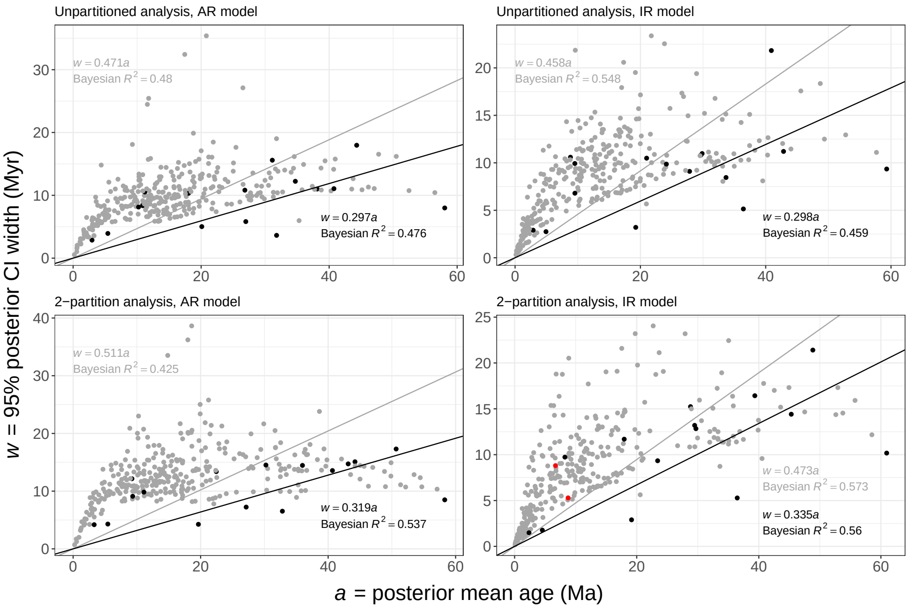

**Figure A.11:** Infinite-site plots of posterior credibility interval (CI) width ($w$) against posterior mean node ages ($a$). Black and gray data points represent the calibrated and uncalibrated nodes, respectively. Red data points in the bottom right panel denote the two nodes whose age estimates did not reach an effective sample size of $> 200$ in the 2-partition IR analysis. Gray and black lines and equations represent Bayesian simple linear regression models through the origin (noninformative priors, 20,000 iterations), fitted to all nodes and the calibrated nodes only, respectively. AR = autocorrelated rates; IE = independent rates.

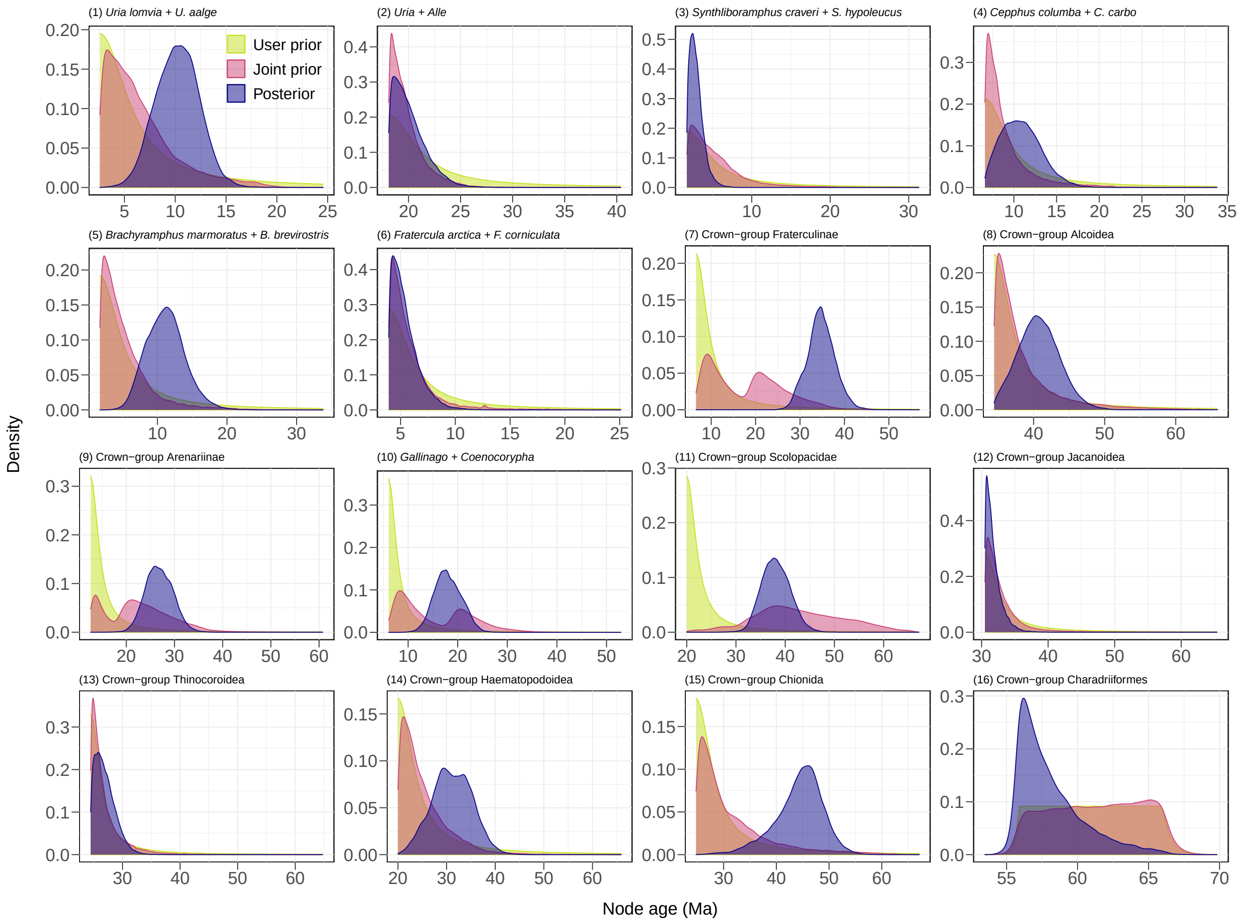

**Figure A.12:** Probability density functions for the ages of the 16 calibrated nodes (cf. Figure 7 in the main text). "User prior" = user-specified calibration densities (truncated Cauchy for calibrations 1 through 15, soft-bounded uniform for calibration 16); "Joint prior" = effective prior resulting from calibration interactions under the AR model; "Posterior" = marginal posterior under the AR model.

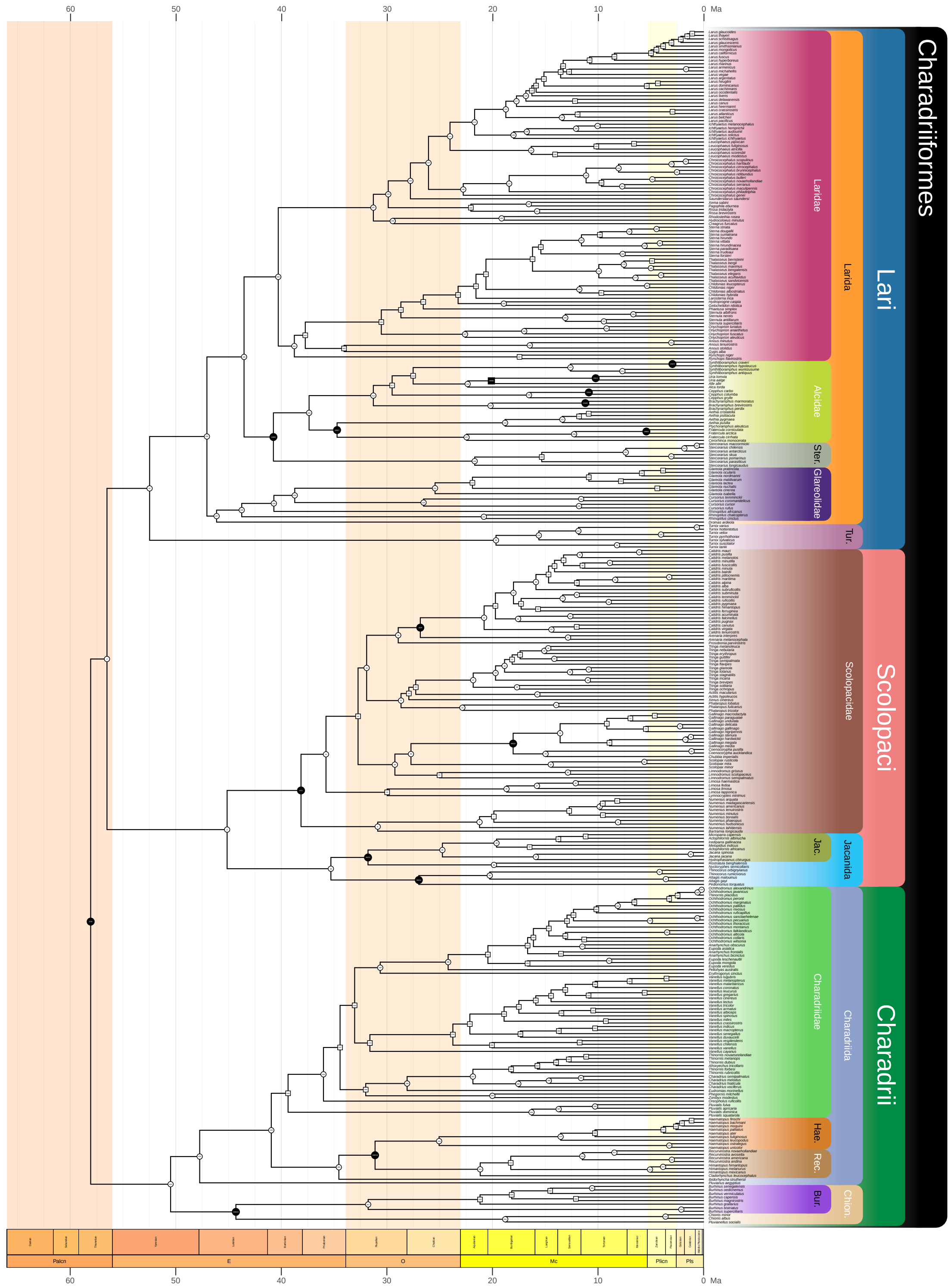

**Figure A.13:** Time-calibrated phylogeny of 336 species of shorebirds based on the Bayesian node-dating analysis of 8 clock-like loci under the autocorrelated-rates relaxed clock (cf. Figure 5 in the main text). Nodes with bootstrap support ≥70% are indicated by circles; nodes with bootstrap support <70% are indicated by squares. Fossil-calibrated nodes are shown in black. Shaded tabs represent higher-level clades; background shading indicates geochronological epochs. Calibration numbers and abbreviations as in Figure 5.

**Table A.1:** Comparison of the ages inferred for major nodes within Charadriiformes under different relaxed clock models and partitioning schemes. Node numbers correspond to those in Figure 5 and Figure A.13. Blue indicates calibrated nodes; red indicates the two age estimates that did not reach an effective sample size of > 200 in the 2-partition independent-rates analysis. AR = autocorrelated rates; IE = independent rates; CI = credibility interval.

| Node | Clade | Analysis | Posterior mean age (Ma) | 95% CI (Ma) |
|---|---|---|---|---|
| 1 | Root | Unpartitioned, IR | 59.31 | [55.42, 64.77] |
| | | Unpartitioned, AR | 58.06 | [54.97, 62.96] |
| | | 2 partitions, IR | 60.95 | [55.85, 66.03] |
| | | 2 partitions, AR | 58.29 | [55.11, 63.61] |
| 2 | (Scolopaci + Lari) | Unpartitioned, IR | 57.68 | [52.77, 63.87] |
| | | Unpartitioned, AR | 56.51 | [51.71, 62.14] |
| | | 2 partitions, IR | 58.53 | [52.65, 64.83] |
| | | 2 partitions, AR | 57.10 | [52.57, 62.92] |
| 97 | Lari | Unpartitioned, IR | 52.77 | [46.56, 59.50] |
| | | Unpartitioned, AR | 52.48 | [47.45, 58.20] |
| | | 2 partitions, IR | 53.77 | [46.64, 61.17] |
| | | 2 partitions, AR | 54.70 | [49.86, 60.61] |
| 3 | Scolopaci | Unpartitioned, IR | 49.37 | [43.27, 55.77] |
| | | Unpartitioned, AR | 45.14 | [39.48, 50.35] |
| | | 2 partitions, IR | 52.62 | [45.50, 59.87] |
| | | 2 partitions, AR | 49.87 | [42.30, 57.23] |
| 242 | Charadrii | Unpartitioned, IR | 48.71 | [39.42, 57.77] |
| | | Unpartitioned, AR | 50.49 | [42.00, 58.20] |
| | | 2 partitions, IR | 55.76 | [47.97, 63.88] |
| | | 2 partitions, AR | 54.45 | [48.60, 60.93] |
| 243 | Charadriida | Unpartitioned, IR | 45.61 | [37.10, 54.67] |
| | | Unpartitioned, AR | 47.73 | [38.53, 55.07] |
| | | 2 partitions, IR | 52.95 | [44.34, 61.66] |
| | | 2 partitions, AR | 53.48 | [47.36, 60.23] |
| 104 | Larida | Unpartitioned, IR | 44.03 | [38.51, 49.97] |
| | | Unpartitioned, AR | 47.05 | [41.63, 52.71] |
| | | 2 partitions, IR | 47.52 | [40.39, 55.05] |
| | | 2 partitions, AR | 52.16 | [47.26, 57.97] |
| 4 | Scolopacidae | Unpartitioned, IR | 42.85 | [37.41, 48.61] |
| | | Unpartitioned, AR | 38.13 | [32.73, 43.73] |
| | | 2 partitions, IR | 45.32 | [38.23, 52.65] |
| | | 2 partitions, AR | 44.21 | [36.61, 51.70] |
| 227 | Glareoloidea | Unpartitioned, IR | 42.70 | [37.10, 49.13] |
| | | Unpartitioned, AR | 46.13 | [40.45, 51.77] |
| | | 2 partitions, IR | 44.94 | [36.56, 53.88] |
| | | 2 partitions, AR | 51.37 | [46.27, 57.38] |

| | | | | |
|---|---|---|---|---|
| 326 | Chionida | Unpartitioned, IR | 40.90 | [30.15, 51.98] |
| | | Unpartitioned, AR | 44.30 | [34.41, 52.38] |
| | | 2 partitions, IR | 48.85 | [37.68, 59.09] |
| | | 2 partitions, AR | 50.67 | [41.97, 59.28] |
| 228 | Glareolidae | Unpartitioned, IR | 40.37 | [34.29, 46.94] |
| | | Unpartitioned, AR | 43.75 | [37.28, 49.93] |
| | | 2 partitions, IR | 40.79 | [32.21, 49.98] |
| | | 2 partitions, AR | 49.85 | [43.75, 56.36] |
| 105 | (Alcoidea + Laridae) | Unpartitioned, IR | 39.53 | [35.72, 43.82] |
| | | Unpartitioned, AR | 43.53 | [38.07, 49.06] |
| | | 2 partitions, IR | 40.53 | [36.04, 45.62] |
| | | 2 partitions, AR | 49.00 | [43.89, 54.67] |
| 83 | Jacanida | Unpartitioned, IR | 38.27 | [32.90, 44.22] |
| | | Unpartitioned, AR | 35.30 | [32.56, 38.53] |
| | | 2 partitions, IR | 42.54 | [33.78, 50.79] |
| | | 2 partitions, AR | 35.04 | [31.17, 40.09] |
| 245 | Charadriidae | Unpartitioned, IR | 37.43 | [31.47, 43.28] |
| | | Unpartitioned, AR | 39.36 | [31.56, 47.05] |
| | | 2 partitions, IR | 39.94 | [33.28, 47.03] |
| | | 2 partitions, AR | 45.64 | [38.21, 52.51] |
| 198 | Alcoidea | Unpartitioned, IR | 36.44 | [34.44, 39.58] |
| | | Unpartitioned, AR | 40.75 | [35.07, 46.13] |
| | | 2 partitions, IR | 36.47 | [34.44, 39.72] |
| | | 2 partitions, AR | 43.15 | [35.12, 49.85] |
| 106 | Laridae | Unpartitioned, IR | 34.98 | [30.15, 39.79] |
| | | Unpartitioned, AR | 40.29 | [34.98, 45.69] |
| | | 2 partitions, IR | 35.04 | [29.56, 40.71] |
| | | 2 partitions, AR | 46.32 | [41.21, 52.07] |
| 84 | Jacanoidea | Unpartitioned, IR | 33.67 | [30.50, 38.95] |
| | | Unpartitioned, AR | 31.79 | [30.50, 34.13] |
| | | 2 partitions, IR | 39.35 | [30.84, 47.27] |
| | | 2 partitions, AR | 32.82 | [30.50, 37.03] |
| 5 | Numeniinae | Unpartitioned, IR | 33.64 | [25.98, 41.12] |
| | | Unpartitioned, AR | 30.87 | [24.43, 37.22] |
| | | 2 partitions, IR | 33.92 | [24.79, 43.73] |
| | | 2 partitions, AR | 36.41 | [28.41, 45.65] |
| 65 | Scolopacinae | Unpartitioned, IR | 33.17 | [27.68, 38.27] |
| | | Unpartitioned, AR | 29.24 | [24.30, 34.73] |
| | | 2 partitions, IR | 35.82 | [30.10, 42.12] |
| | | 2 partitions, AR | 35.36 | [28.96, 42.51] |
| 205 | Alcidae | Unpartitioned, IR | 33.16 | [29.12, 37.17] |
| | | Unpartitioned, AR | 37.37 | [31.68, 43.12] |
| | | 2 partitions, IR | 33.63 | [29.27, 37.99] |
| | | 2 partitions, AR | 38.97 | [32.32, 45.95] |

| | | | | |
|---|---|---|---|---|
| 21 | Tringinae | Unpartitioned, IR | 32.44 | [27.32, 37.49] |
| | | Unpartitioned, AR | 28.66 | [23.81, 33.78] |
| | | 2 partitions, IR | 33.97 | [28.40, 39.78] |
| | | 2 partitions, AR | 32.67 | [25.85, 39.76] |
| 15 | Limosinae | Unpartitioned, IR | 31.91 | [23.14, 39.93] |
| | | Unpartitioned, AR | 29.98 | [23.89, 36.28] |
| | | 2 partitions, IR | 35.82 | [28.47, 43.79] |
| | | 2 partitions, AR | 34.78 | [26.67, 42.96] |
| 24 | Tringini | Unpartitioned, IR | 31.37 | [26.12, 36.25] |
| | | Unpartitioned, AR | 27.92 | [23.22, 33.12] |
| | | 2 partitions, IR | 31.96 | [26.48, 37.84] |
| | | 2 partitions, AR | 30.51 | [23.24, 37.15] |
| 66 | Scolopacini | Unpartitioned, IR | 31.08 | [25.75, 36.33] |
| | | Unpartitioned, AR | 27.72 | [22.52, 32.92] |
| | | 2 partitions, IR | 33.32 | [27.29, 39.79] |
| | | 2 partitions, AR | 33.22 | [26.34, 40.19] |
| 273 | *Vanellus* | Unpartitioned, IR | 30.93 | [25.52, 35.98] |
| | | Unpartitioned, AR | 31.61 | [24.60, 38.80] |
| | | 2 partitions, IR | 32.68 | [26.88, 38.75] |
| | | 2 partitions, AR | 36.77 | [29.17, 45.53] |
| 294 | Charadriinae | Unpartitioned, IR | 30.87 | [25.28, 36.42] |
| | | Unpartitioned, AR | 32.00 | [25.35, 39.18] |
| | | 2 partitions, IR | 33.32 | [27.54, 39.21] |
| | | 2 partitions, AR | 36.61 | [29.34, 45.65] |
| 309 | (Haematopodoidea + *Ibidorhyncha*) | Unpartitioned, IR | 30.23 | [22.26, 37.47] |
| | | Unpartitioned, AR | 34.55 | [26.13, 42.29] |
| | | 2 partitions, IR | 33.18 | [24.28, 41.52] |
| | | 2 partitions, AR | 44.24 | [37.08, 51.46] |
| 40 | Arenariinae | Unpartitioned, IR | 29.85 | [24.25, 35.22] |
| | | Unpartitioned, AR | 26.83 | [21.58, 32.40] |
| | | 2 partitions, IR | 29.49 | [22.67, 35.87] |
| | | 2 partitions, AR | 30.24 | [22.71, 37.23] |
| 206 | Fraterculinae | Unpartitioned, IR | 29.64 | [24.05, 34.85] |
| | | Unpartitioned, AR | 34.72 | [28.32, 40.56] |
| | | 2 partitions, IR | 28.81 | [20.10, 35.33] |
| | | 2 partitions, AR | 35.98 | [28.80, 43.26] |
| 249 | Anarhynchinae | Unpartitioned, IR | 29.52 | [23.97, 34.72] |
| | | Unpartitioned, AR | 30.63 | [24.08, 37.65] |
| | | 2 partitions, IR | 31.98 | [25.81, 38.34] |
| | | 2 partitions, AR | 36.26 | [28.71, 45.42] |
| 214 | Alcinae | Unpartitioned, IR | 29.00 | [24.37, 33.59] |
| | | Unpartitioned, AR | 31.28 | [25.85, 36.34] |
| | | 2 partitions, IR | 29.52 | [23.64, 34.50] |
| | | 2 partitions, AR | 33.44 | [26.87, 39.75] |

| | | | | |
|---|---|---|---|---|
| 327 | Burhinidae | Unpartitioned, IR | 28.97 | [18.88, 38.28] |
| | | Unpartitioned, AR | 31.77 | [22.37, 41.39] |
| | | 2 partitions, IR | 35.06 | [24.45, 46.90] |
| | | 2 partitions, AR | 38.64 | [27.61, 51.42] |
| 93 | Thinocoroidea | Unpartitioned, IR | 27.85 | [24.47, 33.57] |
| | | Unpartitioned, AR | 26.98 | [24.47, 30.29] |
| | | 2 partitions, IR | 29.70 | [24.47, 37.31] |
| | | 2 partitions, AR | 27.14 | [24.47, 31.71] |
| 86 | Jacanidae | Unpartitioned, IR | 27.32 | [21.40, 33.70] |
| | | Unpartitioned, AR | 24.74 | [20.24, 28.84] |
| | | 2 partitions, IR | 30.28 | [22.08, 38.03] |
| | | 2 partitions, AR | 21.54 | [15.59, 27.51] |
| 98 | Turnicidae | Unpartitioned, IR | 26.62 | [18.11, 35.46] |
| | | Unpartitioned, AR | 19.66 | [11.84, 27.59] |
| | | 2 partitions, IR | 23.12 | [14.90, 33.66] |
| | | 2 partitions, AR | 21.47 | [12.33, 32.25] |
| 166 | Sterninae | Unpartitioned, IR | 25.79 | [20.97, 30.98] |
| | | Unpartitioned, AR | 30.53 | [24.59, 36.17] |
| | | 2 partitions, IR | 20.69 | [16.02, 26.14] |
| | | 2 partitions, AR | 35.30 | [28.40, 42.22] |
| 310 | Haematopodoidea | Unpartitioned, IR | 24.15 | [20.00, 29.86] |
| | | Unpartitioned, AR | 31.12 | [23.05, 38.65] |
| | | 2 partitions, IR | 23.43 | [20.00, 29.34] |
| | | 2 partitions, AR | 40.68 | [33.87, 47.44] |
| 27 | *Tringa* | Unpartitioned, IR | 24.05 | [19.30, 28.85] |
| | | Unpartitioned, AR | 21.82 | [17.48, 26.26] |
| | | 2 partitions, IR | 20.85 | [16.39, 25.78] |
| | | 2 partitions, AR | 20.26 | [14.68, 25.77] |
| 42 | *Calidris* | Unpartitioned, IR | 23.69 | [19.45, 28.24] |
| | | Unpartitioned, AR | 20.79 | [16.30, 25.52] |
| | | 2 partitions, IR | 21.06 | [16.64, 26.21] |
| | | 2 partitions, AR | 22.53 | [16.29, 28.65] |
| 69 | (*Gallinago* + *Coenocorypha*) | Unpartitioned, IR | 20.99 | [15.68, 26.17] |
| | | Unpartitioned, AR | 18.04 | [13.01, 23.33] |
| | | 2 partitions, IR | 17.95 | [12.39, 24.08] |
| | | 2 partitions, AR | 22.45 | [15.45, 28.84] |
| 219 | Alcini | Unpartitioned, IR | 20.96 | [18.44, 24.11] |
| | | Unpartitioned, AR | 22.35 | [18.82, 26.22] |
| | | 2 partitions, IR | 20.58 | [18.19, 23.82] |
| | | 2 partitions, AR | 22.07 | [18.50, 26.29] |
| 94 | Thinocoridae | Unpartitioned, IR | 20.15 | [13.96, 26.79] |
| | | Unpartitioned, AR | 20.25 | [15.48, 25.18] |
| | | 2 partitions, IR | 17.94 | [10.70, 25.31] |
| | | 2 partitions, AR | 20.24 | [13.73, 26.55] |

| | | | | |
|---|---|---|---|---|
| 85 | Rostratulidae | Unpartitioned, IR | 20.01 | [11.79, 28.94] |
| | | Unpartitioned, AR | 19.86 | [13.31, 25.95] |
| | | 2 partitions, IR | 22.70 | [11.63, 35.67] |
| | | 2 partitions, AR | 20.76 | [11.83, 29.36] |
| 220 | (*Uria* + *Alle*) | Unpartitioned, IR | 19.26 | [18.10, 21.30] |
| | | Unpartitioned, AR | 20.10 | [18.10, 23.12] |
| | | 2 partitions, IR | 19.15 | [18.10, 21.00] |
| | | 2 partitions, AR | 19.65 | [18.10, 22.36] |
| 334 | Chionoidea | Unpartitioned, IR | 19.20 | [10.21, 29.72] |
| | | Unpartitioned, AR | 18.79 | [9.55, 29.44] |
| | | 2 partitions, IR | 19.76 | [10.05, 33.25] |
| | | 2 partitions, AR | 21.25 | [9.34, 35.13] |
| 107 | Larinae | Unpartitioned, IR | 18.77 | [14.58, 23.09] |
| | | Unpartitioned, AR | 31.28 | [26.57, 36.30] |
| | | 2 partitions, IR | 13.66 | [10.12, 17.72] |
| | | 2 partitions, AR | 36.45 | [31.49, 41.61] |
| 207 | Fraterculini | Unpartitioned, IR | 18.49 | [12.36, 24.99] |
| | | Unpartitioned, AR | 22.45 | [15.67, 29.37] |
| | | 2 partitions, IR | 18.03 | [10.94, 25.81] |
| | | 2 partitions, AR | 29.44 | [20.36, 37.58] |
| 318 | Haematopodidae | Unpartitioned, IR | 17.59 | [11.69, 23.65] |
| | | Unpartitioned, AR | 25.05 | [17.17, 32.32] |
| | | 2 partitions, IR | 15.28 | [9.88, 20.69] |
| | | 2 partitions, AR | 31.74 | [22.62, 40.15] |
| 210 | Aethiini | Unpartitioned, IR | 16.86 | [11.42, 22.36] |
| | | Unpartitioned, AR | 18.81 | [12.54, 25.52] |
| | | 2 partitions, IR | 14.14 | [9.13, 19.97] |
| | | 2 partitions, AR | 18.94 | [11.14, 26.71] |
| 199 | Stercorariidae | Unpartitioned, IR | 16.69 | [10.29, 23.77] |
| | | Unpartitioned, AR | 21.68 | [13.97, 30.04] |
| | | 2 partitions, IR | 13.32 | [6.98, 20.95] |
| | | 2 partitions, AR | 23.33 | [13.70, 34.99] |
| 311 | Recurvirostridae | Unpartitioned, IR | 15.93 | [10.67, 21.08] |
| | | Unpartitioned, AR | 21.16 | [13.86, 28.92] |
| | | 2 partitions, IR | 14.41 | [9.50, 19.68] |
| | | 2 partitions, AR | 34.71 | [27.83, 41.60] |
| 257 | *Ochthodromus* | Unpartitioned, IR | 14.83 | [11.63, 18.16] |
| | | Unpartitioned, AR | 14.66 | [10.68, 18.70] |
| | | 2 partitions, IR | 12.25 | [9.26, 15.35] |
| | | 2 partitions, AR | 17.07 | [11.66, 22.55] |
| 146 | *Chroicocephalus* | Unpartitioned, IR | 10.09 | [6.57, 13.68] |
| | | Unpartitioned, AR | 22.77 | [17.70, 28.27] |
| | | 2 partitions, IR | 6.20 | [3.67, 8.96] |
| | | 2 partitions, AR | 27.94 | [20.66, 34.93] |

| | | | | |
|---|---|---|---|---|
| 226 | (*Cepphus carbo* + *C. columba*) | Unpartitioned, IR | 9.55 | [6.60, 13.40] |
| | | Unpartitioned, AR | 10.87 | [6.67, 15.03] |
| | | 2 partitions, IR | 8.76 | [6.60, 11.89] |
| | | 2 partitions, AR | 11.10 | [6.60, 16.43] |
| 221 | (*Uria lomvia* + *U. aalge*) | Unpartitioned, IR | 9.54 | [4.50, 14.42] |
| | | Unpartitioned, AR | 10.22 | [6.14, 14.27] |
| | | 2 partitions, IR | 6.70 | [3.49, 12.30] |
| | | 2 partitions, AR | 9.34 | [4.81, 13.92] |
| 216 | (*Brachyramphus marmoratus* + *B. brevirostris*) | Unpartitioned, IR | 8.84 | [3.69, 14.28] |
| | | Unpartitioned, AR | 11.20 | [5.96, 16.49] |
| | | 2 partitions, IR | 8.24 | [3.62, 13.36] |
| | | 2 partitions, AR | 9.22 | [3.40, 15.56] |
| 113 | *Larus* | Unpartitioned, IR | 5.88 | [3.94, 7.97] |
| | | Unpartitioned, AR | 18.73 | [15.14, 22.34] |
| | | 2 partitions, IR | 3.24 | [2.16, 4.45] |
| | | 2 partitions, AR | 25.61 | [20.75, 30.57] |
| 209 | (*Fratercula arctica* + *F. corniculata*) | Unpartitioned, IR | 4.93 | [3.92, 6.67] |
| | | Unpartitioned, AR | 5.42 | [3.92, 7.85] |
| | | 2 partitions, IR | 4.54 | [3.92, 5.69] |
| | | 2 partitions, AR | 5.44 | [3.92, 8.22] |
| 335 | Chionidae | Unpartitioned, IR | 3.18 | [0.01, 8.10] |
| | | Unpartitioned, AR | 3.60 | [0.01, 9.02] |
| | | 2 partitions, IR | 4.34 | [0, 12.63] |
| | | 2 partitions, AR | 4.68 | [0, 13.79] |
| 224 | (*Synthliboramphus craveri* + *S. hypoleucus*) | Unpartitioned, IR | 2.89 | [1.73, 4.63] |
| | | Unpartitioned, AR | 2.95 | [1.73, 4.60] |
| | | 2 partitions, IR | 2.36 | [1.73, 3.22] |
| | | 2 partitions, AR | 3.30 | [1.73, 5.92] |

### 5 Image Credits

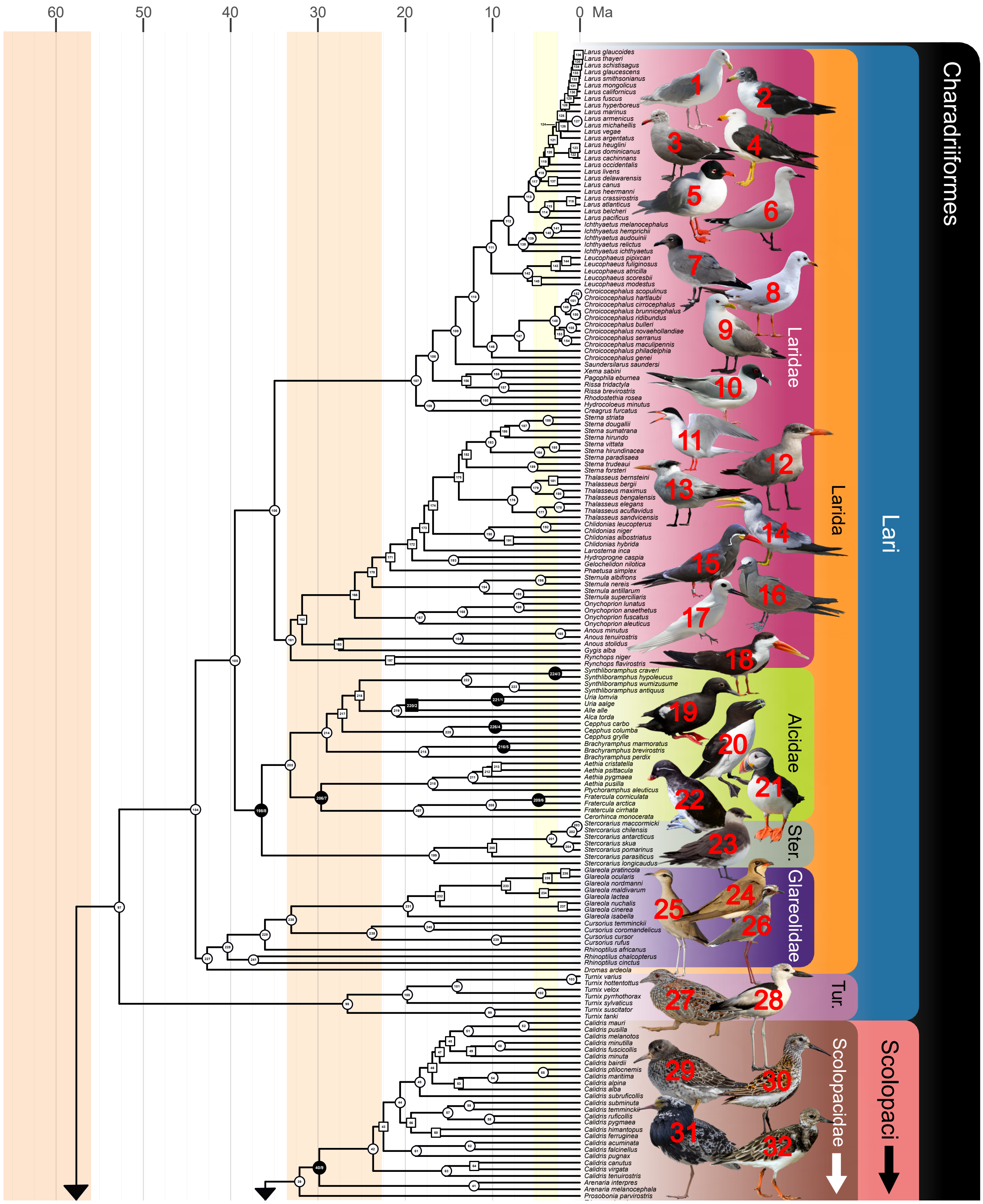

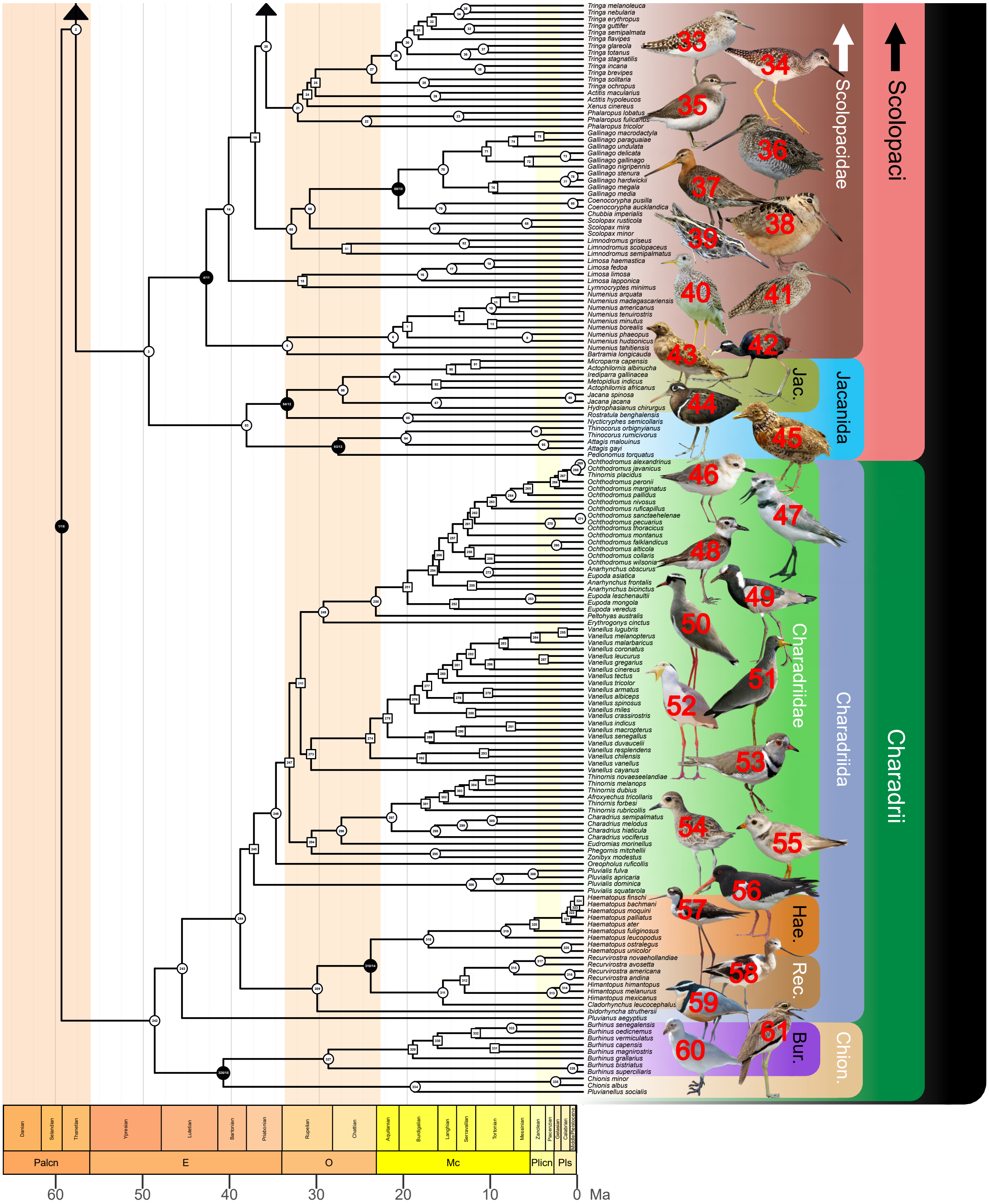

**Figure A.14:** Key to Figure 5 in the main text. Bird image numbers refer to Table A.2.

**Table A.2:** Detailed image credits and information for the bird photographs from Figure 5, numbered as in Figure A.14. The background of each photograph was removed in accordance with its license terms.

| # | Species | URL | Author | License |
|---|---|---|---|---|
| 1 | *Larus glaucescens* | `https://commons.wikimedia.org/wiki/File:Glaucous-winged_gull.jpg` | Drfred55 | CC BY-SA 3.0 |
| 2 | *Larus belcheri* | `https://commons.wikimedia.org/wiki/File:Gaviota_peruana,_Playa_La_Mina,_Paracas,_Ica,_Per%C3%BA.JPG` | Josue Hermoza | CC BY-SA 3.0 |
| 3 | *Larus heermanni* | `https://commons.wikimedia.org/wiki/File:Larus_heermanni_at_Richardson_Bay.jpg` | Frank Schulenburg | CC BY-SA 4.0 |
| 4 | *Larus pacificus* | `https://commons.wikimedia.org/wiki/File:Larus_pacificus_-_Derwent_River_Estuary.jpg` | John Harrison | CC BY-SA 3.0 |
| 5 | *Ichthyaetus melanocephalus* | `https://commons.wikimedia.org/wiki/File:Larus_melanocephalus_-Zwin_(Belgium).jpg` | Michel wal | CC BY-SA 3.0 |
| 6 | *Chroicocephalus bulleri* | `https://commons.wikimedia.org/wiki/File:Black_billed_gull_queenstown_closeup.jpg` | André Richard Chalmers | CC BY-SA 3.0 |
| 7 | *Leucophaeus fuliginosus* | `https://commons.wikimedia.org/wiki/File:Lava_Gull.jpg` | Vince Smith | CC BY 2.0 |
| 8 | *Chroicocephalus ridibundus* | `https://commons.wikimedia.org/wiki/File:Annecy%27s_Lake_-_20111229_-_Larus_ridibundus_01.JPG` | Pierre-Selim Huard | CC BY-SA 3.0 |
| 9 | *Rissa tridactyla* | `https://www.flickr.com/photos/beckymatsubara/17742276573` | Becky Matsubara | CC BY 2.0 |
| 10 | *Creagrus furcatus* | `https://commons.wikimedia.org/wiki/File:Creagrus_furcatus_-Galapagos_Islands-8.jpg` | Sue Cantan | CC BY 2.0 |
| 11 | *Sterna hirundo* | `https://commons.wikimedia.org/wiki/File:Favorite_Spot_-_panoramio.jpg` | NaturesFan1226 | CC BY 3.0 |

| | | | | |
|---|---|---|---|---|
| 12 | *Hydroprogne caspia* | https://commons.wikimedia.org/wiki/File:Caspian_tern_(Hydroprogne_caspia)_non-breeding.jpg | Charles J. Sharp | CC BY-SA 4.0 |
| 13 | *Thalasseus maximus* | https://commons.wikimedia.org/wiki/File:Sterna_maxima_portrait.jpg | Ianaré Sévi | CC BY-SA 3.0 |
| 14 | *Phaetusa simplex* | https://commons.wikimedia.org/wiki/File:Grosschnabel-Seeschwalbe.jpg | Andreas Trepte (www.avi-fauna.info) | CC BY-SA 4.0 |
| 15 | *Larosterna inca* | https://commons.wikimedia.org/wiki/File:Inca_Tern_RWD3.jpg | Dick Daniels (carolinabirds.org) | CC BY-SA 3.0 |
| 16 | *Anous stolidus* | https://commons.wikimedia.org/wiki/File:Brown_Noddy_JCB.jpg | Joseph C. Boone | CC BY-SA 4.0 |
| 17 | *Gygis alba* | https://commons.wikimedia.org/wiki/File:White_tern_with_fish.jpg | Duncan Wright | Public Domain |
| 18 | *Rynchops flavirostris* | https://www.flickr.com/photos/robertmuckley/6493424267 | Robert Muckley | CC BY 2.0 |
| 19 | *Cepphus columba* | https://www.flickr.com/photos/93649757@N07/27424034055 | Jacob McGinnis | CC BY-NC 2.0 |
| 20 | *Alca torda* | https://commons.wikimedia.org/wiki/File:Razorbill_(27971304511).jpg | Melissa McMasters | CC BY 2.0 |
| 21 | *Fratercula arctica* | https://commons.wikimedia.org/wiki/File:Papageitaucher_Fratercula_arctica.jpg | Richard Bartz | CC BY-SA 3.0 |
| 22 | *Aethia psittacula* | https://www.flickr.com/photos/dgovoni/35189010870 | Dave Govoni | CC BY-NC-SA 2.0 |
| 23 | *Stercorarius parasiticus* | https://commons.wikimedia.org/wiki/File:Arctic_skua_(Stercorarius_parasiticus)_on_an_ice_floe,_Svalbard.jpg | Andreas Weith | CC BY-SA 4.0 |
| 24 | *Glareola maldivarum* | https://commons.wikimedia.org/wiki/File:Glareola_maldivarum_-_Beung_Borapet.jpg | John Harrison | CC BY-SA 3.0 |

| | | | | |
|---|---|---|---|---|
| 25 | *Cursorius cursor* | https://commons.wikimedia.org/wiki/File:Cream-coloured_Courser_(4803935963).jpg | Mike Prince | CC BY 2.0 |
| 26 | *Rhinoptilus chalcopterus* | https://commons.wikimedia.org/wiki/File:Bronze-winged_Courser_%28Rhinoptilus_chalcopterus%29_%2813950665425%29.jpg | Bernard Dupont | CC BY-SA 2.0 |
| 27 | *Turnix varius* | https://commons.wikimedia.org/wiki/File:Turnix_varius_-_Castlereigh_nature_reserve.jpg | John Harrison | CC BY-SA 4.0 |
| 28 | *Dromas ardeola* | https://commons.wikimedia.org/wiki/File:Crab_Plover.jpg | David V. Raju | CC BY-SA 4.0 |
| 29 | *Calidris maritima* | https://commons.wikimedia.org/wiki/File:2019-08-13_01_Purple_Sandpiper_(Calidris_maritima),_Reykjavik_Iceland.jpg | Gordon Leggett | CC BY-SA 4.0 |
| 30 | *Calidris alpina* | https://commons.wikimedia.org/wiki/File:Alpenstrandl%C3%A4ufer_(calidris_alpina)_-_Spiekeroog,_Nationalpark_nieders%C3%A4chsisches_Wattenmeer.jpg | Stephan Sprinz | CC BY-SA 4.0 |
| 31 | *Calidris pugnax* | https://commons.wikimedia.org/wiki/File:Brushane%2C_Pulsuj%C3%A4rvi_sameviste%2C_%C3%96vre_Soppero%2C_Torne_lappmark%2C_May_2018_%2844519721192%29.jpg | Hans Norelius | CC BY 2.0 |
| 32 | *Arenaria interpres* | https://commons.wikimedia.org/wiki/File:Ruddy_turnstone_(Arenaria_interpres_morinella).jpg | Charles J. Sharp (Sharp Photography) | CC BY-SA 4.0 |
| 33 | *Tringa glareola* | https://commons.wikimedia.org/wiki/File:Tringa_glareola_-_Laem_Phak_Bia.jpg | John Harrison | CC BY-SA 3.0 |
| 34 | *Tringa flavipes* | https://commons.wikimedia.org/wiki/File:Lesser_Yellowlegs.jpg | Wolfgang Wander | CC BY-SA 3.0 (migrated) |

| | | | | |
|---|---|---|---|---|
| 35 | *Actitis hypoleucos* | https://commons.wikimedia.org/wiki/File:Actitis_hypoleucos_a2.jpg | Alpsdake | CC BY-SA 4.0 |
| 36 | *Gallinago delicata* | https://www.flickr.com/photos/slobirdr/11635736575 | Gregory "Slobirdr" Smith | CC BY-SA 2.0 |
| 37 | *Limosa limosa* | https://commons.wikimedia.org/wiki/File:Black-tailed_Godwit_Uferschnepfe.jpg | Andreas Trepte (www.avi-fauna.info) | CC BY-SA 2.5 |
| 38 | *Scolopax minor* | https://www.flickr.com/photos/79452129@N02/16885281018 | Fyn Kynd | CC BY 2.0 |
| 39 | *Lymnocryptes minimus* | https://commons.wikimedia.org/wiki/File:B%C3%A9cassine_sourde_au_sol.jpg | Clément Burzawa | CC BY-SA 4.0 |
| 40 | *Bartramia longicauda* | https://commons.wikimedia.org/wiki/File:UplandSandpiperOntario.jpg | Johnathan Nightingale | CC BY-SA 3.0 |
| 41 | *Numenius madagascariensis* | https://commons.wikimedia.org/wiki/File:Numenius_madagascariensis_2_-_Stockton_Sandspit.jpg | John Harrison | CC BY-SA 4.0 |
| 42 | *Metopidus indicus* | https://commons.wikimedia.org/wiki/File:Bronze-winged_jacana_(Metopidius_indicus).jpg | Charles J. Sharp (Sharp Photography) | CC BY-SA 4.0 |
| 43 | *Thinocorus orbignyianus* | https://commons.wikimedia.org/wiki/File:Thinocorus_orbignyianus_-_Swedish_Museum_of_Natural_History_-_Stockholm%2C_Sweden_-_DSC00630.JPG | Daderot | CC0 1.0 |
| 44 | *Rostratula benghalensis* | https://commons.wikimedia.org/wiki/File:African_painted_snipe_-_female_-_Kruger_National_Park_%2836204828830%29.jpg | Derek Keats | CC-BY 2.0 |
| 45 | *Pedionomus torquatus* | https://commons.wikimedia.org/wiki/File:Pedionomus_torquatus_-_Swedish_Museum_of_Natural_History_-_Stockholm%2C_Sweden_-_DSC00632.JPG | Daderot | CC0 1.0 |

| | | | | |
|---|---|---|---|---|
| 46 | *Ochthrodomus alexandrinus* | https://commons.wikimedia.org/wiki/File:Charadrius_alexandrinus_-_Laem_Pak_Bia.jpg | John Harrison | CC BY 3.0 |
| 47 | *Anarhynchus frontalis* | https://commons.wikimedia.org/wiki/File:Wrybill_Anarhynchus_frontalis,_New_Zealand.jpg | Renke Lühken | CC BY 2.0 |
| 48 | *Ochthodromus wilsonia* | https://commons.wikimedia.org/wiki/File:Wilson%27s_plover,_little_estero_(34538689103).jpg | Russ Whitehurst | CC BY 2.0 |
| 49 | *Vanellus armatus* | https://commons.wikimedia.org/wiki/File:Blacksmith_Lapwing_(Vanellus_armatus)_(16383437517).jpg | Bernard Dupont | CC BY-SA 2.0 |
| 50 | *Vanellus coronatus* | https://commons.wikimedia.org/wiki/File:Crowned_Lapwing_(Vanellus_coronatus)_(6041345466).jpg | Bernard Dupont | CC BY-SA 2.0 |
| 51 | *Vanellus senegallus* | https://commons.wikimedia.org/wiki/File:Wattled_Plover_Mara_edit3.jpg | Whit Welles | CC BY 3.0 |
| 52 | *Vanellus miles* | https://www.flickr.com/photos/larahsphotography/50324696793 | Larah McElroy | CC BY-NC 2.0 |
| 53 | *Afroxyechus tricollaris* | https://commons.wikimedia.org/wiki/File:Three-banded_plover_%28Charadrius_tricollaris%29.jpg | Charles J. Sharp (Sharp Photography) | CC BY-SA 4.0 |
| 54 | *Pluvialis fulva* | https://commons.wikimedia.org/wiki/File:Pluvialis_fulva_-_Laem_Pak_Bia.jpg | John Harrison | CC BY 3.0 |
| 55 | *Charadrius melodus* | https://commons.wikimedia.org/wiki/File:Charadrius_melodus_-Cape_May,_New_Jersey,_USA-8.jpg | Donald R. Miller | CC BY 2.0 |
| 56 | *Haematopus ostralegus* | https://commons.wikimedia.org/wiki/File:Austernfischer_Haematopus_ostralegus.jpg | Andreas Trepte (www.avi-fauna.info) | CC BY-SA 2.5 |

| | | | | |
|---|---|---|---|---|
| 57 | *Himantopus mexicanus* | https://commons.wikimedia.org/wiki/File:Black-necked_Stilt_-9_100-_%2832500015474%29.jpg | Tim Sackton | CC BY-SA 2.0 |
| 58 | *Recurvirostra americana* | https://www.flickr.com/photos/usfwsmtnprairie/42818815975 | Tom Koerner / USFWS Mountain-Prairie | CC BY 2.0 |
| 59 | *Pluvianus aegyptius* | https://commons.wikimedia.org/wiki/File:Pluvianus_aegyptius_3_Luc_Viatour.jpg | Luc Viatour (lucnix.be) | CC BY-SA 3.0 (migrated) |
| 60 | *Chionis albus* | https://www.flickr.com/photos/mfoubister/42456454022 | Murray Foubister | CC BY-SA 2.0 |
| 61 | *Burhinus vermiculatus* | https://commons.wikimedia.org/wiki/File:Water_Thick-knee_(Burhinus_vermiculatus)_(11668464924).jpg | Bernard Dupont | CC BY-SA 2.0 |

Jarvis, E. D., Mirarab, S., Aberer, A. J., Li, B., Houde, P., Li, C., Ho, S. Y. W., Faircloth, B. C., Nabholz, B., Howard, J. T., Suh, A., Weber, C. C., da Fonseca, R. R., Li, J., Zhang, F., Li, H., Zhou, L., Narula, N., Liu, L., Ganapathy, G., Boussau, B., Bayzid, M. S., Zavidovych, V., Subramanian, S., Gabaldón, T., Capella-Gutiérrez, S., Huerta-Cepas, J., Rekepalli, B., Munch, K., Schierup, M., Lindow, B., Warren, W. C., Ray, D., Green, R. E., Bruford, M. W., Zhan, X., Dixon, A., Li, S., Li, N., Huang, Y., Derryberry, E. P., Bertelsen, M. F., Sheldon, F. H., Brumfield, R. T., Mello, C. V., Lovell, P. V., Wirthlin, M., Schneider, M. P. C., Prosdocimi, F., Samaniego, J. A., Velazquez, A. M. V., Alfaro-Núñez, A., Campos, P. F., Petersen, B., Sicheritz-Ponten, T., Pas, A., Bailey, T., Scofield, P., Bunce, M., Lambert, D. M., Zhou, Q., Perelman, P., Driskell, A. C., Shapiro, B., Xiong, Z., Zeng, Y., Liu, S., Li, Z., Liu, B., Wu, K., Xiao, J., Yinqi, X., Zheng, Q., Zhang, Y., Yang, H., Wang, J., Smeds, L., Rheindt, F. E., Braun, M., Fjeldså, J., Orlando, L., Barker, F. K., Jønsson, K. A., Johnson, W., Koepfli, K.-P., O'Brien, S., Haussler, D., Ryder, O. A., Rahbek, C., Willerslev, E., Graves, G. R., Glenn, T. C., McCormack, J., Burt, D., Ellegren, H., Alström, P., Edwards, S. V., Stamatakis, A., Mindell, D. P., Cracraft, J., Braun, E. L., Warnow, T., Jun, W., Gilbert, M. T. P., and Zhang, G. (2014). Whole-genome analyses resolve early branches in the tree of life of modern birds. *Science*, 346(6215):1320–1331.

Jarzembowski, E. A., Wang, B., Zhang, H., and Fang, Y. (2015). Boring beetles are not necessarily dull: New notocupedins (Insecta: Coleoptera) from the Mesozoic of Eurasia and East Gondwana. *Cretaceous Research*, 52:431–439.

Jehl, J. R. (1968). Relationships in the Charadrii (shorebirds): A taxonomic study based on color patterns of the downy young. *San Diego Society of Natural History Memoirs*, 3:1–54.

Jerzykiewicz, T., Currie, P. J., Fanti, F., and Lefeld, J. (2021). Lithobiotopes of the Nemegt Gobi Basin. *Canadian Journal of Earth Sciences*. 10.1139/cjes-2020-0148.

Jin, F., Zhang, F., Li, Z., Zhang, J., Li, C., and Zhou, Z. (2008). On the horizon of *Protopteryx* and the early vertebrate fossil assemblages of the Jehol Biota. *Chinese Science Bulletin*, 53(18):2820–2827.

Johnson, A. L. A., Valentine, A., Leng, M. J., Sloane, H. J., Schöne, B. R., and Balson, P. S. (2017). Isotopic temperatures from the early and mid-Pliocene of the US Middle Atlantic Coastal Plain, and their implications for the cause of regional marine climate change. *Palaios*, 32(4):250–269.

Kaiser, G. and Dyke, G. J. (2011). Introduction: Changing the questions in avian paleontology. In Dyke, G. J. and Kaiser, G., editors, *Living Dinosaurs: The Evolutionary History of Modern Birds*, pages 3–8. John Wiley & Sons, Chichester, UK.

Kaufman, D. S. and Brigham-Grette, J. (1993). Aminostratigraphic correlations and paleotemperature implications, Pliocene-Pleistocene high-sea-level deposits, northwestern Alaska. *Quaternary Science Reviews*, 12(1):21–33.

Kelly, T. S. (2013). New Hemphillian (late Miocene) rodents from the Coal Valley Formation, Smith Valley, Nevada. *Paludicola*, 9(2):70–96.

Kelly, T. S. and Secord, R. (2009). Biostratigraphy of the Hunter Creek Sandstone, Verdi Basin, Washoe County, Nevada. *Geological Society of America Special Paper*, 447:133–146.

Kempf, O., Matter, A., Burbank, D. W., and Mange, M. (1999). Depositional and structural evolution of a foreland basin margin in a magnetostratigraphic framework: the eastern Swiss Molasse Basin. *International Journal of Earth Sciences*, 88(2):253–275.

Khand, Yo., Badamgarav, D., Ariunchimeg, Ya., and Barsbold, R. (2000). Cretaceous system in Mongolia and its depositional environments. In Okada, H. and Mateer, N. J., editors, *Cretaceous Environments of Asia*, pages 49–79. Elsevier, Amsterdam.

Kim, T., Lee, Y., and Lee, Y.-N. (2018). Fluorapatite diagenetic differences between Cretaceous skeletal fossils of Mongolia and Korea. *Palaeogeography, Palaeoclimatology, Palaeoecology*, 490:579–589.

Kimball, R. T., Oliveros, C. H., Wang, N., White, N. D., Barker, F. K., Field, D. J., Ksepka, D. T., Chesser, R. T., Moyle, R. G., Braun, M. J., Brumfield, R. T., Faircloth, B. C., Smith, B. T., and Braun, E. L. (2019). A phylogenomic supertree of birds. *Diversity*, 11(7):109.

Kimball, R. T., Wang, N., Heimer-McGinn, V., Ferguson, C., and Braun, E. L. (2013). Identifying localized biases in large datasets: A case study using the avian tree of life. *Molecular Phylogenetics and Evolution*, 69(3):1021–1032.

Krantz, D. E. (1991). A chronology of Pliocene sea-level fluctuations: The U.S. Middle Atlantic Coastal Plain record. *Quaternary Science Reviews*, 10(2):163–174.

Ksepka, D. T. and Clarke, J. A. (2015). Phylogenetically vetted and stratigraphically constrained fossil calibrations within Aves. *Palaeontologia Electronica*, 18.1.3FC.

Ksepka, D. T. and Phillips, M. J. (2015). Avian diversification patterns across the K-Pg boundary: influence of calibrations, datasets, and model misspecification. *Annals of the Missouri Botanical Garden*, 100(4):300–328.

Ksepka, D. T., Stidham, T. A., and Williamson, T. E. (2017). Early Paleocene landbird supports rapid phylogenetic and morphological diversification of crown birds after the K–Pg mass extinction. *Proceedings of the National Academy of Sciences*, 114(30):8047–8052.

Kuhl, H., Frankl-Vilches, C., Bakker, A., Mayr, G., Nikolaus, G., Boerno, S. T., Klages, S., Timmermann, B., and Gahr, M. (2020). An unbiased molecular approach using 3'-UTRs resolves the avian family-level tree of life. *Molecular Biology and Evolution*, 38(1):108–127.

Kundrát, M., Nudds, J., Kear, B. P., Lü, J., and Ahlberg, P. (2019). The first specimen of *Archaeopteryx* from the Upper Jurassic Mörnsheim Formation of Germany. *Historical Biology*, 31(1):3–63.

Kurochkin, E. N. (1982). Новый отряд птиц из нижнего мела Монголии [New order of birds from the Lower Cretaceous of Mongolia]. *Doklady Akademii Nauk SSSR*, 262(2):452–455. [in Russian].

Kurochkin, E. N., Dyke, G. J., and Karhu, A. A. (2002). A new presbyornithid bird (Aves, Anseriformes) from the Late Cretaceous of southern Mongolia. *American Museum Novitates*, 3386:1–11.

Kurochkin, E. N., Zelenkov, N. V., Averianov, A. O., and Leshchinskiy, S. V. (2011). A new taxon of birds (Aves) from the Early Cretaceous of Western Siberia, Russia. *Journal of Systematic Palaeontology*, 9(1):109–117.

Leanza, H. A. and Hugo, C. A. (2001). Cretaceous red beds from southern Neuquén Basin (Argentina): age, distribution and stratigraphic discontinuities. VII International Symposium on Mesozoic Terrestrial Ecosystems. *Asociación Paleontológica Argentina, Publicación Especial*, 7:117–122.

Lefèvre, U., Hu, D., Escuillié, F., Dyke, G., and Godefroit, P. (2014). A new long-tailed basal bird from the Lower Cretaceous of north-eastern China. *Biological Journal of the Linnean Society*, 113(3):790–804.

Lenz, O. K., Wilde, V., Mertz, D. F., and Riegel, W. (2015). New palynology-based astronomical and revised $^{40}$Ar/$^{39}$Ar ages for the Eocene maar lake of Messel (Germany). *International Journal of Earth Sciences*, 104(3):873–889.

Li, D., Sullivan, C., Zhou, Z., and Zhang, F. (2010). Basal birds from China: a brief review. *Chinese Birds*, 1(2):83–96.

Li, H. and Yang, J. (2004). Evidence for Cretaceous uplift of the northern Qinghai-Tibetan plateau. *Earth Science Frontiers*, 11(4):345–359. [in Chinese with English abstract].

Li, L., Wang, J.-Q., and Hou, S.-L. (2011). A new ornithurine bird (Hongshanornithidae) from the Jiufotang Formation of Chaoyang, Liaoning, China. *Vertebrata PalAsiatica*, 49(2):195–200. [in Chinese with English abstract].

Lillegraven, J. A. and McKenna, M. C. (1986). Fossil mammals from the "Mesaverde" Formation (Late Cretaceous, Judithian) of the Bighorn and Wind River Basins, Wyoming, with definitions of Late Cretaceous North American land-mammal "ages". *American Museum Novitates*, 2840:1–68.

Linnaeus, C. (1758). *Systema naturae per regna tria naturae, secundum classes, ordines, genera, species, cum characteribus, differentiis, synonimis, locis. Tomus I. Editio decima, reformata.* Laurentius Salvius, Stockholm.

Liu, D., Chiappe, L. M., Zhang, Y., Bell, A. K., Meng, Q., Ji, Q., and Wang, X. (2014). An advanced, new long-legged bird from the Early Cretaceous of the Jehol Group (northeastern China): insights into the temporal divergence of modern birds. *Zootaxa*, 3884(3):253–266.

Liu, Z., Liu, X., and Huang, S. (2017). Cyclostratigraphic analysis of magnetic records for orbital chronology of the Lower Cretaceous Xiagou Formation in Linze, northwestern China. *Palaeogeography, Palaeoclimatology, Palaeoecology*, 481:44–56.

Livezey, B. C. and Zusi, R. L. (2007). Higher-order phylogeny of modern birds (Theropoda, Aves: Neornithes) based on comparative anatomy. II. Analysis and discussion. *Zoological Journal of the Linnean Society*, 149(1):1–95.

Lofgren, D. L., Lillegraven, J. A., Clemens, W. A., Gingerich, P. D., and Williamson, T. E. (2004). Paleocene biochronology: the Puercan through Clarkforkian land mammal ages. In Woodburne, M. O., editor, *Late Cretaceous and Cenozoic Mammals of North America: Biostratigraphy and Geochronology*, pages 43–105. Columbia University Press, New York, NY.

Longrich, N. R. (2009). An ornithurine-dominated avifauna from the Belly River Group (Campanian, Upper Cretaceous) of Alberta, Canada. *Cretaceous Research*, 30(1):161–177.

Longrich, N. R., Tokaryk, T., and Field, D. J. (2011). Mass extinction of birds at the Cretaceous–Paleogene (K–Pg) boundary. *Proceedings of the National Academy of Sciences*, 108(37):15253–15257.

Lucas, S. G. and Kondrashov, P. E. (2004). Early Eocene (Bumbanian) perissodactyls from Mongolia and their biochronological significance. *New Mexico Museum of Natural History and Science Bulletin*, 26:215–220.

Machalski, M. (2018). The Cenomanian ammonite *Schloenbachia varians* (J. Sowerby, 1817) from the Cambridge Greensand of eastern England: Possible sedimentological and taphonomic implications. *Cretaceous Research*, 87:120–125.

Maisch, H. M. (2020). A new species of *Hypolophites* (Chondrichthyes, Myliobatiformes) from the Lower Clayton Limestone Unit of the Midway Group (Paleocene), near Malvern, Arkansas, USA. *Journal of Paleontology*, 94(3):548–556.

Mao, F.-Y., Li, Q., Wang, Y.-Q., and Li, C.-K. (2017). *Taizimylus tongi*, a new eurymylid (Mammalia, Glires) from the upper Paleocene of Xinjiang, China. *Palaeoworld*, 26(3):519–530.

Marivaux, L., Essid, E. M., Marzougui, W., Khayati Ammar, H., Adnet, S., Marandat, B., Merzeraud, G., Tabuce, R., and Vianey-Liaud, M. (2014). A new and primitive species of *Protophiomys* (Rodentia, Hystricognathi) from the late middle Eocene of Djebel el Kébar, Central Tunisia. *Palaeovertebrata*, 38(1):e2.

Marjanović, D. (2021). The making of calibration sausage exemplified by recalibrating the transcriptomic timetree of jawed vertebrates. *Frontiers in Genetics*, 12:535.

Marquillas, R. A., Salfity, J. A., Matthews, S. J., Matteini, M., and Dantas, E. (2011). U-Pb zircon age of the Yacoraite Formation and its significance to the Cretaceous-Tertiary boundary in the Salta basin, Argentina. In Salfity, J. A. and Marquillas, R. A., editors, *Cenozoic Geology of the Central Andes of Argentina*, pages 227–246. SCS Publisher, Salta.

Marsh, O. C. (1872). Notice of a new and remarkable fossil bird. *American Journal of Science, Series 3*, 4(22):344.

Marsh, O. C. (1892). Notes on Mesozoic vertebrate fossils. *American Journal of Science, Series 3*, 44(260):171–176.

Marx, F. G. and Fordyce, R. E. (2015). Baleen boom and bust: a synthesis of mysticete phylogeny, diversity and disparity. *Royal Society Open Science*, 2(4):140434.

Maxwell, E. E., Alexander, S., Bechly, G., Eck, K., Frey, E., Grimm, K., Kovar-Eder, J., Mayr, G., Micklich, N., Rasser, M., Roth-Nebelsick, A., Salvador, R. B., Schoch, R. R., Schweigert, G., Stinnesbeck, W., Wolf-Schwenninger, K., and Ziegler, R. (2016). The Rauenberg fossil Lagerstätte (Baden-Württemberg, Germany): A window into early Oligocene marine and coastal ecosystems of Central Europe. *Palaeogeography, Palaeoclimatology, Palaeoecology*, 463:238–260.

Mayr, G. (2009). *Paleogene Fossil Birds*. Springer-Verlag, Berlin.

Mayr, G. (2011a). Cenozoic mystery birds – on the phylogenetic affinities of bony-toothed birds (Pelagornithidae). *Zoologica Scripta*, 40(5):448–467.

Mayr, G. (2011b). The phylogeny of charadriiform birds (shorebirds and allies) – reassessing the conflict between morphology and molecules. *Zoological Journal of the Linnean Society*, 161(4):916–934.

Mayr, G. (2013). The age of the crown group of passerine birds and its evolutionary significance – molecular calibrations versus the fossil record. *Systematics and Biodiversity*, 11(1):7–13.

Mayr, G. (2014). The origins of crown group birds: molecules and fossils. *Palaeontology*, 57(2):231–242.

Mayr, G. (2015). A new Paleogene procellariiform bird from western North America. *Neues Jahrbuch für Geologie und Paläontologie - Abhandlungen*, 275(1):11–17.

Mayr, G. (2017a). *Avian Evolution: The Fossil Record of Birds and its Paleobiological Significance*. John Wiley & Sons, Chichester, UK.

Mayr, G. (2017b). A small, “wader-like” bird from the Early Eocene of Messel (Germany). *Annales de Paléontologie*, 103(2):141–147.

Mayr, G. (2018). New data on the anatomy and palaeobiology of sandcoleid mousebirds (Aves, Coliiformes) from the early Eocene of Messel. *Palaeobiodiversity and Palaeoenvironments*, 98(4):639–651.

Mayr, G. and Clarke, J. A. (2003). The deep divergences of neornithine birds: a phylogenetic analysis of morphological characters. *Cladistics*, 19(6):527–553.

Mayr, G., De Pietri, V. L., Love, L., Mannering, A. A., and Scofield, R. P. (2017a). A well-preserved new mid-Paleocene penguin (Aves, Sphenisciformes) from the Waipara Greensand in New Zealand. *Journal of Vertebrate Paleontology*, 37(6):e1398169.

Mayr, G., De Pietri, V. L., Love, L., Mannering, A. A., and Scofield, R. P. (2021). Oldest, smallest and phylogenetically most basal pelagornithid, from the early Paleocene of New Zealand, sheds light on the evolutionary history of the largest flying birds. *Papers in Palaeontology*, 7(1):217–233.

Mayr, G., De Pietri, V. L., and Scofield, R. P. (2017b). A new fossil from the mid-Paleocene of New Zealand reveals an unexpected diversity of world's oldest penguins. *Science of Nature*, 104(3):9.

Mayr, G., De Pietri, V. L., Scofield, R. P., and Smith, T. (2019a). A fossil heron from the early Oligocene of Belgium: the earliest temporally well-constrained record of the Ardeidae. *Ibis*, 161(1):79–90.

Mayr, G., De Pietri, V. L., Scofield, R. P., and Worthy, T. H. (2018). On the taxonomic composition and phylogenetic affinities of the recently proposed clade Vegaviidae Agnolín et al., 2017 – neornithine birds from the Upper Cretaceous of the Southern Hemisphere. *Cretaceous Research*, 86:178–185.

Mayr, G., Hervet, S., and Buffetaut, E. (2019b). On the diverse and widely ignored Paleocene avifauna of Menat (Puy-de-Dôme, France): new taxonomic records and unusual soft tissue preservation. *Geological Magazine*, 156(3):572–584.

Mayr, G. and Knopf, C. W. (2007). A stem lineage representative of buttonquails from the Lower Oligocene of Germany – fossil evidence for a charadriiform origin of the Turnicidae. *Ibis*, 149(4):774–782.

Mayr, G. and Scofield, R. P. (2014). First diagnosable non-sphenisciform bird from the early Paleocene of New Zealand. *Journal of the Royal Society of New Zealand*, 44(1):48–56.

Mayr, G. and Scofield, R. P. (2016). New avian remains from the Paleocene of New Zealand: the first early Cenozoic Phaethontiformes (tropicbirds) from the Southern Hemisphere. *Journal of Vertebrate Paleontology*, 36(1):e1031343.

Mayr, G. and Smith, R. (2001). Ducks, rails, and limicoline waders (Aves: Anseriformes, Gruiformes, Charadriiformes) from the lowermost Oligocene of Belgium. *Geobios*, 34(5):547–561.

Mayr, G., Yang, J., De Bast, E., Li, C.-S., and Smith, T. (2013). A *Strigogyps*-like bird from the middle Paleocene of China with an unusual grasping foot. *Journal of Vertebrate Paleontology*, 33(4):895–901.

McCarthy, F. M., Katz, M. E., Kotthoff, U., Browning, J. V., Miller, K. G., Zanatta, R., Williams, R. H., Drljepan, M., Hesselbo, S. P., Bjerrum, C. J., and Mountain, G. S. (2013). Sea-level control of New Jersey margin architecture: Palynological evidence from Integrated Ocean Drilling Program Expedition 313. *Geosphere*, 9(6):1457–1487.

McIver, E. E. (2002). The paleoenvironment of *Tyrannosaurus rex* from southwestern Saskatchewan, Canada. *Canadian Journal of Earth Sciences*, 39(2):207–221.

McLachlan, S. M. S. and Haggart, J. W. (2018). Reassessment of the late Campanian (Late Cretaceous) heteromorph ammonite fauna from Hornby Island, British Columbia, with implications for the taxonomy of the Diplomoceratidae and Nostoceratidae. *Journal of Systematic Palaeontology*, 16(15):1247–1299.

McLachlan, S. M. S., Kaiser, G. W., and Longrich, N. R. (2017). *Maaqwi cascadensis*: A large, marine diving bird (Avialae: Ornithurae) from the Upper Cretaceous of British Columbia, Canada. *PLOS ONE*, 12(12):e0189473.

Megirian, D., Prideaux, G. J., Murray, P. F., and Smit, N. (2010). An Australian land mammal age biochronological scheme. *Paleobiology*, 36(4):658–671.

Meng, F., Gao, S., and Liu, X. (2008). U-Pb zircon geochronology and geochemistry of volcanic rocks of the Yixian Formation in the Lingyuan area, western Liaoning, China. *Geological Bulletin of China*, 27(3):364–373. [in Chinese with English abstract].

Micklich, N., Gregorová, R., Bannikov, A. F., Baciu, D.-S., Grădianu, I., and Carnevale, G. (2016). *Oligoremora rhenana* n. g. n. sp., a new echeneid fish (Percomorpha, Echeneoidei) from the Oligocene of the Grube Unterfeld ("Frauenweiler") clay pit. *PalZ*, 90(3):561–592.

Milne-Edwards, A. (1867–1868). *Recherches anatomiques et paléontologiques pour servir à l'histoire des oiseaux fossiles de la France, Vol. I.* Victor Masson et fils, Paris.

Missiaen, P. (2011). An updated mammalian biochronology and biogeography for the early Paleogene of Asia. *Vertebrata PalAsiatica*, 49(1):29–52.

Mitchell, K. J., Cooper, A., and Phillips, M. J. (2015). Comment on "Whole-genome analyses resolve early branches in the tree of life of modern birds". *Science*, 349(6255):1460–1460.

Moncunill-Solé, B., Isidro, A., Blanco, A., Angelone, C., Rössner, G. E., and Jordana, X. (2019). The most ancient evidence of a diseased lagomorph: Infectious paleopathology in a tibiofibular bone (Middle Miocene, Germany). *Comptes Rendus Palevol*, 18(8):1011–1023.

Morales, J. and Pickford, M. (2018). New *Namalestes* remains from the Ypresian/Lutetian of Black Crow, Namibia. *Communications of the Geological Survey of Namibia*, 18:72–80.

Mourer-Chauviré, C. (2000). A new species of *Ameripodius* (Aves: Galliformes: Quercymegapodiidae) from the Lower Miocene of France. *Palaeontology*, 43(3):481–493.

Mourer-Chauviré, C., Berthet, D., and Hugueney, M. (2004). The late Oligocene birds of the Créchy quarry (Allier, France), with a description of two new genera (Aves: Pelecaniformes: Phalacrocoracidae, and Anseriformes: Anseranatidae). *Senckenbergiana Lethaea*, 84(1):303–315.

Mourer-Chauviré, C., Pickford, M., and Senut, B. (2017). New data on stem group Galliformes, Charadriiformes, and Psittaciformes from the middle Eocene of Namibia. *Contribuciones Científicas del Museo Argentino de Ciencias Naturales "Bernardino Rivadavia"*, 7:99–131.

Musser, G. and Clarke, J. A. (2020). An exceptionally preserved specimen from the Green River Formation elucidates complex phenotypic evolution in Gruiformes and Charadriiformes. *Frontiers in Ecology and Evolution*, 8:559929.

Musser, G., Ksepka, D. T., and Field, D. J. (2019). New material of Paleocene-Eocene *Pellornis* (Aves: Gruiformes) clarifies the pattern and timing of the extant gruiform radiation. *Diversity*, 11(7):102.

Myers, T. S. (2010). Earliest occurrence of the Pteranodontidae (Archosauria: Pterosauria) in North America: new material from the Austin Group of Texas. *Journal of Paleontology*, 84(6):1071–1081.

Nakajima, J., Kobayashi, Y., Chinzorig, T., Tanaka, T., Takasaki, R., Tsogtbaatar, K., Currie, P. J., and Fiorillo, A. R. (2018). Dinosaur tracks at the Nemegt locality: Paleobiological and paleoenvironmental implications. *Palaeogeography, Palaeoclimatology, Palaeoecology*, 494:147–159.

Navalón, G., Meng, Q., Marugán-Lobón, J., Zhang, Y., Wang, B., Xing, H., Liu, D., and Chiappe, L. M. (2018). Diversity and evolution of the Confuciusornithidae: Evidence from a new 131-million-year-old specimen from the Huajiying Formation in NE China. *Journal of Asian Earth Sciences*, 152:12–22.

Nesbitt, S. J. and Clarke, J. A. (2016). The anatomy and taxonomy of the exquisitely preserved Green River Formation (early Eocene) lithornithids (Aves) and the relationships of Lithornithidae. *Bulletin of the American Museum of Natural History*, 406:1–91.

Norell, M. A. and Clarke, J. A. (2001). Fossil that fills a critical gap in avian evolution. *Nature*, 409(6817):181–184.

Obradovich, J. D. (1993). A Cretaceous time scale. In Caldwell, W. G. E. and Kauffman, E. G., editors, *Evolution of the Western Interior Basin. Geological Association of Canada Special Paper No. 39*, pages 379–396. Geological Association of Canada, St. John's, NL.

O'Connor, J. K., Chiappe, L. M., and Bell, A. K. (2011). Pre-modern birds: Avian divergences in the Mesozoic. In Dyke, G. J. and Kaiser, G., editors, *Living Dinosaurs: The Evolutionary History of Modern Birds*, pages 39–114. John Wiley & Sons, Chichester, UK.

O'Connor, J. K., Gao, K.-Q., and Chiappe, L. M. (2010). A new ornithuromorph (Aves: Ornithothoraces) bird from the Jehol Group indicative of higher-level diversity. *Journal of Vertebrate Paleontology*, 30(2):311–321.

O'Connor, J. K., Sun, C., Xu, X., Wang, X., and Zhou, Z. (2012). A new species of *Jeholornis* with complete caudal integument. *Historical Biology*, 24(1):29–41.

O'Connor, J. K., Wang, M., and Hu, H. (2016). A new ornithuromorph (Aves) with an elongate rostrum from the Jehol Biota, and the early evolution of rostralization in birds. *Journal of Systematic Palaeontology*, 14(11):939–948.

O'Connor, J. K. and Zelenkov, N. V. (2013). The phylogenetic position of *Ambiortus*: Comparison with other Mesozoic birds from Asia. *Paleontological Journal*, 47(11):1270–1281.

O'Connor, J. K. and Zhou, Z. (2013). A redescription of *Chaoyangia beishanensis* (Aves) and a comprehensive phylogeny of Mesozoic birds. *Journal of Systematic Palaeontology*, 11(7):889–906.

O'Connor, P. M. and Forster, C. A. (2010). A Late Cretaceous (Maastrichtian) avifauna from the Maevarano Formation, Madagascar. *Journal of Vertebrate Paleontology*, 30(4):1178–1201.

Ogg, J. G. (2012). Geomagnetic polarity time scale. In Gradstein, F. M., Ogg, J. G., Schmitz, M. D., and Ogg, G. M., editors, *The Geologic Time Scale 2012*, pages 85–113. Elsevier, Boston, MA.

Ogg, J. G., Agterberg, F. P., and Gradstein, F. M. (2004). The Cretaceous Period. In Gradstein, F. M., Ogg, J. G., and Smith, A. G., editors, *A Geologic Time Scale 2004*, pages 344–383. Cambridge University Press, Cambridge, UK.

Ogg, J. G., Hinnov, L. A., and Huang, C. (2012). Cretaceous. In Gradstein, F. M., Ogg, J. G., Schmitz, M. D., and Ogg, G. M., editors, *The Geologic Time Scale 2012*, pages 793–853. Elsevier, Boston, MA.

Ohno, M., Hayashi, T., Komatsu, F., Murakami, F., Zhao, M., Guyodo, Y., Acton, G., Evans, H. F., and Kanamatsu, T. (2012). A detailed paleomagnetic record between 2.1 and 2.75 Ma at IODP Site U1314 in the North Atlantic: Geomagnetic excursions and the Gauss-Matuyama transition. *Geochemistry, Geophysics, Geosystems*, 13(5):Q12Z39.

Okada, H. O. and Bukry, D. (1980). Supplementary modification and introduction of code numbers to the low-latitude coccolith biostratigraphic zonation (Bukry, 1973; 1975). *Marine Micropaleontology*, 5:321–325.

Olson, S. L. (2013). A Late Pliocene occurrence of the Thick-billed Murre (Alcidae: *Uria lomvia*) on St. George Island, Pribilofs, Alaska. *Paleontological Journal*, 47(11):1365–1368.

Olson, S. L. and Parris, D. C. (1987). The Cretaceous birds of New Jersey. *Smithsonian Contributions to Paleobiology*, 63:1–22.

Olson, S. L. and Rasmussen, P. C. (2001). Miocene and Pliocene birds from the Lee Creek Mine, North Carolina. In Ray, C. E. and Bohaska, D. J., editors, *Geology and Paleontology of the Lee Creek Mine, North Carolina, III*, pages 233–365. Smithsonian Contributions to Paleobiology 90.

Pacheco, M. A., Battistuzzi, F. U., Lentino, M., Aguilar, R. F., Kumar, S., and Escalante, A. A. (2011). Evolution of modern birds revealed by mitogenomics: Timing the radiation and origin of major orders. *Molecular Biology and Evolution*, 28(6):1927–1942.

Parham, J. F., Donoghue, P. C. J., Bell, C. J., Calway, T. D., Head, J. J., Holroyd, P. A., Inoue, J. G., Irmis, R. B., Joyce, W. G., Ksepka, D. T., Patané, J. S. L., Smith, N. D., Tarver, J. E., van Tuinen, M., Yang, Z., Angielczyk, K. D., Greenwood, J. M., Hipsley, C. A., Jacobs, L., Makovicky, P. J., Müller, J., Smith, K. T., Theodor, J. M., Warnock, R. C. M., and Benton, M. J. (2011). Best practices for justifying fossil calibrations. *Systematic Biology*, 61(2):346–359.

Parmley, D. and Alan, H. J. (2003). *Nebraskophis* Holman from the Late Eocene of Georgia (USA), the oldest known North American colubrid snake. *Acta Zoologica Cracoviensia*, 46(1):1–8.

Parmley, D., Hutchison, J. H., and Parham, J. F. (2006). Diverse turtle fauna from the Late Eocene of Georgia including the oldest records of aquatic testudinoids in southeastern North America. *Journal of Herpetology*, 40(3):343–350.

Parris, D. C. and Hope, S. (2002). New interpretations of the birds from the Navesink and Hornerstown Formations, New Jersey, USA (Aves: Neornithes). In Zhou, Z.-H. and Fu-Cheng, Z., editors, *Proceedings of the 5th International Symposium of the Society of Avian Paleontology and Evolution*, pages 113–124. Science Press, Beijing.

Perez, V. J., Godfrey, S. J. G., Kent, B. W., Weems, R. E., and Nance, J. R. (2018). The transition between *Carcharocles chubutensis* and *Carcharocles megalodon* (Otodontidae, Chondrichthyes): lateral cusplet loss through time. *Journal of Vertebrate Paleontology*, 38(6):e1546732.

Pickford, M. (2015). Cenozoic geology of the Northern Sperrgebiet, Namibia, accenting the Palaeogene. *Communications of the Geological Survey of Namibia*, 16:10–104.

Pickford, M. (2018). Tufamyidae, a new family of hystricognath rodents from the Palaeogene and Neogene of the Sperrgebiet, Namibia. *Communications of the Geological Survey of Namibia*, 19:71–109.

Pickford, M., Senut, B., Mocke, H., Mourer-Chauviré, C., Rage, J.-C., and Mein, P. (2014). Eocene aridity in southwestern Africa: timing of onset and biological consequences. *Transactions of the Royal Society of South Africa*, 69(3):139–144.

Powars, D. S. and Scott, B. T. (1999). The effects of the Chesapeake Bay impact crater on the geological framework and correlation of hydrogeologic units of the lower York-James Peninsula, Virginia. *U.S. Geological Survey Professional Paper*, 1612:1–82.

Prothero, D. R. and Emry, R. J. (2004). The Chadronian, Orellan, and Whitneyan North American land mammal ages. In Woodburne, M. O., editor, *Late Cretaceous and Cenozoic Mammals of North America: Biostratigraphy and Geochronology*, pages 157–168. Columbia University Press, New York, NY.

Prum, R. O., Berv, J. S., Dornburg, A., Field, D. J., Townsend, J. P., Lemmon, E. M., and Lemmon, A. R. (2015). A comprehensive phylogeny of birds (Aves) using targeted next-generation DNA sequencing. *Nature*, 526(7574):569–573.

Racicot, R. A., Deméré, T. A., Beatty, B. L., and Boessenecker, R. W. (2014). Unique feeding morphology in a new prognathous extinct porpoise from the Pliocene of California. *Current Biology*, 24(7):774–779.

Radmacher, W., Pérez-Rodríguez, I., Arz, J. A., and Pearce, M. A. (2014). Dinoflagellate biostratigraphy at the Campanian-Maastrichtian boundary in Zumaia, northern Spain. *Cretaceous Research*, 51:309–320.

Rasmussen, D. T., Olson, S. L., and Simons, E. L. (1987). Fossil birds from the Oligocene Jebel Qatrani Formation, Fayum Province, Egypt. *Smithsonian Contributions to Paleobiology*, 62:1–20.

Rauhut, O. W. M., Foth, C., and Tischlinger, H. (2018). The oldest *Archaeopteryx* (Theropoda: Avialiae [*sic*]): a new specimen from the Kimmeridgian/Tithonian boundary of Schamhaupten, Bavaria. *PeerJ*, 6:e4191.

Reddy, S., Kimball, R. T., Pandey, A., Hosner, P. A., Braun, M. J., Hackett, S. J., Han, K.-L., Harshman, J., Huddleston, C. J., Kingston, S., Marks, B. D., Miglia, K. J., Moore, W. S., Sheldon, F. H., Witt, C. C., Yuri, T., and Braun, E. L. (2017). Why do phylogenomic data sets yield conflicting trees? Data type influences the avian tree of life more than taxon sampling. *Systematic Biology*, 66(5):857–879.

Renne, P. R., Deino, A. L., Hilgen, F. J., Kuiper, K. F., Mark, D. F., Mitchell, W. S., Morgan, L. E., Mundil, R., and Smit, J. (2013). Time scales of critical events around the Cretaceous-Paleogene boundary. *Science*, 339(6120):684–687.

Riamon, S., Tourment, N., and Louchart, A. (2020). The earliest Tyrannida (Aves, Passeriformes), from the Oligocene of France. *Scientific Reports*, 10(1):9776.

Roberts, E. M., Lamanna, M. C., Clarke, J. A., Meng, J., Gorscak, E., Sertich, J. J. W., O'Connor, P. M., Claeson, K. M., and MacPhee, R. D. E. (2014). Stratigraphy and vertebrate paleoecology of Upper Cretaceous–?lowest Paleogene strata on Vega Island, Antarctica. *Palaeogeography, Palaeoclimatology, Palaeoecology*, 402:55–72.

Rocholl, A., Böhme, M., Gilg, H. A., Pohl, J., Schaltegger, U., and Wijbrans, J. (2018). Comment on "A high-precision $^{40}$Ar/$^{39}$Ar age for the Nördlinger Ries impact crater, Germany, and implications for the accurate dating of terrestrial impact events" by Schmieder et al. (Geochimica et Cosmochimica Acta 220 (2018) 146–157). *Geochimica et Cosmochimica Acta*, 238:599–601.

Roux, T. (2002). Deux fossiles d'oiseaux de l'Oligocène inférieur du Luberon. *Courrier scientifique du Parc naturel régional du Luberon*, 6:38–57.

Ruiz-Sánchez, F. J., Murelaga, X., Larrasoaña, J. C., Freudenthal, M., and Garcés, M. (2012). Hypsodont Myomiminae (Gliridae, Rodentia) from five new localities in the Lower Miocene Tudela Formation (Bardenas Reales, Ebro Basin, Spain) and their bearing on the age of the Agenian-Ramblian boundary. *Geodiversitas*, 34(3):645–663.

Sageman, B. B., Singer, B. S., Meyers, S. R., Siewert, S. E., Walaszczyk, I., Condon, D. J., Jicha, B. R., Obradovich, J. D., and Sawyer, D. A. (2014). Integrating 40Ar/39Ar, U-Pb, and astronomical clocks in the Cretaceous Niobrara Formation, Western Interior Basin, USA. *Geological Society of America Bulletin*, 126(7–8):956–973.

Sallam, H. M. and Seiffert, E. R. (2016). New phiomorph rodents from the latest Eocene of Egypt, and the impact of Bayesian "clock"-based phylogenetic methods on estimates of basal hystricognath relationships and biochronology. *PeerJ*, 4:e1717.

Sallam, H. M. and Seiffert, E. R. (2019). Revision of Oligocene '*Paraphiomys*' and an origin for crown Thryonomyoidea (Rodentia: Hystricognathi: Phiomorpha) near the Oligocene–Miocene boundary in Africa. *Zoological Journal of the Linnean Society*, 190(1):352–371.

Schmieder, M., Kennedy, T., Jourdan, F., Buchner, E., and Reimold, W. U. (2018a). A high-precision $^{40}$Ar/$^{39}$Ar age for the Nördlinger Ries impact crater, Germany, and implications for the accurate dating of terrestrial impact events. *Geochimica et Cosmochimica Acta*, 220:146–157.

Schmieder, M., Kennedy, T., Jourdan, F., Buchner, E., and Reimold, W. U. (2018b). Response to comment on "A high-precision $^{40}$Ar/$^{39}$Ar age for the Nördlinger Ries impact crater, Germany, and implications for the accurate dating of terrestrial impact events" by Schmieder et al. (Geochimica et Cosmochimica Acta 220 (2018) 146–157). *Geochimica et Cosmochimica Acta*, 238:602–605.

Schweigert, G. (2007). Ammonite biostratigraphy as a tool for dating Upper Jurassic lithographic limestones from South Germany – first results and open questions. *Neues Jahrbuch für Geologie und Paläontologie - Abhandlungen*, 245(1):117–125.

Secord, R. (2008). The Tiffanian land-mammal age (middle and late Paleocene) in the northern Bighorn Basin, Wyoming. *University of Michigan Papers on Paleontology*, 35:1–192.

Secord, R., Gingerich, P. D., Smith, M. E., Clyde, W. C., Wilf, P., and Singer, B. S. (2006). Geochronology and mammalian biostratigraphy of middle and upper Paleocene continental strata, Bighorn Basin, Wyoming. *American Journal of Science*, 306(4):211–245.

Seeley, H. G. (1876). On the British fossil Cretaceous birds. *Quarterly Journal of the Geological Society*, 32(1-4):496–512.

Seiffert, E. R. (2006). Revised age estimates for the later Paleogene mammal faunas of Egypt and Oman. *Proceedings of the National Academy of Sciences*, 103(13):5000–5005.

Sen, S. (1997). Magnetostratigraphic calibration of the European Neogene mammal chronology. *Palaeogeography, Palaeoclimatology, Palaeoecology*, 133(3):181–204.

Shao, S., Li, L., Yang, Y., and Zhou, C.-F. (2018). Hyperphalangy in a new sinemydid turtle from the Early Cretaceous Jehol Biota. *PeerJ*, 6:e5371.

Sibley, C. G. and Ahlquist, J. E. (1990). *Phylogeny and Classification of Birds: A Study in Molecular Evolution.* Yale University Press, New Haven, CT.

Singer, B. S. (2014). A Quaternary geomagnetic instability time scale. *Quaternary Geochronology*, 21:29–52.

Slack, K. E., Jones, C. M., Ando, T., Harrison, G. L. A., Fordyce, R. E., Arnason, U., and Penny, D. (2006). Early penguin fossils, plus mitochondrial genomes, calibrate avian evolution. *Molecular Biology and Evolution*, 23(6):1144–1155.

Smith, N. A. (2011a). *Systematics and evolution of extinct and extant Pan-Alcidae (Aves, Charadriiformes): combined phylogenetic analyses, divergence estimation, and paleoclimatic interactions.* PhD thesis, University of Texas at Austin.

Smith, N. A. (2011b). Taxonomic revision and phylogenetic analysis of the flightless Mancallinae (Aves, Pan-Alcidae). *ZooKeys*, 91:1–116.

Smith, N. A. (2014). The fossil record and phylogeny of the auklets (Pan-Alcidae, Aethiini). *Journal of Systematic Palaeontology*, 12(2):217–236.

Smith, N. A. (2015). Sixteen vetted fossil calibrations for divergence dating of Charadriiformes (Aves, Neognathae). *Palaeontologia Electronica*, 18.1.4FC.

Smith, N. A. and Clarke, J. A. (2015). Systematics and evolution of the Pan-Alcidae (Aves, Charadriiformes). *Journal of Avian Biology*, 46(2):125–140.

Smith, N. D. and Ksepka, D. T. (2015). Five well-supported fossil calibrations within the "Waterbird" assemblage (Tetrapoda, Aves). *Palaeontologia Electronica*, 18.1.7FC.

Smith, P. E., Evensen, N. M., York, D., Chang, M.-M., Jin, F., Li, J.-L., Cumbaa, S., and Russell, D. (1995). Dates and rates in ancient lakes: $^{40}$Ar–$^{39}$Ar evidence for an Early Cretaceous age for the Jehol Group, northeast China. *Canadian Journal of Earth Sciences*, 32(9):1426–1431.

Sprain, C. J., Renne, P. R., Clemens, W. A., and Wilson, G. P. (2018). Calibration of chron C29r: New high-precision geochronologic and paleomagnetic constraints from the Hell Creek region, Montana. *Geological Society of America Bulletin*, 130(9–10):1615–1644.

Steininger, F. F. (1999). Chronostratigraphy, geochronology and biochronology of the Miocene "European Land Mammal Mega-Zones (ELMMZ)" and the Miocene "Mammal-Zones (MN-Zones)". In Rössner, G. E. and Heissig, K., editors, *The Miocene Land Mammals of Europe*, pages 9–24. Verlag Dr. Friedrich Pfeil, Munich.

Steininger, F. F., Bernor, R. L., and Fahlbusch, V. (1989). European Neogene marine/continental chronologic correlations. In Lindsay, E. H., Fahlbusch, V., and Mein, P., editors, *European Neogene Mammal Chronology*, pages 15–46. Springer, Boston, MA.

Stervander, M., Ryan, P. G., Melo, M., and Hansson, B. (2019). The origin of the world's smallest flightless bird, the Inaccessible Island Rail *Atlantisia rogersi* (Aves: Rallidae). *Molecular Phylogenetics and Evolution*, 130:92–98.

Stidham, T. A., Lofgren, D. L., Farke, A. A., Paik, M., and Choi, R. (2014). A lithornithid (Aves: Palaeognathae) from the Paleocene (Tiffanian) of southern California. *PaleoBios*, 31(1):1–7.

Stidham, T. A. and Smith, N. A. (2015). An ameghinornithid-like bird (Aves, Cariamae, ?Ameghinornithidae) from the early Oligocene of Egypt. *Palaeontologia Electronica*, 18.1.5A.

Strauch, J. G. (1978). The phylogeny of the Charadriiformes (Aves): a new estimate using the method of character compatibility analysis. *Transactions of the Zoological Society of London*, 34(3):263–345.

Suh, A. (2016). The phylogenomic forest of bird trees contains a hard polytomy at the root of Neoaves. *Zoologica Scripta*, 45(S1):50–62.

Sun, G., Dilcher, D. L., Wang, H., and Chen, Z. (2011). A eudicot from the Early Cretaceous of China. *Nature*, 471(7340):625–628.

Tambussi, C. P., Degrange, F. J., De Mendoza, R. S., Sferco, E., and Santillana, S. (2019). A stem anseriform from the early Palaeocene of Antarctica provides new key evidence in the early evolution of waterfowl. *Zoological Journal of the Linnean Society*, 186(3):673–700.

Tedford, R. H., Albright III, L. B., Barnosky, A. D., Ferrusquia-Villafranca, I., Hunt Jr., R. M., Storer, J. E., Swisher III, C. C., Voorhies, M. R., Webb, S. D., and Whistler, D. P. (2004). Mammalian biochronology of the Arikareean through Hemphillian interval (Late Oligocene through Early Pliocene epochs). In Woodburne, M. O., editor, *Late Cretaceous and Cenozoic Mammals of North America: Biostratigraphy and Geochronology*, pages 169–231. Columbia University Press, New York, NY.

Thorn, K. M., Hutchinson, M. N., Lee, M. S. Y., Brown, N. J., Camens, A. B., and Worthy, T. H. (2021). A new species of *Proegernia* from the Namba Formation in South Australia and the early evolution and environment of Australian egerniine skinks. *Royal Society Open Science*, 8(2):201686.

Ting, S., Vowen, G. J., Koch, P. L., Clyde, W. C., Wang, Y., Wang, Y., and McKenna, M. C. (2003). Biostratigraphic, chemostratigraphic, and magnetostratigraphic study across the Paleocene-Eocene boundary in the Hengyang Basin, Hunan, China. In *Causes and consequences of globally warm climates in the early Paleogene: Geological Society of America Special Paper 369*, pages 521–535. Geological Society of America, Boulder, CO.

Titov, V. V. and Tesakov, A. S. (2013). Late Miocene (Turolian) vertebrate faunas from southern European Russia. In Wang, X., Flynn, L. J., and Fortelius, M., editors, *Fossil Mammals of Asia: Neogene Biostratigraphy and Chronology*, pages 538–545. Columbia University Press, New York, NY.

Tokaryk, T. T., L., C. S., and Storer, J. E. (1997). Early Late Cretaceous birds from Saskatchewan, Canada: the oldest diverse avifauna known from North America. *Journal of Vertebrate Paleontology*, 17(1):172–176.

Underwood, C. J., King, C., and Steurbaut, E. (2013). Eocene initiation of Nile drainage due to East African uplift. *Palaeogeography, Palaeoclimatology, Palaeoecology*, 392:138 – 145.

Vandenberghe, N., Hilgen, F. J., Speijer, R. P., Ogg, J. G., Gradstein, F. M., Hammer, Ø., Hollis, C. J., and Hooker, J. J. (2012). The Paleogene Period. In Gradstein, F. M., Ogg, J. G., Schmitz, M. D., and Ogg, G. M., editors, *The Geologic Time Scale 2012*, pages 855–921. Elsevier, Boston, MA.

Velez-Juarbe, J. (2017). *Eotaria citrica*, sp. nov., a new stem otariid from the "Topanga" formation of Southern California. *PeerJ*, 5:e3022.

Vendrasco, M. J., Eernisse, D. J., Powell, C. L. I., and Fernandez, C. Z. (2012). Polyplacophora (Mollusca) from the San Diego Formation: A remarkable assemblage of fossil chitons from the Pliocene of Southern California. *Contributions in Science*, 520:15–72.

Wagner, H. M., Riney, B. O., Deméré, T. A., and Prothero, D. R. (2001). Magnetic stratigraphy and land mammal biochronology of the nonmarine facies of the Pliocene San Diego Formation, San Diego County, California. In Prothero, D. R., editor, *Magnetic Stratigraphy of the Pacific Coast Cenozoic*, pages 359–368. Society of Economic Paleontologists and Mineralogists, Pacific Section, Special Publication 91, Los Angeles, CA.

Walker, C. A. and Dyke, G. J. (2009). Euenantiornithine birds from the Late Cretaceous of El Brete (Argentina). *Irish Journal of Earth Sciences*, 27:15–62.

Wang, M., Li, Z., Liu, Q., and Zhou, Z. (2020a). Two new Early Cretaceous ornithuromorph birds provide insights into the taxonomy and divergence of Yanornithidae (Aves: Ornithothoraces). *Journal of Systematic Palaeontology*, 18(21):1805–1827.

Wang, M., O'Connor, J. K., Pan, Y., and Zhou, Z. (2017a). A bizarre Early Cretaceous enantiornithine bird with unique crural feathers and an ornithuromorph plough-shaped pygostyle. *Nature Communications*, 8(1):14141.

Wang, M., O'Connor, J. K., Zhou, S., and Zhou, Z. (2020b). New toothed Early Cretaceous ornithuromorph bird reveals intraclade diversity in pattern of tooth loss. *Journal of Systematic Palaeontology*, 18(8):631–645.

Wang, M., Zheng, X., O'Connor, J. K., Lloyd, G. T., Wang, X., Wang, Y., Zhang, X., and Zhou, Z. (2015a). The oldest record of ornithuromorpha [*sic*] from the early cretaceous [*sic*] of China. *Nature Communications*, 6(1):6987.

Wang, M. and Zhou, Z. (2018). A new confuciusornithid (Aves: Pygostylia) from the Early Cretaceous increases the morphological disparity of the Confuciusornithidae. *Zoological Journal of the Linnean Society*, 185(2):417–430.

Wang, M., Zhou, Z., O'Connor, J. K., and Zelenkov, N. V. (2014a). A new diverse enantiornithine family (Bohaiornithidae fam. nov.) from the Lower Cretaceous of China with information from two new species. *Vertebrata PalAsiatica*, 52(1):31–76.

Wang, M., Zhou, Z., and Zhou, S. (2016a). A new basal ornithuromorph bird (Aves: Ornithothoraces) from the Early Cretaceous of China with implication for morphology of early Ornithuromorpha. *Zoological Journal of the Linnean Society*, 176(1):207–223.

Wang, R., Wang, Y., and Hu, D. (2015b). Discovery of a new ornithuromorph genus, *Juehuaornis* gen. nov. from Lower Cretaceous of western Liaoning, China. *Global Geology*, 34(1):7–11. [in Chinese with English abstract].

Wang, X., Cau, A., Kundrát, M., Chiappe, L. M., Ji, Q., Wang, Y., Li, T., and Wu, W. (2020c). A new advanced ornithuromorph bird from Inner Mongolia documents the northernmost geographic distribution of the Jehol paleornithofauna in China. *Historical Biology*. doi:10.1080/08912963.2020.1731805.

Wang, X., Chiappe, L. M., Teng, F., and Ji, Q. (2013a). *Xinghaiornis lini* (Aves: Ornithothoraces) from the Early Cretaceous of Liaoning: an example of evolutionary mosaic in early birds. *Acta Geologica Sinica (English Edition)*, 87(3):686–689.

Wang, X., Huang, J., Hu, Y., Liu, X., Peteya, J., and Clarke, J. A. (2018). The earliest evidence for a supraorbital salt gland in dinosaurs in new Early Cretaceous ornithurines. *Scientific Reports*, 8(1):3969.

Wang, X., Huang, J., Kundrát, M., Cau, A., Liu, X., Wang, Y., and Ju, S. (2020d). A new jeholornithiform exhibits the earliest appearance of the fused sternum and pelvis in the evolution of avialan dinosaurs. *Journal of Asian Earth Sciences*, 199:104401.

Wang, X., Ji, Q., Teng, F., and Jin, K. (2013b). A new species of *Yanornis* (Aves: Ornithurae) from the Lower Cretaceous strata of Yixian, Liaoning Province. *Geological Bulletin of China*, 32(4):601–606.

Wang, X., Ju, S., and Wang, Y. (2019a). First zircon U-Pb ages of the Pigeon Hill fossil locality of the Jehol Biota in the Greater Khingan Mountains, Inner Mongolia. *Acta Geologica Sinica (English Edition)*, 93(4):1142–1145.

Wang, X., O'Connor, J. K., Zheng, X., Wang, M., Hu, H., and Zhou, Z. (2014b). Insights into the evolution of rachis dominated tail feathers from a new basal enantiornithine (Aves: Ornithothoraces). *Biological Journal of the Linnean Society*, 113(3):805–819.

Wang, Y., Hu, H., O'Connor, J. K., Wang, M., Xu, X., Zhou, Z., Wang, X., and Zheng, X. (2017b). A previously undescribed specimen reveals new information on the dentition of *Sapeornis chaoyangensis*. *Cretaceous Research*, 74:1–10.

Wang, Y., Olsen, P. E., Sha, J., Yao, X., Liao, H., Pan, Y., Kinney, S., Zhang, X., and Rao, X. (2016b). Stratigraphy, correlation, depositional environments, and cyclicity of the Early Cretaceous Yixian and ?Jurassic-Cretaceous Tuchengzi formations in the Sihetun area (NE China) based on three continuous cores. *Palaeogeography, Palaeoclimatology, Palaeoecology*, 464:110–133.

Wang, Y.-M., O'Connor, J. K., Li, D.-Q., and You, H.-L. (2013c). Previously unrecognized ornithuromorph bird diversity in the Early Cretaceous Changma Basin, Gansu Province, northwestern China. *PLOS ONE*, 8(10):e77693.

Wang, Y.-Q., Li, Q., Bai, B., Jin, X., Mao, F.-Y., and Meng, J. (2019b). Paleogene integrative stratigraphy and timescale of China. *Science China Earth Sciences*, 62(1):287–309.

Wang, Y.-Q., Meng, J., Beard, C. K., Li, Q., Ni, X.-J., Gebo, D. L., Bai, B., Jin, X., and Li, P. (2010). Early Paleogene stratigraphic sequences, mammalian evolution and its response to environmental changes in Erlian Basin, Inner Mongolia, China. *Science China Earth Sciences*, 53(12):1918–1926.

Ward, L. W. and Andrews, G. W. (2008). Stratigraphy of the Calvert, Choptank, and St. Marys Formations (Miocene) in the Chesapeake Bay area, Maryland and Virginia. *Virginia Museum of Natural History Memoir*, 9:1–170.

Watanabe, J., Matsuoka, H., and Hasegawa, Y. (2016). Two species of *Uria* (Aves: Alcidae) from the Pleistocene of Shiriya, northeast Japan, with description and body mass estimation of a new species. *Bulletin of the Gunma Museum of Natural History*, 20:59–72.

Weems, R. E., Edwards, L. E., and Landacre, B. (2017). Geology and biostratigraphy of the Potomac River cliffs at Stratford Hall, Westmoreland County, Virginia. In Bailey, C. M. and Jaye, S., editors, *From the Blue Ridge to the Beach: Geological Field Excursions across Virginia*, pages 125–152. Geological Society of America, Boulder, CO.

Weems, R. E. and George, R. A. (2013). Amphibians and nonmarine turtles from the Miocene Calvert Formation of Delaware, Maryland, and Virginia (USA). *Journal of Paleontology*, 87(4):570–588.

Weishampel, D. B., Fastovsky, D. E., Watabe, M., Varricchio, D., Jackson, F., Tsogtbaatar, K., and Barsbold, R. (2008). New oviraptorid embryos from Bugin-Tsav, Nemegt Formation (Upper Cretaceous), Mongolia, with insights into their habitat and growth. *Journal of Vertebrate Paleontology*, 28(4):1110–1119.

West, A. R., Torres, C. R., Case, J. A., Clarke, J. A., O'Connor, P. M., and Lamanna, M. C. (2019). An avian femur from the Late Cretaceous of Vega Island, Antarctic Peninsula: removing the record of cursorial landbirds from the Mesozoic of Antarctica. *PeerJ*, 7:e7231.

Westerhold, T., Röhl, U., Wilkens, R. H., Gingerich, P. D., Clyde, W. C., Wing, S. L., Bowen, G. J., and Kraus, M. J. (2018). Synchronizing early Eocene deep-sea and continental records – cyclostratigraphic age models for the Bighorn Basin Coring Project drill cores. *Climate of the Past*, 14(3):303–319.

Westgate, J. W. (2012). Palaeoecology of a primate-friendly, middle Eocene community from Laredo, Texas and a review of stratigraphic occurrences of Paleogene land mammals across the Gulf Coastal Plain, USA. *Palaeobiodiversity and Palaeoenvironments*, 92(4):497–505.

Wiest, L. A., Buynevich, I. V., Grandstaff, D. E., Terry, Dennis O., J., Maza, Z. A., and Lacovara, K. J. (2016). Ichnological evidence for endobenthic response to the K–Pg event, New Jersey, U.S.A. *Palaios*, 31(5):231–241.

Wijnker, E. and Olson, S. L. (2009). A revision of the fossil genus *Miocepphus* and other Miocene Alcidae (Aves: Charadriiformes) of the Western North Atlantic Ocean. *Journal of Systematic Palaeontology*, 7(4):471–487.

Wing, S. L., Bao, H., and Koch, P. L. (1999). An early Eocene cool period? Evidence for continental cooling during the warmest part of the Cenozoic. In Huber, B. T., Macleod, K. G., and Wing, S. L., editors, *Warm Climates in Earth History*, page 197–238. Cambridge University Press, Cambridge, UK.

Woodburne, M. O., MacFadden, B. J., Case, J. A., Springer, M. S., Pledge, N. S., Power, J. D., Woodburne, J. M., and Springer, K. B. (1994). Land mammal biostratigraphy and magnetostratigraphy of the Etadunna Formation (late Oligocene) of South Australia. *Journal of Vertebrate Paleontology*, 13(4):483–515.

Worthy, T. H., Degrange, F. J., Handley, W. D., and Lee, M. S. Y. (2017). The evolution of giant flightless birds and novel phylogenetic relationships for extinct fowl (Aves, Galloanseres). *Royal Society Open Science*, 4(10):170975.

Xue, X. (1995). *Qinornis paleocenica*—a Paleocene bird discovered in China. *Courier Forschungsinstitut Senckenberg*, 181:89–93.

Yang, W., Li, S., and Jiang, B. (2007). New evidence for cretaceous age of the feathered dinosaurs of Liaoning: zircon U–Pb SHRIMP dating of the Yixian Formation in Sihetun, northeast China. *Cretaceous Research*, 28(2):177–182.

Yang, Z. (2020). User Guide. PAML: Phylogenetic Analysis by Maximum Likelihood. Version 4.9j (February 2020). Available from `http://abacus.gene.ucl.ac.uk/software/pamlDOC.pdf`.

Yao, X., Liao, C.-C., Sullivan, C., and Xu, X. (2019). A new transitional therizinosaurian theropod from the Early Cretaceous Jehol Biota of China. *Scientific Reports*, 9(1):5026.

Yonezawa, T., Segawa, T., Mori, H., Campos, P. F., Hongoh, Y., Endo, H., Akiyoshi, A., Kohno, N., Nishida, S., Wu, J., Jin, H., Adachi, J., Kishino, H., Kurokawa, K., Nogi, Y., Tanabe, H., Mukoyama, H., Yoshida, K., Rasoamiaramanana, A., Yamagishi, S., Hayashi, Y., Yoshida, A., Koike, H., Akishinonomiya, F., Willerslev, E., and Hasegawa, M. (2017). Phylogenomics and morphology of extinct paleognaths reveal the origin and evolution of the ratites. *Current Biology*, 27(1):68–77.

You, H.-l., Lamanna, M. C., Harris, J. D., Chiappe, L. M., O'Connor, J., Ji, S.-a., Lü, J.-c., Yuan, C.-x., Li, D.-q., Zhang, X., Lacovara, K. J., Dodson, P., and Ji, Q. (2006). A nearly modern amphibious bird from the Early Cretaceous of northwestern China. *Science*, 312(5780):1640–1643.

Yuri, T., Kimball, R. T., Harshman, J., Bowie, R. C. K., Braun, M. J., Chojnowski, J. L., Han, K.-L., Hackett, S. J., Huddleston, C. J., Moore, W. S., Reddy, S., Sheldon, F. H., Steadman, D. W., Witt, C. C., and Braun, E. L. (2013). Parsimony and model-based analyses of indels in avian nuclear genes reveal congruent and incongruent phylogenetic signals. *Biology*, 2(1):419–444.

Zanno, L. E. and Makovicky, P. J. (2013). No evidence for directional evolution of body mass in herbivorous theropod dinosaurs. *Proceedings of the Royal Society B*, 280(1751):20122526.

Zeebe, R. E. and Lourens, L. J. (2019). Solar System chaos and the Paleocene–Eocene boundary age constrained by geology and astronomy. *Science*, 365(6456):926–929.

Zelenkov, N. V. (2018). A swan-sized anseriform bird from the late Paleocene of Mongolia. *Journal of Vertebrate Paleontology*, 38(6):e1531879.

Zelenkov, N. V. and Kurochkin, E. N. (2015). Класс Aves [Class Aves]. In Kurochkin, E. N., Lopatin, A. V., and Zelenkov, N. V., editors, *Ископаемые позвоночные России и сопредельных стран: Справочник для палеонтологов, биологов и геологов. Ископаемые рептилии и птицы, Часть 3 [Fossil Vertebrates of Russia and Adjacent Countries: The Reference Book for Paleontologists, Biologists and Geologists. Fossil Reptiles and Birds, Part 3]*, pages 86–290. GEOS, Moscow, Russia. [in Russian].

Zelenkov, N. V. and Panteleyev, A. V. (2015). Three bird taxa (Aves: Anatidae, Phasianidae, Scolopacidae) from the Late Miocene of the Sea of Azov (Southwestern Russia). *Paläontologische Zeitschrift*, 89(3):515–527.

Zelenkov, N. V., Panteleyev, A. V., and De Pietri, V. L. (2017). Late Miocene rails (Aves: Rallidae) from southwestern Russia. *Palaeobiodiversity and Palaeoenvironments*, 97(4):791–805.

Zelenkov, N. V., Volkova, N. V., and Gorobets, L. V. (2016). Late Miocene buttonquails (Charadriiformes, Turnicidae) from the temperate zone of Eurasia. *Journal of Ornithology*, 157(1):85–92.

Zhang, F., Zhou, Z., and Benton, M. J. (2008). A primitive confuciusornithid bird from China and its implications for early avian flight. *Science in China Series D: Earth Sciences*, 51(5):625–639.

Zhang, H., Liu, X., Yuan, H., Hu, Z., and Diqu, C. (2006). U-Pb isotopic age of the lower Yixian Formation in Lingyuan of western Liaoning and its significance. *Geological Review*, 52(1):63–71. [in Chinese with English abstract].

Zhang, L., Yang, Y., Zhang, L., Guo, S., Wang, W., and Zheng, S. (2007). Precious fossil-bearing beds of the Lower Cretaceous Jiufotang Formation in western Liaoning Province, China. *Acta Geologica Sinica (English Edition)*, 81(3):357–364.

Zheng, X., O'Connor, J. K., Wang, X., Wang, Y., and Zhou, Z. (2018). Reinterpretation of a previously described Jehol bird clarifies early trophic evolution in the Ornithuromorpha. *Proceedings of the Royal Society B*, 285(1871):20172494.

Zheng, X., O'Connor, J. K., Wang, X., Zhang, X., and Wang, Y. (2014). New information on Hongshanornithidae (Aves: Ornithuromorpha) from a new subadult specimen. *Vertebrata PalAsiatica*, 52(2):217–232.

Zhong, Y., Huyskens, M. H., Yin, Q.-Z., Wang, Y., Ma, Q., and Xu, Y.-G. (2021). High-precision geochronological constraints on the duration of 'Dinosaur Pompeii' and the Yixian Formation. *National Science Review*, 8(6):nwab063.

Zhou, S., O'Connor, J. K., and Wang, M. (2014a). A new species from an ornithuromorph (Aves: Ornithothoraces) dominated locality of the Jehol Biota. *Chinese Science Bulletin*, 59(36):5366–5378.

Zhou, S., Zhou, Z., and O'Connor, J. K. (2012). A new basal beaked ornithurine bird from the Lower Cretaceous of western Liaoning, China. *Vertebrata PalAsiatica*, 50(1):9–24.

Zhou, S., Zhou, Z., and O'Connor, J. K. (2013). Anatomy of the basal ornithuromorph bird *Archaeorhynchus spathula* from the Early Cretaceous of Liaoning, China. *Journal of Vertebrate Paleontology*, 33(1):141–152.

Zhou, S., Zhou, Z., and O'Connor, J. K. (2014b). A new piscivorous ornithuromorph from the Jehol Biota. *Historical Biology*, 26(5):608–618.

Zhou, Z., Barrett, P. M., and Hilton, J. (2003). An exceptionally preserved Lower Cretaceous ecosystem. *Nature*, 421(6925):807–814.

Zhou, Z., Fucheng, Z., and Li, Z. (2009). A new basal ornithurine bird (*Jianchangornis microdonta* gen. et sp. nov.) from the Lower Cretaceous of China. *Vertebrata PalAsiatica*, 47(4):299–310.

Zhou, Z., Fucheng, Z., and Li, Z. (2010). A new Lower Cretaceous bird from China and tooth reduction in early avian evolution. *Proceedings of the Royal Society B*, 277(1679):219–227.

Zhou, Z. and Zhang, F. (2001). Two new ornithurine birds from the Early Cretaceous of western Liaoning, China. *Chinese Science Bulletin*, 46(15):1258–1264.

Zhou, Z. and Zhang, F. (2002). Largest bird from the Early Cretaceous and its implications for the earliest avian ecological diversification. *Naturwissenschaften*, 89(1):34–38.

Zhou, Z. and Zhang, F. (2005). Discovery of an ornithurine bird and its implication for Early Cretaceous avian radiation. *Proceedings of the National Academy of Sciences*, 102(52):18998–19002.

Zhou, Z. and Zhang, F. (2006). A beaked basal ornithurine bird (Aves, Ornithurae) from the Lower Cretaceous of China. *Zoologica Scripta*, 35(4):363–373.

Zhou, Z. and Zhang, F. (2007). Mesozoic birds of China—a synoptic review. *Frontiers of Biology in China*, 2(1):1–14.

Zhu, R., Pan, Y., Shi, R., Liu, Q., and Li, D. (2007). Palaeomagnetic and $^{40}Ar/^{39}Ar$ dating constraints on the age of the Jehol Biota and the duration of deposition of the Sihetun fossil-bearing lake sediments, northeast China. *Cretaceous Research*, 28(2):171–176.

Zusi, R. L. (1984). A functional and evolutionary analysis of rhynchokinesis in birds. *Smithsonian Contributions to Zoology*, 395:1–40.
